# Mild/Asymptomatic Maternal SARS-CoV-2 Infection Leads to Immune Paralysis in Fetal Circulation and Immune Dysregulation in Fetal-Placental Tissues

**DOI:** 10.1101/2023.05.10.540233

**Authors:** Brianna M. Doratt, Suhas Sureshchandra, Heather True, Monica Rincon, Nicole Marshall, Ilhem Messaoudi

**Affiliations:** Department of Microbiology, Immunology and Molecular Genetics, University of Kentucky, Lexington KY 40536; Department of Physiology and Biophysics, School of Medicine, University of California, Irvine CA 92697; Institute for Immunology, University of California, Irvine CA 92697; Department of Pharmaceutical Sciences, University of Kentucky, Lexington KY 40536; Maternal-Fetal Medicine, Oregon Health and Science University, Portland OR 97239

**Keywords:** COVID-19, SARS-CoV-2, placenta, chorionic villi, umbilical cord blood, Hofbauer cells

## Abstract

Few studies have addressed the impact of maternal mild/asymptomatic SARS-CoV-2 infection on the developing neonatal immune system. In this study, we analyzed umbilical cord blood and placental chorionic villi from newborns of unvaccinated mothers with mild/asymptomatic SARS-CoV-2 infection during pregnancy using flow cytometry, single-cell transcriptomics, and functional assays. Despite the lack of vertical transmission, levels of inflammatory mediators were altered in cord blood. Maternal infection was also associated with increased memory T, B cells, and non-classical monocytes as well as increased activation. However, *ex vivo* responses to stimulation were attenuated. Finally, within the placental villi, we report an expansion of fetal Hofbauer cells and infiltrating maternal macrophages and rewiring towards a heightened inflammatory state. In contrast to cord blood monocytes, placental myeloid cells were primed for heightened antiviral responses. Taken together, this study highlights dysregulated fetal immune cell responses in response to mild maternal SARS-CoV-2 infection during pregnancy.

## INTRODUCTION

To date, over 255,000 pregnant women in the United States have been infected with severe acute respiratory syndrome coronavirus 2 (SARS-CoV-2)^1^. Although most pregnant women experience asymptomatic or mild coronavirus disease 2019 (COVID-19), those who experience severe disease are at a significantly higher risk for admission to the intensive care unit, mechanical ventilation, and pre-term birth ^2–4^. While some studies have described the presence of SARS-CoV-2 RNA in placental villi, including maternal macrophages and Hofbauer cells (HBC) ^5–7^, vertical transmission is extremely rare ^8–11^. Nevertheless, SARS-CoV-2 infection during pregnancy has been shown to alter frequencies of macrophage and effector T-cell subsets and induce a pro-inflammatory environment at the maternal-fetal interface (MFI), specifically within the maternal decidua ^12–14^. Moreover, pregnant women with severe COVID-19 are more likely to give birth to newborns with morbidities including respiratory distress syndrome ^15, 16^, hyperbilirubinemia ^16–18^, sepsis ^16, 19–21^, infections requiring antibiotic treatments ^22^, and admission to the neonatal intensive care unit (NICU) ^16, 20, 23^.

Mechanistic underpinnings explaining these adverse outcomes are only beginning to emerge. Recent studies have reported significantly elevated NK cell frequencies in UCB from neonates born to pregnant women who have recovered from SARS-CoV-2 infection compared to those born to mothers with ongoing infection at delivery ^24^. Furthermore, umbilical cord blood (UCB) NK cells from neonates born to mothers with active SARS-CoV-2 infection or those who recovered display increased expression of DNAX accessory molecule 1 (DNAM-1) ^24^, an NK cell-activating receptor essential for the recognition and killing of virus-infected cells ^25^. Asymptomatic or mild maternal SARS-CoV-2 infection detected at delivery also results in an altered inflammatory milieu in fetal circulation, including increased UCB plasma levels of IL-1β, IL-6, IL-8, IL-18, IL-33, IFNγ, caspase 1, nuclear factor of activated T cells (NFATC3), and CCL21 ^26–28^. Additionally, T_H_2 responses are dampened in infants born to mothers with infection in the second and third trimesters ^28^. Cases of very high anti-SARS-CoV-2 IgG concentrations (>5871.07 U/mL) detected in UCB were associated with higher frequencies of fetal neutrophils and cytotoxic T cells ^27^.

Bulk RNA sequencing of UCB revealed that mild/asymptomatic maternal SARS-CoV-2 infection in the third trimester is associated with the upregulation of genes responsible for antimicrobial responses and down-regulation of genes enriched for phagocytosis, complement activation, and extracellular matrix organization ^26^. Additionally, UCB monocytes exhibited upregulation of IFN-stimulated genes (ISG) and MHC class I and II genes ^26^. Single-cell analysis of UCB from newborns of mothers with mild COVID-19 in the third trimester revealed transcriptional changes that correlated with activation of plasmacytoid dendritic cells (pDCs), activation and exhaustion of NK cells, and clonal expansion of fetal T cells ^29^. While there are clear disruptions in the UCB immune landscape with maternal SARS-CoV-2 infection, the functional implications of these changes remain largely unknown.

Our previous studies have shown extensive remodeling of decidua (maternal placental compartment) obtained from pregnant women with mild/asymptomatic SARS-CoV-2 infection ^12^, including altered frequencies of decidual macrophages, regulatory T cells (Tregs), and activated T cells. Furthermore, antigen presentation and type I IFN signaling were attenuated in decidual macrophages, while pathways associated with cytokine signaling and cell killing were upregulated in decidual T cells. While abnormal placental pathologies have been reported with maternal SARS-CoV-2 infection, including inflammation and necrosis ^30–32^, few studies have addressed how maternal SARS-CoV-2 impacts the immune landscape of villous tissues (fetal placental compartment) ^26, 33–35^. Placental SARS-CoV-2 infection is associated with the recruitment of maternal monocytes and macrophages to villous tissues and increased frequency of fetal HBCs that express PD-L1, a possible mechanism to prevent immune cell-driven placental damage ^36^. Finally, a recent study reported a significant downregulation of genes responsible for Type 1 interferon and IL-6/IL-1β cytokine responses in the chorionic villous regardless of maternal COVID-19 severity, the gestational timing of infection, gestational age at delivery, pre-pregnancy BMI, or mode of delivery (cesarean versus vaginal delivery) ^34^.

Despite these observations, our understanding of the impact of asymptomatic/mild maternal SARS-CoV-2 infection on the immune landscape of fetal placental tissues and circulation remains incomplete due to a lack of studies that examined paired samples and where transcriptional analyses were coupled with functional assays. In this study, we used a combination of single-cell RNA sequencing and functional assays to address this gap in knowledge. Our data show that mild/asymptomatic maternal SARS-CoV-2 infection leads to heightened basal activation but dysfunctional responses of both innate and adaptive branches in circulation. This dysregulation extends to the fetal placental compartment (chorionic villi), as shown by the increased infiltration of regulatory maternal monocytes/macrophages to the fetal compartment, HBC activation, and impaired responses of villous myeloid cells to antimicrobial stimulation.

## METHODS

### Cohort characteristics

This study was approved by the Institutional Ethics Review Boards of Oregon Health & Science University and the University of Kentucky. Placental chorionic villi and UCB samples from 41 healthy, pregnant participants without SARS-CoV-2 infection or vaccination who had an uncomplicated, singleton pregnancy and 18 pregnant participants with asymptomatic (n=8) or mild (n=10) SARS-CoV-2 infection, but otherwise healthy pregnancies, were collected. Participants were classified as having mild SARS-CoV-2 infection if they experienced mild respiratory symptoms accompanied by a positive COVID test, while participants were classified as experiencing an asymptomatic infection if they tested positive during the mandatory COVID testing upon admission to labor and delivery and reported no symptoms. Importantly, all nasal swabs from newborns of SARS-CoV-2 infected participants as well as placental chorionic villi tissue samples tested negative for SARS-CoV-2 by qPCR. Controls were participants who did not experience COVID symptoms or report a positive COVID test at any time during their pregnancy receiving care at the same facility. The characteristics of the cohort are outlined in Table 1.

**Table 1:**
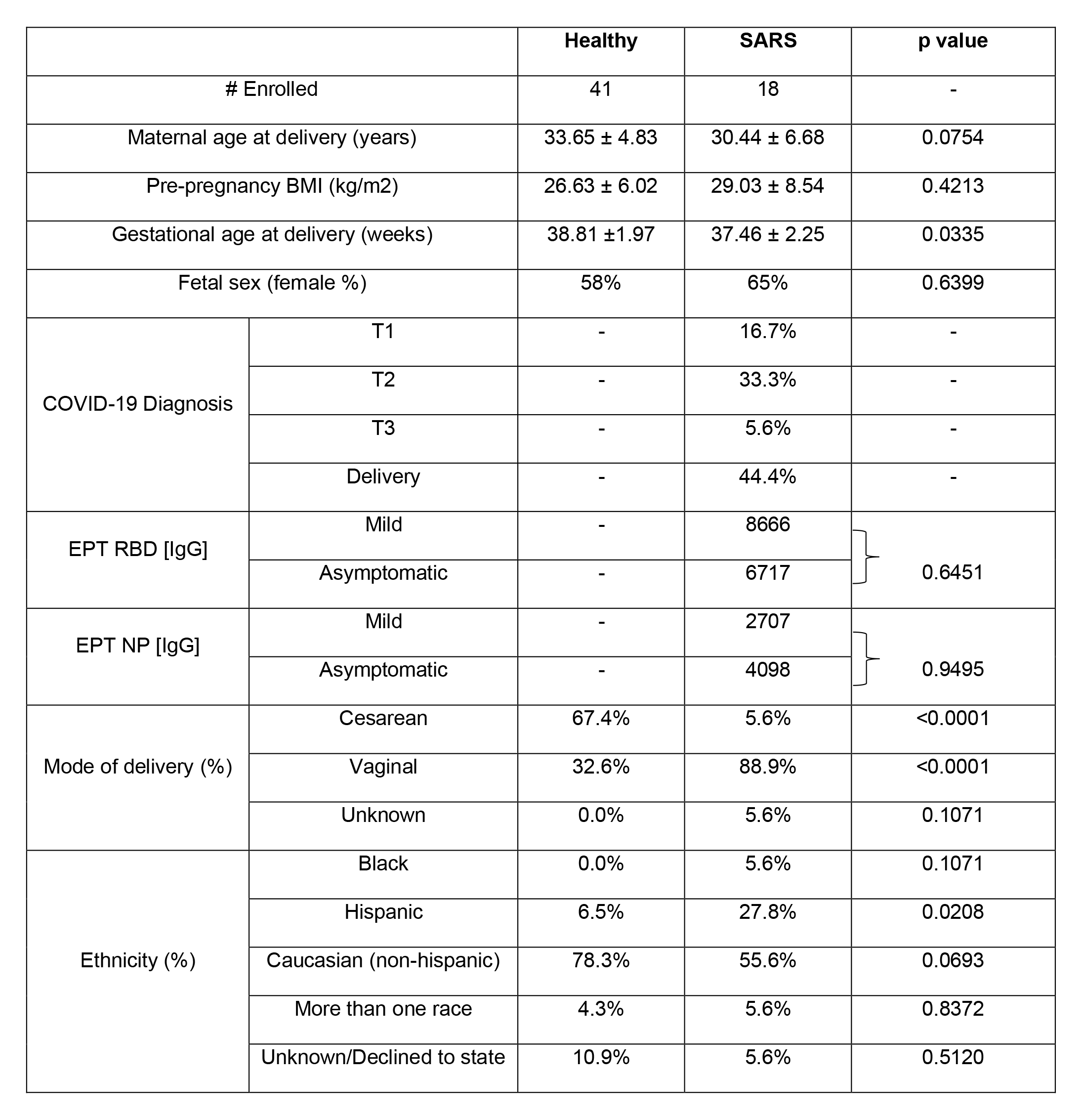
Description of cohort characteristics.

|  |  | Healthy | SARS | p value |
| --- | --- | --- | --- | --- |
| # Enrolled |  | 41 | 18 | - |
| Maternal age at delivery (years) |  | 33.65 ± 4.83 | 30.44 ± 6.68 | 0.0754 |
| Pre-pregnancy BMI (kg/m2) |  | 26.63 ± 6.02 | 29.03 ± 8.54 | 0.4213 |
| Gestational age at delivery (weeks) |  | 38.81 ± 1.97 | 37.46 ± 2.25 | 0.0335 |
| Fetal sex (female %) |  | 58% | 65% | 0.6399 |
| COVID-19 Diagnosis | T1 | - | 16.7% | - |
|  | T2 | - | 33.3% | - |
|  | T3 | - | 5.6% | - |
|  | Delivery | - | 44.4% | - |
| EPT RBD [IgG] | Mild | - | 8666 | } 0.6451 |
|  | Asymptomatic | - | 6717 |  |
| EPT NP [IgG] | Mild | - | 2707 | } 0.9495 |
|  | Asymptomatic | - | 4098 |  |
| Mode of delivery (%) | Cesarean | 67.4% | 5.6% | <0.0001 |
|  | Vaginal | 32.6% | 88.9% | <0.0001 |
|  | Unknown | 0.0% | 5.6% | 0.1071 |
| Ethnicity (%) | Black | 0.0% | 5.6% | 0.1071 |
|  | Hispanic | 6.5% | 27.8% | 0.0208 |
|  | Caucasian (non-hispanic) | 78.3% | 55.6% | 0.0693 |
|  | More than one race | 4.3% | 5.6% | 0.8372 |
|  | Unknown/Declined to state | 10.9% | 5.6% | 0.5120 |

### Blood processing

Whole blood samples were collected in EDTA vacutainer tubes. Complete blood counts were obtained by a Cell-Dyn Emerald 22 (Abbott, Abbott Park, Illinois). UCB mononuclear cells (UCBMC) and plasma were isolated after whole-blood centrifugation over LymphoPrep in SepMate tubes (STEMCELL Technologies, Vancouver, BC) following manufacturers’ protocols. Plasma was stored at –80°C until analysis. UCBMC were cryopreserved using 10% DMSO/FBS and Mr. Frosty Nalgene Freezing containers (Thermo Fisher Scientific, Waltham, Massachusetts) at –80°C overnight and then transferred to a cryogenic unit until analysis.

### Placenta processing

Fetal chorionic villi were separated from maternal decidua and immediately immersed in RPMI supplemented with 10% FBS, 1% penicillin-streptomycin, and 1% L-glutamine (GeminiBio, Sacramento, California). Samples were processed within 24 hours of collection. Chorionic villi were first washed thoroughly in HBSS to remove contaminating blood, then minced into approximately 0.2-0.3mm^3^ cubes, followed by enzymatic digestion at 37°C for 1 hour in R3 media (RPMI 1640 with 3% FBS, 1% Penicillin-Streptomycin, 1% L-glutamine, and 1M HEPES) supplemented with 0.5 mg/mL collagenase IV (Sigma-Aldrich, Saint Louis, Missouri). The disaggregated cell suspension was passed through tissue strainers to eliminate large tissue chunks. Cells were pelleted and passed sequentially through 100-, 70-, and 40-µm cell sieves. Red blood cells were lysed using RBC lysis buffer (155 mM NH4Cl, 12 mM NaHCO3, 0.1 mM EDTA in double distilled water). The cell suspension was then layered on discontinuous 60% and 40% percoll gradients (Sigma-Aldrich, Saint Louis, Missouri) and centrifuged for 30 minutes at 930xg with the brakes off. Immune cells at the interface of 40% and 60% gradients were collected, counted, and cryopreserved as described above for UCBMC for future analysis. SARS-CoV-2 viral loads were assessed in placental tissues using qPCR as previously described ^12^.

### ELISA

End-point titers (EPT) against the SARS-CoV-2 receptor binding domain of the spike protein (RBD) and nucleocapsid protein (NP) were determined using standard ELISA as recently described ^37^. Plates were coated with 500 ng/mL RBD or 1 μg/mL NP (GenScript, Piscataway, New Jersey), and heat-inactivated plasma (1:50 in blocking buffer) was added in 3-fold dilutions. Responses were visualized by adding HRP anti-human IgG (BD Pharmingen, San Diego, California) followed by o-Phenylenediamine dihydrochloride (Thermo Fisher Scientific, Waltham, Massachusetts). Batch differences were minimized by normalizing to a positive control sample run on each plate. EPTs were calculated using log-log transformation of the linear portion of the curve and 0.1 OD units as the cut-off.

### Plasma Luminex

Levels of immune mediators in plasma, cell culture supernatant following RSV or *E. coli* stimulation, and resting HBC cell culture supernatant were measured using a human, premixed 45-plex panel (R&D Systems, Inc. Minneapolis, Minnesota). Immune mediators in cell culture supernatant following CD3/CD28 bead stimulation were measured using a human premixed CD8+ T Cell Human 17-plex panel (Millipore, Temecula, California). All Luminex assays were analyzed using a MAGPIX Instrument and xPONENT software (Luminex, Austin, Texas).

### Phenotyping

1-2 x 10^6^ UCBMC were stained using antibodies against CD4, CD8b, CCR7, CD45RA, CD19, CD27, IgD, and KLRG1 to delineate naïve and memory T and B cell populations ^38^. Cells were then fixed (Fixation buffer; BioLegend, San Diego, CA), permeabilized (Permeabilization wash buffer; BioLegend, San Diego, CA), and stained intracellularly for the proliferation marker Ki-67 (BioLegend, San Diego, CA). A second set of samples were stained using antibodies against CD3, CD20, HLA-DR, CD14, CD11c, CD123, CD56, and CD16 to delineate monocytes, myeloid dendritic cells (mDC); plasmacytoid dendritic cells (pDC) and natural killer (NK) cell subsets ^39, 40^. All flow cytometry samples were acquired with the Attune NxT instrument (ThermoFisher Scientific, Waltham, Massachusetts) and analyzed using FlowJo 10.5 (TreeStar, Ashland, Oregon).

Villous leukocytes were stained with CD45 (pan leukocyte marker), CD14, HLA-DR, FOLR2, CD9, and CCR2 to delineate HBCs (CD14^+^HLA-DR^-^FOLR2^+^CCR2^-^) and placenta associated maternal macrophages (PAMMs; PAMM1a: CD14^+^HLADR^+^FOLR2^-^CD9^+^CCR2^low/int^) and monocytes (PAMM1b: CD14^+^HLADR^+^FOLR2^-^CD9^-^/intCCR2^+^) and infiltrating maternal decidual macrophages (PAMM2: CD14^+^HLA-DR^hi^FOLR2^hi^) as previously described ^41^.

### Ex vivo cell stimulation

For T cell stimulations, 1×10^6^ UCBMC were cultured for 24 hours at 37C in RPMI supplemented with 10% FBS in the presence or absence of anti-CD3/CD28 beads (ThermoFisher Scientific, Waltham, Massachusetts). After 24 hours, the cells were spun down and the supernatants were collected for analysis by Human T Cell 17-plex panel (Sigma-Aldrich, Saint Louis, Missouri). For NK cell stimulation, 1×10^6^ UCBMC were stimulated for 6 hours at 37°C in RPMI supplemented with 10% FBS in the presence or absence of 0.5 μg/ml PMA and 5 μg/ml ionomycin (InvivoGen, San Diego, California). CD107a antibodies were added at the beginning of stimulation; Brefeldin A (BioLegend, San Diego, CA) was added after 1 hour incubation. Cells were stained for CD3, CD20, CD16, CD56, and HLA-DR, fixed, permeabilized, and stained intracellularly for IL-2, TNFα, MIP-1β, and IFNγ.

For monocyte/macrophages responses, CD14+ cells were FACS sorted from UCBMC or villous leukocytes and cultured for 16 h at 37°C in the absence/presence of either RSV (MOI 1) or E. coli (6×10^5^ cfu/well). Production of immune mediators in the supernatants was assessed using a human 45-plex (R&D Systems, Inc. Minneapolis, Minnesota).

### B cell purification and stimulation methods

B cells were purified from UCBMC using MACS CD20+ microbeads (Miltenyi Biotec, Waltham, MA). 50-100,000 B cells were plated per well and stimulated using a TLR agonist cocktail containing LPS (100 μg/μL), R848 (10 μg/mL), and ODN2216 (5μg/mL) in RP10 media. Control wells received RP10 + 0.4% DMSO. After stimulation for 24 hours, cells were surfaced stained with antibodies against CD3, CD20, HLA-DR, IgD, CD27, CD40, CD83, CD86, CD80, CD69, and IgG (BioLegend, San Diego, CA).

### 3’ multiplexed single-cell RNA sequencing

Freshly thawed UCBMCs (1-2×10^6^ cells) were stained with Ghost Violet 540 (Tonbo Biosciences, San Diego, CA) for 30 min at 4C in the dark before being incubated with Fc blocker (Human TruStain FcX, Biolegend, San Diego, California) in PBS with 1% BSA for 10 min at 4C. Cells were surface stained with CD45-FITC (HI30, BioLegend, San Diego, California) for 30 min at 4C in the dark. Samples were then washed twice in PBS with 0.04% BSA and incubated with individual CellPlex oligos (CMO) (10X Genomics, Pleasanton, California) per manufacturer’s instructions. Pellets were washed three times in PBS with 1% BSA, resuspended in 300 uL FACS buffer, and sorted on BD FACS Aria Fusion into RPMI (supplemented with 30% FBS). Sorted live CD45+ cells were counted in triplicates on a TC20 Automated Cell Counter (BioRad, Hercules, California), washed, and resuspended in PBS with 0.04% BSA in a final concentration of 1500 cells/µL. Single-cell suspensions were then immediately loaded on the 10X Genomics Chromium Controller with a loading target of 20,000 cells.

Freshly thawed villous leukocytes (1-2×10^6^ cells) were stained with Ghost Violet 540 (Tonbo Biosciences, San Diego, CA) for 30 min at 4C in the dark before being incubated with Fc blocker (Human TruStain FcX, BioLegend, San Diego, California) in PBS with 1% BSA for 10 min at 4C. Finally, cells were surface stained with HLA-DR, CD14, CCR2, and FOLR2 (BioLegend, San Diego, California) for 30 min at 4C in the dark. Samples were then washed twice and incubated with individual TotalSeq B antibodies (HTO) (BioLegend, San Diego, California) per manufacturer’s instructions. Pellets were resuspended in 300 uL FACS buffer and sorted on BD FACS Aria Fusion into RPMI (supplemented with 30% FBS). Sorted live CD14+CCR2+ and CD14+CCR2- cells were counted in triplicates on a TC20 Automated Cell Counter (BioRad, Hercules, California), washed, and resuspended in PBS with 0.04% BSA in a final concentration of 1500 cells/µL. Single-cell suspensions were then immediately loaded on the 10X Genomics Chromium Controller with a loading target of 20,000 cells.

All libraries were generated using the V3.1 chemistry for gene expression and Single Cell 3ʹ Feature Barcode Library Kit per the manufacturer’s instructions (10X Genomics, Pleasanton, California). Libraries were sequenced on Illumina NovaSeq 6000 with a sequencing target of 30,000 gene expression reads and 5,000 feature barcoding reads per cell.

### Single-cell RNA-Seq data analysis

Raw reads were aligned and quantified using Cell Ranger (version 6.0.2, 10X Genomics, Pleasanton, California) against the human reference genome (GRCh38) using the *multi-* option. Seurat (version 4.0) was used for downstream analysis. Cell doublets were removed by retaining droplets with a single CMO or HTO signal. Additionally, ambient RNA and dying cells were removed by filtering out droplets with less than 200 detected genes and greater than 20% mitochondrial gene expression, respectively. Data objects from controls and SARS+ groups were integrated using Seurat. Data normalization and variance stabilization were performed on the integrated object using the *NormalizeData* and *ScaleData* functions in Seurat, where a regularized negative binomial regression was corrected for differential effects of mitochondrial and ribosomal gene expression levels. Dimensionality reduction was performed using *RunPCA* function to obtain the first 30 principal components and clusters visualized using Seurat’s *RunUMAP* function. Cell types were assigned to individual clusters using *FindAllMarkers* function with a log2 fold change cutoff of at least 0.4, FDR<0.05, and using a known catalog of well-characterized scRNA markers for human PBMC and villous leukocytes (Supplemental Table 1) ^42^. Differential gene expression analysis was performed using MAST function in Seurat. Only statistically significant genes maintaining an FDR<0.05 and a log2 fold change ± 0.25 for UCBMC or 0.4 for villous leukocytes were included in downstream analyses. Module scores for specific pathways/gene sets were incorporated cluster-wise using the *AddModuleScores* function (Supplemental Table 2). Functional enrichment was performed using Metascape ^43^.

### Statistical Analyses

Data sets were first assessed for normality using Shapiro Wilk test and equality of variances using the Levene test. Group differences between datasets normally distributed were tested using an unpaired t-test (for datasets with equal variances) or an unpaired t-test with Welch’s correction (for cases with unequal variances). Datasets not normally distributed were subjected to non-parametric Mann-Whitney test. All statistical analyses were conducted in Prism version 9.4.1 (GraphPad)

## RESULTS

### Maternal SARS-CoV-2 infection leads to increased systemic fetal inflammation and frequency of myeloid cells

UCB and placental chorionic villous tissues were collected at delivery from newborns of mothers who tested positive for SARS-CoV-2 during pregnancy (mild) or at the time of delivery (asymptomatic) or had no COVID symptoms (controls) and receiving care at OHSU. Controls were mostly participants who delivered by scheduled cesarean due to challenges associated with recruitment during the pandemic, hence the higher number of cesarean sections in the control group (67.4%, p<0.0001) (Table 1). Cohort characteristics can be found in Table 1. Maternal age at delivery, pre-pregnancy BMI, and fetal sex were comparable between both groups (Table 1). Gestational age at delivery was significantly lower with maternal SARS-CoV-2 infection (p=0.0335), consistent with findings of increased rates of early labor in SARS-CoV-2 pregnancies ^16, 20, 23^. A greater proportion of pregnant participants with SARS-CoV-2 infection identified as Hispanic (27.8%, p=0.0208), in line with the increased incidence of SARS-CoV-2 in this population44.

All but one of the neonates of participants with SARS-CoV-2 infection had detectable IgG antibodies directed against spike protein receptor binding domain (RBD) at birth, albeit lower than maternal IgG titers (Figure 1A). Additionally, all but 2 dyads had detectable antibodies against nucleocapsid protein (NP), and maternal/neonatal titers were comparable between the two groups (Figure 1A). We observed no differences in antibody (IgG) titers against receptor binding domain (RBD) or nucleocapsid protein (NP) between participants in the mild and asymptomatic groups (Table 1). Therefore, all subsequent comparisons were performed on UCB and fetal placental tissues from newborns of SARS-CoV-2-positive (maternal SARS+) and SARS-CoV-2-naïve participants (control).

**Figure 1:**
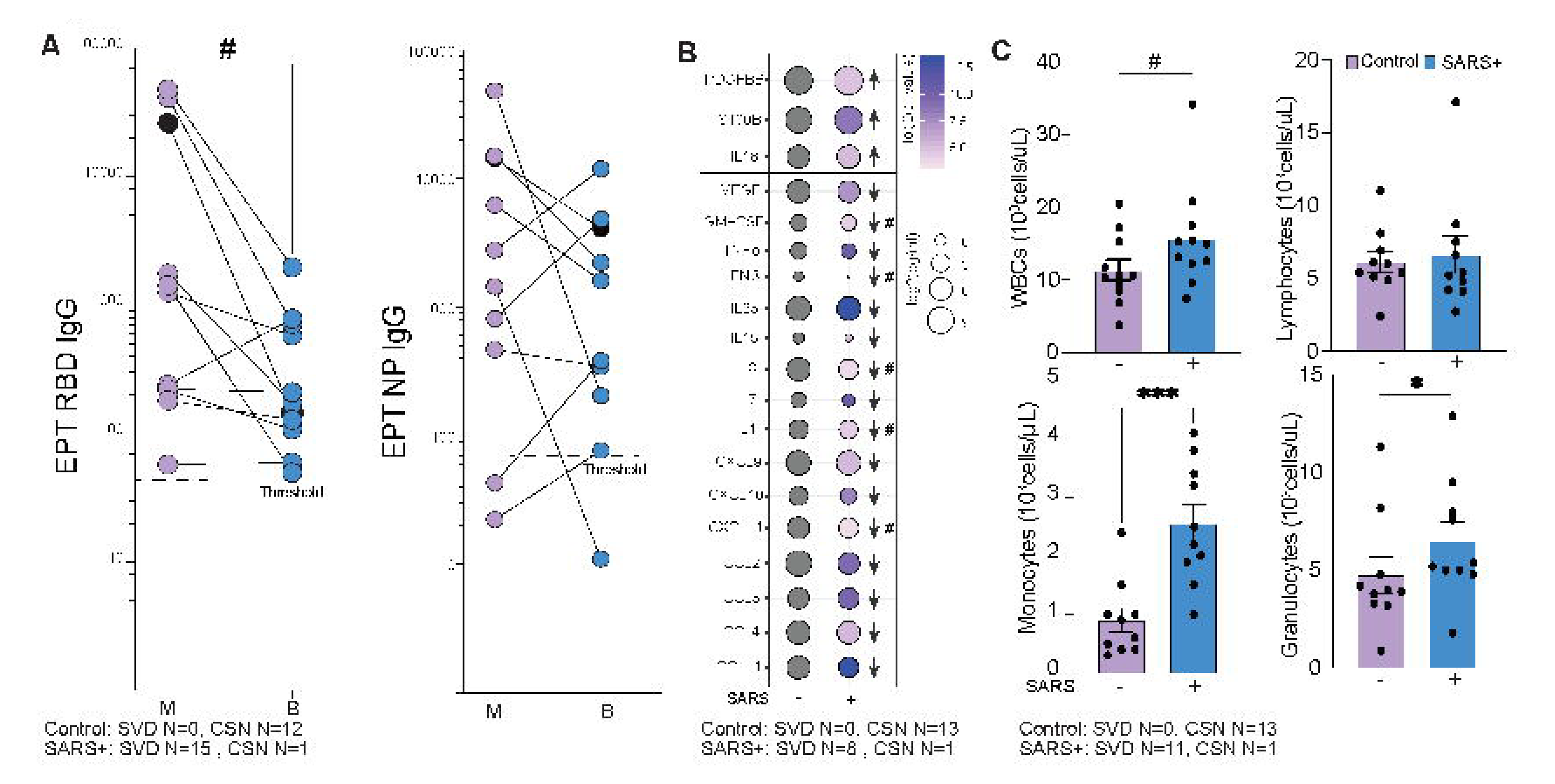
Maternal SARS Infection alters the frequency of circulating immune cells and immune mediators. (A) Maternal and umbilical cord blood (UCB) anti-RBD (left) and anti-NP (right) endpoint titers (EPT) from SARS-CoV-2 infected mothers (M) and their offspring (B). All samples are from mothers with a history of mild infection, except for the sample denoted by the black circle, which was asymptomatic and tested positive at delivery. (B) Bubble plot comparing UCB plasma immune mediators from control and maternal SARS+ group. Size represents analyte concentration (pg/mL), whereas color represents statistical significance. (C) UCB complete blood cell counts, including white blood cell (WBC) (top left), lymphocyte (top right), monocyte (bottom left), and granulocyte (bottom right) proportions from control and maternal SARS+ groups. SVD = standard vaginal delivery, CSN = cesarean section. (#=p<0.1, *=p<0.05, ***=p<0.0001).

Interestingly, maternal SARS-CoV-2 infection altered immune mediators in cord blood (Figure 1B). Specifically, concentrations of several chemokines important for the recruitment of both innate immune cells and lymphocytes (CXCL8, CXCL9, CXCL10, CCL4, CCL3, CXCL11, and CCL11) were lower in the maternal SARS+ group (Figure 1B). Moreover, levels of several antiviral and pro-inflammatory mediators, notably IFNβ, TNFα, IL-23 (Th17), and IL-15 (NK cell activation), were also lower. Levels of growth factor VEGF, anti-inflammatory regulator IL-1RA, and lymphocyte survival factor IL-7 were dampened in the maternal SARS+ group. In contrast, levels of S100B, a neurobiochemical marker for CNS injury, PDGF-BB, which regulates cell growth, and IL18 were increased (Figure 1B and Table 2). Complete blood cell counts from UCB of newborns in the maternal SARS+ group show increased numbers of total white blood cells driven by elevated monocyte and granulocyte numbers (Figure 1C).

**Table 2:**
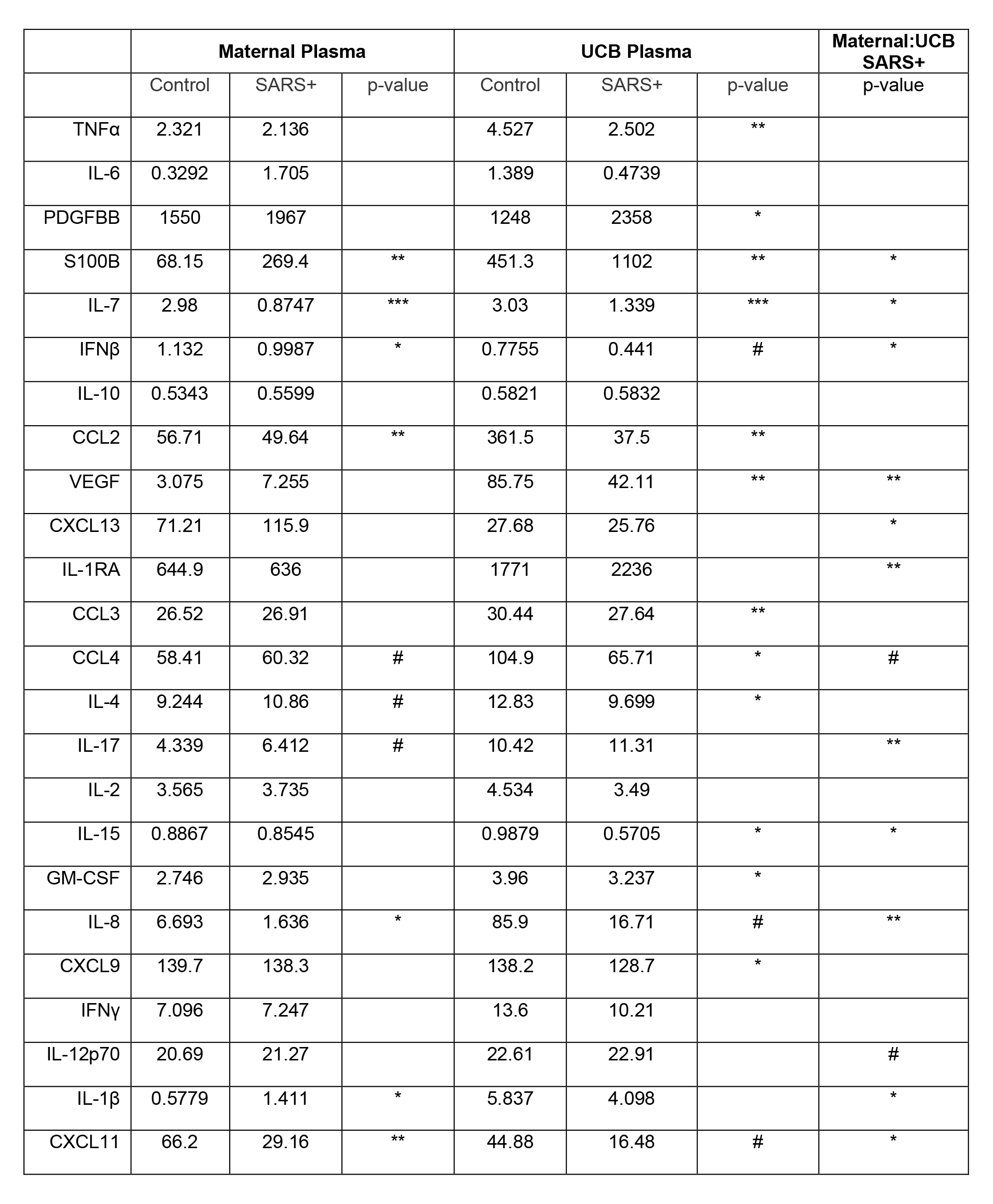
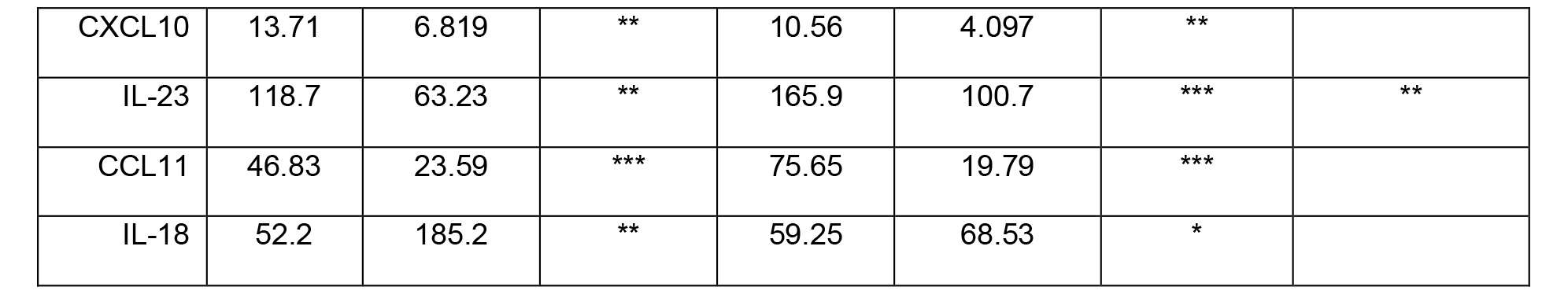
Maternal-fetal Luminex dyads. Concentration (pg/mL) of maternal and UCB plasma immune mediators from controls and maternal SARS+ groups. (#=p<0.1, *=p<0.05, **=p<0.01, ***=p<0.001, ****=p<0.0001)

|  | Maternal Plasma |  |  | UCB Plasma |  |  | Maternal:UCB<br>SARS+ |
| --- | --- | --- | --- | --- | --- | --- | --- |
|  | Control | SARS+ | p-value | Control | SARS+ | p-value | p-value |
| TNFα | 2.321 | 2.136 |  | 4.527 | 2.502 | ** |  |
| IL-6 | 0.3292 | 1.705 |  | 1.389 | 0.4739 |  |  |
| PDGFBB | 1550 | 1967 |  | 1248 | 2358 | * |  |
| S100B | 68.15 | 269.4 | ** | 451.3 | 1102 | ** | * |
| IL-7 | 2.98 | 0.8747 | *** | 3.03 | 1.339 | *** | * |
| IFNβ | 1.132 | 0.9987 | * | 0.7755 | 0.441 | # | * |
| IL-10 | 0.5343 | 0.5599 |  | 0.5821 | 0.5832 |  |  |
| CCL2 | 56.71 | 49.64 | ** | 361.5 | 37.5 | ** |  |
| VEGF | 3.075 | 7.255 |  | 85.75 | 42.11 | ** | ** |
| CXCL13 | 71.21 | 115.9 |  | 27.68 | 25.76 |  | * |
| IL-1RA | 644.9 | 636 |  | 1771 | 2236 |  | ** |
| CCL3 | 26.52 | 26.91 |  | 30.44 | 27.64 | ** |  |
| CCL4 | 58.41 | 60.32 | # | 104.9 | 65.71 | * | # |
| IL-4 | 9.244 | 10.86 | # | 12.83 | 9.699 | * |  |
| IL-17 | 4.339 | 6.412 | # | 10.42 | 11.31 |  | ** |
| IL-2 | 3.565 | 3.735 |  | 4.534 | 3.49 |  |  |
| IL-15 | 0.8867 | 0.8545 |  | 0.9879 | 0.5705 | * | * |
| GM-CSF | 2.746 | 2.935 |  | 3.96 | 3.237 | * |  |
| IL-8 | 6.693 | 1.636 | * | 85.9 | 16.71 | # | ** |
| CXCL9 | 139.7 | 138.3 |  | 138.2 | 128.7 | * |  |
| IFNγ | 7.096 | 7.247 |  | 13.6 | 10.21 |  |  |
| IL-12p70 | 20.69 | 21.27 |  | 22.61 | 22.91 |  | # |
| IL-1β | 0.5779 | 1.411 | * | 5.837 | 4.098 |  | * |
| CXCL11 | 66.2 | 29.16 | ** | 44.88 | 16.48 | # | * |
| CXCL10 | 13.71 | 6.819 | ** | 10.56 | 4.097 | ** |  |
| IL-23 | 118.7 | 63.23 | ** | 165.9 | 100.7 | *** | ** |
| CCL11 | 46.83 | 23.59 | *** | 75.65 | 19.79 | *** |  |
| IL-18 | 52.2 | 185.2 | ** | 59.25 | 68.53 | * |  |

### Maternal SARS-CoV-2 infection alters the frequency of circulating immune cells, suggestive of a heightened activation state

To uncover the changes within the fetal immune compartment in response to maternal SARS-CoV-2 infection, we performed single-cell RNA sequencing (scRNA-Seq) on UCBMC. We identified 16 unique immune cell clusters (Figure 2A and Supplemental Figure 1A) that were annotated using established gene markers for adult PBMC (Figure 2B and Supplemental Table 1). Within the lymphoid clusters, B cells were identified based on high expression of *MS4A1*, *CD79A*, and *IGHD*, while T cell subsets were defined based on the expression level of *CD3D*, *CD8B*, *IL7R*, and *CCR7* (Figure 2B). NK cell subsets were identified based on the high expression of *GZMA* and *NKG7* (Figure 2B). Monocyte clusters (classical, intermediate, and non-classical) were identified based on the expression of *CD14*, *HLA-DRA*, *S100A8*, *IL1B*, and *FCGR3*. Both subsets of DCs were identified – mDCs (expressing high *CD1C*) and pDCs (expressing high *IL3RA*) (Figure 2B). Additionally, stem cells (expressing *CD34*), proliferating cells (expressing *MKI67*), and a cluster of contaminating erythroid cells (expressing *HBB*) were identified (Figure 2B).

**Figure 2:**
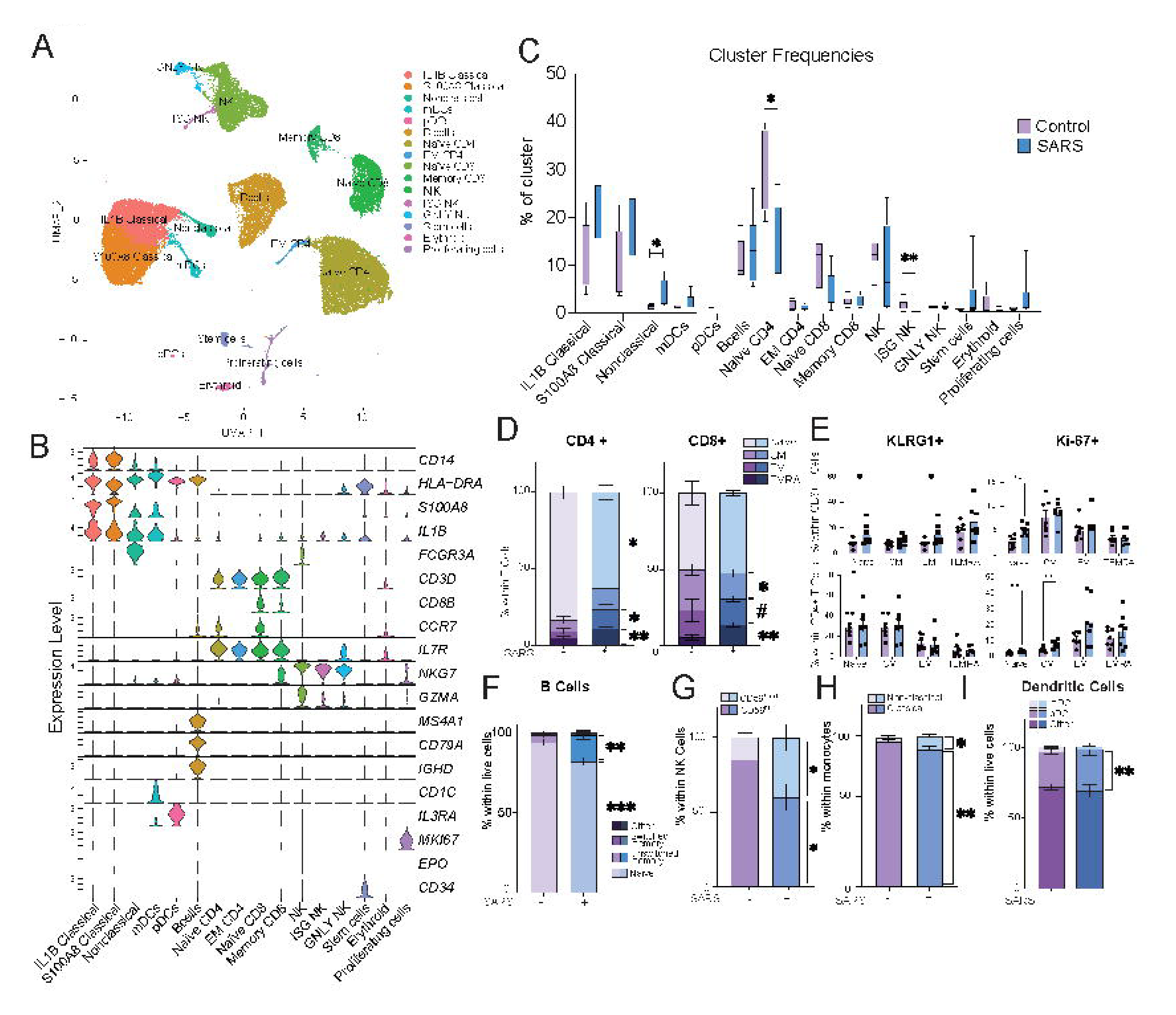
Impact of maternal asymptomatic/mild SARS-CoV-2 infection on phenotype and frequencies of cord blood immune cells. (A) Uniform manifold approximation and projection (UMAP) representation of 42,486 live immune cells from UCBMC of control and maternal SARS+ groups (N=4/group) showing 16 clusters. (B) Violin plots of marker genes used for cluster identification. (C) Box and whiskers plot comparing cluster frequencies in control and maternal SARS+ groups. (D) Stacked bar graphs of UCB CD4+ and CD8+ T cell subset frequencies in control and maternal SARS+ groups by flow cytometry. (E) Bar graphs comparing KLRG1 and Ki- 67 expression within CD4+ and CD8+ T cells between control and maternal SARS+ groups. (F-I) Stacked bar graphs comparing (F) B cell, (G) CD56bright/dim NK cell, (H) non-classical and classical monocyte, and (I) Dendritic cell subsets between control and maternal SARS+ group. (#=p<0.1, *=p<0.05, **=p<0.01, ***=p<0.001, ***=p<0.0001).

Despite the lack of differences in the total number of circulating lymphocytes (Figure 1C), maternal SARS-CoV-2 infection resulted in decreased frequencies of naïve CD4+ T cells and NK cells with high interferon signature (NK ISG) (Figure 2C). On the other hand, and in line with the increased numbers of circulating total monocytes measured by CBC (Figure 1C), the proportion of non-classical monocytes increased in the maternal SARS+ group (Figure 2C). We validated these observations using flow cytometry in a larger number of samples. This analysis confirmed the reduction of naïve CD4 T cells but also revealed a concomitant expansion of both effector and terminally differentiated effector memory (EM and TEMRA) CD4+ and CD8+ T cells (Figure 2D). Furthermore, expression of the activation marker KLRG1 was elevated in naïve and effector memory CD8+ T cells but not CD4+ T cells (Figure 2E), whereas expression of the proliferation marker Ki67 was increased in naïve CD4 and CD8 T cells as well as CM CD4 T cells in the maternal SARS+ group (Figure 2E). Similarly, a shift from naïve to unswitched memory B cell subsets was detected (Figure 2F). Finally, an expansion of immunoregulatory CD56^bright^ NK cells, non-classical monocytes, and pDCs (Figure 2G- I) were observed in the maternal SARS+ group.

### Maternal SARS-CoV-2 infection results in aberrant activation of fetal lymphocytes

Given the observed shift toward memory for T and B cells, we used the scRNA-Seq data to interrogate gene expression patterns associated with lymphocyte activation. Within B cells, SARS-CoV-2 infection was associated with increased scores of cytokine signaling and cell migration modules (Supplemental Figure 1B and Supplemental Table 2). Differentially expressed genes (DEG) with maternal SARS-CoV-2 infection mapped to the regulation of protein kinase activity and immunoglobulin receptor binding gene ontology (GO) terms (Supplemental Figure 1C) and include downregulated genes such as *FCRLA, MZB1, IGLC1/2/3, IGKC, and CD79B* (Supplemental Figure 1D). Given the downregulation of these key genes, we next tested the impact of maternal SARS-Cov-2 infection on functional B cell responses. Despite the increased frequency of memory subsets, B cells from the SARS+ group were less responsive to stimulation with TLR agonist cocktail ^45^, indicated by lack of induction of CD40 and dampened expression of HLA-DR and CD83 (Supplemental Figure 1E).

Within CD8 T cell clusters, there was an increase in transcriptional signatures of cell migration, cytotoxicity, and cytokine signaling with maternal infection (Figure 3A and Supplemental Table 2). DEGs within the memory CD8+ T cell compartment were in line with increased potential for cytotoxicity (*IL32*, *GZMK*, *KLRC2*, *KLRD1*, *NKG7*), inflammation (*S100A4*, *S100A9*, *S100A10*), survival/differentiation of activated lymphocytes (*CD8A, CD27, CD3E)*, and antiviral signaling (*IFITM1*) (Figure 3B). Within CD4+ T cell clusters, module scores associated with cell migration, cytokine signaling, Treg, and T_H_1 phenotype were increased in both naïve and EM subsets (Figure 3A and Supplemental Table 2). DEG analysis within naïve CD4 revealed increased transcript levels of genes associated with cell cycle (*CDK6*), consistent with elevated proliferation of naïve CD4 T cells in the maternal SARS+ group. On the other hand, EM CD4+ T cells had increased expression of genes associated with ATP synthesis and mitochondrial homeostasis (*ATP2B1, TSPO),* and T cell activation/signaling *(TNFRSF18, TRDC, TRAC, NFKBID)* (Figure 3B).

**Figure 3:**
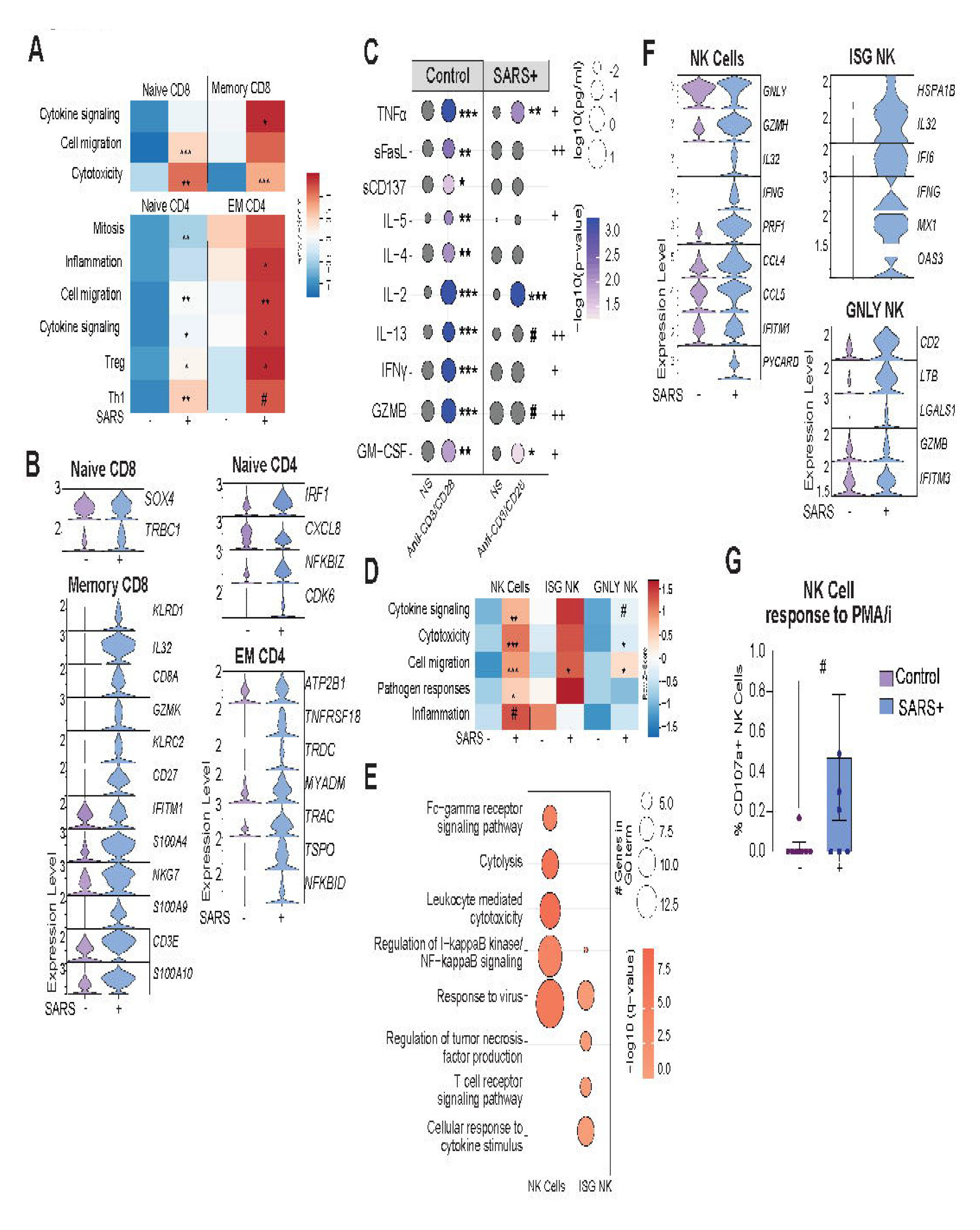
The impact of maternal SARS-CoV-2 infection on fetal lymphocytes and NK cells. (A) Heatmap of module scores within T cell clusters for the terms indicated. (B) Violin plot comparing normalized transcript counts of select statistically significant DEG within the indicated T cell cluster. (C) Bubble plot comparing secreted levels of immune mediators in cell culture supernatants following stimulation of UCBMC from control and maternal SARS+ groups with anti-CD3/CD28. The bubble size represents the analyte concentration (pg/mL), whereas the color represents the level of statistical significance compared to non-stimulated cells. Statistical significance between stimulated control and maternal SARS+ groups are indicated by plus signs (+=p<0.05, ++=p<0.01). (D) Heatmap of module scores within NK cell clusters for the terms indicated. (E) Bubble plot comparing functional enrichment of DEG relative to controls within ISG NK cell and NK cell clusters. The bubble size represents the number of genes mapping to the term, whereas the color represents the level of statistical significance. (F) Violin plot of select statistically significant DEG within the shown NK cell clusters. (G) Bar graph of total NK cell responses to PMA/ionomycin stimulation. (#=p<0.1, *=p<0.05, **=p<0.01, ***=p<0.001).

To interrogate the biological consequences of the changes in activation and transcriptional landscape, UCBMC from both groups were stimulated with anti-CD3/CD28 beads for 24 hours. T cells from controls generated a robust response as indicated by increased levels of canonical immune mediators (TNFα, sFASL, sCD137, IL-4, IL-5, IL-2, IL-13, IFNγ, GZMB, GM-CSF) (Figure 3C). On the other hand, T cells from the maternal SARS+ group responded poorly to polyclonal stimulation, indicated by the dampened secretion of both T_H_1 cytokines (IFNγ, GM-CSF) and T_Η_2 cytokines (IL-5, IL-13), and cytotoxic (GZMB) mediators (Figure 3C). These data suggest that heightened maternal inflammation consequent to SARS-CoV-2 infection reprograms neonatal lymphocytes leading to increased activation at baseline but their inability to respond to *ex vivo* stimulations.

### Maternal SARS-CoV-2 infection enhances fetal NK cell activation

As described for T cells, maternal SARS-CoV-2 infection was associated with increased scores of modules associated with cytotoxicity, cytokine signaling, cell migration, anti-viral and bacterial pathogen responses, and inflammation in NK cell clusters (Figure 3D and Supplemental Table 2). Moreover, gene expression changes in NK cell cluster with maternal SARS-CoV-2 infection enriched to GO terms associated with Fc-gamma receptor signaling, cytolysis, leukocyte-mediated cytotoxicity, regulation of NF-κB signaling, and viral responses (Figure 3E). This included increased expression of genes such as *GNLY*, *GZMH*, *IL32*, *IFNG*, *PRF1, IFITM1*, *IFI6, CCL5, and PYCARD* across the multiple NK cell subsets (Figure 3F). In line with these observations, an increase in the expression of degranulation marker CD107a by NK cells in response to PMA-ionomycin stimulation was observed in the maternal SARS+ group by flow cytometry (Figure 3G), suggesting increased NK cell activity. No significant differences were seen in the expression of MIP1β, IL-2, TNFα, or IFNγ by NK cells in response to stimulation (data not shown).

### Myeloid cells from babies born to mothers with asymptomatic/mild SARS-CoV-2 are hyper-responsive to bacterial TLR ligands

Increased immune activation at baseline was also evident within monocytes as indicated by increased module scores for cytokine signaling in the *IL1B* and *S100A8* classical monocyte clusters (Figure 4A and Supplementary Table 2). Functional enrichment of DEG revealed an over-representation of GO terms associated with responses to cytokines and regulation of immune responses within the *IL1B* cluster (Figure 4B). While chemokine expression was increased in this subset, the expression of MHC class II molecules was reduced in the maternal SARS+ group, as was the expression of several ISG (Figure 4C). These transcriptional patterns suggest a state of immune regulation in monocytes. To test this hypothesis, we assessed markers of monocyte activation using flow cytometry. While expression of CD16, TLR4, and CCR2 was increased in line with immune activation, the frequency of regulatory marker CD62L+ increased while that of co-stimulatory molecules CD83 and CD86, chemokine receptor CCR7, M1-like marker TREM1, and CSF1R decreased on monocytes, indicative of immune regulation ^46^ (Figure 4D). To test this hypothesis, UCBMC were stimulated with RSV or *E. coli* overnight and secreted factors were measured using Luminex. While both groups responded to RSV, induction of RANTES, IL-12p70, GROa, and Eotaxin was significantly attenuated in the maternal SARS+ group (Figure 4E and Table 3). In contrast, upon stimulation with *E. coli*, secreted levels of TNFα and IL1RA were significantly higher in the maternal SARS+ group (Figure 4F and Table 3). Collectively, these data suggest the rewiring of fetal monocytes towards a state of tolerance to viral antigens but enhanced responses to bacterial ligands.

**Figure 4:**
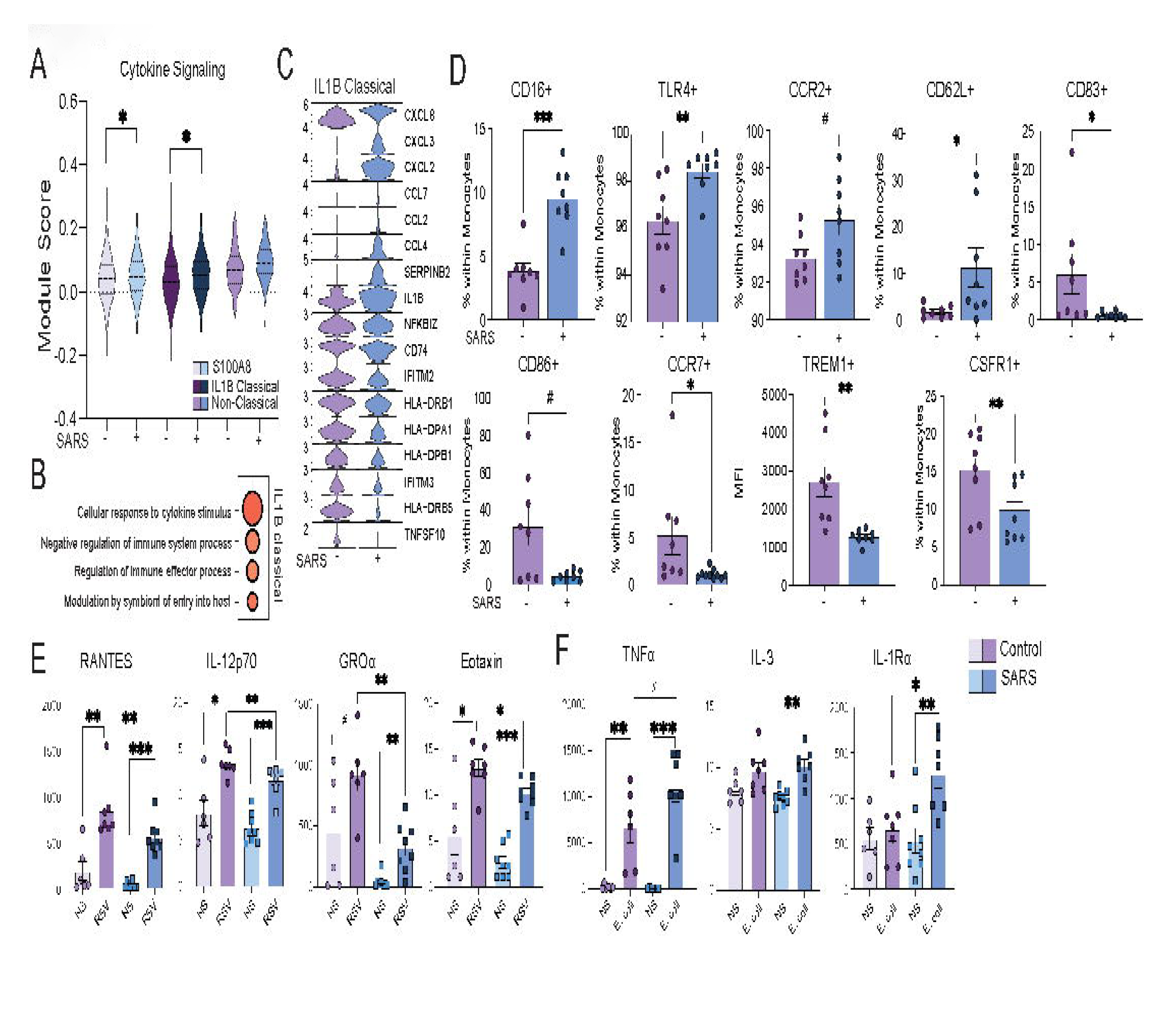
The impact of maternal SARS-CoV-2 infection on fetal myeloid cells. (A) Violin plot of module scores within clusters of monocyte subsets associated with cytokine and chemokine signaling. (B) Bubble plot of functional enrichment of top genes within the IL-1B classical monocyte cluster. The bubble size represents the number of genes mapping to the term, whereas the color represents the level of statistical significance. (C) Violin plots of select statistically significant DEG within non-classical and IL-1B classical monocytes. (D) Bar graphs of significant differences in UCB monocyte activation phenotypes by maternal infection status. (E-F) Scatterplot comparing select immune mediators secreted in culture supernatants of UCBMC stimulated overnight with (E) RSV or (F) *E. coli* in control and maternal SARS+ groups. (#=p<0.1, *=p<0.05, **=p<0.01, ***=p<0.001).

**Table 3:** Antiviral and antimicrobial responses of UCB Monocytes. Concentration (pg/mL) of immune mediators from cell culture supernatant of cells from controls and maternal SARS+ groups stimulated with RSV or *E. coli.* (#=p<0.1, *=p<0.05, **=p<0.01, ***=p<0.001, ****=p<0.0001)

| RSV | Control |  |  | SARS+ |  |  | RSV<br>Control:SARS+ |
| --- | --- | --- | --- | --- | --- | --- | --- |
|  | NS | RSV | p-value | NS | RSV | p-value | p-value |
| VEGF | 8.644 | 833 | ** | 16.44 | 793.2 | *** |  |
| TNFα | 178.9 | 169.7 | | 29.82 | 57.4 | # | # |
| RANTES | 186.4 | 844.1 | ** | 35.3 | 555.4 | *** | ** |
| PD-L1 | 25.21 | 41.11 | ** | 20.29 | 39.03 | *** |  |
| PDGF-AB/BB | 0.4893 | 7.286 | ** | 0.08323 | 7.142 | *** |  |
| PDGF-AA | 51.88 | 936.7 | ** | 50.96 | 938.2 | *** |  |
| MIP-3β | 17.23 | 22.23 | # | 11.24 | 17.09 | *** | |
| MIP-1β | 698.4 | 932.9 | | 181.4 | 600.5 | * | |
| MCP-1 | 4245 | 5346 |  | 708.2 | 8343 | ** |  |
| IP-10 | 0.4971 | 46.97 | * | 0.2228 | 100.1 | *** |  |
| IL-6 | 333.9 | 54125 | ** | 138.4 | 51648 | *** |  |
| IL-1Rα | 556.9 | 1010 | | 533.5 | 1385 | * | |
| IL-1α | 25.5 | 67.27 | # | 2.928 | 63.65 | *** | |
| IL-15 | 0.7462 | 2.72 | ** | 0.5025 | 2.551 | *** |  |
| IL-12p70 | 7.992 | 13.44 | * | 6.334 | 11.59 | *** | ** |
| IL-10 | 23.97 | 58.47 | # | 9.503 | 28.69 | ** |  |
| IFNγ | 30.71 | 32.25 | | 26.32 | 29.74 | * | |
| IFNβ | 0.4652 | 53.06 | ** | 0.2315 | 51 | *** | |
| IFNα | 1.957 | 14.72 | ** | 1.644 | 13.51 | *** | |
| GROα | 452.5 | 921.5 | # | 46.99 | 326.4 | ** | ** |
| GROβ | 212 | 1104 | ** | 32.27 | 847.6 | *** | |
| GM-CSF | 59.65 | 88.57 | # | 10.1 | 48.09 | ** | # |
| G-CSF | 79.49 | 135.5 | * | 3.994 | 26.05 | *** | * |
| Fractalkaline | 90.66 | 140.6 | * | 70.89 | 127 | *** |  |
| flt-3L | 44.01 | 63.11 | * | 32.34 | 60.44 | *** |  |
| FGFβ | 1.88 | 358.4 | ** | 0.8137 | 348.3 | *** | |
| Eotaxin | 5.595 | 12.95 | * | 2.704 | 10.27 | *** | * |
| EGF | 6.887 | 9.028 | * | 6.446 | 8.342 | *** |  |
| CD40L | 785.6 | 992.3 | # | 627.7 | 871.7 | *** |  |

| *E. coli* | Control |  |  | SARS+ |  |  | *E. coli*<br>Control:SARS+ |
| --- | --- | --- | --- | --- | --- | --- | --- |
|  | NS | *E.coli* | p-value | NS | *E.coli* | p-value | p-value |
| TRAIL | 2.534 | 9.728 | ** | 0.6544 | 9.556 | *** |  |
| TNFα | 178.9 | 6698 | ** | 29.82 | 10915 | *** | # |
| TGFα | 2.587 | 7.336 | * | 1.968 | 7.375 | *** | |
| RANTES | 186.4 | 157.7 |  | 35.3 | 82.93 | * |  |
| PD-L1 | 25.21 | 51.51 | ** | 20.29 | 49.84 | *** |  |
| PDGF-AB/BB | 0.4893 | 4.055 | ** | 0.08323 | 4.093 | *** |  |
| MIP-3β | 17.23 | 9.119 | * | 11.24 | 9.028 | * | |
| MIP-3α | 3.852 | 85.88 | ** | 2.903 | 53.48 | *** | |
| MIP-1β | 698.4 | 41542 | ** | 181.4 | 44049 | *** | |
| MIP-1α | 734 | 3252 | ** | 190.5 | 3423 | *** | |
| IP-10 | 0.4971 | 4.463 | ** | 0.2228 | 11.2 | *** |  |
| IL-6 | 333.9 | 16161 | ** | 138.4 | 15194 | *** |  |
| IL-3 | 8.183 | 9.78 | # | 7.599 | 10.23 | ** |  |
| IL-1Rα | 556.9 | 659.5 | | 533.5 | 1263 | ** | * |
| IL-1β | 75.76 | 1974 | ** | 11.92 | 3538 | *** | |
| IL-1α | 25.5 | 139.7 | * | 2.928 | 150.2 | *** | |
| IL-15 | 0.7462 | 2.48 | ** | 0.5025 | 2.317 | *** |  |
| IL-12p70 | 7.992 | 18.47 | ** | 6.334 | 19.41 | *** |  |
| IL-10 | 23.97 | 980.6 | ** | 9.503 | 1413 | *** |  |
| IFNγ | 30.71 | 38.78 | # | 26.32 | 41.08 | *** | |
| IFNβ | 0.4652 | 1.881 | ** | 0.2315 | 2.141 | *** | |
| IFNα | 1.957 | 5.337 | ** | 1.644 | 5.187 | *** | |
| GZMB | 28.72 | 26.92 |  | 18.3 | 24.45 | * |  |
| GROα | 452.5 | 842.2 | | 46.99 | 725.4 | *** | |
| GROβ | 212 | 4996 | ** | 32.27 | 5298 | *** | |
| GM-CSF | 59.65 | 1334 | ** | 10.1 | 2044 | *** |  |
| Fractalkaline | 90.66 | 245.5 | ** | 70.89 | 247.3 | *** |  |
| FLT-3L | 44.01 | 84.86 | ** | 32.34 | 82.96 | *** |  |
| FGFb | 1.88 | 10.46 | ** | 0.8137 | 20.02 | *** |  |
| Eotaxin | 5.595 | 15.8 | ** | 2.704 | 16.48 | *** |  |
| EGF | 6.887 | 10.43 | ** | 6.446 | 10.57 | *** |  |
| CD40L | 785.6 | 1620 | ** | 627.7 | 1606 | *** |  |

Finally, within the stem cell cluster, module scores for cytokine signaling, cell migration, and mitosis were increased, suggesting an altered differentiation program (Supplemental Figure 1I and Supplemental Table 2). Interestingly, differential gene expression analysis of the “proliferating cells” subset showed an over-representation of GO terms associated with inflammatory responses, wound healing, and regulation of viral processes (Supplemental Figure 1J) with increased expression of *LYZ*, *CRIP1*, *CD52*, *LGALS1*, and *S100A8* suggesting that these cells may be myeloid in nature (Supplemental Figure 1K).

### Maternal SARS-CoV-2 infection is associated with increased frequency and activation of fetal Hofbauer cells

Given the observed changes in circulating fetal immune cells and our recently described changes in decidual leukocytes with maternal SARS-CoV-2 infection ^12^, we next interrogated the impact of maternal SARS-CoV-2 infection on the immune landscape of chorionic villi (fetal side of the placenta). No viral RNA was detected in any of the villous tissue samples as measured by qPCR. Since immune cells in the villi are predominantly myeloid, we sorted CCR2+CD14+ (monocytes and monocyte-derived macrophages) and CCR2-CD14+ (tissue-resident macrophages) from frozen villous leukocytes and performed scRNA-seq on multiplexed controls (n=8) and maternal SARS+ samples (n=6). Dimensionality reduction and clustering revealed 10 unique cell clusters that contained cells from both groups (Figure 5A and Supplemental Figure 2A). These clusters were annotated (Figure 5B and Supplemental Figure 2B) based on markers previously described for the first-trimester villous immune landscape ^42^. HBCs were defined based on high levels of *FOLR2* and low levels of *HLA-DRA* with a proliferating HBC cluster also expressing high levels of *MKI67*. Placenta-associated maternal macrophages and monocytes (PAMM) clusters were identified based on the relative expression of *CD14*, *CCR2*, *CD9*, *HLA-DRA,* and *FOLR2*. In addition, other maternal infiltrating macrophages were detected and identified based on relative expression of *HLA-DRA*, *CCL4*, *APOE*, *IL1B*, *CCL20*, *CXCL10*, and *ISGs* (Figure 5B and Supplemental Figure 2B).

**Figure 5:**
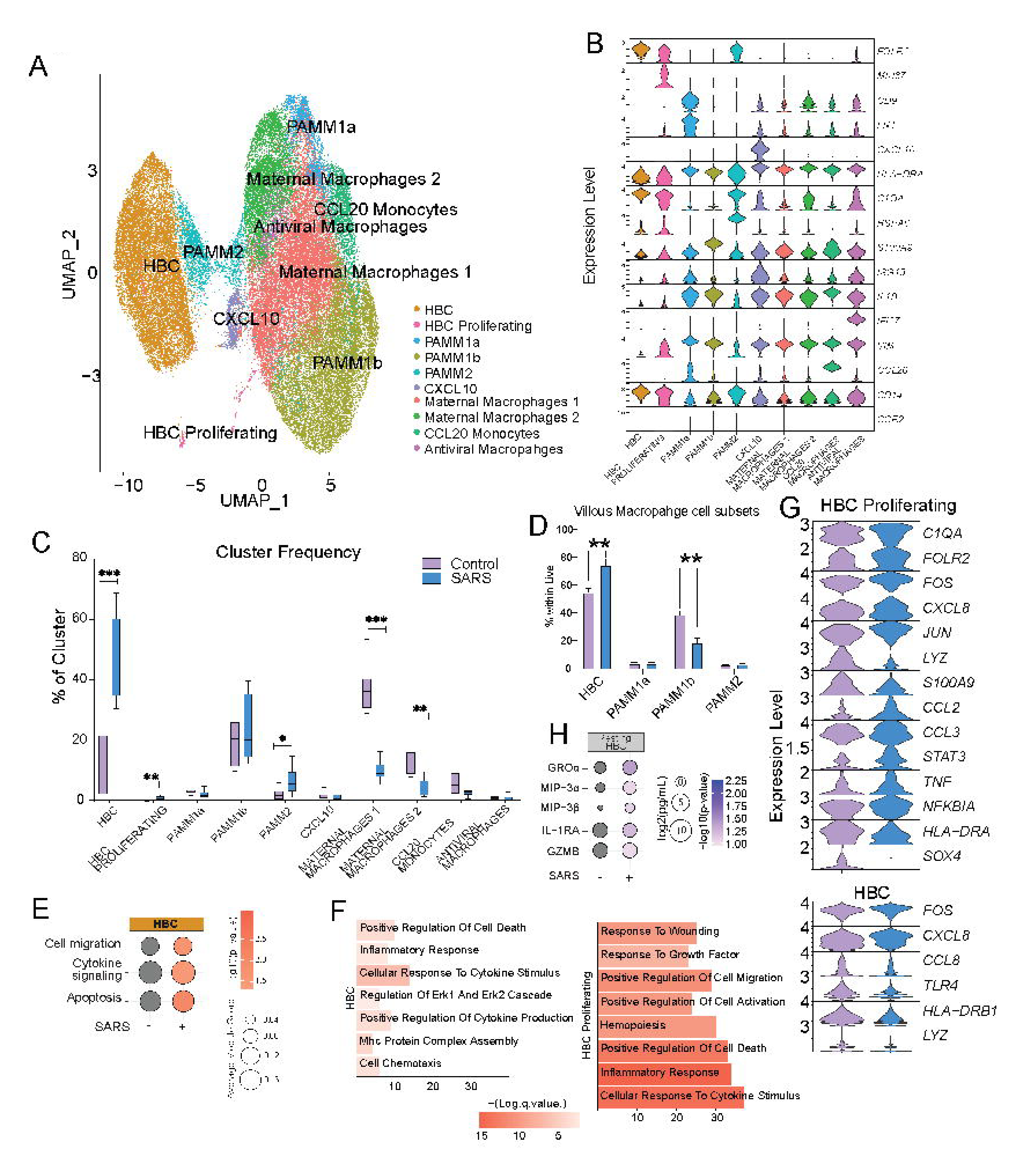
Impact of maternal SARS-CoV-2 infection on immune cells in the villous compartment. (A) Uniform Manifold Approximation and Projection (UMAP) of 48,553 immune cells within the villous compartment showing 10 clusters. (B) Violin plots of marker genes that were used for cluster annotation. (C) Box and whisker plots comparing relative cluster frequencies by infection status. (D) Bar graph comparing villous monocyte/macrophage subsets identified by flow cytometry within the live gate. (E) Bubble plot comparing module scores within HBC clusters for the terms indicated. The bubble size represents the average module score, whereas the color represents the level of statistical significance. (F) Barplot of GO terms identified in Metascape for DEG between controls and maternal SARS+ groups from the indicated cluster. The length of the bar indicates the number of genes associated with the term and the color intensity represents statistical significance. (G) Violin plots of select statistically significant DEG within the indicated cluster. (H) Bubble plot comparing secreted levels of immune factors by resting HBCs. The bubble size represents analyte concentration, whereas the color represents the level of statistical significance. (*=p<0.05, **=p<0.01, ***=p<0.001).

While maternal SARS-CoV-2 infection was associated with elevated frequencies of resting and proliferating HBC and PAMM-2 cells (infiltrating decidual macrophages), additional subsets of infiltrating maternal macrophages were decreased in the maternal SARS+ group (Figure 5C). The increased frequency of HBC was confirmed by flow cytometry (Figure 5D and Supplemental Figure 2C). Furthermore, module scores of gene signatures associated with cell migration, cytokine signaling, and apoptosis were elevated in HBC in the maternal SARS+ group (Figure 5E and Supplemental Table 2). Interestingly, DEG in both HBC subsets in the maternal SARS+ group mapped to pathways associated with inflammatory and cytokine responses (Figure 5F). These included both cytokines/chemokines (*CXCL8, CCL2, TNF*) and canonical transcription factors (*FOS, JUN, STAT3, NFKBIA)* associated with macrophage activation (Figure 5G). To test whether fetal Hofbauer cells were activated with maternal SARS-CoV-2 infection, we cultured purified HBC (CD14+FOLR2+HLA-DR-) for 16 hours and measured secreted levels of cytokines and chemokines at baseline. Indeed, maternal SARS-CoV-2 infection was associated with increased secretion of immune factors associated with myeloid cell recruitment (MIP-3α, MIP-3β) and activation (GROα and IL-1RA) (Figure 5H).

### Single-cell analysis of term placental villi reveals fetal macrophage adaptations to maternal SARS-CoV-2 infection

Flow analyses of macrophage populations within placental villi revealed a decrease in the frequency of maternal macrophages (PAMM1b) in the maternal SARS+ group, while PAMM1a (maternal monocyte) frequencies remained unchanged (Figure 5D). However, both populations exhibited altered module scores for anti-viral and bacterial defenses, cell and cytokine signaling, cell migration, apoptosis, and inflammation (Supplemental Figure 2D and Supplemental Table 2). DEGs within the PAMM1a subset mapped to GO terms such as cell activation, cell death/apoptotic signaling, and vessel morphogenesis (Supplemental Figure 2E) and included an increase in the expression of *APOE*, *FN1*, *FCGR2B,* and *JUNB* (Supplemental Figure 2F). On the other hand, DEG in PAMM1b subset mapped to GO terms associated with immune activation, cytokine production, and immune effector processes (Supplemental Figure 2E). We observed up-regulation of *ATF4*, *CD55*, *EREG*, *FCN1*, *THBS1*, and class-I MHC molecules (*HLA-A*, *HLA-F*) and down-regulation of complement transcripts (*C1QA*, and *C1QB*) (Supplemental Figure 2F) in the maternal SARS+ group. Finally, while flow analyses revealed no differences in the proportion of PAMM2 cells (Figure 5D), maternal SARS-CoV-2 infection was associated with increased module scores for cell signaling, migration, and inflammation (Supplemental Figure 2D and Supplemental Table 2). Importantly, the maternal SARS+ group was associated with downregulation of *IL1B*, *HLA-DRA*, *S100A8/9*, *CXCR4*, *IFI30*, *TREM1/2* and upregulation of *C1QA*, *CCL2*, and *CSF1R* (Supplemental Figure 2F).

In addition to canonical macrophage populations residing in the placental chorionic villi, we identified additional clusters - a CXCL10high cluster, two maternal macrophage clusters, a CCL10high monocyte cluster, and an antiviral macrophage cluster (Figure 5B and Supplemental Figure 2B). All infiltrating clusters had increased cell migration and cytokine signaling modules in the maternal SARS+ group (Supplemental Figure 3A and Supplementary Table 2). A consistent theme across these monocyte/macrophage subsets was the altered expression of genes involved in anti-microbial responses, inflammatory responses, and antigen processing and presentation (Supplemental Figure 3B, 3C, and Supplementary Table 2). Gene markers associated with immune activation were elevated in different myeloid subsets – neutrophil chemoattractant CXCL8 in infiltrating maternal macrophages, interferon-stimulated genes (IRF1, IFI6) in CCL20 monocytes, and alarmins (S100A8, S100A9) in antiviral macrophage clusters. We, therefore, posit that an elevated baseline activation state might alter their functional responses to pathogens. We tested this hypothesis by purifying the CD14+ compartment from chorionic villi and stimulating them with viral and bacterial PAMPs. Our analysis of supernatants demonstrated significantly higher levels of pro-inflammatory IL-1α, Flt-3L, and MCP-1 following viral TLR ligand stimulation but no differences in secreted cytokines in response to bacterial PAMPs (Supplemental Figure 3D).

## DISCUSSION

The Developmental Origins of Health and Disease (DOHaD) hypothesis postulates that fetal exposure to environmental insults (such as poor nutrition, infection, chemicals, or hormonal perturbations) during critical periods of development and growth influences organ system development and susceptibility to diseases in later life ^47^. Infectious diseases provoke the maternal immune system ^48^, which in turn impacts the risk for disease risk in offspring ^49^. Immune cell ontogeny in early life is particularly vulnerable to maternal infection, as shown by the higher mortality risk from infectious disease in HIV-exposed but uninfected infants ^50^ potentially due to altered neonatal Th17 and Treg immune balance ^51^. Additionally, infection of placental HBCs by ZIKV leads to the production of type 1 interferons and pro-inflammatory cytokines and chemokines, eliciting placental inflammation, poor placental perfusion, and poor fetal outcomes^52^. Malaria in pregnancy is associated with dysregulation of placental development and preterm birth, with an increased risk of mortality due to complications such as acute respiratory illness and sepsis ^53^. Similarly, SARS-CoV-2 also provokes maternal immune activation as indicated by increased levels of systemic immune mediators ^26, 54^. While most studies of COVID-19 in pregnancy have focused on severe cases, there is growing evidence suggesting that mild maternal SARS-CoV-2 alters inflammatory responses at the MFI. Therefore, there is a critical need to understand the impact of mild/asymptomatic maternal SARS-CoV-2 infection on the immune landscape of fetal chorionic villous tissues and fetal circulation.

Studies included herein used UCB, a practical surrogate for newborn blood ^55, 56^. Despite the lack of vertical transmission, levels of several chemokines and cytokines necessary for anti-microbial responses were reduced in UCB plasma in the maternal SARS+ group. These observations differ from data reported for non-gravid adult SARS-CoV-2 infection where levels of VEGF, GM-CSF, TNFα, IL-23, IL-4, IL-7, CXCL9, CCL2, CCL3, CCL4, and CCL11 are all elevated^57–59^. Our data also differ from those reported in a recent study where a lack of differences in concentration of many of these markers (except a modest increase in IFNα, in the SARS+ group) were noted ^60^. In contrast, S100B, IL-18, and PDGFBB were elevated in our study. These factors are linked to neurologic insults in newborns. S100B is a marker of neurological complications ^61^ while IL-18 and PDGF-BB levels are increased in circulation after traumatic spinal cord or brain injury to repair vascular dysfunction ^62–64^. Indeed, maternal SARS-CoV-2 infection during pregnancy is linked to cases of perinatal brain injury and a greater rate of neurodevelopmental diagnoses in the first year of life ^65–67^. Consistent with our findings, IL-18 secretion has been shown to be elevated in the circulation of neonates born to individuals with asymptomatic SARS-CoV-2 infection and plays an important role in fetal cortical injury and adverse neurobehavioral outcomes ^28, 64, 68^. These results indicate a generally immunosuppressive environment in UCB plasma in the maternal SARS+ group compared to controls, aside from elevated markers of brain injury consistent with neurodevelopmental issues reported in previous reports.

Here, we report an increased number of white blood cells characterized by an increased frequency of monocytes and granulocytes in UCB. These observations are in line with increased monocyte frequencies in adults with severe SARS-CoV-2 infection, driven by an expansion of intermediate and non-classical monocyte subsets ^69^ as well as infants younger than 1 year of age with mild COVID-19 ^70^. We observed an expansion of non-classical monocytes measured by both flow cytometry and scRNA-Seq. Additionally, cytokine signaling pathways were activated in classical monocyte subsets, further reflected by increased expression of genes responsible for monocyte recruitment and cytokine signaling in the IL-1β classical monocyte subset. Interestingly, expression of HLA-DR and interferon-stimulated genes (ISG) was reduced in classical monocytes from the maternal SARS+ group. This is in contrast to earlier studies that reported upregulation of interferon-stimulated genes (ISG) and MHC genes in UCB monocytes with maternal SARS-CoV-2 infection ^71^. These discrepancies in findings may be due to the emergence of the more severe delta SARS-CoV-2 variant during sample collection for this study that was not present when the prior study was completed ^71^. Additionally, the timing of infection could potentially influence inflammation in UCB. For example, the prior study included pregnant participants that were infected with SARS-CoV-2 exclusively in the third trimester, whereas this study included participants infected with the virus over the course of pregnancy ^71^.

Dysregulated monocyte responses linked to COVID-19 pathogenesis and ensuing cytokine storm during infection in adults ^72^. Our results show that frequencies of CD16+, TLR4+, and CCR2+ monocytes were increased in UCB in the maternal SARS+ group. We also noted increased expression of CD62L on UCB monocytes, consistent with reports of higher proportions of CD62L-positive monocytes in adults with COVID-19 ^73–75^. On the other hand, decreased expression of co-stimulatory molecules CD83 and CD86 suggests a more regulatory monocyte phenotype. This hypothesis is further supported by the decreased expression of receptors important for recruiting DCs and activation of T cells (CCR7), amplification of inflammation (TREM1), and proliferation (CSF1R). Alterations in monocyte activation state may contribute to dysregulated antimicrobial responses. Indeed, UCB monocytes generated an increased response to stimulation with *E. coli*, in line with increased expression for the LPS receptor TLR4. However, monocyte responses to RSV were suppressed. We have previously shown the opposite trend with aged adults with COVID-19, where innate immune signaling was preferentially geared towards antiviral responses ^75^, indicating that exposure to SARS-CoV-2 *in utero* has a distinct impact on innate immune responses compared to infection in later life. Minimal studies exist on the impact of maternal SARS-CoV-2 infection on newborn susceptibility to pathogens in early life. However, acute respiratory distress syndrome and pneumonia-like symptoms are more frequent in newborns of mothers with SARS-CoV-2 during pregnancy ^76^.

A healthy newborn’s adaptive immune system is primarily comprised of naïve lymphocytes with limited immune memory and effector function ^77, 78^. However, our analysis of UCB revealed accelerated lymphocyte maturation indicated by the increased relative abundance of memory cells and proportionally less naïve T and B cells, increased expression of the proliferation marker Ki67, and effector marker KLRG1. Infectious exposures can broadly impact the developing T cell compartment and elicit pathogen-specific T-cell responses. For example, malaria-specific CD4+ T cell responses in UCB correlate with protection against malaria infection in early life ^79, 80^, suggesting that priming of pathogen-specific CD4+ T cell responses *in utero* can confer protection later in life. Naïve T cells can also acquire memory T-cell markers and functional properties during cytokine-driven proliferation independent of antigen encounter ^81^. Previous studies report that asymptomatic maternal SARS-CoV-2 infection resulted in dampened T_H_1 and T_H_17 responses and reduced T cell repertoire diversity that does not extend to neonatal circulation ^14^. Here, stimulation of neonatal T cells from the maternal SARS+ group resulted in a dampened response to anti-CD3/CD28 stimulation. Poor T cell responses may result in impaired anti-microbial defenses. The mechanisms by which maternal SARS-CoV-2 infection dysregulates T cells in early life may be cell-intrinsic, as transcriptional analysis of the lymphocytes showed upregulation of mitosis, cytokine signaling, inflammation, and migration pathways. Future studies should address the molecular underpinnings (epigenetic and signaling) of these expanded yet functionally impaired fetal T cells and identify their unique phenotypes and antigen specificities.

Normally, early-life B cell responses to antigens are muted and exhibit distinct gene expression profiles with limited B cell activation compared to adults ^82^. Other studies have shown that maternal SARS-CoV-2 infection in the third trimester has no significant impact on the frequency of CD19+ B cells ^83^. However, the frequencies of memory B cell subsets and their functional capacity were not addressed. Our results show significant alterations in UCB humoral immunity in the maternal SARS+ group. Despite the increased frequency of memory subsets, UCB B cells from the maternal SARS+ group were less responsive to stimulation, as shown by decreased expression of co-stimulatory (CD40) and activation markers (CD83), indicating an early activation of UCB B cells in the SARS+ group.

Here, we report a decrease in the frequency of ISG-expressing NK cells but an expansion of cytokine-producing CD56^Bright^ NK cells in UCB. These observations align with the increase in expression of genes responsible for type 2 interferon responses, cytolytic functions, and monocyte activation and recruitment. Moreover, UCB NK cells from the maternal SARS+ group expressed higher levels of degranulation molecules, indicating the heightened cytotoxic potential of these cells. Our data align with other studies that reported a decrease in NK cell frequencies in adults with SARS-CoV-2 infection as well as in neonates of mothers with SARS-CoV-2 infection during pregnancy, but an activated phenotype ^84, 85^. Increased activation of fetal NK cells is perhaps a compensatory mechanism against dampened T cell responses.

Maternal SARS-CoV-2 infection has been shown to compromise placental function as shown by the increased risk of pre-eclampsia ^86^, abnormal placental histopathologic changes indicative of hypoxia ^87^, and placental inflammation ^88–90^. However, the impact of SARS-CoV-2 infection on the immune landscape at the maternal-fetal interface has been relatively understudied. The human placenta is comprised of maternal (decidua) and fetal (chorionic villous) tissues, each with unique immune repertoires ^91^. Studies evaluating how decidual leukocytes are altered by maternal SARS-CoV-2 infection during gestations are inconsistent. Early reports suggested that COVID-19 infection in the first trimester did not alter leukocyte frequencies within the decidua ^92^. Other studies have demonstrated increased macrophages, NK cells, and T cells, accompanied by elevated expression of various cytokines (IL-6, IL-8, IL-10, TNFα) with maternal SARS-CoV-2 infection in the first trimester ^92, 93^. Our previous studies show significant perturbations induced by maternal SARS-CoV-2 infection in the decidua ^12, 38^, including attenuated antigen presentation and viral signaling pathways, reduced frequencies of tissue-resident decidual macrophages, and upregulated cytokine/chemokine signaling in monocyte-derived decidual macrophages ^12, 38^.

The maternal decidua is in direct and/or indirect contact with fetal membranes, placental villi, and maternal circulation. Therefore, perturbations in the maternal circulation and decidua tissues likely expand into fetal villous tissues and, thus, fetal circulation ^94, 95^. The fetus-derived chorionic villous is comprised exclusively of macrophages, a major population being Hofbauer cells (HBC) that secrete factors important for placental angiogenesis and remodeling but also offer protection from bacterial pathogens^42, 96^. HBC also expand in numbers with adverse pregnancy outcomes ^97–99^. Our single cell and flow cytometry data revealed an increase in the frequency of HBCs in chorionic villi compared to uninfected negative controls, consistent with previous reports ^100^. Furthermore, our analysis indicates increased activation of HBC in the maternal SARS+ group as suggested by the increase in the expression of genes associated with migration, cytokine signaling, and apoptosis in HBCs, as well as increased secretion of inflammatory chemoattractants (GROα, MIP-3α, MIP-3β) and mediators of cytotoxicity (IL-1RA and GZMB). Given that HBs play a central role in pathogen sensing and host defense and the absence of active infection in the placenta, these findings suggest non-specific activation in response to inflammatory signals from the maternal compartment.

In addition to fetal HBC, the placental villi harbor additional maternal monocyte/macrophage subsets that assist in placental repair mechanisms and possibly the prevention of microbial transmission ^42, 96^. Among these, are PAMM1a cells (maternal macrophages), which did not vary in numbers with maternal infection but exhibited transcriptional signatures associated with cell activation, cell death, and vessel morphogenesis. These observations suggest activation of the infiltrating maternal macrophages to repair possible placental structural damage caused by SARS-CoV-2. PAMM1b cells are less abundant in healthy villous tissues and are transcriptionally comparable to adult circulating classical monocytes ^42^. In addition to their expansion with maternal SARS-CoV-2 infection, we observed a significant rewiring of their transcriptional states suggestive of enhanced activation and immune effector function. In this study, we also report five additional clusters of macrophages, arguably, new cell states of infiltrating maternal macrophages driven by chemokine expression (*CXCL10*, *CCL10*) that exhibited elevated expression of alarmins, ISGs, *NFKB1*, and MHC-I molecules.

Interestingly, frequencies of infiltrating decidual macrophage (PAMM2), the bona fide placenta-resident macrophages of maternal origin, did not vary with maternal SARS-CoV-2 infection. This contrasts with their elevated frequencies in the decidual compartment, as previously described in SARS-CoV-2 infected mothers ^38^. However, infection resulted in the down-regulation of several genes involved in host defense and anti-viral immunity. This observation is in line with reports showing dampened expression of genes important for antiviral innate immunity (*IFNB*, *IFIT1*, *MXA*) and cytokine responses (*IL6*, *IL1B*) in chorionic villous tissues by qPCR regardless of gestational age during infection ^34, 101^. Taken together, these findings suggest that maternal SARS-CoV-2 infection triggers opposing adaptations within different maternal myeloid cells residing in the fetal chorionic villi – with heightened activation and antiviral state in infiltrating maternal monocyte/macrophages and suppression of cytokine signaling and antigen-presentation pathways in rare infiltrating decidual macrophages. We argue that this enhanced antiviral state contributes to an augmented response to viral TLRs. Our findings suggest that fetal immune cells are not fully protected from inflammatory signals from mild/asymptomatic maternal SARS-CoV-2 infection. More importantly, this study highlights the unique functional adaptations within circulating and tissue-resident fetal myeloid and lymphoid cells in response to an ongoing/resolving maternal viral infection.

## AUTHOR CONTRIBUTIONS

Conceptualization, S.S., N.E.M., and I.M.; methodology, S.S., N.E.M. and I.M.; investigation, B.D., S.S., H.T., and N.M; writing, B.D., S.S., H.T., N.E.M, and I.M.; funding acquisition, N.E.M, and I.M.; participant enrollment, M.R. and N.E.M. All authors have read and approved the final draft of the manuscript.

## ACKNOWLEDGMENTS

We are grateful to all participants in the study. We thank the MFM Research Unit at OHSU for sample collection and Allen Jankeel, Michael Z. Zulu, Gouri Ajith, Isaac Cinco, and Hannah Debray at UCI for assistance with tissue processing. We thank Dr. Jennifer Atwood at the UCIT Institute for Immunology Flow Cytometry Core for assistance with FACS sorting, and imaging flow cytometry, and Dr. Melanie Oakes at the UCI Genomics Research and Technology Hub (GRT Hub) for assistance with 10x library preparation and sequencing.

## COMPETING INTERESTS

The authors declare that there is no conflict of interest regarding the publication of this article.

**Supplemental Table 1: Gene markers for human PBMC and villous leukocytes**

**Supplemental Table 2: Module scores for human PBMC and villous leukocytes**

**Supplemental Figure 1:**
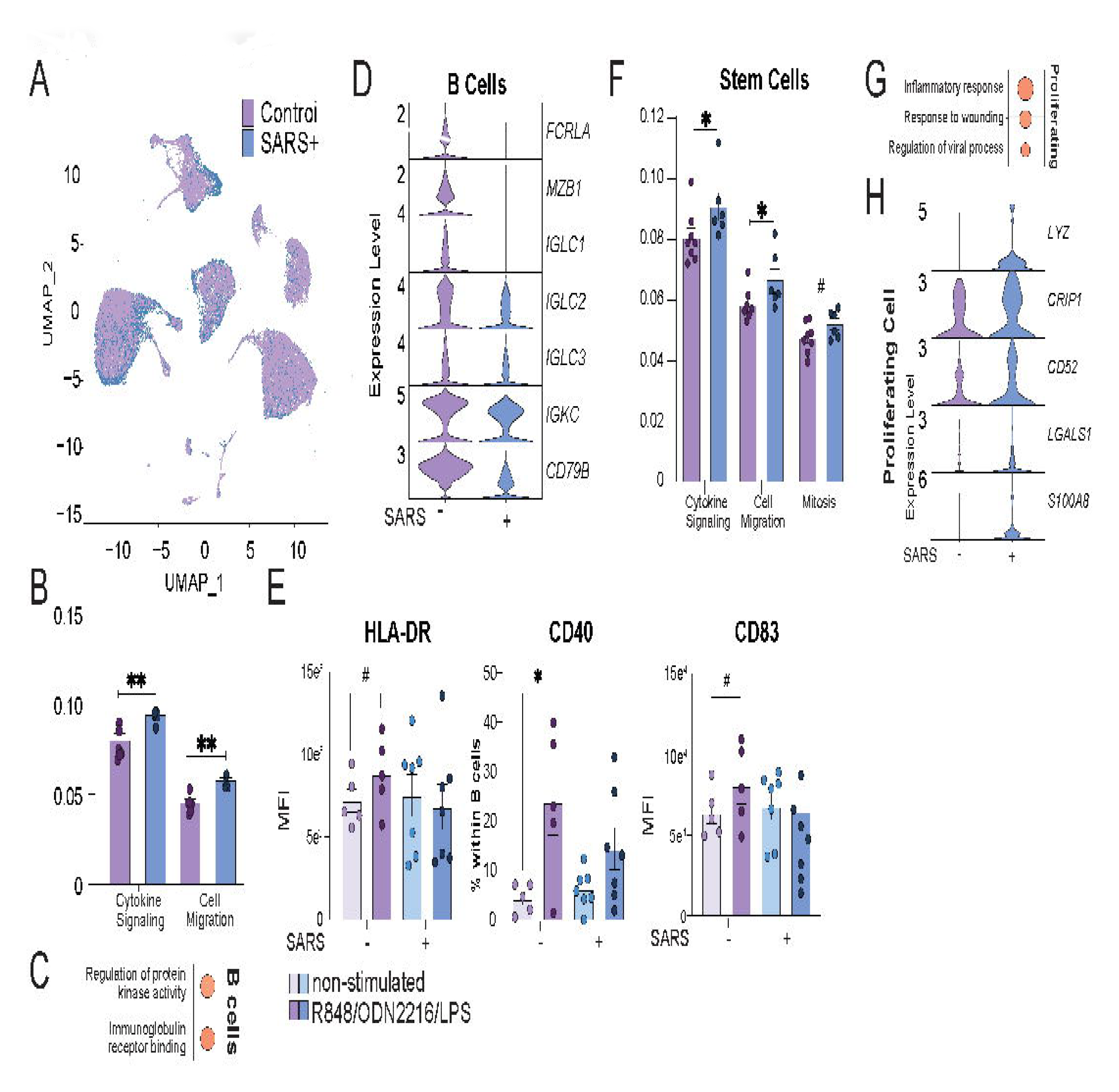
UCB Immune cell clusters by infection status and module scores. (A) UMAP of UCB immune cells colored by control (purple) and maternal SARS+ (blue) groups. (B) Bar graph comparing module scores within the B cell cluster for the terms indicated. (C) Bubble plot of functional enrichment of top genes within the B cell cluster. The bubble size represents the number of genes mapping to the term, whereas the color represents the level of statistical significance. (D) Violin plot of select statistically significant DEG within the B cell subset. (E) Bar graphs comparing B cell responses to stimulation with R848, ODN2216, and LPS (F) Bar graph of module scores within the stem cell cluster for the terms indicated. (G) Bubble plot of functional enrichment of top genes within the proliferating cell clusters. The bubble size represents the number of genes mapping to the term, whereas the color represents the level of statistical significance. (H) Violin plot comparing normalized transcript counts of statistically significant DEG within the proliferating cell subset. (#=p<0.1, *=p<0.05, **=p<0.01, ***=p<0.001).

**Supplemental Figure 2:**
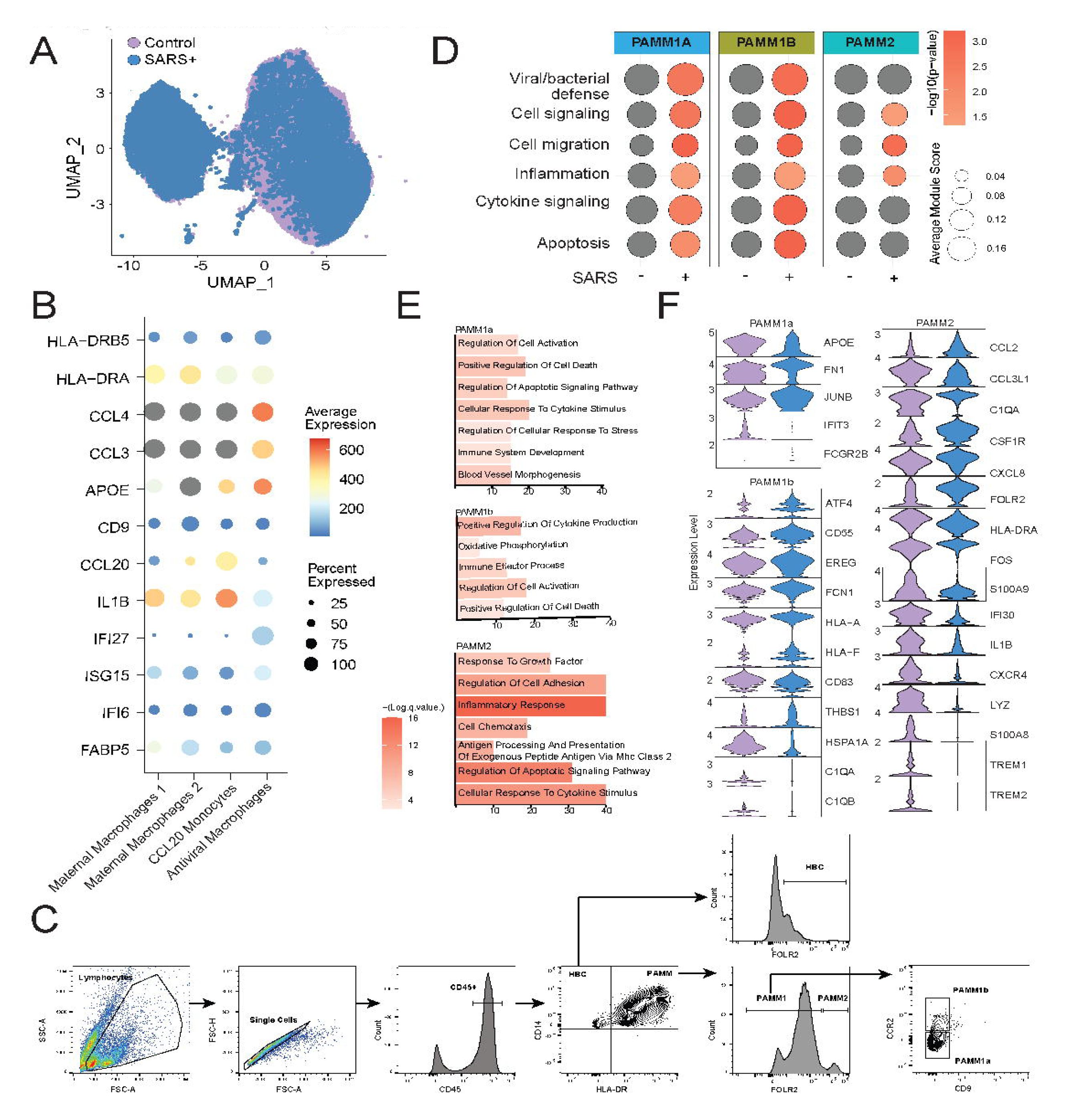
Impact of maternal SARS-CoV-2 infection on the immune landscape of the chorionic villous PAMMs. (A) UMAP highlighting villous immune cells from control and maternal SARS+ groups. (B) Bubble plot of additional marker genes used for cluster identification. The bubble size represents the amount of gene expression, whereas color represents the average gene expression. (C) Gating strategy to identify villous immune cell subsets. (D) Bubble plot of module scores within PAMM clusters for the terms indicated. The bubble size represents the average module score, whereas the color represents the level of statistical significance. (E) Barplot of GO terms identified in Metascape for DEG between controls and maternal SARS+ groups from the indicated cluster. Length of the bar indicated the number of genes associated with the term and the color represents the level of statistical significance. (F) Violin plots of select statistically significant DEG within indicated clusters.

**Supplemental Figure 3:**
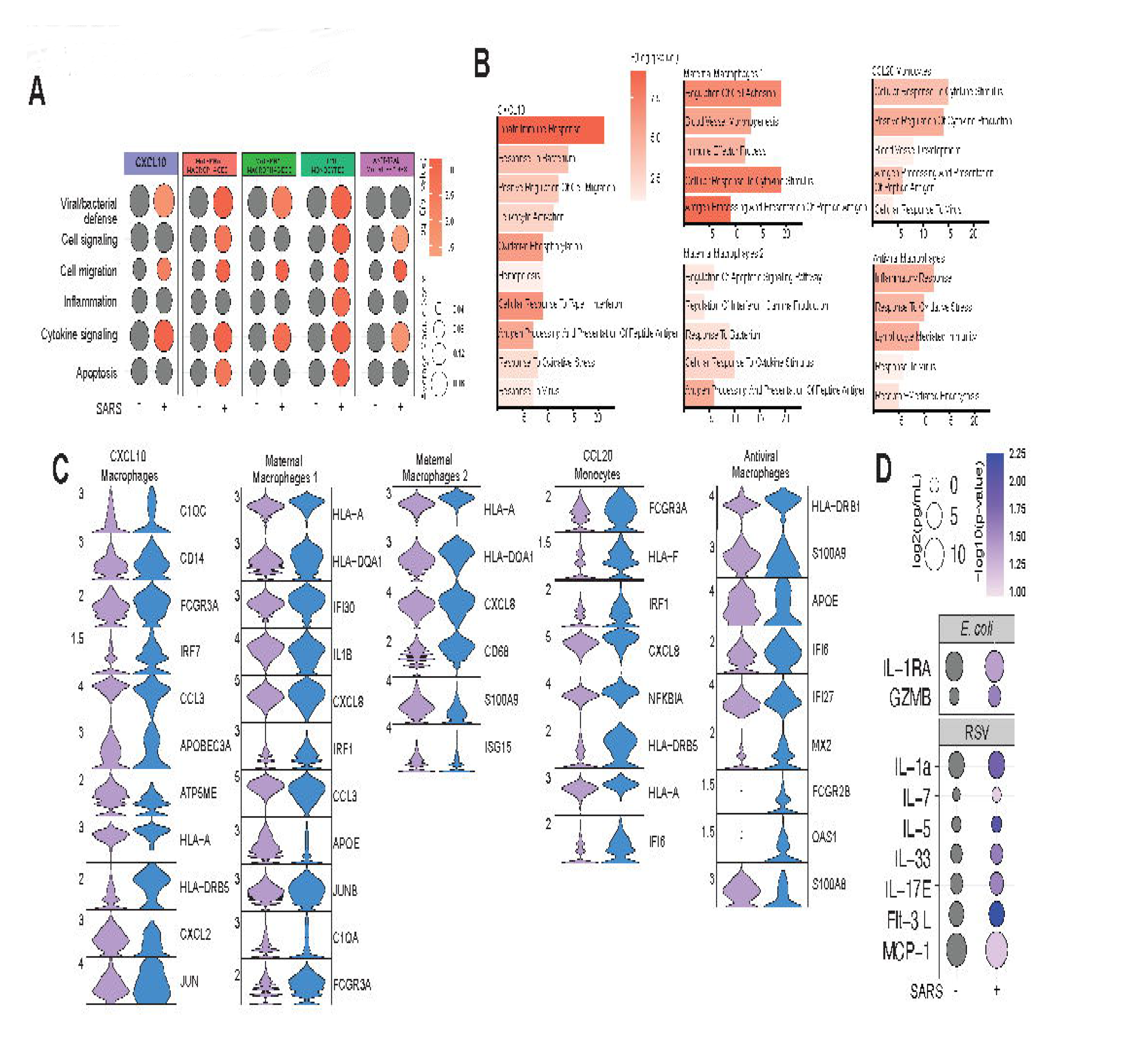
Impact of maternal SARS-CoV-2 infection on the immune landscape of the infiltrating myeloid cells in chorionic villi. (A) Bubble plot comparing module scores within infiltrating myeloid cell clusters for the terms indicated. The bubble size represents the average module score, whereas the color represents the level of statistical significance. (B) Barplot of GO terms identified in Metascape for DEG between controls and maternal SARS+ groups from the indicated cluster. Length of the bar indicated the number of genes associated with the term and the color represents the level of statistical significance. (C) Violin plots of select statistically significant DEG within indicated clusters. (D) Bubble plot comparing immune mediators produced CD45+CD14+ macrophages in response to *E.* coli (top) or RSV stimulation. The bubble size represents analyte concentration, whereas the color represents the level of statistical significance.

