## Supplemental Table 3 for "Mild/Asymptomatic Maternal SARS-CoV-2 Infection Leads to Immune Paralysis in Fetal Circulation and Immune Dysregulation in Fetal-Placental Tissues": Sup Table 3 Module scores.pdf

| Column1 |
| --- |
| LTF |
| TF |
| CAMP |
| JCHAIN |
| DEFB1 |
| DEFA1B |
| DEFA4 |
| IGHM |
| HIST1H2BJ |
| RNASE6 |
| RNASE7 |
| CTSG |
| HLA-E |
| PLA2G6 |
| SEMG2 |
| SLPI |
| PLA2G1B |
| FAU |
| BPIFA1 |
| AHSG |
| ITIH4 |
| INS |
| FN1 |
| LBP |
| HP |
| SCARB1 |
| NECTIN2 |
| ICAM1 |
| SENP7 |
| MX1 |
| PHB2 |
| RNF135 |
| CLPB |
| OAS1 |
| ZDHHC1 |
| PHB |
| AKAP1 |
| NLRP1 |
| IFIT1 |
| CXCL10 |
| PGLYRP3 |
| PGLYRP2 |
| GPX1 |
| DDB1 |

| |
|---|
| PLG |
| ELANE |
| GFI1 |
| ZC3H12A |
| TNFRSF1B |
| CCR5 |
| CD86 |
| BCL10 |
| PDE4B |
| WNT5A |
| CD80 |
| PLSCR4 |
| TNF |
| NFKBIL1 |
| TNFAIP3 |
| SERPINE1 |
| CDC73 |
| IL6 |
| RHOA |
| TNFSF4 |
| RIPK2 |
| VIM |
| MRC1 |
| CD36 |
| HMGB2 |
| XBP1 |
| MAPK8 |
| TRAF6 |
| GIT1 |
| JAK2 |
| TNIP1 |
| PDCD4 |
| PLAA |
| NR1D1 |
| ABCA1 |
| CMPK2 |
| IL10 |
| RARA |
| PLSCR3 |
| NR1H3 |
| NR1H4 |
| NKG2D |
| STAP1 |
| CTR9 |
| mml-mir-224 |

| |
|---|
| TRIM41 |
| NOD2 |
| LY96 |
| IL12B |
| NUGGC |
| TICAM1 |
| NFKBIB |
| NFKB1 |
| MAPK14 |
| AXL |
| DAB2IP |
| ADAM9 |
| mml-mir-342 |
| PAF1 |
| IRF8 |
| MALT1 |
| ZFP36 |
| PDCD1LG2 |
| CDK4 |
| NOS2 |
| RELA |
| SBNO2 |
| mml-mir-431 |
| mml-mir-433 |
| HAVCR2 |
| CHMP5 |
| IRF3 |
| TNIP3 |
| NLRP7 |
| PYCARD |
| CD180 |
| TLR4 |
| LDOC1 |
| CACTIN |
| ANKRD1 |
| IL37 |
| CD14 |
| UPF1 |
| ACOD1 |
| CXCL1 |
| CXCL6 |
| CXCL8 |
| AKAP8 |
| ABL1 |
| SIGLEC1 |

| |
|---|
| HRG |
| CFHR5 |
| F2 |
| REG3G |
| TPCN2 |
| TPCN1 |
| CTSL |
| HYAL2 |
| GAS6 |
| SFTPD |
| FGB |
| DYNLT1 |
| GAPDH |
| PF4 |
| NUCKS1 |
| SAP30 |
| SAP30BP |
| IFI27 |
| FBXL2 |
| TMEM41B |
| FMR1 |
| PSMC3 |
| RAB9A |
| APCS |
| PTX3 |
| PRKN |
| ZBED1 |
| CCNK |
| CCL8 |
| VAPA |
| TRIM11 |
| TRIM26 |
| TRIM15 |
| PML |
| TRIM21 |
| TRIM28 |
| TRIM35 |
| TRIM8 |
| TRIM13 |
| MID2 |
| TRIM32 |
| MARCHF2 |
| HDAC1 |
| TARDBP |
| ZNF639 |

| |
|---|
| JUN |
| POU2F3 |
| TFAP4 |
| INPP5K |
| REST |
| HMGA2 |
| SPINK5 |
| ITGAV |
| CH25H |
| TRIM10 |
| FCN3 |
| TRIM59 |
| SNX3 |
| GSN |
| FCN1 |
| MYD88 |
| SCNN1B |
| AZU1 |
| TUSC2 |
| NLRP6 |
| DHCR24 |
| PSMB8 |
| MAMU-DRA |
| MAML1 |
| YTHDC2 |
| TBC1D20 |
| PPIB |
| ZNF502 |
| CAV2 |
| CSF1R |
| IGF2R |
| PC |
| CFL1 |
| APOE |
| TAF11 |
| RRP1B |
| SNW1 |
| LEF1 |
| CHD1 |
| HPN |
| SMARCB1 |
| CDK9 |
| CCNT1 |
| CTDP1 |
| EP300 |

| |
|---|
| SP1 |
| SMARCA4 |
| PGC |
| KLK3 |
| KLK5 |
| KLK8 |
| SYK |
| PRF1 |
| CLEC7A |
| IFNG |
| FCER2 |
| F2RL1 |
| TMPRSS2 |
| CD74 |
| TRIM38 |
| CD4 |
| TMPRSS4 |
| P4HB |
| EPS15 |
| PIKFYVE |
| LRRC15 |
| ACE2 |
| FUCA2 |
| EXOC2 |
| EXOC7 |
| HS3ST5 |
| TRIM22 |
| TRIM6 |
| TRIM68 |
| TRIM14 |
| S100A14 |
| S100A8 |
| DUSP10 |
| CD96 |
| IL12RB2 |
| IL23R |
| PTAFR |
| SNCA |
| LIAS |
| NOCT |
| ADH5 |
| NFKBIA |
| PTGFR |
| MAPKAPK3 |
| PRDX3 |

| |
|---|
| TRIB1 |
| IL12A |
| TAB2 |
| PELI1 |
| ADAM17 |
| TIRAP |
| CEBPB |
| MAPKAPK2 |
| CPS1 |
| SMAD6 |
| PTGIR |
| CCR7 |
| FER |
| CD6 |
| F2R |
| RPS6KA3 |
| MAPK1 |
| SLC11A1 |
| KCNJ8 |
| GCH1 |
| ERBIN |
| SRR |
| IDO1 |
| CYP27B1 |
| NOS1 |
| IRAK1 |
| PLCG2 |
| UMOD |
| IRAK3 |
| TNFRSF11A |
| PTGER1 |
| TBXA2R |
| PTGER4 |
| IL1B |
| OTUD5 |
| MAPK3 |
| WDR83 |
| CHMP2B |
| CHMP7 |
| CHMP4C |
| CHMP2A |
| CHMP4B |
| CHMP6 |
| VPS4B |
| XPR1 |

| |
|---|
| AGTR1 |
| CLDN1 |
| GPR15 |
| CLEC5A |
| NECTIN4 |
| SIVA1 |
| DAG1 |
| CTSB |
| TYRO3 |
| NRP1 |
| AVPR1B |
| VAMP8 |
| NPC1 |
| GRK2 |
| SMPD1 |
| CLDN6 |
| INSR |
| SLC3A2 |
| UVRAG |
| AVP |
| CLEC4G |
| CD209 |
| VPS18 |
| DPP4 |
| ARL8B |
| DDX39B |
| THOC7 |
| THOC5 |
| THOC3 |
| THOC6 |
| THOC1 |
| THOC2 |
| RAB7A |
| VPS37B |
| IST1 |
| TSG101 |
| HSP90AB1 |
| NECTIN1 |
| CD81 |

| Column1 |
| --- |
| ZC3H12A |
| ZBTB7B |
| RC3H1 |
| IL2 |
| IL4 |
| TBX21 |
| SMAD7 |
| RC3H2 |
| FOXP3 |
| LOXL3 |
| IL27RA |
| NXPE3 |
| MALT1 |
| IL23A |
| PRKCQ |
| NLRP10 |
| CLEC7A |
| CARD9 |
| CLEC6A |
| CCR2 |
| BATF |
| ENTPD7 |
| RORA |
| SLAMF6 |
| IL6 |
| IRF4 |
| STAT3 |
| TRAF3IP2 |
| NOTCH1 |
| PHB |

| Column1 |
| --- |
| ARG1 |
| TBX21 |
| IFNB1 |
| HLX |
| BCL6 |
| SOCS5 |
| ANXA1 |
| PRKCQ |
| PRKCZ |
| DENND1B |
| GATA3 |
| RSAD2 |
| IL4 |
| CD81 |
| RARA |
| IL4R |
| IL18 |
| BATF |
| BCL3 |

| Column1 |
| --- |
| IL4R |
| IL33 |
| HAVCR2 |
| IL1RL1 |
| TBX21 |
| IL18 |
| IL1R1 |
| IL18R1 |
| IL1B |
| HLX |
| RIPK2 |
| SOCS5 |
| ANXA1 |
| CCR2 |
| PLA2G4A |
| IL23R |
| NLRP10 |
| SLC11A1 |
| IL12B |
| IL27RA |
| IL12RB1 |
| IL23A |
| IL27 |
| TCIRG1 |
| SEMA4A |
| LEF1 |
| TMEM98 |
| RELB |
| MTOR |
| SPN |
| STAT6 |
| TRAF6 |
| IL18BP |
| BCL3 |
| HRAS |

| Column1 |
| --- |
| EPHB6 |
| FYN |
| BTN3A1 |
| ABL1 |
| BCL10 |
| TNFRSF21 |
| JCHAIN |
| MAMU-DOA |
| PSMB8 |
| MAMU-DOB |
| RIPK2 |
| MAMU-DRB1 |
| IGHM |
| CTSH |
| HLA-E |
| RAG1 |
| CD4 |
| CTSL |
| SH2D1A |
| NKG2D |
| CD3E |
| CD3D |
| PRKCB |
| JAM3 |
| TSC1 |
| TEC |
| IFNB1 |
| CD7 |
| CD79A |
| C1QBP |
| ITK |
| BMX |
| CTSS |
| BTK |
| TNFRSF11A |
| ANXA1 |
| GPR183 |
| CD79B |
| DOCK2 |
| INS |
| RPL22 |
| BRAF |
| LEF1 |
| PRR7 |

| |
|---|
| BCL11B |
| RHOA |
| SLAMF1 |
| CARD11 |
| CCL26 |
| ADAM8 |
| CXCL16 |
| CCL7 |
| CCL16 |
| CCL23 |
| CCL18 |
| CCL20 |
| CX3CL1 |
| CCL17 |
| MSMP |
| PRKDC |
| IKZF1 |
| LY6D |
| SPI1 |
| TOX |
| CBFB |
| RELB |
| GBA |
| GATA3 |
| TBX21 |
| SPNS2 |
| WNK1 |
| FUT7 |
| ARTN |
| EXT1 |
| CRTAM |
| IMPDH2 |
| HELLS |
| HPRT1 |
| PURA |
| TCIRG1 |
| F11R |
| ITGAL |
| DOCK8 |
| ARG1 |
| RC3H1 |
| SCRIB |
| PRNP |
| IDO1 |
| PDCD1LG2 |

| |
|---|
| CD274 |
| LRRC32 |
| PRKAR1A |
| CD160 |
| TNFRSF14 |
| TRIM27 |
| SHH |
| IHH |
| GLI3 |
| CBLB |
| TYRO3 |
| AXL |
| MERTK |
| PADI2 |
| FBXO7 |
| GCSAM |
| MIA3 |
| ADTRP |
| AKT1 |
| WASL |
| LST1 |
| IL2 |
| SOX11 |
| FOXP3 |
| FGL2 |
| IFNL1 |
| PLA2G2D |
| TIGIT |
| ILDR2 |
| VTCN1 |
| PTPN22 |
| GPNMB |
| LAX1 |
| CTSG |
| PAG1 |
| PELI1 |
| LAG3 |
| SOCS6 |
| TNFAIP8L2 |
| IL4I1 |
| TGFB1 |
| HAVCR2 |
| HFE |
| SMAD7 |
| CD74 |

| |
|---|
| DTX1 |
| NRARP |
| IL27RA |
| PTPRC |
| IL7R |
| KLRD1 |
| PPP3CB |
| NOD2 |
| CLEC4G |
| APOD |
| LRCH1 |
| IL20RB |
| GLMN |
| CD86 |
| CD80 |
| DLG5 |
| DLG1 |
| MAD1L1 |
| CEBPB |
| SDC4 |
| NDFIP1 |
| PTPN6 |
| VSIG4 |
| FOXJ1 |
| PAWR |
| MARCHF7 |
| HHLA2 |
| AGER |
| IGF2 |
| IL12B |
| GPAM |
| IGF1 |
| STAT5B |
| RPS3 |
| IL12RB1 |
| IL18 |
| PYCARD |
| IL23A |
| FADD |
| IGFBP2 |
| TMIGD2 |
| CD81 |
| SKAP1 |
| SASH3 |
| SIRT1 |

| |
|---|
| ITPKB |
| ADA |
| SYK |
| ZAP70 |
| NKAP |
| CCR2 |
| CD28 |
| CLEC7A |
| INPP5D |
| NCKAP1L |
| JAM2 |
| MADCAM1 |
| MYD88 |
| FGF10 |
| IL12A |
| ZNF335 |
| TLR4 |
| CYRIB |
| CD46 |
| MAMU-DRA |
| PCK1 |
| LCK |
| CD47 |
| PRKCQ |
| THY1 |
| SIRPG |
| FCHO1 |
| MALT1 |
| TNFSF11 |
| CCDC88B |
| CORO1A |
| PDCD1 |
| LGALS14 |
| LGALS13 |
| WNT5A |
| OXSR1 |
| ADAM10 |
| ADAM17 |
| TMEM102 |
| STK39 |
| TNFSF14 |
| CCR7 |
| MAP3K7 |
| B2M |
| TRAF6 |

| |
|---|
| FZD5 |
| TRAF2 |
| RHOH |
| ZMIZ1 |
| XBP1 |
| TAPBPL |
| IL7 |
| IL4 |
| ZBTB1 |
| MDK |
| IL1RL2 |
| PIK3R6 |
| BAD |
| VNN1 |
| EGR3 |
| RASGRP1 |
| CD99L2 |
| CD1D |
| STX7 |
| CD1E |
| MR1 |
| SLC22A13 |
| FBXO38 |
| AIF1 |
| APP |
| CCL21 |
| ITGB3 |
| CXCL10 |
| NCK1 |
| ZP4 |
| VCAM1 |
| SPTA1 |
| IL6 |
| HES1 |
| IL15 |
| TFRC |
| JAK2 |
| NCK2 |
| IL21 |
| CD6 |
| DNAJA3 |
| CD40LG |
| JAK3 |
| TNFSF13B |
| EFNB1 |

| |
|---|
| LEP |
| SLC7A1 |
| CD209 |
| IL1B |
| IL6ST |
| TYK2 |
| DHPS |
| BACH2 |
| DUSP10 |
| CD48 |
| ADCY7 |
| IRF7 |
| GSN |
| SLC8B1 |
| IKZF3 |
| RNF41 |
| GCSAML |
| STK10 |
| MSN |
| BCL6 |
| SPN |
| RAB5C |
| SIT1 |
| WAS |
| TRPM4 |
| CYP26B1 |
| SLC46A2 |
| SOS2 |
| SOS1 |
| CLPTM1 |
| FOXN1 |
| LMO1 |
| CRK |
| ECM1 |
| CTNNB1 |
| LGALS3 |
| TNFSF18 |
| RAC2 |
| CTLA4 |
| IL27 |
| NECTIN2 |
| CD2 |
| SLAMF6 |
| TRAF3IP2 |
| GJA1 |

| |
|---|
| TREML2 |
| AZI2 |
| CD8A |
| CD44 |
| SLA2 |
| SMAD3 |
| NLRC3 |
| VAV1 |
| DPP4 |
| CD1C |
| F2RL1 |
| PSEN1 |
| LCP1 |
| APBB1IP |
| ICAM1 |
| SLC12A2 |
| PLEC |
| CXCL11 |
| CD5 |
| EFNB3 |
| EFNB2 |
| ICOS |
| LEPR |
| RUNX2 |
| KIT |
| PKNOX1 |
| JAG2 |
| LFNG |
| RAG2 |
| KDELR1 |
| SP3 |
| CHD7 |
| PTPN2 |
| SOX4 |
| LMBR1L |
| NHEJ1 |
| EGR1 |
| PREX1 |
| BCL2 |
| PATZ1 |
| DLL4 |
| WNT4 |
| RABL3 |
| NCAPH2 |
| PSMB11 |

| |
|---|
| ZFP36L2 |
| TP53 |
| WNT1 |
| MAFB |
| LIG4 |
| FZD7 |
| JMJD6 |
| ZFP36L1 |
| SEMA4A |
| FCER1G |
| MYO1G |
| CTSC |
| EMP2 |
| PRF1 |
| GZMM |
| BTN3A3 |
| JAG1 |
| S1PR1 |
| GPR15 |
| ITGB7 |
| CTPS1 |
| NCSTN |
| CCND3 |
| CD151 |
| FKBP1B |
| PSMB10 |
| RC3H2 |
| ARMC5 |
| SLC11A1 |

| Column1 |
| --- |
| CCR7 |
| OXSR1 |
| WNK1 |
| STK39 |
| ACKR4 |
| CCR9 |
| XCR1 |
| CCR1 |
| CCR3 |
| CCR2 |
| CCRL2 |
| CCR6 |
| ACKR1 |
| CCR4 |
| CCL26 |
| CCR8 |
| FOXC1 |
| CXCL12 |
| ACKR3 |
| CXCR2 |
| PTK2B |
| CCL7 |
| GPR75 |
| CCL16 |
| CCL23 |
| CCL18 |
| CCL20 |
| GPR35 |
| CX3CL1 |
| CCL17 |
| CXCL11 |
| CXCL10 |
| CCR10 |
| CXCR4 |
| CXCL1 |
| CXCR5 |
| CXCR3 |
| TREM2 |
| WBP1L |
| EDN2 |
| IL5RA |
| GREM2 |
| CSF3R |
| IL17RB |

| |
|---|
| LEPR |
| IL6R |
| IRF5 |
| IL1RAP |
| TRAF3IP2 |
| IL20RA |
| IFNGR1 |
| KIT |
| SH2B2 |
| IL6 |
| GHR |
| IFNGR2 |
| IFNAR1 |
| IL2RB |
| DUOX1 |
| CSF2RB |
| CSF1R |
| JAK2 |
| IL7 |
| IFNK |
| SLC1A1 |
| CD44 |
| IL7R |
| FER |
| TNFRSF1A |
| SOCS1 |
| IL13RA2 |
| IL13RA1 |
| NUMBL |
| CNTFR |
| STAT4 |
| SOCS5 |
| IFNB1 |
| JAK1 |
| ZC3H15 |
| STAT5B |
| JAK3 |
| IL17RE |
| IL4R |
| IL12RB1 |
| IL21R |
| RELA |
| IL1RAPL2 |
| IL1R2 |
| LIFR |

| |
|---|
| IL1R1 |
| IRAK3 |
| STAT3 |
| STAT2 |
| EPOR |
| IL1A |
| IL1B |
| IL37 |
| IRAK4 |
| IL6ST |
| TYK2 |
| PRLR |
| EREG |
| PF4 |
| STAT6 |
| TNFSF11 |
| BAD |
| IRF6 |
| MAMU-A |
| IRF9 |
| IRF4 |
| IRF1 |
| TP53 |
| IFNG |
| STAT1 |
| NMI |
| IRF8 |
| IRF7 |
| PSMB2 |
| PSMA5 |
| MYD88 |
| PSMB1 |
| PSMA6 |
| PSMB9 |
| PSMB8 |
| IKBKB |
| PLCB1 |
| PSMB5 |
| RPS6KA5 |
| TRAF6 |
| PSMA7 |
| PSMA1 |
| PSMB7 |
| PSMB4 |
| PSMB3 |

| |
|---|
| EGR1 |
| IRAK1 |
| PSMA8 |
| IRAK2 |
| PSMA4 |
| RPS6KA4 |
| MAPK3 |
| IL15 |
| IL15RA |
| CD4 |
| TRAF5 |
| IL17F |
| IL17A |
| IKBKE |
| IL17RA |
| NOTCH1 |
| TRAF2 |
| AKT1 |
| PDGFB |
| IL18 |
| IL18R1 |
| IL18RAP |
| IL2RA |
| MX1 |
| OAS1 |
| OASL |
| OAS2 |
| SYK |
| IL33 |
| IL34 |
| SPI1 |
| SRC |
| CTR9 |
| YAP1 |
| SMAD4 |
| CHMP4C |
| ZFYVE19 |
| SLIT2 |
| PADI2 |
| ROBO1 |
| SLIT3 |
| RNF113A |
| SH2B3 |
| PTPRC |
| ECM1 |

| |
|---|
| IL36RN |
| ARG1 |
| OTOP1 |
| PTPN2 |
| PARP14 |
| OTUD4 |
| IL1RN |
| ADIPOQ |
| TRAIP |
| HIST1H2BJ |
| GPS2 |
| APOA1 |
| XIAP |
| NAIP |
| NR1H4 |
| F2RL1 |
| PELI3 |
| RFFL |
| CCDC3 |
| NOL3 |
| PPP2CB |
| GAS6 |
| PYDC1 |
| PIAS4 |
| MUL1 |
| DCST1 |
| YTHDF2 |
| ADAR |
| ISG15 |
| METTL3 |
| SAMHD1 |
| YTHDF3 |
| OAS3 |
| MMP12 |
| TTLL12 |
| NLRC5 |
| CACTIN |
| HIF1A |
| EDN1 |
| RIPK2 |
| CD74 |
| TRIM44 |
| AXL |
| PAFAH1B1 |
| HPX |

| |
|---|
| MED1 |
| PARP9 |
| TRIM32 |
| GFI1 |
| RIPK1 |
| C1QTNF4 |
| UBE2K |
| ADAM17 |
| PRKN |
| CASP4 |
| WNT5A |
| TRIM56 |
| ZBP1 |
| TRIM6 |
| LSM14A |
| TRIM41 |
| MAVS |
| STING1 |
| TBK1 |
| IRF3 |
| FADD |
| USP27X |
| HSP90AB1 |
| VRK2 |
| NKIRAS1 |
| HIPK1 |
| SPATA2 |
| OTULIN |
| CYLD |
| SHARPIN |
| NKIRAS2 |
| PYCARD |
| SPHK1 |
| TNFRSF1B |
| TNF |
| AIM2 |
| NFKBIA |
| TNFSF18 |
| FOXO3 |
| RRAGA |
| TRAF3 |
| ACTN4 |
| ILK |
| CARD14 |
| CDIP1 |

| |
|---|
| TNFRSF17 |
| EDA2R |
| TNFSF13B |
| PLVAP |
| UMOD |
| TXNDC17 |
| TNFRSF11A |
| TNFRSF13C |
| CD70 |
| IRF2 |
| IFI27 |
| IFNAR2 |
| FFAR2 |
| MBL2 |
| FCN1 |
| TIFA |
| ALPK1 |
| MYO1G |
| CSF2 |
| CDC42 |
| ITGA10 |
| ITGA9 |
| ADAM15 |
| ITGB2 |
| ZYX |
| CCN2 |
| FGR |
| PTN |
| FYB2 |
| ADAMTS4 |
| ITGB8 |
| RCC2 |
| ADAMTS1 |
| CDH17 |
| ADAM10 |
| ITGB1 |
| NRP1 |
| SEMA7A |
| FERMT1 |
| HPS5 |
| MPIG6B |
| ITGB1BP1 |
| THY1 |
| CIB1 |
| ITGA11 |

| |
|---|
| ITGB4 |
| FERMT2 |
| FN1 |
| ITGA2B |
| ITGA5 |
| BCAR1 |
| ITGA2 |
| ITGB5 |
| ILKAP |
| ITGA7 |
| TEC |
| CUL3 |
| FUT8 |
| CD40LG |
| ITGAV |
| MADCAM1 |
| ITGB3 |
| PTPRA |
| COL3A1 |
| ZNF304 |
| TLN1 |
| PLEK |
| CIB2 |
| FYB1 |
| ITGAE |
| PTPN11 |
| ITGB7 |
| PRAM1 |
| ADAMTS10 |
| ABL1 |
| FERMT3 |
| VAV1 |
| ITGB6 |
| ITGA6 |
| BCL10 |
| PTAFR |
| MTDH |
| CD6 |
| MAPK1 |
| LBP |
| LY96 |
| TICAM1 |
| PRKCE |
| MAPK14 |
| TGFB1 |

| |
|---|
| LYN |
| MALT1 |
| PLCG2 |
| TLR4 |
| CD14 |
| CSF1 |
| DOK1 |
| TLR5 |
| TLR10 |
| TLR1 |
| TLR6 |
| TLR2 |
| TNIP1 |
| TLR9 |
| TLR8 |
| TLR3 |
| TNIP3 |
| MKKS |
| BBS4 |
| APPL1 |
| HAVCR2 |
| PHACTR4 |
| CTNNA1 |
| SLC2A10 |
| LTF |
| DEFB114 |
| NFKBIL1 |
| TRIB1 |
| LACRT |
| STAP1 |
| TKFC |
| RIOK3 |
| DHX58 |
| C1QBP |
| CD300A |
| CD300LF |
| SARM1 |
| GPATCH3 |
| UFD1 |
| NLRX1 |
| NPLOC4 |
| SEC14L1 |
| RNF125 |
| MFHAS1 |
| NOD2 |

| |
|---|
| TNFAIP3 |
| RAB7B |
| NR1D1 |
| DAB2IP |
| BPIFB1 |
| GRAMD4 |
| TYRO3 |
| ARRB2 |
| LGR4 |
| MFSD3 |
| NLRP6 |
| NOD1 |
| INAVA |
| LACC1 |
| CLEC4E |
| APPL2 |
| DMTN |
| LIMS1 |
| LIMS2 |
| LOXL3 |
| EMP2 |
| CD63 |
| FLNA |
| BMP6 |
| LY86 |
| SASH1 |
| SCIMP |
| CD180 |
| PRKCA |
| ANKRD17 |
| DDX60 |
| ZDHHC5 |
| PUM1 |
| TRIM15 |
| PUM2 |
| ZCCHC3 |
| USP15 |
| TIRAP |
| PJA2 |
| CYBA |
| PTPN22 |
| FLOT1 |
| PELI1 |
| WDFY1 |
| NR1H3 |

| |
|---|
| IFI35 |
| RSAD2 |
| SLC15A4 |
| TASL |
| DDX3X |
| RTN4 |
| TIMP1 |
| ANGPT1 |
| NOP53 |
| FCRL3 |
| CD36 |
| ESR1 |
| PHB2 |
| RNF135 |
| CLPB |
| PHB |
| NKG2D |
| KLRD1 |
| CLEC7A |
| CLEC6A |
| CLDN18 |
| EXT1 |
| PIK3AP1 |
| RFTN1 |
| UNC93B1 |
| COLEC12 |
| S100A14 |
| TRIL |
| TLR7 |
| EPG5 |
| MAPKAPK3 |
| LRRC19 |
| MAPKAPK2 |
| RPS6KA3 |
| AP3B1 |
| RAB11FIP2 |
| CD40 |

| Column1 |
| --- |
| DYNLT1 |
| IFIT1 |
| APCS |
| PTX3 |
| KPNA7 |
| NFKBIA |
| BRAP |
| KPNA3 |
| NUP58 |
| KPNA1 |
| KPNA2 |
| TNPO2 |
| AKAP5 |
| CCR7 |
| PCNT |
| TPR |
| PDCD10 |
| PCM1 |
| ZFAND1 |
| PRR5L |
| ICE1 |
| OAZ3 |
| PLK3 |
| RIPOR1 |
| CEP290 |
| CEP131 |
| KIF20B |
| B3GAT3 |
| ERGIC3 |
| OAZ2 |
| OAZ1 |
| MCOLN2 |
| TRPV4 |
| SEC16B |
| SLC35D3 |
| TMEM30A |
| TMEM30B |
| CD81 |
| SORL1 |
| SAR1B |
| SAR1A |
| TM9SF4 |
| SLC51B |
| GSK3B |

| |
|---|
| CTDSPL2 |
| SFN |
| BAG3 |
| PPM1A |
| RBM22 |
| YWHAE |
| RAPGEF3 |
| TCF7L2 |
| CAMK1 |
| PRKACA |
| GAS6 |
| EMD |
| XPO4 |
| ZC3H12A |
| PRKCD |
| ECT2 |
| HCLS1 |
| TARDBP |
| SMO |
| PTPN22 |
| HYAL2 |
| PRKD1 |
| SHH |
| HDAC3 |
| ZPR1 |
| TRIM28 |
| JUP |
| IFNG |
| CHP2 |
| CDH1 |
| MAVS |
| PSEN1 |
| MAPK14 |
| TGFB1 |
| UBR5 |
| IPO5 |
| DMAP1 |
| LEP |
| FLNA |
| PIK3R1 |
| GLI3 |
| YWHAG |
| YWHAQ |
| YWHAB |
| YWHAZ |

| |
|---|
| PDCD5 |
| TCAF1 |
| FYN |
| FIS1 |
| MIEF1 |
| CEMIP |
| ERBB2 |
| KCNB1 |
| CIB1 |
| AKT2 |
| CDK5 |
| MYO1C |
| MIEF2 |
| CDK5R1 |
| PRNP |
| C2CD5 |
| MFF |
| HRAS |
| TOMM7 |
| FBXW7 |
| HSPA1L |
| PRKAA1 |
| EDEM1 |
| BCAP31 |
| EDEM2 |

| Column1 |
|---|
| LYST |
| RNF19B |
| CD2 |
| FCGR3 |
| PLEKHM2 |
| KIF5B |
| GZMB |
| ULBP3 |
| PTPN6 |
| NKG2D |
| PRDX1 |
| IL18 |
| CEBPG |
| HLA-E |
| ARRB2 |
| KLRD1 |
| CLEC12B |
| HAVCR2 |
| CD160 |
| SLAMF6 |
| NCR3 |
| IL12A |
| IL21 |
| LAG3 |
| SH2D1A |
| RASGRP1 |
| CD226 |
| AP1G1 |
| CRTAM |
| STAT5B |
| NECTIN2 |
| IL18RAP |
| CADM1 |
| VAV1 |
| IL12B |
| SERPINB9 |
| LEP |
| PIK3R6 |
| CD96 |
| CLNK |
| BAG6 |
| IL18R1 |
| CD244 |
| IFNB1 |

| |
|---|
| UNC13D |
| VAMP7 |
| CORO1A |
| PBX1 |
| RABL3 |
| TUSC2 |
| PTPRC |
| HECTD1 |
| TYRO3 |
| NFIL3 |
| TOX |
| SP3 |
| AXL |
| KAT7 |
| MERTK |
| ELF4 |
| FGR |
| RHBDD3 |
| PIBF1 |
| PGLYRP3 |
| PGLYRP2 |
| BLOC1S6 |
| TICAM1 |
| BLOC1S3 |
| CXCL14 |
| CCL7 |
| IL15 |
| ZBTB1 |
| GAS6 |
| JAK2 |
| IL23A |
| TYK2 |
| DCAF15 |
| PRDM1 |

| Column1 |
|---|
| SPI1 |
| STAP1 |
| CENPC |
| CSF2 |
| CSF2RB |
| JAK2 |
| CSF1 |
| CSF1R |
| DOK1 |
| CD74 |
| TREM2 |
| PTPN2 |
| CXCR4 |
| ANGPT1 |
| WNT5A |
| CD36 |
| SEMA7A |
| PLCG2 |
| SIRT1 |
| TLR4 |
| MCOLN2 |
| TRPV4 |
| SENP7 |
| MX1 |
| PHB2 |
| RNF135 |
| CLPB |
| OAS1 |
| ZDHHC1 |
| PHB |
| AKAP1 |
| NLRP1 |
| IFIT1 |
| CXCL10 |
| SFPQ |
| TOMM70 |
| PYHIN1 |
| AIM2 |
| IFI16 |
| CGAS |
| PRKDC |
| ZNFX1 |
| SIN3A |
| TLR9 |

| |
|---|
| RBM14 |
| CASP6 |
| MAVS |
| HEXIM1 |
| STING1 |
| TBK1 |
| ZCCHC3 |
| PYCARD |
| XRCC6 |
| PQBP1 |
| XRCC5 |
| PSPC1 |
| BCL10 |
| DEFB114 |
| CX3CR1 |
| CLEC7A |
| RARRES2 |
| CARD9 |
| CLEC4E |
| CLEC4D |
| CLEC6A |
| ETV3 |
| AKT1 |
| CD4 |
| ZFP36L2 |
| MAPK1 |
| PDE1B |
| ZFP36 |
| NPR2 |
| FER |
| DCSTAMP |
| TRIM58 |
| S100A8 |
| TTC4 |
| TRIM11 |
| TRIM45 |
| TLR5 |
| TRIM46 |
| MASP2 |
| ADAM15 |
| DEFB112 |
| DEFB110 |
| DEFB113 |
| ANKRD17 |
| IRF5 |

| |
|---|
| TLR10 |
| TLR1 |
| TLR6 |
| IFI6 |
| FGR |
| VNN1 |
| MYD88 |
| FYN |
| APCS |
| ARHGEF2 |
| CAMP |
| FCER1G |
| TRIM56 |
| CLEC5A |
| JCHAIN |
| TRIM50 |
| TRIM33 |
| TRIM26 |
| TRIM15 |
| TRIM10 |
| TRIM40 |
| FBXO9 |
| TLR3 |
| C7 |
| LGALS3 |
| C6 |
| DEFB128 |
| DEFB108B |
| IFI27 |
| DEFB134 |
| RIPK2 |
| DEFB135 |
| COLEC10 |
| TRIM36 |
| BLK |
| IGHE |
| IGHM |
| DEFB125 |
| DEFB115 |
| DEFB116 |
| PTX3 |
| DEFB118 |
| SFTPD |
| RNASE6 |
| HMGB2 |

| |
|---|
| DEFB119 |
| INAVA |
| DEFB121 |
| RNASE7 |
| APOBEC3G |
| DEFB122 |
| DEFB123 |
| PML |
| POLR3A |
| TRIM59 |
| RPP21 |
| TMEM131L |
| REL |
| TRIM63 |
| TLR2 |
| TRIM54 |
| DDX58 |
| TRIM22 |
| SERINC3 |
| TRIM5 |
| TIRAP |
| TRIM27 |
| TRIM6 |
| DAPK1 |
| SLPI |
| TRAF3 |
| TRIM60 |
| TOLLIP |
| TRIM68 |
| TRIM21 |
| OTULIN |
| SYK |
| SAMHD1 |
| SH2D1A |
| TRIM28 |
| STYK1 |
| TRIM14 |
| TRIM55 |
| C1S |
| SKP2 |
| PJA2 |
| C1R |
| G3BP1 |
| KRT16 |
| CYBB |

| |
|---|
| MFHAS1 |
| TRIM35 |
| LBP |
| ELF4 |
| NOD2 |
| RIOK3 |
| LY96 |
| CYLD |
| TRIM65 |
| OASL |
| TIFA |
| ALPK1 |
| OAS3 |
| OAS2 |
| TICAM1 |
| TRIM3 |
| DHX58 |
| IPO7 |
| SERINC5 |
| IFI35 |
| CYBA |
| CYBC1 |
| NMI |
| HMGB1 |
| NLRC5 |
| LYN |
| SLA |
| MALT1 |
| C5 |
| COLEC12 |
| TRIM8 |
| IRAK1 |
| DEFB132 |
| MBL2 |
| MID1 |
| BST2 |
| PYDC1 |
| TRIM13 |
| C9 |
| MID2 |
| TLR7 |
| TLR8 |
| TRIM32 |
| FADD |
| IRAK4 |

| |
|---|
| CD14 |
| DDX3X |
| G3BP2 |
| PGLYRP1 |
| NLRC4 |
| BPIFB3 |
| C3 |
| BPIFA1 |
| KCNH7 |
| LTF |
| DEFB1 |
| DEFA1B |
| DEFA4 |
| HIST1H2BJ |
| IL4 |
| APOA4 |
| PLA2G1B |
| FAU |
| BPIFB1 |
| EDN2 |
| PLA2G3 |
| IL13 |
| CRTC3 |
| PLA2G10 |
| SLC11A1 |
| TMEM106A |
| JMJD6 |
| SUCNR1 |
| DYSF |
| IFNG |
| PRKCE |
| IL33 |
| SBNO2 |
| HAVCR2 |
| TAFA4 |
| EDNRB |
| NUP85 |
| IL34 |
| CCR2 |
| JUN |
| SNCA |
| TNF |
| IFNGR1 |
| AGER |
| APP |

| |
|---|
| C5AR1 |
| CLU |
| AZU1 |
| GBA |
| CX3CL1 |
| HYAL2 |
| ADAM10 |
| ADAM9 |
| CCL26 |
| PDGFB |
| CTSG |
| CALCA |
| CCL7 |
| CCL16 |
| CCL23 |
| CCL18 |
| PTPRO |
| CCL20 |
| CCL17 |
| FLT1 |
| MSMP |
| ANXA1 |
| TNFSF11 |
| JAML |
| PECAM1 |
| TGFB1 |
| USP15 |
| FAM3A |
| MUL1 |
| TNFAIP3 |
| TREX1 |
| LYAR |
| ATG12 |
| TYRO3 |
| NLRX1 |
| ATG5 |
| TRAFD1 |
| IRAK3 |
| DRD2 |
| CACTIN |
| CD200 |
| ADGRF5 |
| LRFN5 |
| NR1H3 |
| VSIG4 |

| |
|---|
| MMP28 |
| EMILIN1 |
| SYT11 |
| NR1D1 |
| CST7 |
| LDLR |
| FN1 |
| CD33 |
| CCN3 |
| DUSP1 |
| PLCB1 |
| SLIT2 |
| CRP |
| ATG3 |
| ADIPOQ |
| PIP4P2 |
| SNX3 |
| RACK1 |
| CNN2 |
| PRTN3 |
| CD300A |
| NCF2 |
| LEPR |
| ITGB2 |
| EIF2AK1 |
| C1H1orf43 |
| ELMO1 |
| ANXA3 |
| PLD4 |
| TUSC2 |
| RAB11FIP2 |
| MET |
| MESD |
| NCF4 |
| CEBPE |
| CDC42SE2 |
| UNC13D |
| TM9SF4 |
| ELMO2 |
| AXL |
| ANXA11 |
| CEACAM4 |
| IRF8 |
| CDC42SE1 |
| MERTK |

| |
|---|
| GAS6 |
| LEP |
| ELANE |
| MYO7A |
| MYO1G |
| ICAM5 |
| ABL1 |
| LRP1 |
| CORO1A |
| VAV1 |
| CDC42 |
| AIF1 |
| ARHGAP12 |
| MSR1 |
| MYH9 |
| ADGRB1 |
| ABCA1 |
| MFGE8 |
| CLCN3 |
| BIN2 |
| GSN |
| GULP1 |
| ARHGAP25 |
| TULP1 |
| TUB |
| SRPX |
| CLN3 |
| RAB7A |
| TMEM175 |
| RAB7B |
| RAB39A |
| PIKFYVE |
| RAB20 |
| RAB34 |
| ISL1 |
| RASGRP1 |
| IL12B |
| IL18 |
| IL23A |
| PAEP |
| IL1B |
| POLR3C |
| PLSCR2 |
| POLR3D |
| ADAM8 |

| |
|---|
| COCH |
| SLC15A4 |
| POLR3G |
| FPR2 |
| TASL |
| POLR3F |
| POLR3B |
| EREG |
| PLA2G4A |
| THBS1 |
| KARS1 |
| HAMP |
| IL4R |
| IL1RL1 |
| AKIRIN1 |
| CMKLR1 |
| MSTN |
| CXCL17 |
| MDK |
| C3AR1 |
| MAPK3 |
| MMP14 |
| TTBK1 |
| CTSC |
| MMP8 |
| HAS2 |
| NR4A3 |
| CD44 |
| S100A14 |
| CCR1 |
| PLA2G7 |
| SERPINE1 |
| LGMN |
| CCL1 |
| MOSPD2 |
| CREB3 |
| ANO6 |
| IL12A |
| AHSG |
| CD47 |
| SOD1 |
| APOA2 |
| IL15 |
| IL2RB |
| IL15RA |

| |
|---|
| APOA1 |
| LMAN2 |
| DOCK2 |
| CAMK1D |
| CALR |
| F2RL1 |
| NCKAP1L |
| APPL2 |
| LRP8 |
| DUSP10 |
| APPL1 |
| TKFC |
| N4BP1 |
| NFE2L2 |
| APOE |
| RORA |
| MYO18A |
| MST1 |
| PTK2B |
| CD81 |
| CD9 |
| SPHK1 |
| P2RY12 |
| PTPRC |
| HCK |
| ITGAV |
| PRKCG |
| LETMD1 |

| Column1 |
| --- |
| E2F8 |
| SHCBP1L |
| MSX1 |
| STRA8 |
| PLCB1 |
| NUF2 |
| MAD1L1 |
| CENPC |
| KIF2C |
| NDC80 |
| CENPE |
| CDT1 |
| MIS12 |
| MAPRE1 |
| CHAMP1 |
| CCNE2 |
| ATM |
| MEIOC |
| CCNE1 |
| ATRX |
| RAD51 |
| MAD2L1BP |
| HSF2BP |
| SPO11 |
| BRIP1 |
| BRME1 |
| FANCD2 |
| UBA3 |
| CIB1 |
| MYH9 |
| SPIRE1 |
| SPIRE2 |
| MOS |
| NAA50 |
| NIPBL |
| CLASP2 |
| KPNB1 |
| CLASP1 |
| NUSAP1 |
| GPSM2 |
| PKHD1 |
| GJA1 |
| HTT |
| MCPH1 |

| |
|---|
| FGF10 |
| ITGB1 |
| PAX6 |
| CENPA |
| SPRY1 |
| INSC |
| UBXN2B |
| SPDL1 |
| NUMA1 |
| PLK1 |
| SPRY2 |
| DCTN1 |
| NDE1 |
| NSFL1C |
| NDEL1 |
| PAFAH1B1 |
| KAT5 |
| CDK5RAP2 |
| MISP |
| ZW10 |
| CHMP7 |
| CHMP2A |
| CHMP4B |
| EPS8 |
| CTDP1 |
| UBE2S |
| SPAST |
| MLH1 |
| TTK |
| ACSL6 |
| NCAPH2 |
| TRIP13 |
| CDC25B |
| MEIOB |
| MLH3 |
| RAD51C |
| WEE2 |
| TOP2A |
| SYCP2 |
| MARF1 |
| IQGAP3 |
| TRIM71 |
| E2F3 |
| EIF4E |
| GPR132 |

| |
|---|
| CACUL1 |
| LATS1 |
| HINFP |
| MYC |
| ZNF324 |
| PLK2 |
| SKP2 |
| TFDP3 |
| EIF4EBP1 |
| TAF10 |
| PPP3CA |
| USP37 |
| RBBP8 |
| RPS6KB1 |
| E2F1 |
| PLK3 |
| CDKN1B |
| POLE |
| SPDYA |
| RB1 |
| ACVR1B |
| CDK4 |
| PIM2 |
| LATS2 |
| PHF8 |
| CCND1 |
| NES |
| TPD52L1 |
| CCNA2 |
| CDK14 |
| ARPP19 |
| TAF2 |
| CCNY |
| CHEK2 |
| USH1C |
| BRSK2 |
| CALM3 |
| FBXL15 |
| FBXL7 |
| PPM1D |
| MASTL |
| CDK1 |
| WNT10B |
| CDC25C |
| FBXL6 |

| |
|---|
| KDM8 |
| FBXL12 |
| KAT14 |
| FOXM1 |
| BRSK1 |
| FBXL3 |
| CCNB2 |
| YTHDC2 |
| OVOL1 |
| NDC1 |
| MAEL |
| SYCP1 |
| MEI4 |
| IHO1 |
| REC8 |
| SUN1 |
| MSH4 |
| TERB2 |
| STAG3 |
| C7H14orf39 |
| TERB1 |
| KASH5 |
| RAD21L1 |
| TEX15 |
| HORMAD1 |
| SPATA22 |
| MRE11 |
| MAJIN |
| SIRT7 |
| DSCC1 |
| MAU2 |
| TEX11 |
| M1AP |
| BRDT |
| FOXJ3 |
| UBR2 |
| SLC25A31 |
| TDRD9 |
| ING2 |
| BRCA2 |
| AGO4 |
| ASZ1 |
| FIGNL1 |
| TESMIN |
| KIF18A |

| |
|---|
| TDRKH |
| CYP26B1 |
| TEX14 |
| TAF1L |
| FANCA |
| CKS2 |
| PSMD13 |
| ESPL1 |
| FKBP6 |
| NBN |
| TUBGCP6 |
| AURKA |
| HORMAD2 |
| PDIK1L |
| SMC3 |
| BOLL |
| STK35 |
| SMC1A |
| ZNF805 |
| PSMA8 |
| SMC1B |
| ACTR3 |
| FAM9C |
| ACTR2 |
| NSUN2 |
| EDNRA |
| EDN1 |
| SMC4 |
| SMC2 |
| NCAPD3 |
| SGO1 |
| C14H11orf80 |
| SLX4 |
| TEX12 |
| ERCC1 |
| RAD1 |
| SGO2 |
| BUB1B |
| ASPM |
| WASHC5 |
| CHFR |
| INCENP |
| TUBG1 |
| TERF1 |
| UBE2B |

| |
|---|
| TACC3 |
| CDC23 |
| ANAPC1 |
| CDC27 |
| CDC16 |
| SETD2 |
| CENPF |
| NEK4 |
| RNF2 |
| PHF13 |
| CENPW |
| TUBB2B |
| DYNC1H1 |
| USP16 |
| GEM |
| AZI2 |
| NCAPD2 |
| ABRAXAS2 |
| CLTCL1 |
| DIS3L2 |
| CDC6 |
| TUBA8 |
| USP3 |
| RGS14 |
| RECQL5 |
| ZNF830 |
| SNX9 |
| SNX33 |
| TUBA3C |
| WAPL |
| KIF18B |
| E2F4 |
| TUBA3D |
| WEE1 |
| CUL3 |
| NUDT15 |
| CLTC |
| TADA3 |
| SNX18 |
| TUBA1C |
| SKA2 |
| XRCC2 |
| CENPT |
| PRDM5 |
| CETN1 |

| |
|---|
| TADA2A |
| TBCD |
| CDCA5 |
| SKA1 |
| PPP1R12A |
| TUBA4A |
| TTYH1 |
| SKA3 |
| IK |
| TUBB4A |
| ZWILCH |
| ZWINT |
| CCND3 |
| LZTS1 |
| CCNG1 |
| CCNB1 |
| TIGAR |
| CCNI2 |
| CCNA1 |
| CCNI |
| CCNJ |
| KIF11 |
| NUP62 |
| NCAPG |
| NCAPH |
| AKAP8 |
| AKAP8L |
| KATNB1 |
| RHOA |
| ARF1 |
| SETMAR |
| CLSPN |
| TOPBP1 |
| TICRR |
| NAE1 |
| MCM4 |
| POLA1 |
| CDC45 |
| RTF2 |
| RPA2 |
| SDE2 |
| WAC |
| PRKDC |
| GIGYF2 |
| RPS27L |

| |
|---|
| TP53 |
| FBXO31 |
| RFWD3 |
| DTL |
| SYF2 |
| ABRAXAS1 |
| DONSON |
| RINT1 |
| FOXN3 |
| RPP21 |
| TAOK1 |
| CDK5RAP3 |
| UIMC1 |
| CDC14B |
| TAOK2 |
| BLM |
| NOP53 |
| CDKN1A |
| BRCA1 |
| BABAM2 |
| MRNIP |
| BABAM1 |
| HMGA2 |
| TAOK3 |
| FZR1 |
| FOXO4 |
| BRCC3 |
| INTS3 |
| INIP |
| NABP1 |
| NABP2 |
| TIPIN |
| RAD17 |
| ATF2 |
| MSH2 |
| RAD9A |
| CDCA8 |
| PSRC1 |
| CHMP2B |
| KIFC1 |
| KIF14 |
| NUDC |
| BOD1 |
| CHMP4C |
| RAB11A |

| |
|---|
| PINX1 |
| RRS1 |
| KIF22 |
| EML4 |
| DCTN2 |
| CHMP6 |
| CHMP5 |
| ANKRD53 |
| VPS4B |
| PIBF1 |
| EML3 |
| BECN1 |
| KLHDC8B |
| RNF103 |
| SMPD3 |
| REEP4 |
| ANKLE2 |
| CDC20 |
| PDS5A |
| POGZ |
| PTTG1 |
| HASPIN |
| PDS5B |
| NSL1 |
| NEK2 |
| NCAPG2 |
| DSN1 |
| ZNF207 |
| CENPK |
| SPAG5 |
| BUB3 |
| KNSTRN |
| INO80 |
| STAG1 |
| SPICE1 |
| TASOR |
| MAP9 |
| TPX2 |
| STAG2 |
| LSM14A |
| ARHGEF10 |
| KIF2A |
| CEP192 |
| KIF3B |
| GOLGA2 |

| |
|---|
| TOGARAM2 |
| KIFC2 |
| FLNA |
| MYBL2 |
| CCDC61 |
| AAAS |
| TPR |
| APC |
| KNTC1 |
| AURKB |
| BUB1 |
| VCP |
| KIF4B |
| KIF4A |
| RACGAP1 |
| PCNT |
| STIL |
| DLGAP5 |
| EFHC1 |
| PTPA |
| CENPH |
| CKAP5 |
| EFHC2 |
| CEP126 |
| WDR62 |
| TACC1 |
| EML1 |
| INTS13 |
| TACC2 |
| DCTN6 |
| MECP2 |
| SPC25 |
| RTEL1 |
| PCNA |
| DNA2 |
| RGCC |
| CTDSPL |
| CDC73 |
| GPNMB |
| ZNF655 |
| DACT1 |
| PKD2 |
| JADE1 |
| DLG1 |
| KLF4 |

| |
|---|
| RBL1 |
| RBL2 |
| EZH2 |
| CDK2AP2 |
| BRD7 |
| CDKN2B |
| KANK2 |
| ACVR1 |
| FBXO7 |
| MYO16 |
| CTDSP2 |
| E2F7 |
| BCL2 |
| DCUN1D3 |
| INHBA |
| NACC2 |
| PSMB2 |
| MIIP |
| PSMA5 |
| PSMB1 |
| PSMA6 |
| PSMB9 |
| PSMB8 |
| PSMB5 |
| PSMA7 |
| PSMA1 |
| USP47 |
| PSMB7 |
| PSMB4 |
| PSMB3 |
| RAD21 |
| PSMA4 |
| NAA10 |
| TNKS |
| NPPC |
| NPR2 |
| LIF |
| FBXO5 |
| NANOS2 |
| ANGEL2 |
| BRINP3 |
| TNF |
| HECA |
| BRINP2 |
| BTG2 |

| |
|---|
| FOXC1 |
| GAS1 |
| BTG3 |
| SCRIB |
| TRIM35 |
| NLE1 |
| BTG1 |
| PTPN3 |
| SLC9A3R1 |
| BTG4 |
| BRINP1 |
| ABL1 |
| ZFP36L2 |
| ZFP36L1 |
| CTNNB1 |
| FZD3 |
| SMARCA5 |
| BAZ1B |
| PPP1R10 |
| NME6 |
| MTBP |
| BMP7 |
| TOM1L2 |
| BMP4 |
| TOM1L1 |
| LIG1 |
| HNRNPU |
| KAT2B |
| NEUROG1 |
| PHB2 |
| LSM10 |
| UBE2E2 |
| EIF4G1 |
| ANKRD17 |
| AIF1 |
| SASS6 |
| AKT1 |
| KMT2E |
| HYAL1 |
| ADAMTS1 |
| EGFR |
| MBLAC1 |
| MEPCE |
| RRM2 |
| PLRG1 |

| |
|---|
| ADAM17 |
| CPSF3 |
| RRM1 |
| LSM11 |
| RPTOR |
| CUL4B |
| STOX1 |
| CUL4A |
| TFDP1 |
| CENPJ |
| ANXA1 |
| DDX3X |
| PBX1 |
| CDC7 |
| RCC2 |
| APP |
| SIN3A |
| MTA3 |
| DBF4B |
| RRM2B |
| RAD51B |
| NSMCE2 |
| SLF2 |
| MACROH2A1 |
| SLF1 |
| PIWIL2 |
| LFNG |
| TOPAZ1 |
| WNT4 |
| WNT5A |
| SIRT2 |
| MAPK15 |
| PKN2 |
| ASNS |
| SMOC2 |
| FOXA1 |
| NKX3-1 |
| TAL1 |
| USP2 |
| LGMN |
| SHB |
| MDM2 |
| HSF1 |
| STAT5B |
| POLDIP2 |

| |
|---|
| USP22 |
| SPHK1 |
| PRKCA |
| MEIS2 |
| PTPN11 |
| KLHL18 |
| TMOD3 |
| DYNC1LI1 |
| GEN1 |
| XRCC3 |
| PCID2 |
| HES1 |
| RANBP1 |
| MAP3K20 |
| ANAPC4 |
| ANAPC5 |
| ANAPC11 |
| HOXA13 |
| PDGFB |
| FGF8 |
| LRP5 |
| IGF2 |
| INS |
| EDN3 |
| EGF |
| IGF1 |
| INSR |
| TGFA |
| IL1A |
| IL1B |
| CD28 |
| EREG |
| EPGN |
| RAD51AP1 |
| ZPR1 |
| BTBD8 |
| RNF212 |
| CCNB1IP1 |
| RAD54B |
| SYCE3 |
| RNF212B |
| SHOC1 |
| PSMC3IP |
| CNTD1 |
| CDKN1C |

| |
|---|
| SENP2 |
| APPL1 |
| CDKN2C |
| MNAT1 |
| ID2 |
| CDK7 |
| TM4SF5 |
| PTPN6 |
| CCNH |
| ECD |
| APPL2 |
| PKD1 |
| CDKN2D |
| KCNH5 |
| CTC1 |
| CCNB3 |
| YTHDF2 |
| GPR3 |
| PDE3A |
| CALR |
| ZMPSTE24 |
| RPA3 |
| TP73 |
| THAP1 |
| FOXG1 |
| TTC28 |
| PTCH1 |
| IQGAP1 |
| SDCBP |
| ASCL1 |
| CDC26 |
| GMNN |
| CYLD |
| ZNF268 |
| TTLL12 |
| MCIDAS |
| SIRT1 |
| AATF |
| CDK10 |
| AFAP1L2 |
| RPRM |
| XPC |
| UBD |
| DDB1 |
| ATAD5 |

| |
|---|
| TFAP4 |
| ERCC3 |
| ERCC2 |
| LAMTOR1 |
| USP44 |
| LCMT1 |
| E4F1 |
| CEP85 |
| NEK6 |
| RIOK2 |
| HECW2 |
| CDC42 |
| CUL9 |
| CUL7 |
| CAV2 |
| MKI67 |
| BORA |
| KIF20B |
| OBSL1 |
| CCDC8 |
| FBXW5 |
| CEP97 |
| CCSAP |
| DRG1 |
| GNAI1 |
| RAE1 |
| PARP3 |
| FSD1 |
| BACH1 |
| CDK18 |
| KLF11 |
| NPAT |
| E2F6 |
| CDK17 |
| CDK16 |
| HFM1 |
| CENPS-CORT |
| FANCM |

| Column1 |
| --- |
| MCOLN2 |
| TRPV4 |
| AHSG |
| ITIH4 |
| INS |
| FN1 |
| LBP |
| HP |
| F3 |
| EIF2AK1 |
| VNN1 |
| TREM1 |
| NLRP6 |
| ACVR1 |
| B4GALT1 |
| CD6 |
| ELANE |
| IL31RA |
| THBS1 |
| TNF |
| HIF1A |
| F2R |
| TIMP1 |
| IL1A |
| CXCR6 |
| CCR1 |
| CCR3 |
| CCR2 |
| TNFRSF1B |
| CCR5 |
| CSF1 |
| NXPE3 |
| PTGS2 |
| MTOR |
| PTAFR |
| KNG1 |
| TLR1 |
| TLR6 |
| TRAF3IP2 |
| MYD88 |
| BMP6 |
| TAC1 |
| ACKR1 |
| SELP |

| |
|---|
| SMAD1 |
| CCL24 |
| CCR4 |
| SCYL3 |
| CCL26 |
| KIT |
| TNFRSF4 |
| ACKR2 |
| LIAS |
| CHI3L1 |
| BMPR1B |
| IL6 |
| AKT1 |
| IL17F |
| IL17A |
| CCL28 |
| HYAL3 |
| HYAL1 |
| TUSC2 |
| PTGFR |
| PXK |
| AGER |
| POLB |
| EPHA2 |
| ADAM8 |
| CSF1R |
| NRROS |
| BMP2 |
| REL |
| CMKLR1 |
| ELF3 |
| TIRAP |
| CXCL12 |
| PLAA |
| MAPKAPK2 |
| IL13 |
| TOLLIP |
| S1PR3 |
| PTGIR |
| CCR7 |
| CALCA |
| ALOX5 |
| HRH4 |
| CXCL14 |
| PTGDR |

| |
|---|
| CRH |
| TCIRG1 |
| BDKRB2 |
| PJA2 |
| CCL19 |
| CRHBP |
| C5AR2 |
| CYP26B1 |
| LOXL3 |
| FASN |
| KRT16 |
| TNFRSF1A |
| MFHAS1 |
| NFE2L2 |
| CCL27 |
| AGTR2 |
| MS4A2 |
| AP3B1 |
| SLC11A1 |
| CD40 |
| CCL7 |
| NFKBIB |
| NKIRAS2 |
| CCL8 |
| CCL13 |
| CELA1 |
| CCL1 |
| PTGS1 |
| CCL5 |
| RARRES2 |
| CCL16 |
| CD40LG |
| AXL |
| CCL23 |
| CCL18 |
| IL17C |
| CYBA |
| TGFB1 |
| HMGB1 |
| IL17RE |
| MEFV |
| IDO1 |
| PLP1 |
| RELA |
| C5 |

| |
|---|
| CCL20 |
| IL18 |
| LIPA |
| CCL22 |
| CCL17 |
| NLRP1 |
| GPR32 |
| UMOD |
| MAP2K3 |
| STAT3 |
| IRAK2 |
| PYCARD |
| SPHK1 |
| IL23A |
| ACER3 |
| SCN9A |
| TLR4 |
| ECM1 |
| TLR7 |
| TLR8 |
| TBXA2R |
| PTGER4 |
| IL1B |
| DPEP1 |
| IL37 |
| IL36G |
| IL36A |
| ANXA1 |
| IL36B |
| IL36RN |
| IL1F10 |
| AFAP1L2 |
| IL1RN |
| CXCL11 |
| CXCL10 |
| CD14 |
| CCL25 |
| IL17B |
| PPBP |
| PF4 |
| CXCL1 |
| CXCR3 |
| C3 |
| IL17D |
| CSRP3 |

| |
|---|
| ITGB6 |
| IL5RA |
| IL20RB |
| NOTCH2 |
| RBPJ |
| HMGB2 |
| RASGRP1 |
| NOTCH1 |
| CYSLTR1 |
| WDR83 |
| SCNN1B |
| PTN |
| NINJ1 |
| FFAR2 |
| MDK |
| S100A8 |
| S100A9 |
| TRIM55 |
| JAM3 |
| CX3CL1 |
| FUT7 |
| ALOX5AP |
| ASH1L |
| IL4 |
| TNFAIP3 |
| FOXP3 |
| IL10 |
| ABI3BP |
| ZC3H12A |
| APOD |
| CHID1 |
| PDCD4 |
| PLA2G10 |
| ADCY7 |
| F2 |
| ABCD1 |
| MACIR |
| APPL2 |
| IL1R2 |
| NLRP7 |
| PPARA |
| NR1D2 |
| GHSR |
| RABGEF1 |
| ENPP3 |

| |
|---|
| SOD1 |
| IL22RA2 |
| ADIPOQ |
| C1QTNF12 |
| ADORA1 |
| CCN3 |
| HGF |
| ISL1 |
| LRFN5 |
| TYRO3 |
| GPER1 |
| GATA3 |
| RORA |
| NT5E |
| PTGIS |
| ADA |
| NLRX1 |
| IL2 |
| FFAR4 |
| NR1D1 |
| MMP26 |
| NDFIP1 |
| SMAD3 |
| GPS2 |
| APOA1 |
| FOXF1 |
| OTULIN |
| NR1H3 |
| NR1H4 |
| LPCAT3 |
| CDH5 |
| MVK |
| FPR2 |
| PROC |
| PTPN2 |
| SHARPIN |
| SOCS5 |
| WFDC1 |
| PBK |
| NFKB1 |
| HAMP |
| CXCL17 |
| TNFAIP8L2 |
| RB1 |
| TNFAIP6 |

| |
|---|
| APOE |
| ZFP36 |
| NLRP12 |
| FNDC4 |
| NLRC3 |
| SOCS3 |
| KRT1 |
| METRNL |
| GPX1 |
| MKRN2 |
| ADCY4 |
| VIP |
| PSMA1 |
| GPR17 |
| GHRH |
| ADCY3 |
| ADCY2 |
| NOD2 |
| PSMB4 |
| ADCY8 |
| IL12B |
| ADCY5 |
| GIP |
| ADCY6 |
| ADCY1 |
| GIT1 |
| SIGLEC10 |
| TREM2 |
| IGF1 |
| DUSP10 |
| GRN |
| C2CD4B |
| C2CD4A |
| PARK7 |
| LTA |
| ADORA2B |
| KPNA6 |
| APPL1 |
| GPSM3 |
| PLA2G3 |
| ANKRD42 |
| mml-mir-324 |
| IL17RA |
| CLEC7A |
| TICAM1 |

| |
|---|
| IL17RC |
| CARD9 |
| CAMK2N1 |
| SUCNR1 |
| CD47 |
| WNT5A |
| DEFB114 |
| TLR10 |
| SNCA |
| SERPINE1 |
| CLOCK |
| NAPEPLD |
| TLR3 |
| NFKBIA |
| SETD4 |
| LPL |
| TNFSF4 |
| HYAL2 |
| APP |
| RIPK1 |
| FABP4 |
| OSM |
| TLR2 |
| CEBPB |
| ETS1 |
| TLR9 |
| NLRP10 |
| IFNG |
| TRADD |
| IFI35 |
| STAT5B |
| IL33 |
| NMI |
| CCN4 |
| IL1RL1 |
| LILRA5 |
| GPR4 |
| LDLR |
| CD81 |
| KARS1 |
| CD28 |
| MMP8 |
| PLCG2 |
| CHIA |
| PBXIP1 |

| |
|---|
| IL4R |
| DNASE1L3 |
| RHBDD3 |
| DNASE1 |
| PLA2G2D |
| BAP1 |
| PLD4 |
| PLD3 |
| MAPK14 |
| NOS2 |
| LEP |
| DAGLB |
| BCL6 |
| MCPH1 |
| BST1 |
| DUOXA2 |
| DUOXA1 |
| CMA1 |
| SEMA7A |
| ESR1 |
| SPATA2 |
| JAK2 |
| TNIP1 |
| LRRC19 |
| CASP4 |
| AKNA |
| DROSHA |
| PGLYRP2 |
| MAS1 |
| CYLD |
| STING1 |
| LACC1 |
| SETD6 |
| LYN |
| SBNO2 |
| FANCD2 |
| IRF3 |
| IL1R1 |
| ABHD12 |
| IL1RL2 |
| RICTOR |
| ADAMTS12 |
| PIK3AP1 |
| STK39 |
| TNC |

| |
|---|
| FANCA |
| ALOX15 |
| LETMD1 |
| PTGES |
| CD200R1 |
| CD200R1L |
| DAGLA |
| HMOX1 |

| Column1 |
| --- |
| BST2 |
| KIT |
| CLEC7A |
| SCIMP |
| PLCG2 |
| DHX36 |
| DDX1 |
| TICAM1 |
| KIFBP |
| TLR3 |
| TLR9 |
| TLR4 |
| TRPM2 |
| CCL21 |
| CCL19 |
| TRPM4 |
| GPR183 |
| ANO6 |
| CDC42 |
| EXT1 |
| ALOX5 |
| DOCK8 |
| EPS8 |
| AZI2 |
| TBK1 |
| BCL3 |
| PYCARD |
| SLAMF1 |
| DOCK2 |
| ARHGEF5 |
| CCR7 |
| BATF3 |
| NOTCH2 |
| CSF2 |
| RBPJ |
| BATF |
| TGFBR2 |
| IRF4 |
| TRAF6 |
| UBD |
| IL4 |
| SPI1 |
| DHRS2 |
| CAMK4 |

| |
|---|
| LTBR |
| PSEN1 |
| TGFB1 |
| RELB |
| DCSTAMP |
| BATF2 |
| IL10 |
| HAVCR2 |
| LAG3 |
| IRF8 |
| BLK |
| LYN |
| CCR6 |
| IL12A |
| C1QBP |
| GAS6 |
| CALR |

| Column1 |
| --- |
| FCGR3 |
| PTPRC |
| IL7R |
| KLRD1 |
| PPP3CB |
| CD1D |
| STX7 |
| CD1E |
| MAMU-DOB |
| MAMU-DRA |
| B2M |
| IL12A |
| HLA-E |
| CYRIB |
| IL12B |
| IL23A |
| FADD |
| MR1 |
| SLC22A13 |
| AGER |
| NECTIN2 |
| CTSH |
| CTSC |
| EMP2 |
| HPRT1 |
| PRF1 |
| GZMM |
| JAGN1 |
| GZMB |
| SRGN |
| GZMA |
| STXBP2 |
| LYST |
| RNF19B |
| CD2 |
| PLEKHM2 |
| KIF5B |
| ULBP3 |
| PTPN6 |
| NKG2D |
| PRDX1 |
| IL18 |
| CEBPG |
| IL13 |

| |
|---|
| IL4 |
| ARRB2 |
| CLEC12B |
| HAVCR2 |
| CD5L |
| SPI1 |
| STAP1 |
| CD160 |
| SLAMF6 |
| NCR3 |
| IL21 |
| LAG3 |
| SH2D1A |
| RASGRP1 |
| CD226 |
| AP1G1 |
| CRTAM |
| STAT5B |
| IL18RAP |
| CADM1 |
| VAV1 |
| SERPINB9 |
| CR1L |
| LEP |
| PIK3R6 |
| DNASE1L3 |
| DNASE1 |

| Column1 |
| --- |
| ZC3H12A |
| APOD |
| CHID1 |
| PDCD4 |
| PLA2G10 |
| ADCY7 |
| F2 |
| ABCD1 |
| MACIR |
| MEFV |
| APPL2 |
| IL1R2 |
| NLRP7 |
| PPARA |
| PYCARD |
| KPNA6 |
| APPL1 |
| MYD88 |
| TNF |
| IL6 |
| IL17F |
| IL17A |
| GPSM3 |
| PLA2G3 |
| ANKRD42 |
| HIF1A |
| mml-mir-324 |
| CD6 |
| IL17RA |
| CLEC7A |
| NOD2 |
| TICAM1 |
| IL17RC |
| CARD9 |
| STAT3 |
| TLR4 |
| IL17B |
| BAP1 |
| PLD4 |
| ALOX5 |
| PLD3 |
| MAPK14 |
| NOS2 |
| LEP |

| |
|---|
| HFE |
| SMAD7 |
| FOXP3 |
| ARG1 |
| TBX21 |
| IFNB1 |
| MAP3K7 |
| B2M |
| TRAF6 |
| SASH3 |
| FZD5 |
| MALT1 |
| TRAF2 |
| IL18 |
| IL1R1 |
| IL18R1 |
| IL1B |
| PRKCZ |
| DENND1B |
| GATA3 |
| RSAD2 |
| IL4 |
| CD81 |
| CCR2 |
| TRPM4 |
| CCR9 |
| CCR1 |
| CCR3 |
| CCR5 |
| CCR6 |
| CCR4 |
| CCR8 |
| ACKR2 |
| CCR7 |
| ACKR3 |
| GPR75 |
| CCR10 |
| CXCR5 |
| CXCR3 |
| CXCR6 |
| CXCR2 |
| CXCR1 |
| GPR35 |
| CXCR4 |
| CX3CR1 |

| |
|---|
| CCRL2 |
| CCL26 |
| CCL19 |
| CCL7 |
| CCL16 |
| CCL23 |
| CCL18 |
| CCL20 |
| CCL17 |
| MSMP |
| CCL27 |
| CCL11 |
| NARS1 |
| CNIH4 |
| DEFB1 |
| CCL21 |
| E2F8 |
| OXSR1 |
| RHOA |
| FOXC1 |
| DUSP1 |
| SLC12A2 |
| RIPOR2 |
| DOCK8 |
| LOX |
| WNK1 |
| SH2B3 |
| LRCH1 |
| STK39 |
| CRNN |
| LDLRAP1 |
| MME |
| HCLS1 |
| GBP7 |
| GBP3 |
| HAX1 |
| TRAF3IP2 |
| POU4F2 |
| CSF1R |
| CSF3 |
| IL13 |
| FOXF1 |
| LEF1 |
| TCIRG1 |
| POU4F1 |

| |
|---|
| NFAT5 |
| FOXH1 |
| DPYSL3 |
| CDK9 |
| PID1 |
| TPR |
| PYHIN1 |
| RO60 |
| PDE12 |
| AXL |
| IFIT2 |
| IFIT3 |
| GAS6 |
| UBE2G2 |
| AIM2 |
| IFI16 |
| UBE2K |
| PNPT1 |
| TRIM6 |
| IRF1 |
| HTRA2 |
| OAS1 |
| STAT1 |
| STING1 |
| CDC34 |
| CDC42 |
| EPRS1 |
| CD58 |
| STXBP3 |
| VAMP3 |
| WNT5A |
| ZYX |
| AIF1 |
| SLC26A6 |
| VIM |
| MRC1 |
| KIF5B |
| TLR2 |
| GAPDH |
| RAB7B |
| DAPK1 |
| MYO1C |
| CDC42EP4 |
| IL12B |
| VPS26B |

| |
|---|
| STXBP4 |
| RPS6KB1 |
| GSN |
| IRF8 |
| IL12RB1 |
| AQP4 |
| STX4 |
| RAB20 |
| WAS |
| STXBP1 |
| STX8 |
| ACTR3 |
| ACTG1 |
| ACTR2 |
| RC3H1 |
| HYAL3 |
| HYAL1 |
| HYAL2 |
| MAPK11 |
| HAS2 |
| NKX3-1 |
| RORA |
| PTGIS |
| NR1D1 |
| CD40 |
| NFKB1 |
| USP10 |
| MAPK13 |
| DAB2IP |
| RELA |
| SFRP1 |
| SOX9 |
| ADAMTS12 |
| CACTIN |
| ANKRD1 |
| UPF1 |
| ADAMTS7 |
| CXCL8 |
| TANK |
| SBNO2 |
| CD47 |
| ALOX15 |
| IL18RAP |
| JAK2 |
| MCM2 |

| |
|---|
| HSP90AB1 |
| IMPDH2 |
| PML |
| HSPA5 |
| NFIL3 |
| TCF7 |
| ALAD |
| FASN |
| RUFY4 |
| CDK4 |
| DCSTAMP |
| KEAP1 |
| CORO1A |
| ST3GAL6 |
| SELPLG |
| PHB |
| PTPRT |
| YBX1 |
| HDGF |
| RAD23B |
| LSP1 |
| ATIC |
| BTK |
| GIPC1 |
| P4HB |
| STIP1 |
| EGR1 |
| DHX9 |
| GBA |
| TNFRSF21 |
| MYOG |
| CHI3L1 |
| GSDME |
| NFKBIA |
| AKT1 |
| IKBKB |
| ZFAND6 |
| RIPK1 |
| GPER1 |
| FABP4 |
| CHUK |
| MYOD1 |
| SLC2A4 |
| TCL1A |
| CIB1 |

| |
|---|
| YBX3 |
| CRHBP |
| ZFP36L2 |
| HMHB1 |
| NFE2L2 |
| BIRC2 |
| BAG4 |
| MAPK1 |
| INPP5K |
| SMPD3 |
| TRADD |
| ERBIN |
| BRCA1 |
| RPS3 |
| ZFP36 |
| SIRT1 |
| ZFP36L1 |
| GPD1 |
| NPNT |
| MAPK3 |
| IFIT1 |
| CCL24 |
| CCL28 |
| CXCL12 |
| CXCL16 |
| CCL4L1 |
| CXCL14 |
| CCL2 |
| CCL8 |
| CCL13 |
| CCL1 |
| CCL5 |
| CCL14 |
| CCL3 |
| CCL22 |
| CX3CL1 |
| CXCL13 |
| CXCL11 |
| CXCL10 |
| CXCL9 |
| CCL25 |
| CXCL5 |
| PPBP |
| PF4 |
| CXCL1 |

| |
|---|
| PF4V1 |
| CXCL6 |
| ACKR4 |
| XCR1 |
| S100A14 |
| ACKR1 |
| PTK2B |
| TREM2 |
| WBP1L |
| ITCH |
| IL5RA |
| CSF3R |
| MPL |
| IL12RB2 |
| LEPR |
| IL6R |
| IFNGR1 |
| GHR |
| IFNGR2 |
| IL2RB |
| CSF2RB |
| CD74 |
| FZD4 |
| CD44 |
| IL7R |
| IL13RA2 |
| IL13RA1 |
| CNTFR |
| IL9R |
| IL4R |
| IL21R |
| LIFR |
| OSMR |
| EPOR |
| IL6ST |
| PRLR |
| ANKRD24 |
| IL15 |
| IFNK |
| IL21 |
| IL9 |
| IFNA13 |
| IFNA8 |
| IFNA6 |
| IFNA14 |

| |
|---|
| IFNA16 |
| IFNA10 |
| IFNW1 |
| EDN2 |
| GREM2 |
| IL17RB |
| IRF5 |
| IL1RAP |
| IL20RA |
| KIT |
| SH2B2 |
| IFNAR1 |
| DUOX1 |
| IL7 |
| SLC1A1 |
| FER |
| TNFRSF1A |
| SOCS1 |
| NUMBL |
| STAT4 |
| SOCS5 |
| JAK1 |
| ZC3H15 |
| STAT5B |
| JAK3 |
| IL17RE |
| IL1RAPL2 |
| IRAK3 |
| STAT2 |
| IL1A |
| IL37 |
| IRAK4 |
| TYK2 |
| EREG |
| STAT6 |
| TNFSF11 |
| BAD |
| DLG1 |
| DOCK2 |
| MSN |
| PRF1 |
| IFNG |
| IRF6 |
| MAMU-A |
| IRF9 |

| |
|---|
| IRF4 |
| TP53 |
| NMI |
| IRF7 |
| IL1RL2 |
| IL1RL1 |
| IL36RN |
| IL1RN |
| IL36G |
| IL36A |
| IL36B |
| IL1F10 |
| PSMB2 |
| PSMA5 |
| PSMB1 |
| PSMA6 |
| PSMB9 |
| PSMB8 |
| PLCB1 |
| PSMB5 |
| RPS6KA5 |
| PSMA7 |
| PSMA1 |
| PSMB7 |
| PSMB4 |
| PSMB3 |
| IRAK1 |
| PSMA8 |
| IRAK2 |
| PSMA4 |
| RPS6KA4 |
| IL11 |
| IL23R |
| IL12A |
| IL15RA |
| SYK |
| CD4 |
| IL17RD |
| IL17REL |
| TRAF5 |
| IKBKE |
| NOTCH1 |
| PDGFB |
| IL2RA |
| IL2 |

| |
|---|
| ECM1 |
| IL20 |
| IL22RA2 |
| IL23A |
| IL27RA |
| IL27 |
| MX1 |
| OASL |
| OAS2 |
| IFNL1 |
| IL3 |
| RNF41 |
| IL33 |
| IL34 |
| PIBF1 |
| IL5 |
| SDCBP |
| ADAM17 |
| ERAP1 |
| SPI1 |
| SRC |
| CTR9 |
| YAP1 |
| SMAD4 |
| SHCBP1L |
| ECT2 |
| MAP9 |
| PDCD6IP |
| ANLN |
| EFHC1 |
| APC |
| CUL7 |
| SON |
| TRIM36 |
| RAB35 |
| LZTS2 |
| CEP55 |
| CENPA |
| CKAP2 |
| SPTBN1 |
| EFHC2 |
| SNX9 |
| SNX33 |
| PLK1 |
| RTKN |

| |
|---|
| ANK3 |
| CHMP4B |
| KIF20A |
| SNX18 |
| MYH10 |
| KIF4B |
| STAMBP |
| MITD1 |
| USP8 |
| UNC119 |
| KIF4A |
| ZFYVE26 |
| RACGAP1 |
| CIT |
| BBS4 |
| INCENP |
| SPAST |
| NUSAP1 |
| CHMP4C |
| ZFYVE19 |
| KLHDC8B |
| CNTROB |
| SLIT2 |
| PADI2 |
| ROBO1 |
| SLIT3 |
| RNF113A |
| PTPRC |
| OTOP1 |
| PTPN2 |
| PARP14 |
| OTUD4 |
| TIGIT |
| SLAMF1 |
| CMKLR1 |
| IL10 |
| ARRB2 |
| THBS1 |
| C1QBP |
| LILRA5 |
| TLR8 |
| GHSR |
| RABGEF1 |
| TNFAIP3 |
| SYT11 |

| |
|---|
| PTPN22 |
| HGF |
| NLRX1 |
| TLR9 |
| ORM1 |
| PTPN6 |
| INPP5D |
| NR1H4 |
| C5AR2 |
| NCKAP1L |
| BANK1 |
| ARRB1 |
| NLRP12 |
| HAVCR2 |
| NLRC3 |
| FOXJ1 |
| CD96 |
| BST2 |
| KLF4 |
| APOA1 |
| MAPK7 |
| CD34 |
| DEFB114 |
| NFKBIL1 |
| ILRUN |
| ADIPOQ |
| DICER1 |
| TRIM27 |
| RARA |
| VSIR |
| GSTP1 |
| CIDEA |
| LBP |
| IGF1 |
| BCL3 |
| ARG2 |
| CLEC4A |
| CD33 |
| GPR18 |
| MC1R |
| AKAP8 |
| CD274 |
| TRAIP |
| HIST1H2BJ |
| GPS2 |

| |
|---|
| XIAP |
| NAIP |
| F2RL1 |
| PELI3 |
| RFFL |
| CCDC3 |
| NOL3 |
| PPP2CB |
| PYDC1 |
| PIAS4 |
| MUL1 |
| DCST1 |
| YTHDF2 |
| ADAR |
| ISG15 |
| METTL3 |
| SAMHD1 |
| YTHDF3 |
| OAS3 |
| MMP12 |
| TTLL12 |
| NLRC5 |
| TRIM32 |
| MCOLN2 |
| TRPV4 |
| MAVS |
| ADCYAP1 |
| LPL |
| MBP |
| CHIA |
| AIRE |
| TLR3 |
| RIPK2 |
| AGER |
| APP |
| ADORA2B |
| EIF2AK2 |
| FFAR2 |
| C5 |
| TLR7 |
| EDN1 |
| WNT3A |
| GLMN |
| BCL10 |
| TLR5 |

| |
|---|
| HHLA2 |
| FGR |
| SOD1 |
| CLEC5A |
| TMF1 |
| FLOT1 |
| BATF |
| BTN3A1 |
| AGPAT1 |
| HTR2B |
| PELI1 |
| ADRA2A |
| LURAP1 |
| INS |
| IL26 |
| SLC11A1 |
| CLEC9A |
| RAB5C |
| CRTAM |
| ITK |
| NFAM1 |
| PANX1 |
| CLEC6A |
| RGCC |
| PTGER4 |
| CADM1 |
| TMIGD2 |
| CCDC88B |
| AGPAT2 |
| PLA2R1 |
| TNFRSF14 |
| INAVA |
| SPHK2 |
| HK1 |
| LACC1 |
| SCIMP |
| TRIM44 |
| PAFAH1B1 |
| PLCG2 |
| DHX36 |
| SETD2 |
| TBK1 |
| IRF3 |
| DDX3X |
| ZBTB20 |

| |
|---|
| POLR3C |
| TOMM70 |
| POLR3D |
| HMGB2 |
| POLR3A |
| POLR3G |
| RNF135 |
| RIOK3 |
| DHX58 |
| POLR3F |
| POLR3B |
| PTPN11 |
| KCNH7 |
| CD160 |
| CD2 |
| PDE4B |
| SLAMF6 |
| LTA |
| CD244 |
| TNFSF4 |
| ISL1 |
| PDE4D |
| CYRIB |
| SCRIB |
| NKG2D |
| RASGRP1 |
| CD3E |
| SLC7A5 |
| ZFPM1 |
| HRAS |
| FADD |
| CEBPG |
| CD14 |
| ABL1 |
| HPX |
| MED1 |
| PARP9 |
| HSPB1 |
| CD36 |
| GSDMD |
| TRIM16 |
| P2RX7 |
| STMP1 |
| TMEM106A |
| NLRP1 |

| |
|---|
| AZU1 |
| USP50 |
| CASP8 |
| NLRC4 |
| PANX2 |
| PANX3 |
| TMED10 |
| CD46 |
| IL20RB |
| TUSC2 |
| CD40LG |
| CD83 |
| CD28 |
| LTB |
| MDK |
| IDO1 |
| HLA-E |
| TIRAP |
| ZBTB7B |
| PRKCQ |
| OSM |
| TGFB1 |
| VTCN1 |
| RUNX1 |
| CARD11 |
| PRKD2 |
| ANXA1 |
| STOML2 |
| CSF2 |
| CEBPB |
| EPX |
| TLR1 |
| ARHGEF2 |
| NOD1 |
| SETD4 |
| C1QTNF4 |
| F2R |
| MMP8 |
| CYBA |
| POU2AF1 |
| POU2F2 |
| GFI1 |
| PARK7 |
| APOA2 |
| SERPINE1 |

| |
|---|
| F3 |
| RAB1A |
| GDF2 |
| PRG3 |
| ELANE |
| AFAP1L2 |
| FCN1 |
| DDIT3 |
| SEMA7A |
| NUP62 |
| KIF20B |
| DDX1 |
| KIFBP |
| CLNK |
| CD226 |
| ADAM8 |
| EPHB2 |
| TNFRSF8 |
| NFATC4 |
| MAPKAPK2 |
| SPN |
| LY96 |
| CLU |
| PSEN1 |
| FRMD8 |
| PIK3R1 |
| ARFGEF2 |
| PRKN |
| CASP4 |
| TRIM15 |
| CGAS |
| G3BP1 |
| DHX33 |
| ZCCHC3 |
| PQBP1 |
| TRIM56 |
| ZBP1 |
| LSM14A |
| TRIM41 |
| USP27X |
| VRK2 |
| MAST2 |
| ANKRD53 |
| LTF |
| UBE2J1 |

| |
|---|
| ANGPT1 |
| NKIRAS1 |
| HIPK1 |
| SPATA2 |
| OTULIN |
| CYLD |
| SHARPIN |
| NKIRAS2 |
| SPHK1 |
| ITIH4 |
| SRF |
| TIMP4 |
| MAPKAPK3 |
| ALDH1A2 |
| REL |
| BCL2L1 |
| TMEM102 |
| PPP3CB |
| AVPR2 |
| BCL2 |
| COL3A1 |
| MCL1 |
| TNFRSF11A |
| LAMP3 |
| MX2 |
| IFNAR2 |
| SHFL |
| XAF1 |
| SNCA |
| UBD |
| TRIM21 |
| CIITA |
| GCH1 |
| KYNU |
| CALCOCO2 |
| CYP27B1 |
| SP100 |
| SELE |
| YTHDC2 |
| SMPD1 |
| IGBP1 |
| PRKCA |
| SHANK3 |
| ADAM10 |
| MAP4K3 |

| |
|---|
| ADAM9 |
| AFF3 |
| CH25H |
| SHMT2 |
| TNFSF10 |
| SIVA1 |
| FASLG |
| TRAF3 |
| TNFSF12 |
| BABAM2 |
| TNFSF13B |
| TRIM37 |
| TRAF1 |
| EDA |
| TNFSF8 |
| TRAF4 |
| TNFSF15 |
| TNFSF9 |
| CD70 |
| TNFSF14 |
| TNFSF18 |
| TNFRSF1B |
| FOXO3 |
| RRAGA |
| ACTN4 |
| ILK |
| CARD14 |
| CDIP1 |
| TNFRSF17 |
| EDA2R |
| PLVAP |
| UMOD |
| TXNDC17 |
| TNFRSF13C |
| IRF2 |
| IFI27 |

| Column1 |
| --- |
| WDFY4 |
| HLA-E |
| CLEC4A |
| PSMB11 |
| IRF1 |
| PAX1 |
| TNFSF8 |
| EOMES |
| TOX |
| BCL2 |
| CCR9 |
| GPR18 |
| HFE |
| CD274 |
| ZBTB7B |
| SOCS1 |
| VSIR |
| NCKAP1L |
| FADD |
| CBFB |
| CCR2 |
| PTPN22 |
| SART1 |
| CRTAM |
| MAPK8IP1 |

| Column1 |
|---|
| TGFB1 |
| LTF |
| TF |
| CAMP |
| JCHAIN |
| DEFB1 |
| DEFA1B |
| DEFA4 |
| IGHM |
| HIST1H2BJ |
| RNASE6 |
| RNASE7 |
| CTSG |
| HLA-E |
| PLA2G6 |
| SEMG2 |
| SLPI |
| PLA2G1B |
| FAU |
| BPIFA1 |
| RARRES2 |
| IL36RN |
| FAM3A |
| S100A12 |
| S100A9 |
| PGLYRP4 |
| HRG |
| MUC7 |
| DEFB127 |
| DEFB118 |
| RPL30 |
| GAPDH |
| F2 |
| CCL13 |
| REG3G |
| CXCL10 |
| PGLYRP1 |
| PF4 |
| CXCL1 |
| CXCL6 |
| DEFA6 |
| DEFB4 |
| CST9 |
| BCL3 |

| |
|---|
| AZU1 |
| EPHB2 |
| TXLNA |
| FCRL1 |
| LAT2 |
| LAX1 |
| IL4 |
| PHB2 |
| MS4A1 |
| RASGRP1 |
| PRKCB |
| CD40 |
| BANK1 |
| CD22 |
| PHB |
| MALT1 |
| CD79A |
| CXCR5 |
| IFNB1 |
| TRAF3IP2 |
| BTK |
| CYP7B1 |
| HSD3B7 |
| CH25H |
| GAS6 |
| TNFSF13B |
| CD320 |
| CR2 |
| RABL3 |
| NTRK1 |
| BCL6 |
| GON4L |
| FZD9 |
| RBPJ |
| NKX2-3 |
| ONECUT1 |
| PTPRC |
| CDH17 |
| CARD11 |
| DCAF1 |
| TSHR |
| RAG2 |
| RAG1 |
| GPS2 |
| IL9 |

| |
|---|
| TCIRG1 |
| SP3 |
| EZH2 |
| ZBTB1 |
| MSH2 |
| PTPN2 |
| DNAJB9 |
| CD40LG |
| JAK3 |
| YY1 |
| DCLRE1C |
| NHEJ1 |
| PLCG2 |
| BCL2 |
| ADGRG3 |
| PIK3R1 |
| CEBPG |
| EP300 |
| ZBTB7A |
| CD79B |
| PRKDC |
| TP53 |
| CD19 |
| CTPS1 |
| PRKCD |
| LEF1 |
| IL7R |
| SHB |
| ABL1 |
| PLCL2 |
| GAPT |
| CD180 |
| TLR4 |
| SPI1 |
| IRF8 |
| ADA |
| POU2AF1 |
| ITFG2 |
| KLHL6 |
| UNC13D |
| CCR2 |
| NOTCH2 |
| AIRE |
| TNFRSF21 |
| TNF |

| |
|---|
| LTA |
| TFEB |
| GATA3 |
| BMI1 |
| ALOX5 |
| SH2D1A |
| NOTCH1 |
| FOXJ1 |
| GPR183 |
| TFE3 |
| EXO1 |
| CD81 |
| IRF2BP2 |
| KIT |
| BCL10 |
| MYD88 |
| FCER1G |
| CSF2RB |
| CD74 |
| INPP5D |
| IL13RA2 |
| IL4R |
| CARD9 |
| IRF7 |
| DLL1 |
| LFNG |
| MFNG |
| DOCK11 |
| PTK2B |
| POU2F2 |
| ST3GAL1 |
| TNFAIP3 |
| TBC1D10C |
| SFRP1 |
| RC3H1 |
| BLK |
| IL10 |
| ATM |
| CD300A |
| LYN |
| PAWR |
| PKN1 |
| CTLA4 |
| FCRL3 |
| LPXN |

| |
|---|
| PTPN6 |
| PARP3 |
| THOC1 |
| NDFIP1 |
| FOXP3 |
| ITM2A |
| ACOD1 |
| IL6 |
| IGHE |
| PELI1 |
| NOD2 |
| TICAM1 |
| SH3KBP1 |
| MMP14 |
| XBP1 |
| TAPBPL |
| PPP2R3C |
| IL7 |
| SYK |
| NCKAP1L |
| STAT5B |
| PCID2 |
| ATP11C |
| BAD |
| WNT3A |
| VAV3 |
| CD38 |
| TFRC |
| IL2 |
| IL21 |
| IRS2 |
| ATAD5 |
| SASH3 |
| CDKN1A |
| NFATC2 |
| STAP1 |
| SLC39A10 |
| NFAM1 |
| ZP4 |
| CCR7 |
| HPX |
| FCER2 |
| CD226 |
| NECTIN2 |
| HMCES |

| |
|---|
| MAD2L2 |
| PAXIP1 |
| SHLD1 |
| KMT5B |
| TP53BP1 |
| EXOSC3 |
| SHLD3 |
| RIF1 |
| KMT5C |
| MLH1 |
| NSD2 |
| STAT6 |
| TBX21 |
| CD28 |
| GATA6 |
| THEMIS2 |
| TLR9 |
| IKZF3 |
| ZFP36L2 |
| ZFP36L1 |
| MZB1 |
| GCSAML |
| GCSAM |
| SPNS2 |
| SUPT6H |
| SLC15A4 |
| IL27RA |

| Column1 |
| --- |
| BLK |
| LYN |
| SIRT1 |
| YWHAG |
| SFN |
| YWHAQ |
| YWHAB |
| YWHAE |
| YWHAZ |
| IL10 |
| CD274 |
| PDCD1 |
| IDO1 |
| LGALS14 |
| LGALS13 |
| CD36 |
| TYRO3 |
| ADGRB1 |
| RARA |
| MFGE8 |
| NR1H3 |
| TGM2 |
| AXL |
| ITGAV |
| MARCO |
| MERTK |
| ITGB3 |
| GAS6 |
| JMJD6 |
| XKR8 |
| RHOH |
| MEGF10 |
| RHOG |
| RHOBTB1 |
| THBS1 |
| BECN1 |
| LAPTM5 |
| PDCD10 |
| STK25 |
| DDIT3 |
| SCN2A |
| MAP3K5 |
| GPX1 |
| ARL6IP5 |

| |
|---|
| PML |
| DIABLO |
| ZNF622 |
| GSKIP |
| MELK |
| STK24 |
| BCL2 |
| CYP1B1 |
| PDK1 |
| PINK1 |
| PARK7 |
| PTGS2 |
| EPO |
| YBX3 |
| BDKRB2 |
| DNAJA1 |
| VNN1 |
| HSPB1 |
| SOD2 |
| BAG5 |
| NFE2L2 |
| NME5 |
| NOL3 |
| MAPK7 |
| HIF1A |
| PRKN |
| SPG11 |
| C3 |
| UTP11 |
| STPG1 |
| HRG |
| TNFRSF8 |
| FOXL2 |
| NOTCH2 |
| TNFSF10 |
| ECT2 |
| CCN1 |
| BCL10 |
| GADD45A |
| HOXA5 |
| VAV3 |
| DFFA |
| RPS6KA2 |
| SLIT2 |
| CASP2 |

| |
|---|
| IRF5 |
| BCLAF1 |
| SFRP4 |
| ATF6 |
| JUN |
| OMA1 |
| CASP3 |
| TNF |
| BCL6 |
| SPDEF |
| PDCD2 |
| RRP1B |
| CASP9 |
| C2H3orf38 |
| BAK1 |
| CTNNB1 |
| IP6K2 |
| QRICH1 |
| UNC5C |
| HTT |
| IL6 |
| ALDH1A3 |
| DDX20 |
| SCIN |
| MOAP1 |
| RBM5 |
| FASLG |
| APC |
| RIPK2 |
| BNIP3 |
| NET1 |
| FOXA1 |
| PLEKHN1 |
| RIPK1 |
| BMP7 |
| ALDH1A2 |
| NFATC4 |
| TP53INP1 |
| CCAR2 |
| PRKDC |
| SYCE3 |
| ITGB1 |
| KCNMA1 |
| PHLDA3 |
| MAPK8 |

| |
|---|
| ING3 |
| SAV1 |
| BMP2 |
| RARB |
| SFRP2 |
| KLF11 |
| STK3 |
| FOXO3 |
| PRMT2 |
| MTCH2 |
| LATS1 |
| PTPA |
| UBD |
| ING4 |
| STK4 |
| VAV2 |
| TGFBR1 |
| DAPK1 |
| FANK1 |
| BCL2L1 |
| FLCN |
| DUSP6 |
| TOP2A |
| MAP3K10 |
| HIP1R |
| NF1 |
| INPP5D |
| BID |
| GADD45G |
| SPHK2 |
| BARD1 |
| PRR7 |
| SHQ1 |
| CASP6 |
| BOK |
| MMP9 |
| SCRIB |
| HTRA2 |
| GRIN2A |
| BAX |
| BNIP3L |
| ING5 |
| CLIP3 |
| CTNNBL1 |
| POU4F1 |

| |
|---|
| TP53 |
| MTCH1 |
| TFAP4 |
| ATM |
| TRIM35 |
| OSGIN1 |
| NCOA1 |
| ERCC3 |
| SEPTIN4 |
| BMP4 |
| BIN1 |
| RACK1 |
| CRADD |
| ZNF268 |
| GPLD1 |
| RASSF2 |
| CAMK1D |
| AIFM2 |
| FRZB |
| CLU |
| SLC27A4 |
| HMOX1 |
| APBB1 |
| WNT10B |
| SOX4 |
| E2F1 |
| SMPD1 |
| PHLDA2 |
| BCL2L11 |
| NOTCH1 |
| PSEN1 |
| CCAR1 |
| ARHGEF6 |
| HTRA4 |
| DAB2IP |
| ACVR1C |
| TNFRSF12A |
| RNF122 |
| GZMA |
| PHLDA1 |
| REST |
| MLLT11 |
| NR4A1 |
| FOXO1 |
| PIP5KL1 |

| |
|---|
| SFRP1 |
| RBCK1 |
| WNT11 |
| C1QBP |
| PDCD5 |
| CSRNP3 |
| B4GALT1 |
| IFIT2 |
| UBE2Z |
| STK17A |
| BMF |
| TFPT |
| EMILIN2 |
| TCTN3 |
| HMGA2 |
| ARHGEF7 |
| PYCARD |
| MCL1 |
| SIAH1 |
| ADAMTSL4 |
| NEUROD1 |
| SUDS3 |
| DCUN1D3 |
| ANKRD1 |
| DNM1L |
| FADD |
| CALHM2 |
| FAM162A |
| NEURL1 |
| ZBTB16 |
| CTLA4 |
| DDX3X |
| RARG |
| DAPK2 |
| NLRC4 |
| ABL1 |
| USP27X |
| BAD |
| GADD45B |
| VAV1 |
| MAP3K20 |
| VDR |
| TNFRSF1A |
| PRKCD |
| TPD52L1 |

| |
|---|
| TP63 |
| CTSH |
| NKX3-1 |
| RPP21 |
| JAK2 |
| CTSC |
| IL19 |
| APAF1 |
| TRIM65 |
| IFNB1 |
| NGFR |
| RPS3 |
| MAPK9 |
| TRAF7 |
| INHBB |
| LTK |
| TIGAR |
| ZC3H12A |
| EIF4G1 |
| FAF1 |
| AKR1C3 |
| ACOX2 |
| CIDEB |
| AIFM1 |
| PRNP |
| MT3 |
| CDKN1B |
| PHB |
| AXIN2 |
| ACER2 |
| TRIM13 |
| RAMP3 |
| LRP1 |
| NCK1 |
| SERINC3 |
| NCK2 |
| PTPN2 |
| PRKCI |
| CD160 |
| PDCD4 |
| CD248 |
| CD40 |
| CD40LG |
| ITGA4 |
| RGCC |

| |
|---|
| ANO6 |
| TREM2 |
| TP53BP2 |
| DLC1 |
| PTGIS |
| SIRT2 |
| FAP |
| G0S2 |
| AGT |
| HYAL2 |
| PTPRC |
| PAK2 |
| TNFSF12 |
| LTBR |
| AGTR2 |
| CYLD |
| WWOX |
| DEDD2 |
| ITM2C |
| TRAF2 |
| CTNNA1 |
| SRPX |
| PPP1CA |
| PPP2R1B |
| INHBA |
| SKIL |
| PEA15 |
| BMPR1B |
| MAL |
| ZSWIM2 |
| STK17B |
| PLA2G1B |
| PLA2R1 |
| S100A8 |
| S100A9 |
| LCK |
| IL20RA |
| FIS1 |
| GSDME |
| STYXL1 |
| PRKRA |
| PLAGL2 |
| SLC9A3R1 |
| BCAP31 |
| PLEKHF1 |

| |
|---|
| MSX1 |
| UBB |
| MYC |
| RPS7 |
| FBH1 |
| RAD9A |
| NACC2 |
| PIAS4 |
| GSN |
| HOXA13 |
| ADIPOQ |
| PCSK9 |
| TGFB2 |
| TFAP2B |
| ATF4 |
| RAPSN |
| BACE1 |
| ASCL1 |
| EPHA7 |
| PIK3CD |
| PIK3CB |
| ANXA1 |
| SFPQ |
| SOD1 |
| NOX1 |
| FBXW7 |
| CDKN1A |
| LRRK2 |
| IL12A |
| MAP2K4 |
| IFNG |
| IL12B |
| RBM10 |
| WNT5A |
| ADAM8 |
| ZC3H8 |
| ATF3 |
| TIMP3 |
| CAPN10 |
| MFN2 |
| E2F3 |
| PJVK |
| PEAR1 |
| FCN3 |
| SCARB1 |

| |
|---|
| FCN1 |
| UBQLN1 |
| P4HB |

RSAD2
CDH26
STOML2
CD81
BRAF
NKX2-3
PAX1
ARMC5
CRK
CD4
TARM1
TWSG1
CD160
CBFB
VSIR
CD274
ARG2
RC3H1
MAMU-DRA
ZBTB7B
SOCS1
SASH3
NCKAP1L
CD83
TGFBR2
MYB
SHB
AGER
GATA3
IL12B
PTGER4
