## Supplemental Table 4 for "Mild/Asymptomatic Maternal SARS-CoV-2 Infection Leads to Immune Paralysis in Fetal Circulation and Immune Dysregulation in Fetal-Placental Tissues": Sup Table 4 Villous cluster markers.pdf

| p_val | avg_log2FC | pct.1 | pct.2 | p_val_adj | cluster | gene |
|---|---|---|---|---|---|---|
| 0 | 0.90881624 | 0.941 | 0.636 | 0 | 0 | GPR183 |
| 0 | 0.8250506 | 0.897 | 0.539 | 0 | 0 | C15orf48 |
| 0 | 0.81764018 | 0.959 | 0.882 | 0 | 0 | CCL4L2 |
| 0 | 0.77585166 | 0.98 | 0.894 | 0 | 0 | CCL3L1 |
| 0 | 0.77296888 | 0.955 | 0.618 | 0 | 0 | OLR1 |
| 0 | 0.74345087 | 0.986 | 0.776 | 0 | 0 | HLA-DPB1 |
| 0 | 0.7024596 | 0.902 | 0.574 | 0 | 0 | G0S2 |
| 0 | 0.70008666 | 0.718 | 0.475 | 0 | 0 | MT2A |
| 0 | 0.69357262 | 0.943 | 0.644 | 0 | 0 | HLA-DQB1 |
| 0 | 0.6905138 | 0.99 | 0.831 | 0 | 0 | HLA-DPA1 |
| 0 | 0.64957465 | 0.998 | 0.955 | 0 | 0 | HLA-DRA |
| 0 | 0.62063184 | 0.84 | 0.487 | 0 | 0 | HLA-DQA1 |
| 0 | 0.5993855 | 0.987 | 0.794 | 0 | 0 | IFI30 |
| 0 | 0.59249442 | 0.8 | 0.52 | 0 | 0 | CXCR4 |
| 0 | 0.56237737 | 0.978 | 0.736 | 0 | 0 | BCL2A1 |
| 0 | 0.54837607 | 0.999 | 0.988 | 0 | 0 | RPS19 |
| 0 | 0.54302107 | 0.995 | 0.911 | 0 | 0 | S100A11 |
| 0 | 0.53912403 | 0.996 | 0.917 | 0 | 0 | HLA-DRB1 |
| 0 | 0.53080986 | 0.969 | 0.82 | 0 | 0 | PLIN2 |
| 0 | 0.51935732 | 0.999 | 0.994 | 0 | 0 | RPS27 |
| 0 | 0.48975003 | 0.999 | 0.994 | 0 | 0 | RPS29 |
| 0 | 0.48683623 | 0.573 | 0.379 | 0 | 0 | NR4A3 |
| 0 | 0.48624504 | 0.829 | 0.546 | 0 | 0 | UPP1 |
| 0 | 0.48345125 | 0.957 | 0.788 | 0 | 0 | TNFAIP3 |
| 0 | 0.47794174 | 0.657 | 0.478 | 0 | 0 | ISG15 |
| 0 | 0.46801978 | 0.964 | 0.642 | 0 | 0 | S100A10 |
| 0 | 0.46650175 | 0.962 | 0.669 | 0 | 0 | LGALS1 |
| 0 | 0.46604052 | 0.999 | 0.994 | 0 | 0 | RPS28 |
| 0 | 0.46361848 | 0.999 | 0.994 | 0 | 0 | RPL28 |
| 0 | 0.46312563 | 0.818 | 0.601 | 0 | 0 | VEGFA |
| 0 | 0.46115959 | 0.999 | 0.97 | 0 | 0 | SRGN |
| 0 | 0.45990822 | 0.498 | 0.255 | 0 | 0 | FN1 |
| 0 | 0.45410411 | 0.811 | 0.593 | 0 | 0 | C1orf162 |
| 0 | 0.4513362 | 0.996 | 0.976 | 0 | 0 | RPS21 |
| 0 | 0.4487985 | 0.954 | 0.703 | 0 | 0 | CD44 |
| 0 | 0.44511593 | 0.977 | 0.795 | 0 | 0 | PLAUR |
| 0 | 0.44478473 | 0.981 | 0.711 | 0 | 0 | S100A4 |
| 0 | 0.44473642 | 0.972 | 0.736 | 0 | 0 | IL1B |
| 0 | 0.4436427 | 0.996 | 0.975 | 0 | 0 | RPS4X |
| 0 | 0.4431849 | 0.998 | 0.982 | 0 | 0 | RPL18A |
| 0 | 0.44141904 | 0.983 | 0.679 | 0 | 0 | S100A6 |
| 0 | 0.44081145 | 0.996 | 0.991 | 0 | 0 | CCL3 |

| | | | | | | |
|---|---|---|---|---|---|---|
| 0 | 0.43977867 | 0.998 | 0.987 | 0 | 0 | RPS27A |
| 0 | 0.43789712 | 0.849 | 0.574 | 0 | 0 | LGALS3 |
| 0 | 0.42172587 | 0.996 | 0.973 | 0 | 0 | ATP5F1E |
| 0 | 0.42112922 | 0.999 | 0.986 | 0 | 0 | RPS15 |
| 0 | 0.41984692 | 0.999 | 0.989 | 0 | 0 | RPL30 |
| 0 | 0.418608 | 1 | 0.999 | 0 | 0 | RPL41 |
| 0 | 0.41846603 | 0.99 | 0.719 | 0 | 0 | VIM |
| 0 | 0.41550165 | 1 | 1 | 0 | 0 | FTH1 |
| 0 | 0.41525715 | 1 | 0.998 | 0 | 0 | RPL10 |
| 0 | 0.41475449 | 0.745 | 0.568 | 0 | 0 | RASGEF1B |
| 0 | 0.4124776 | 0.799 | 0.584 | 0 | 0 | RGS1 |
| 0 | 0.41179803 | 0.999 | 0.994 | 0 | 0 | RPL34 |
| 0 | 0.40905736 | 0.815 | 0.542 | 0 | 0 | XIST |
| 0 | 0.40825772 | 0.999 | 0.995 | 0 | 0 | RPS12 |
| 0 | 0.40520323 | 0.998 | 0.983 | 0 | 0 | RPL36 |
| 0 | 0.40358937 | 0.84 | 0.63 | 0 | 0 | DUSP2 |
| 0 | 0.40325233 | 0.999 | 0.993 | 0 | 0 | RPL11 |
| 0 | 0.40114018 | 0.965 | 0.873 | 0 | 0 | RPS5 |
| 0 | 0.39712457 | 0.922 | 0.789 | 0 | 0 | RPSA |
| 0 | 0.39696153 | 0.723 | 0.498 | 0 | 0 | SERPINB9 |
| 0 | 0.39223129 | 0.999 | 0.993 | 0 | 0 | RPS18 |
| 0 | 0.39144339 | 0.493 | 0.276 | 0 | 0 | CD9 |
| 0 | 0.39128135 | 0.996 | 0.966 | 0 | 0 | RPL8 |
| 0 | 0.39125018 | 0.999 | 0.992 | 0 | 0 | RPS8 |
| 0 | 0.39118551 | 0.997 | 0.973 | 0 | 0 | RPL18 |
| 0 | 0.389811 | 0.997 | 0.978 | 0 | 0 | RPL35 |
| 0 | 0.38952986 | 0.998 | 0.981 | 0 | 0 | UBA52 |
| 0 | 0.38815299 | 0.723 | 0.508 | 0 | 0 | INSIG1 |
| 0 | 0.387686 | 0.981 | 0.916 | 0 | 0 | CD83 |
| 0 | 0.38382279 | 0.927 | 0.752 | 0 | 0 | EMP3 |
| 0 | 0.38019783 | 0.998 | 0.991 | 0 | 0 | RPS14 |
| 0 | 0.37856012 | 0.783 | 0.634 | 0 | 0 | NAP1L1 |
| 0 | 0.37789741 | 0.801 | 0.627 | 0 | 0 | COMMD6 |
| 0 | 0.37769873 | 0.996 | 0.977 | 0 | 0 | RPL35A |
| 0 | 0.37754287 | 0.694 | 0.496 | 0 | 0 | CTSH |
| 0 | 0.37233952 | 0.836 | 0.579 | 0 | 0 | SERPINA1 |
| 0 | 0.37060831 | 0.985 | 0.923 | 0 | 0 | RPL10A |
| 0 | 0.36677129 | 0.919 | 0.756 | 0 | 0 | CDKN1A |
| 0 | 0.36598986 | 0.689 | 0.428 | 0 | 0 | CD52 |
| 0 | 0.36528265 | 0.999 | 0.989 | 0 | 0 | CD74 |
| 0 | 0.36448749 | 0.828 | 0.629 | 0 | 0 | ANXA2 |
| 0 | 0.3640414 | 0.441 | 0.254 | 0 | 0 | SORL1 |
| 0 | 0.36391869 | 1 | 0.998 | 0 | 0 | TMSB10 |

| | | | | | | |
|---|---|---|---|---|---|---|
| 0 | 0.36340932 | 0.999 | 0.98 | 0 | 0 | SOD2 |
| 0 | 0.36336536 | 0.997 | 0.985 | 0 | 0 | FAU |
| 0 | 0.36270598 | 0.995 | 0.959 | 0 | 0 | BTG1 |
| 0 | 0.3618914 | 0.999 | 0.996 | 0 | 0 | RPL39 |
| 0 | 0.35790366 | 0.984 | 0.886 | 0 | 0 | SH3BGRL3 |
| 0 | 0.35236861 | 0.913 | 0.78 | 0 | 0 | TYMP |
| 0 | 0.34902317 | 0.837 | 0.715 | 0 | 0 | MIF |
| 0 | 0.34700477 | 0.998 | 0.988 | 0 | 0 | RPS15A |
| 0 | 0.34323196 | 0.544 | 0.336 | 0 | 0 | APOBEC3A |
| 0 | 0.34150433 | 0.909 | 0.781 | 0 | 0 | NME2 |
| 0 | 0.34143873 | 0.999 | 0.991 | 0 | 0 | RPLP2 |
| 0 | 0.34104458 | 0.849 | 0.687 | 0 | 0 | HLA-DMA |
| 0 | 0.33966549 | 0.972 | 0.908 | 0 | 0 | RPS17 |
| 0 | 0.33455789 | 0.688 | 0.49 | 0 | 0 | BHLHE40 |
| 0 | 0.33371178 | 0.464 | 0.293 | 0 | 0 | PEA15 |
| 0 | 0.32963318 | 0.929 | 0.816 | 0 | 0 | LST1 |
| 0 | 0.32798516 | 0.889 | 0.712 | 0 | 0 | COTL1 |
| 0 | 0.32790343 | 0.96 | 0.881 | 0 | 0 | COX7C |
| 0 | 0.3273389 | 0.952 | 0.843 | 0 | 0 | ZFAS1 |
| 0 | 0.32513735 | 0.998 | 0.987 | 0 | 0 | RPL12 |
| 0 | 0.32509261 | 0.996 | 0.977 | 0 | 0 | PSAP |
| 0 | 0.32477229 | 0.79 | 0.589 | 0 | 0 | GPCPD1 |
| 0 | 0.32322871 | 0.999 | 0.993 | 0 | 0 | RPL32 |
| 0 | 0.3231882 | 0.995 | 0.97 | 0 | 0 | SERF2 |
| 0 | 0.32276866 | 0.999 | 0.991 | 0 | 0 | RPS2 |
| 0 | 0.32215835 | 0.919 | 0.831 | 0 | 0 | GPX4 |
| 0 | 0.32108157 | 0.999 | 0.996 | 0 | 0 | RPS23 |
| 0 | 0.32060967 | 0.946 | 0.867 | 0 | 0 | EEF1B2 |
| 0 | 0.32057233 | 0.997 | 0.967 | 0 | 0 | RPS3 |
| 0 | 0.31938772 | 0.673 | 0.435 | 0 | 0 | CAPG |
| 0 | 0.318962 | 0.998 | 0.99 | 0 | 0 | RPL26 |
| 0 | 0.31810585 | 0.997 | 0.983 | 0 | 0 | RPS13 |
| 0 | 0.31681848 | 0.998 | 0.985 | 0 | 0 | RPS11 |
| 0 | 0.31663717 | 0.841 | 0.531 | 0 | 0 | EREG |
| 0 | 0.31639107 | 0.997 | 0.972 | 0 | 0 | RPS7 |
| 0 | 0.3128763 | 0.998 | 0.988 | 0 | 0 | RPL38 |
| 0 | 0.31200086 | 0.996 | 0.98 | 0 | 0 | RPL21 |
| 0 | 0.31057817 | 0.999 | 0.994 | 0 | 0 | RPL37A |
| 0 | 0.30837546 | 0.998 | 0.989 | 0 | 0 | RPL9 |
| 0 | 0.30441185 | 0.995 | 0.974 | 0 | 0 | RPL24 |
| 0 | 0.30274628 | 0.997 | 0.981 | 0 | 0 | RPL29 |
| 0 | 0.30214642 | 0.999 | 0.997 | 0 | 0 | RPS24 |
| 0 | 0.30174552 | 0.998 | 0.985 | 0 | 0 | RPS9 |

| | | | | | | |
|---|---|---|---|---|---|---|
| 0 | 0.30101244 | 0.996 | 0.979 | 0 | 0 | RPL36A |
| 0 | 0.29119407 | 0.711 | 0.501 | 0 | 0 | PPIF |
| 0 | 0.28695684 | 0.996 | 0.979 | 0 | 0 | RPL14 |
| 0 | 0.28469999 | 0.716 | 0.453 | 0 | 0 | CRIP1 |
| 0 | 0.28445326 | 1 | 0.997 | 0 | 0 | RPL13 |
| 0 | 0.28382576 | 0.998 | 0.988 | 0 | 0 | RPS3A |
| 0 | 0.28333774 | 0.997 | 0.981 | 0 | 0 | RPL17 |
| 0 | 0.28122095 | 0.853 | 0.592 | 0 | 0 | TIMP1 |
| 0 | 0.27863426 | 1 | 0.999 | 0 | 0 | RPLP1 |
| 0 | 0.27862068 | 0.999 | 0.995 | 0 | 0 | RPL37 |
| 0 | 0.27787702 | 0.999 | 0.992 | 0 | 0 | RPL27A |
| 0 | 0.27592726 | 0.995 | 0.982 | 0 | 0 | RPS25 |
| 0 | 0.27340479 | 0.994 | 0.964 | 0 | 0 | PFDN5 |
| 0 | 0.27115319 | 0.998 | 0.986 | 0 | 0 | RPL15 |
| 0 | 0.26877798 | 0.994 | 0.969 | 0 | 0 | RPL22 |
| 0 | 0.26377321 | 1 | 1 | 0 | 0 | TMSB4X |
| 0 | 0.25530192 | 0.993 | 0.967 | 0 | 0 | HLA-C |
| 0 | 0.25331312 | 0.999 | 0.99 | 0 | 0 | RPS16 |
| 1.99E-305 | 0.33637251 | 0.353 | 0.191 | 4.83E-301 | 0 | SDS |
| 1.72E-304 | 0.26368958 | 0.99 | 0.954 | 4.18E-300 | 0 | RPL27 |
| 2.20E-303 | 0.33494208 | 0.542 | 0.365 | 5.34E-299 | 0 | SNX10 |
| 1.62E-302 | 0.49091963 | 0.671 | 0.499 | 3.93E-298 | 0 | AC253572.2 |
| 3.81E-298 | 0.3043891 | 0.623 | 0.438 | 9.25E-294 | 0 | FGL2 |
| 3.62E-297 | 0.26926571 | 0.995 | 0.975 | 8.77E-293 | 0 | RPL3 |
| 3.89E-297 | 0.36171038 | 0.797 | 0.654 | 9.43E-293 | 0 | CHMP1B |
| 2.05E-295 | 0.43106857 | 0.637 | 0.475 | 4.96E-291 | 0 | ZNF331 |
| 9.62E-288 | 0.28460076 | 0.379 | 0.217 | 2.33E-283 | 0 | CD69 |
| 1.25E-284 | 0.31858173 | 0.976 | 0.948 | 3.02E-280 | 0 | MT-ND4L |
| 4.80E-284 | 0.34883103 | 0.394 | 0.237 | 1.16E-279 | 0 | MIR155HG |
| 1.34E-277 | 0.26971618 | 0.947 | 0.859 | 3.25E-273 | 0 | ATP5MG |
| 2.86E-276 | 0.27182991 | 0.944 | 0.849 | 6.94E-272 | 0 | MAP3K8 |
| 1.87E-272 | 0.30298001 | 0.792 | 0.635 | 4.53E-268 | 0 | MXD1 |
| 3.63E-272 | 0.35730219 | 0.703 | 0.564 | 8.80E-268 | 0 | TNFAIP8 |
| 5.36E-271 | 0.28589279 | 0.932 | 0.847 | 1.30E-266 | 0 | TOMM7 |
| 8.27E-270 | 0.26457584 | 0.965 | 0.911 | 2.01E-265 | 0 | RPLP0 |
| 1.57E-267 | 0.27255992 | 0.838 | 0.676 | 3.80E-263 | 0 | GSTP1 |
| 1.67E-262 | 0.30289493 | 0.312 | 0.17 | 4.05E-258 | 0 | HLA-DQA2 |
| 7.22E-262 | 0.27407996 | 0.92 | 0.841 | 1.75E-257 | 0 | EEF1D |
| 5.23E-252 | 0.31400489 | 0.68 | 0.526 | 1.27E-247 | 0 | CLEC2B |
| 2.90E-251 | 0.30055733 | 0.621 | 0.462 | 7.02E-247 | 0 | LIMS1 |
| 9.35E-250 | 0.29502889 | 0.711 | 0.573 | 2.27E-245 | 0 | NDUFB2 |
| 1.38E-248 | 0.29678595 | 0.613 | 0.449 | 3.33E-244 | 0 | FAM107B |
| 3.69E-247 | 0.28108347 | 0.738 | 0.552 | 8.94E-243 | 0 | PTGS2 |

| | | | | | | |
|---|---|---|---|---|---|---|
| 4.03E-245 | 0.32110473 | 0.453 | 0.305 | 9.76E-241 | 0 | GRASP |
| 2.78E-244 | 0.33114827 | 0.739 | 0.61 | 6.74E-240 | 0 | LDHA |
| 5.43E-242 | 0.32499757 | 0.71 | 0.54 | 1.32E-237 | 0 | RGCC |
| 4.57E-241 | 0.30335028 | 0.673 | 0.524 | 1.11E-236 | 0 | TAGAP |
| 3.64E-240 | 0.27195207 | 0.537 | 0.366 | 8.83E-236 | 0 | LSP1 |
| 9.82E-240 | 0.26550384 | 0.806 | 0.671 | 2.38E-235 | 0 | EIF3K |
| 1.21E-238 | 0.31191017 | 0.627 | 0.471 | 2.94E-234 | 0 | PHACTR1 |
| 2.26E-237 | 0.30814026 | 0.633 | 0.503 | 5.47E-233 | 0 | PPT1 |
| 6.60E-237 | 0.28143813 | 0.739 | 0.608 | 1.60E-232 | 0 | PSME1 |
| 1.27E-231 | 0.26524699 | 0.756 | 0.625 | 3.08E-227 | 0 | COX7B |
| 1.05E-230 | 0.25448675 | 0.984 | 0.953 | 2.54E-226 | 0 | LAPTM5 |
| 5.79E-229 | 0.26881358 | 0.815 | 0.694 | 1.40E-224 | 0 | ATP6V1F |
| 6.46E-228 | 0.29264496 | 0.576 | 0.437 | 1.57E-223 | 0 | CIB1 |
| 5.75E-227 | 0.2871258 | 0.542 | 0.391 | 1.39E-222 | 0 | STX11 |
| 6.82E-226 | 0.29326791 | 0.6 | 0.454 | 1.65E-221 | 0 | PSME2 |
| 2.75E-222 | 0.2555414 | 0.968 | 0.919 | 6.67E-218 | 0 | PFN1 |
| 5.96E-222 | 0.28975283 | 0.437 | 0.286 | 1.45E-217 | 0 | CYTOR |
| 1.68E-217 | 0.54017054 | 0.963 | 0.941 | 4.06E-213 | 0 | CCL4 |
| 1.20E-214 | 0.30064488 | 0.423 | 0.279 | 2.91E-210 | 0 | BIRC3 |
| 1.05E-213 | 0.27839563 | 0.417 | 0.275 | 2.54E-209 | 0 | VAMP5 |
| 1.74E-213 | 0.29148891 | 0.658 | 0.517 | 4.22E-209 | 0 | NFKB1 |
| 3.76E-212 | 0.27066992 | 0.728 | 0.591 | 9.11E-208 | 0 | PDE4B |
| 1.93E-211 | 0.27015239 | 0.744 | 0.606 | 4.68E-207 | 0 | FOSL2 |
| 5.80E-207 | 0.28130753 | 0.731 | 0.605 | 1.41E-202 | 0 | LITAF |
| 1.33E-205 | 0.25917498 | 0.39 | 0.251 | 3.23E-201 | 0 | C3 |
| 1.90E-203 | 0.34156159 | 0.56 | 0.448 | 4.61E-199 | 0 | CALHM6 |
| 3.45E-203 | 0.27295106 | 0.53 | 0.382 | 8.36E-199 | 0 | LTA4H |
| 1.74E-200 | 0.2728637 | 0.639 | 0.512 | 4.23E-196 | 0 | PLXDC2 |
| 1.50E-191 | 0.30735946 | 0.55 | 0.425 | 3.64E-187 | 0 | PFKFB3 |
| 2.43E-191 | 0.26389426 | 0.768 | 0.659 | 5.90E-187 | 0 | DSE |
| 2.96E-180 | 0.29891949 | 0.757 | 0.671 | 7.19E-176 | 0 | CFLAR |
| 2.28E-177 | 0.2508646 | 0.775 | 0.68 | 5.53E-173 | 0 | MBNL1 |
| 7.54E-177 | 0.25811378 | 0.744 | 0.645 | 1.83E-172 | 0 | LCP2 |
| 1.37E-176 | 0.26041894 | 0.903 | 0.823 | 3.32E-172 | 0 | SGK1 |
| 4.23E-171 | 0.25898386 | 0.605 | 0.484 | 1.03E-166 | 0 | RIPK2 |
| 1.95E-157 | 0.27437701 | 0.651 | 0.524 | 4.72E-153 | 0 | LMNA |
| 5.80E-157 | 0.25086257 | 0.499 | 0.383 | 1.41E-152 | 0 | JPT1 |
| 1.12E-151 | 0.2538675 | 0.474 | 0.351 | 2.72E-147 | 0 | HLA-DRB5 |
| 2.41E-150 | 0.2514029 | 0.818 | 0.728 | 5.85E-146 | 0 | ATF3 |
| 1.59E-148 | 0.291821 | 0.577 | 0.475 | 3.86E-144 | 0 | MAP3K2 |
| 8.40E-140 | 0.28613225 | 0.8 | 0.71 | 2.04E-135 | 0 | PMAIP1 |
| 2.99E-120 | 0.28842524 | 0.721 | 0.625 | 7.24E-116 | 0 | AC103591.3 |
| 5.15E-117 | 0.27614348 | 0.409 | 0.308 | 1.25E-112 | 0 | AL355075.4 |

| | | | | | | |
|---|---|---|---|---|---|---|
| 2.06E-107 | 0.31403818 | 0.394 | 0.3 | 4.98E-103 | 0 | AC245014.3 |
| 9.54E-76 | 0.33296228 | 0.253 | 0.184 | 2.31E-71 | 0 | IFIT3 |
| 9.21E-59 | 0.52253441 | 0.349 | 0.289 | 2.23E-54 | 0 | IFIT2 |
| 0 | 3.43165095 | 0.999 | 0.262 | 0 | 1 | SELENOP |
| 0 | 3.22945248 | 0.997 | 0.353 | 0 | 1 | RNASE1 |
| 0 | 3.15797383 | 0.99 | 0.237 | 0 | 1 | F13A1 |
| 0 | 3.03110457 | 0.996 | 0.349 | 0 | 1 | DAB2 |
| 0 | 2.97635492 | 0.969 | 0.226 | 0 | 1 | PLTP |
| 0 | 2.94646971 | 0.968 | 0.097 | 0 | 1 | LYVE1 |
| 0 | 2.87425168 | 0.991 | 0.239 | 0 | 1 | MAF |
| 0 | 2.6921101 | 0.954 | 0.123 | 0 | 1 | MRC1 |
| 0 | 2.5003707 | 0.999 | 0.577 | 0 | 1 | C1QA |
| 0 | 2.45916114 | 0.99 | 0.41 | 0 | 1 | LGMN |
| 0 | 2.35042703 | 0.991 | 0.393 | 0 | 1 | C1QC |
| 0 | 2.32012608 | 0.974 | 0.288 | 0 | 1 | STAB1 |
| 0 | 2.31754433 | 0.955 | 0.174 | 0 | 1 | FOLR2 |
| 0 | 2.28554179 | 0.999 | 0.944 | 0 | 1 | JUN |
| 0 | 2.27207105 | 0.994 | 0.483 | 0 | 1 | C1QB |
| 0 | 2.2589374 | 0.983 | 0.756 | 0 | 1 | KLF2 |
| 0 | 2.14762411 | 0.978 | 0.57 | 0 | 1 | RHOB |
| 0 | 2.14151978 | 0.748 | 0.083 | 0 | 1 | DDX3Y |
| 0 | 2.0210778 | 0.757 | 0.152 | 0 | 1 | CCL2 |
| 0 | 2.01759988 | 0.509 | 0.046 | 0 | 1 | CCL8 |
| 0 | 1.95094988 | 0.696 | 0.142 | 0 | 1 | PDK4 |
| 0 | 1.94577692 | 0.936 | 0.277 | 0 | 1 | MS4A4A |
| 0 | 1.87423555 | 0.992 | 0.744 | 0 | 1 | HSPA1B |
| 0 | 1.87106771 | 0.983 | 0.645 | 0 | 1 | CTSC |
| 0 | 1.83398632 | 0.979 | 0.699 | 0 | 1 | HSPB1 |
| 0 | 1.83074448 | 0.966 | 0.799 | 0 | 1 | HSPA5 |
| 0 | 1.82730517 | 0.925 | 0.346 | 0 | 1 | CD163 |
| 0 | 1.71571031 | 0.93 | 0.448 | 0 | 1 | CSF1R |
| 0 | 1.70908982 | 0.88 | 0.195 | 0 | 1 | VSIG4 |
| 0 | 1.70815507 | 0.973 | 0.75 | 0 | 1 | MAFB |
| 0 | 1.68215158 | 0.782 | 0.2 | 0 | 1 | NEU1 |
| 0 | 1.64318351 | 0.801 | 0.361 | 0 | 1 | CITED2 |
| 0 | 1.6271877 | 0.946 | 0.494 | 0 | 1 | KCTD12 |
| 0 | 1.62471819 | 0.912 | 0.502 | 0 | 1 | BTG2 |
| 0 | 1.59014623 | 0.992 | 0.837 | 0 | 1 | CD14 |
| 0 | 1.57378997 | 0.906 | 0.544 | 0 | 1 | HSPH1 |
| 0 | 1.56995275 | 0.997 | 0.952 | 0 | 1 | ITM2B |
| 0 | 1.55569729 | 0.999 | 0.896 | 0 | 1 | HSPA1A |
| 0 | 1.55374321 | 0.822 | 0.126 | 0 | 1 | GPR34 |
| 0 | 1.54282046 | 0.799 | 0.09 | 0 | 1 | NRP1 |

| | | | | | | |
|---|---|---|---|---|---|---|
| 0 | 1.53682833 | 0.895 | 0.512 | 0 | 1 | CD36 |
| 0 | 1.53051528 | 0.875 | 0.579 | 0 | 1 | EGR1 |
| 0 | 1.51794224 | 0.963 | 0.805 | 0 | 1 | TRA2B |
| 0 | 1.51001575 | 0.876 | 0.486 | 0 | 1 | ARRDC3 |
| 0 | 1.50794718 | 0.999 | 0.976 | 0 | 1 | FOS |
| 0 | 1.4928438 | 0.843 | 0.182 | 0 | 1 | SLCO2B1 |
| 0 | 1.48330575 | 0.801 | 0.145 | 0 | 1 | PMP22 |
| 0 | 1.46847473 | 0.778 | 0.052 | 0 | 1 | COLEC12 |
| 0 | 1.4641757 | 0.86 | 0.247 | 0 | 1 | LPAR6 |
| 0 | 1.44958204 | 0.814 | 0.169 | 0 | 1 | ITPR2 |
| 0 | 1.43325557 | 0.862 | 0.185 | 0 | 1 | FRMD4B |
| 0 | 1.42579136 | 0.951 | 0.745 | 0 | 1 | DNAJA1 |
| 0 | 1.42236634 | 0.898 | 0.336 | 0 | 1 | ADAP2 |
| 0 | 1.41382301 | 0.987 | 0.858 | 0 | 1 | FCGRT |
| 0 | 1.41316298 | 0.794 | 0.168 | 0 | 1 | ATP1B1 |
| 0 | 1.38609778 | 0.737 | 0.037 | 0 | 1 | SCN9A |
| 0 | 1.37323289 | 0.819 | 0.484 | 0 | 1 | IFRD1 |
| 0 | 1.36562267 | 0.906 | 0.388 | 0 | 1 | SNX6 |
| 0 | 1.35192528 | 0.76 | 0.105 | 0 | 1 | WWP1 |
| 0 | 1.34992581 | 0.745 | 0.045 | 0 | 1 | EGFL7 |
| 0 | 1.34659271 | 0.915 | 0.404 | 0 | 1 | MEF2C |
| 0 | 1.33955773 | 0.742 | 0.05 | 0 | 1 | LILRB5 |
| 0 | 1.33769344 | 0.765 | 0.21 | 0 | 1 | A2M |
| 0 | 1.32775584 | 0.916 | 0.713 | 0 | 1 | HSP90B1 |
| 0 | 1.32153221 | 0.758 | 0.082 | 0 | 1 | ITSN1 |
| 0 | 1.29457607 | 0.715 | 0.177 | 0 | 1 | DNAJB4 |
| 0 | 1.2767243 | 0.919 | 0.612 | 0 | 1 | HERPUD1 |
| 0 | 1.27604588 | 0.912 | 0.485 | 0 | 1 | RBPJ |
| 0 | 1.27277465 | 0.817 | 0.219 | 0 | 1 | MTSS1 |
| 0 | 1.26760081 | 0.725 | 0.171 | 0 | 1 | MAN1A1 |
| 0 | 1.26376249 | 0.855 | 0.309 | 0 | 1 | PLD3 |
| 0 | 1.25294366 | 0.948 | 0.902 | 0 | 1 | KLF4 |
| 0 | 1.24874127 | 0.773 | 0.281 | 0 | 1 | CPEB4 |
| 0 | 1.23548767 | 0.749 | 0.1 | 0 | 1 | GAS6 |
| 0 | 1.22717777 | 0.719 | 0.064 | 0 | 1 | FSCN1 |
| 0 | 1.2116264 | 1 | 0.986 | 0 | 1 | HSP90AA1 |
| 0 | 1.21154934 | 0.988 | 0.994 | 0 | 1 | KLF6 |
| 0 | 1.20341026 | 0.625 | 0.064 | 0 | 1 | SLC40A1 |
| 0 | 1.19868709 | 0.69 | 0.043 | 0 | 1 | HPGDS |
| 0 | 1.19477578 | 0.639 | 0.223 | 0 | 1 | BAG3 |
| 0 | 1.19422558 | 0.744 | 0.11 | 0 | 1 | SLC9A9 |
| 0 | 1.18784564 | 0.741 | 0.32 | 0 | 1 | NRP2 |
| 0 | 1.18773878 | 0.795 | 0.398 | 0 | 1 | NUFIP2 |

| | | | | | | |
|---|---|---|---|---|---|---|
| 0 | 1.18314839 | 0.9 | 0.651 | 0 | 1 | SRSF7 |
| 0 | 1.18206312 | 0.971 | 0.794 | 0 | 1 | DNAJB1 |
| 0 | 1.16430125 | 0.744 | 0.225 | 0 | 1 | GOLGA4 |
| 0 | 1.16328559 | 0.952 | 0.751 | 0 | 1 | SPP1 |
| 0 | 1.16216599 | 0.763 | 0.331 | 0 | 1 | TLR4 |
| 0 | 1.15096042 | 0.761 | 0.149 | 0 | 1 | MKNK1 |
| 0 | 1.14655788 | 0.695 | 0.174 | 0 | 1 | MYLIP |
| 0 | 1.1451189 | 0.788 | 0.242 | 0 | 1 | C3AR1 |
| 0 | 1.14495929 | 0.948 | 0.747 | 0 | 1 | MS4A7 |
| 0 | 1.14289357 | 0.919 | 0.785 | 0 | 1 | IER5 |
| 0 | 1.13802813 | 0.714 | 0.12 | 0 | 1 | C2 |
| 0 | 1.13013135 | 0.728 | 0.136 | 0 | 1 | SIGLEC1 |
| 0 | 1.12662825 | 0.974 | 0.901 | 0 | 1 | IER2 |
| 0 | 1.12618329 | 0.624 | 0.039 | 0 | 1 | ENPP2 |
| 0 | 1.11740655 | 0.885 | 0.671 | 0 | 1 | HSPD1 |
| 0 | 1.1096522 | 0.868 | 0.708 | 0 | 1 | RGS2 |
| 0 | 1.10812768 | 0.955 | 0.74 | 0 | 1 | FCGR2A |
| 0 | 1.1020996 | 0.625 | 0.044 | 0 | 1 | IGF1 |
| 0 | 1.10172937 | 0.703 | 0.132 | 0 | 1 | CD59 |
| 0 | 1.10066231 | 0.317 | 0.029 | 0 | 1 | FCGBP |
| 0 | 1.09827436 | 0.396 | 0.024 | 0 | 1 | RND3 |
| 0 | 1.0982443 | 0.856 | 0.728 | 0 | 1 | OTUD1 |
| 0 | 1.09075008 | 0.72 | 0.25 | 0 | 1 | CLTC |
| 0 | 1.08120545 | 0.995 | 0.951 | 0 | 1 | ZFP36L1 |
| 0 | 1.05603917 | 0.725 | 0.206 | 0 | 1 | GYPC |
| 0 | 1.03729465 | 0.631 | 0.033 | 0 | 1 | CD28 |
| 0 | 1.03182189 | 0.682 | 0.152 | 0 | 1 | PYGL |
| 0 | 1.02486572 | 0.634 | 0.091 | 0 | 1 | FUCA1 |
| 0 | 1.02330602 | 0.623 | 0.077 | 0 | 1 | GATM |
| 0 | 1.01892211 | 0.61 | 0.11 | 0 | 1 | NFIA |
| 0 | 1.00056928 | 0.624 | 0.235 | 0 | 1 | DDIT3 |
| 0 | 0.99538882 | 0.93 | 0.835 | 0 | 1 | HNRNPU |
| 0 | 0.98762063 | 0.639 | 0.095 | 0 | 1 | SPRED1 |
| 0 | 0.98123312 | 0.836 | 0.426 | 0 | 1 | TMEM176B |
| 0 | 0.97886435 | 0.52 | 0.191 | 0 | 1 | AC091271.1 |
| 0 | 0.97442074 | 0.874 | 0.519 | 0 | 1 | WSB1 |
| 0 | 0.97287349 | 0.994 | 0.956 | 0 | 1 | HSP90AB1 |
| 0 | 0.96232687 | 0.815 | 0.431 | 0 | 1 | CREG1 |
| 0 | 0.95736083 | 0.85 | 0.451 | 0 | 1 | IFI16 |
| 0 | 0.95553956 | 0.935 | 0.89 | 0 | 1 | VMP1 |
| 0 | 0.95400601 | 0.957 | 0.91 | 0 | 1 | CCNL1 |
| 0 | 0.95266561 | 0.482 | 0.175 | 0 | 1 | ZBTB10 |
| 0 | 0.94468378 | 0.977 | 0.912 | 0 | 1 | UBB |

| | | | | | | |
|---|---|---|---|---|---|---|
| 0 | 0.94458406 | 0.935 | 0.692 | 0 | 1 | EIF4A2 |
| 0 | 0.94106876 | 0.718 | 0.363 | 0 | 1 | NABP1 |
| 0 | 0.94089532 | 0.997 | 0.99 | 0 | 1 | DDX5 |
| 0 | 0.94043393 | 0.761 | 0.238 | 0 | 1 | NCF4 |
| 0 | 0.93455688 | 0.65 | 0.228 | 0 | 1 | DNAJB11 |
| 0 | 0.92660253 | 0.596 | 0.07 | 0 | 1 | RGL1 |
| 0 | 0.92420788 | 0.669 | 0.13 | 0 | 1 | FCHSD2 |
| 0 | 0.92402845 | 0.539 | 0.028 | 0 | 1 | CD163L1 |
| 0 | 0.92015268 | 0.564 | 0.088 | 0 | 1 | CR1 |
| 0 | 0.91784327 | 0.919 | 0.827 | 0 | 1 | DDX3X |
| 0 | 0.91771131 | 0.675 | 0.173 | 0 | 1 | MCOLN1 |
| 0 | 0.91430379 | 0.98 | 0.942 | 0 | 1 | JUNB |
| 0 | 0.90638047 | 0.348 | 0.045 | 0 | 1 | CH25H |
| 0 | 0.90545586 | 0.594 | 0.101 | 0 | 1 | SESN1 |
| 0 | 0.90372874 | 0.517 | 0.039 | 0 | 1 | CD209 |
| 0 | 0.90230673 | 0.679 | 0.174 | 0 | 1 | PLXND1 |
| 0 | 0.89590084 | 0.584 | 0.067 | 0 | 1 | OLFML2B |
| 0 | 0.88691328 | 0.972 | 0.98 | 0 | 1 | FOSB |
| 0 | 0.88374642 | 0.681 | 0.179 | 0 | 1 | EPS15 |
| 0 | 0.88180392 | 0.725 | 0.381 | 0 | 1 | TLE4 |
| 0 | 0.87896396 | 0.504 | 0.154 | 0 | 1 | DNAJA4 |
| 0 | 0.87830589 | 0.577 | 0.082 | 0 | 1 | ARHGAP5 |
| 0 | 0.87564761 | 0.748 | 0.425 | 0 | 1 | YME1L1 |
| 0 | 0.86610904 | 0.583 | 0.096 | 0 | 1 | TPM1 |
| 0 | 0.86564806 | 0.723 | 0.236 | 0 | 1 | CYTH4 |
| 0 | 0.8642462 | 0.611 | 0.112 | 0 | 1 | EPS8 |
| 0 | 0.86086095 | 0.877 | 0.533 | 0 | 1 | SON |
| 0 | 0.85973117 | 0.993 | 0.993 | 0 | 1 | DUSP1 |
| 0 | 0.85969854 | 0.89 | 0.697 | 0 | 1 | EIF5 |
| 0 | 0.85815155 | 0.74 | 0.377 | 0 | 1 | FCGR1A |
| 0 | 0.85333365 | 0.991 | 0.96 | 0 | 1 | HNRNPA2B1 |
| 0 | 0.84995294 | 0.827 | 0.478 | 0 | 1 | MFSD1 |
| 0 | 0.84937336 | 0.457 | 0.038 | 0 | 1 | TMIGD3 |
| 0 | 0.8434249 | 0.74 | 0.308 | 0 | 1 | SNX2 |
| 0 | 0.836862 | 0.719 | 0.397 | 0 | 1 | RAP2B |
| 0 | 0.83476071 | 0.541 | 0.092 | 0 | 1 | FAM13A |
| 0 | 0.83107617 | 0.758 | 0.468 | 0 | 1 | AMD1 |
| 0 | 0.82072919 | 1 | 1 | 0 | 1 | MALAT1 |
| 0 | 0.81034281 | 0.872 | 0.671 | 0 | 1 | FUS |
| 0 | 0.80949358 | 0.748 | 0.357 | 0 | 1 | LAMP1 |
| 0 | 0.80919986 | 0.675 | 0.3 | 0 | 1 | PNN |
| 0 | 0.80667324 | 0.951 | 0.795 | 0 | 1 | HSPA8 |
| 0 | 0.80642291 | 0.493 | 0.027 | 0 | 1 | PCDH12 |

| | | | | | | |
|---|---|---|---|---|---|---|
| 0 | 0.80309092 | 0.675 | 0.341 | 0 | 1 | PPP1R10 |
| 0 | 0.8009532 | 0.831 | 0.549 | 0 | 1 | SFPQ |
| 0 | 0.8009439 | 0.984 | 0.965 | 0 | 1 | GADD45B |
| 0 | 0.80085605 | 0.605 | 0.167 | 0 | 1 | GASK1B |
| 0 | 0.79458482 | 0.683 | 0.299 | 0 | 1 | ANKRD12 |
| 0 | 0.79003129 | 0.54 | 0.168 | 0 | 1 | AL499604.1 |
| 0 | 0.78720261 | 0.734 | 0.323 | 0 | 1 | RASSF4 |
| 0 | 0.7834043 | 0.655 | 0.256 | 0 | 1 | ATP6V0A1 |
| 0 | 0.78093011 | 0.713 | 0.297 | 0 | 1 | ARHGAP18 |
| 0 | 0.77505443 | 0.586 | 0.115 | 0 | 1 | SPATS2L |
| 0 | 0.77367252 | 0.689 | 0.291 | 0 | 1 | NAIP |
| 0 | 0.77159551 | 0.492 | 0.106 | 0 | 1 | CEMIP2 |
| 0 | 0.77043741 | 0.65 | 0.239 | 0 | 1 | ADAM9 |
| 0 | 0.76796657 | 0.731 | 0.389 | 0 | 1 | OGFRL1 |
| 0 | 0.7614261 | 0.651 | 0.243 | 0 | 1 | 1-Mar |
| 0 | 0.75701307 | 0.763 | 0.401 | 0 | 1 | ARGLU1 |
| 0 | 0.75387056 | 0.646 | 0.293 | 0 | 1 | CHORDC1 |
| 0 | 0.74875715 | 0.526 | 0.094 | 0 | 1 | TM2D2 |
| 0 | 0.74490544 | 0.437 | 0.02 | 0 | 1 | NAV3 |
| 0 | 0.73990604 | 0.608 | 0.185 | 0 | 1 | PIK3R1 |
| 0 | 0.73749626 | 0.801 | 0.539 | 0 | 1 | RUNX1 |
| 0 | 0.73706666 | 0.615 | 0.229 | 0 | 1 | ATRX |
| 0 | 0.73490565 | 0.96 | 0.832 | 0 | 1 | MS4A6A |
| 0 | 0.73453978 | 0.664 | 0.275 | 0 | 1 | BMP2K |
| 0 | 0.73394646 | 0.664 | 0.318 | 0 | 1 | TCP1 |
| 0 | 0.73310046 | 0.491 | 0.062 | 0 | 1 | FAM20A |
| 0 | 0.7323343 | 0.885 | 0.757 | 0 | 1 | CYBB |
| 0 | 0.73189329 | 0.578 | 0.183 | 0 | 1 | ATM |
| 0 | 0.73180769 | 0.62 | 0.221 | 0 | 1 | CYFIP1 |
| 0 | 0.72956177 | 0.419 | 0.036 | 0 | 1 | CCND1 |
| 0 | 0.72262642 | 0.833 | 0.605 | 0 | 1 | CLK1 |
| 0 | 0.72043297 | 0.928 | 0.828 | 0 | 1 | CEBPD |
| 0 | 0.71952557 | 0.657 | 0.266 | 0 | 1 | TMEM176A |
| 0 | 0.71715107 | 0.648 | 0.235 | 0 | 1 | RRBP1 |
| 0 | 0.71639521 | 0.686 | 0.307 | 0 | 1 | MPEG1 |
| 0 | 0.71551947 | 0.452 | 0.029 | 0 | 1 | IGFBP4 |
| 0 | 0.70986941 | 0.652 | 0.246 | 0 | 1 | CCR1 |
| 0 | 0.70798199 | 0.294 | 0.147 | 0 | 1 | CCL5 |
| 0 | 0.70563222 | 0.445 | 0.124 | 0 | 1 | BHLHE41 |
| 0 | 0.70487455 | 0.521 | 0.344 | 0 | 1 | TFRC |
| 0 | 0.70273877 | 0.452 | 0.05 | 0 | 1 | SLC7A8 |
| 0 | 0.70161525 | 0.626 | 0.31 | 0 | 1 | ZFAND2A |
| 0 | 0.70151253 | 0.71 | 0.361 | 0 | 1 | GNG10 |

| | | | | | | |
|---|---|---|---|---|---|---|
| 0 | 0.70092435 | 0.502 | 0.079 | 0 | 1 | RPS4Y1 |
| 0 | 0.69902473 | 0.473 | 0.052 | 0 | 1 | PDGFC |
| 0 | 0.69687567 | 0.746 | 0.422 | 0 | 1 | CLTA |
| 0 | 0.69573214 | 0.937 | 0.777 | 0 | 1 | CALM2 |
| 0 | 0.69310924 | 1 | 1 | 0 | 1 | MT-CYB |
| 0 | 0.69264941 | 0.536 | 0.13 | 0 | 1 | SGMS1 |
| 0 | 0.68923024 | 0.875 | 0.635 | 0 | 1 | TUBA1B |
| 0 | 0.68214611 | 1 | 1 | 0 | 1 | MT-ATP6 |
| 0 | 0.681606 | 0.726 | 0.395 | 0 | 1 | TACC1 |
| 0 | 0.67951094 | 0.718 | 0.379 | 0 | 1 | N4BP2L2 |
| 0 | 0.67642365 | 0.632 | 0.314 | 0 | 1 | BRD2 |
| 0 | 0.67622791 | 0.625 | 0.218 | 0 | 1 | AP1B1 |
| 0 | 0.67482448 | 0.497 | 0.271 | 0 | 1 | P2RY13 |
| 0 | 0.67381198 | 0.461 | 0.026 | 0 | 1 | TNFRSF25 |
| 0 | 0.6723714 | 0.579 | 0.182 | 0 | 1 | TTYH3 |
| 0 | 0.6719929 | 0.502 | 0.093 | 0 | 1 | PLA2G15 |
| 0 | 0.67039275 | 0.789 | 0.565 | 0 | 1 | ZFP36L2 |
| 0 | 0.66791098 | 0.58 | 0.157 | 0 | 1 | AP2A2 |
| 0 | 0.66315246 | 0.701 | 0.424 | 0 | 1 | DDX24 |
| 0 | 0.66172016 | 0.539 | 0.127 | 0 | 1 | GNG2 |
| 0 | 0.66156225 | 0.424 | 0.041 | 0 | 1 | HRH1 |
| 0 | 0.66051084 | 0.802 | 0.547 | 0 | 1 | HNRNPH1 |
| 0 | 0.66048667 | 0.657 | 0.29 | 0 | 1 | RB1 |
| 0 | 0.65991419 | 0.558 | 0.19 | 0 | 1 | MPHOSPH8 |
| 0 | 0.65960363 | 0.561 | 0.172 | 0 | 1 | STOM |
| 0 | 0.65825465 | 0.501 | 0.102 | 0 | 1 | BIN1 |
| 0 | 0.65779157 | 0.742 | 0.43 | 0 | 1 | PNISR |
| 0 | 0.65113293 | 0.688 | 0.341 | 0 | 1 | ALOX5AP |
| 0 | 0.65089566 | 0.555 | 0.195 | 0 | 1 | ETNK1 |
| 0 | 0.6508723 | 0.674 | 0.34 | 0 | 1 | LRP1 |
| 0 | 0.64970434 | 0.491 | 0.069 | 0 | 1 | TBC1D14 |
| 0 | 0.64429572 | 0.617 | 0.222 | 0 | 1 | COLGALT1 |
| 0 | 0.64362039 | 0.44 | 0.069 | 0 | 1 | GIMAP5 |
| 0 | 0.64229962 | 0.433 | 0.083 | 0 | 1 | CCDC144A |
| 0 | 0.64003452 | 0.982 | 0.936 | 0 | 1 | CTSB |
| 0 | 0.63620133 | 0.704 | 0.391 | 0 | 1 | RSRP1 |
| 0 | 0.63592779 | 0.642 | 0.27 | 0 | 1 | TNFRSF1A |
| 0 | 0.63384125 | 0.484 | 0.083 | 0 | 1 | FRMD4A |
| 0 | 0.6291296 | 0.574 | 0.21 | 0 | 1 | STX7 |
| 0 | 0.62649613 | 0.666 | 0.308 | 0 | 1 | SERPINB6 |
| 0 | 0.62640019 | 0.765 | 0.635 | 0 | 1 | TSC22D3 |
| 0 | 0.62568391 | 0.574 | 0.235 | 0 | 1 | MACF1 |
| 0 | 0.62505878 | 0.449 | 0.033 | 0 | 1 | RAB3IL1 |

| | | | | | | |
|---|---|---|---|---|---|---|
| 0 | 0.62492629 | 0.525 | 0.229 | 0 | 1 | IER5L |
| 0 | 0.62291386 | 0.514 | 0.204 | 0 | 1 | EMB |
| 0 | 0.62235229 | 0.643 | 0.318 | 0 | 1 | NUCKS1 |
| 0 | 0.62136883 | 0.665 | 0.397 | 0 | 1 | AFF4 |
| 0 | 0.62083263 | 0.615 | 0.223 | 0 | 1 | ARHGAP4 |
| 0 | 0.6203075 | 0.796 | 0.589 | 0 | 1 | HMOX1 |
| 0 | 0.61688928 | 0.592 | 0.224 | 0 | 1 | NASP |
| 0 | 0.61623784 | 0.655 | 0.337 | 0 | 1 | PRPF38B |
| 0 | 0.61524236 | 0.706 | 0.393 | 0 | 1 | LAIR1 |
| 0 | 0.61388798 | 0.579 | 0.221 | 0 | 1 | DCTN4 |
| 0 | 0.61298841 | 0.437 | 0.058 | 0 | 1 | PEAK1 |
| 0 | 0.61209744 | 0.602 | 0.253 | 0 | 1 | NORAD |
| 0 | 0.60867768 | 0.675 | 0.359 | 0 | 1 | ZFHX3 |
| 0 | 0.60741955 | 0.377 | 0.019 | 0 | 1 | LARGE1 |
| 0 | 0.60733461 | 0.448 | 0.052 | 0 | 1 | SCN1B |
| 0 | 0.60669935 | 0.44 | 0.043 | 0 | 1 | FGFR1 |
| 0 | 0.60399167 | 0.521 | 0.149 | 0 | 1 | CCDC50 |
| 0 | 0.60239168 | 0.492 | 0.147 | 0 | 1 | TCF4 |
| 0 | 0.60050037 | 0.572 | 0.224 | 0 | 1 | NPL |
| 0 | 0.59930718 | 0.592 | 0.25 | 0 | 1 | NKTR |
| 0 | 0.59877416 | 0.431 | 0.055 | 0 | 1 | WLS |
| 0 | 0.59843235 | 0.461 | 0.087 | 0 | 1 | DIP2A |
| 0 | 0.59666797 | 0.522 | 0.166 | 0 | 1 | TGFBR2 |
| 0 | 0.59573163 | 0.372 | 0.019 | 0 | 1 | FGF13 |
| 0 | 0.59558682 | 0.81 | 0.6 | 0 | 1 | XBP1 |
| 0 | 0.59522434 | 0.463 | 0.11 | 0 | 1 | IDH1 |
| 0 | 0.59396537 | 0.451 | 0.103 | 0 | 1 | ABHD3 |
| 0 | 0.59370641 | 0.471 | 0.102 | 0 | 1 | FCHO2 |
| 0 | 0.59266417 | 0.513 | 0.187 | 0 | 1 | CHCHD7 |
| 0 | 0.5905564 | 0.416 | 0.024 | 0 | 1 | PLEKHG5 |
| 0 | 0.59031747 | 0.432 | 0.06 | 0 | 1 | ATP8B4 |
| 0 | 0.58882849 | 0.509 | 0.146 | 0 | 1 | NFIC |
| 0 | 0.58823811 | 0.384 | 0.075 | 0 | 1 | HHEX |
| 0 | 0.58822471 | 0.399 | 0.025 | 0 | 1 | CD200R1 |
| 0 | 0.58674325 | 0.614 | 0.274 | 0 | 1 | RBMS1 |
| 0 | 0.58609959 | 0.503 | 0.174 | 0 | 1 | FCGR1B |
| 0 | 0.58605147 | 0.41 | 0.048 | 0 | 1 | SLC18B1 |
| 0 | 0.58567086 | 0.423 | 0.044 | 0 | 1 | ITGB5 |
| 0 | 0.58397546 | 0.545 | 0.196 | 0 | 1 | PCM1 |
| 0 | 0.58356896 | 0.611 | 0.296 | 0 | 1 | HEXA |
| 0 | 0.58294901 | 0.867 | 0.641 | 0 | 1 | SRRM2 |
| 0 | 0.58230803 | 0.475 | 0.112 | 0 | 1 | TMEM106A |
| 0 | 0.58139424 | 0.566 | 0.269 | 0 | 1 | AKAP9 |

| | | | | | | |
|---|---|---|---|---|---|---|
| 0 | 0.58110945 | 0.57 | 0.239 | 0 | 1 | PEBP1 |
| 0 | 0.57846291 | 0.68 | 0.396 | 0 | 1 | BCLAF1 |
| 0 | 0.57819825 | 0.498 | 0.137 | 0 | 1 | FEZ2 |
| 0 | 0.57719398 | 0.767 | 0.514 | 0 | 1 | LAP3 |
| 0 | 0.57274693 | 0.615 | 0.333 | 0 | 1 | C6orf62 |
| 0 | 0.57052023 | 0.479 | 0.173 | 0 | 1 | GOLGB1 |
| 0 | 0.56970428 | 0.391 | 0.097 | 0 | 1 | Z99127.4 |
| 0 | 0.56936299 | 0.447 | 0.09 | 0 | 1 | ST6GAL1 |
| 0 | 0.56792505 | 0.406 | 0.052 | 0 | 1 | CMKLR1 |
| 0 | 0.56687086 | 0.549 | 0.259 | 0 | 1 | IDI1 |
| 0 | 0.56585585 | 0.342 | 0.021 | 0 | 1 | CCDC152 |
| 0 | 0.56557095 | 0.691 | 0.419 | 0 | 1 | NCOA4 |
| 0 | 0.56548617 | 0.381 | 0.031 | 0 | 1 | MTUS1 |
| 0 | 0.56545473 | 0.547 | 0.22 | 0 | 1 | DOK2 |
| 0 | 0.56510162 | 0.999 | 0.999 | 0 | 1 | H3F3B |
| 0 | 0.56508406 | 0.523 | 0.222 | 0 | 1 | LUC7L3 |
| 0 | 0.56347749 | 0.48 | 0.121 | 0 | 1 | WDFY3 |
| 0 | 0.56247131 | 0.869 | 0.665 | 0 | 1 | RNASET2 |
| 0 | 0.56166967 | 0.552 | 0.238 | 0 | 1 | GIMAP4 |
| 0 | 0.56163502 | 0.533 | 0.225 | 0 | 1 | USP36 |
| 0 | 0.55756831 | 0.387 | 0.131 | 0 | 1 | TSPYL2 |
| 0 | 0.55707642 | 0.423 | 0.059 | 0 | 1 | ARHGAP12 |
| 0 | 0.55454216 | 0.491 | 0.16 | 0 | 1 | FTX |
| 0 | 0.55416559 | 0.373 | 0.017 | 0 | 1 | FGL1 |
| 0 | 0.55164827 | 0.354 | 0.067 | 0 | 1 | AC022868.2 |
| 0 | 0.5496521 | 0.38 | 0.019 | 0 | 1 | IL12RB2 |
| 0 | 0.54667516 | 0.801 | 0.562 | 0 | 1 | DDX17 |
| 0 | 0.54574424 | 0.576 | 0.281 | 0 | 1 | CREM |
| 0 | 0.54411373 | 0.685 | 0.441 | 0 | 1 | ELF1 |
| 0 | 0.542882 | 0.578 | 0.251 | 0 | 1 | UNC93B1 |
| 0 | 0.54032245 | 0.454 | 0.129 | 0 | 1 | MLEC |
| 0 | 0.54008257 | 0.621 | 0.32 | 0 | 1 | RNF13 |
| 0 | 0.53908983 | 0.611 | 0.324 | 0 | 1 | SNHG32 |
| 0 | 0.53724021 | 0.43 | 0.105 | 0 | 1 | TLR7 |
| 0 | 0.5366985 | 0.365 | 0.027 | 0 | 1 | AC129507.1 |
| 0 | 0.53619669 | 0.397 | 0.04 | 0 | 1 | FARP1 |
| 0 | 0.53524053 | 0.306 | 0.071 | 0 | 1 | ZNF503 |
| 0 | 0.53287228 | 0.398 | 0.046 | 0 | 1 | RCN3 |
| 0 | 0.53219211 | 0.473 | 0.183 | 0 | 1 | SNHG12 |
| 0 | 0.53215753 | 1 | 1 | 0 | 1 | MT-CO3 |
| 0 | 0.53086842 | 0.47 | 0.154 | 0 | 1 | EIF4G3 |
| 0 | 0.52987184 | 0.493 | 0.288 | 0 | 1 | INTS6 |
| 0 | 0.52968385 | 0.794 | 0.551 | 0 | 1 | CD164 |

| | | | | | | |
|---|---|---|---|---|---|---|
| 0 | 0.52901 | 0.684 | 0.386 | 0 | 1 | HNMT |
| 0 | 0.52874896 | 0.353 | 0.027 | 0 | 1 | KCNJ5 |
| 0 | 0.52832591 | 1 | 1 | 0 | 1 | MT-ND1 |
| 0 | 0.52799203 | 0.446 | 0.111 | 0 | 1 | STMN1 |
| 0 | 0.52632856 | 0.545 | 0.223 | 0 | 1 | LTC4S |
| 0 | 0.52555969 | 0.862 | 0.713 | 0 | 1 | RTN4 |
| 0 | 0.52522983 | 0.493 | 0.219 | 0 | 1 | GCC2 |
| 0 | 0.52456231 | 0.385 | 0.061 | 0 | 1 | SRGAP1 |
| 0 | 0.52454071 | 0.699 | 0.45 | 0 | 1 | RNF149 |
| 0 | 0.52200419 | 0.495 | 0.175 | 0 | 1 | AKR1B1 |
| 0 | 0.52054333 | 0.484 | 0.163 | 0 | 1 | SERPING1 |
| 0 | 0.5196814 | 0.679 | 0.401 | 0 | 1 | SCAF11 |
| 0 | 0.51965653 | 0.53 | 0.218 | 0 | 1 | PCF11 |
| 0 | 0.51948343 | 0.624 | 0.476 | 0 | 1 | TRIB1 |
| 0 | 0.51935377 | 0.551 | 0.251 | 0 | 1 | BOD1L1 |
| 0 | 0.5189833 | 0.667 | 0.406 | 0 | 1 | RBM25 |
| 0 | 0.51838611 | 0.492 | 0.197 | 0 | 1 | POLR2J3 |
| 0 | 0.51838251 | 0.755 | 0.49 | 0 | 1 | CD99 |
| 0 | 0.51822673 | 0.419 | 0.146 | 0 | 1 | HEXIM1 |
| 0 | 0.5178007 | 0.731 | 0.469 | 0 | 1 | SF3B1 |
| 0 | 0.51743079 | 0.344 | 0.021 | 0 | 1 | NAV2 |
| 0 | 0.51737019 | 0.483 | 0.164 | 0 | 1 | PKN2 |
| 0 | 0.51494222 | 0.438 | 0.175 | 0 | 1 | ADRB2 |
| 0 | 0.51468833 | 0.592 | 0.326 | 0 | 1 | CD84 |
| 0 | 0.51368275 | 0.515 | 0.203 | 0 | 1 | SNX5 |
| 0 | 0.51281676 | 0.501 | 0.196 | 0 | 1 | WASHC4 |
| 0 | 0.51223274 | 0.442 | 0.14 | 0 | 1 | TSPYL1 |
| 0 | 0.51028449 | 0.804 | 0.582 | 0 | 1 | CTSL |
| 0 | 0.50989264 | 0.553 | 0.25 | 0 | 1 | WASF2 |
| 0 | 0.50951325 | 0.361 | 0.029 | 0 | 1 | NFATC2 |
| 0 | 0.50926844 | 0.693 | 0.425 | 0 | 1 | RNF213 |
| 0 | 0.50907038 | 0.338 | 0.053 | 0 | 1 | SNHG14 |
| 0 | 0.50670026 | 0.537 | 0.297 | 0 | 1 | POU2F2 |
| 0 | 0.50655705 | 0.675 | 0.42 | 0 | 1 | TGOLN2 |
| 0 | 0.50558217 | 0.439 | 0.133 | 0 | 1 | TSPAN4 |
| 0 | 0.50480005 | 0.647 | 0.38 | 0 | 1 | SF1 |
| 0 | 0.5046042 | 0.444 | 0.143 | 0 | 1 | TNS3 |
| 0 | 0.50368882 | 0.669 | 0.383 | 0 | 1 | SRSF11 |
| 0 | 0.50318243 | 0.382 | 0.048 | 0 | 1 | RIMKLB |
| 0 | 0.50272049 | 0.527 | 0.21 | 0 | 1 | SCAMP2 |
| 0 | 0.50182507 | 0.412 | 0.121 | 0 | 1 | ARID4A |
| 0 | 0.50175455 | 0.62 | 0.321 | 0 | 1 | AAK1 |
| 0 | 0.50044396 | 0.372 | 0.038 | 0 | 1 | TMEM37 |

| | | | | | | |
|---|---|---|---|---|---|---|
| 0 | 0.5003318 | 0.519 | 0.234 | 0 | 1 | PRPF4B |
| 0 | 0.4996616 | 0.829 | 0.656 | 0 | 1 | CALR |
| 0 | 0.4992637 | 0.476 | 0.19 | 0 | 1 | ZNF106 |
| 0 | 0.49882525 | 0.381 | 0.099 | 0 | 1 | GIMAP7 |
| 0 | 0.49830538 | 0.617 | 0.338 | 0 | 1 | STAT3 |
| 0 | 0.49758591 | 0.467 | 0.156 | 0 | 1 | MYO5A |
| 0 | 0.49583356 | 0.477 | 0.197 | 0 | 1 | RESF1 |
| 0 | 0.49569091 | 0.414 | 0.13 | 0 | 1 | BDP1 |
| 0 | 0.49546888 | 0.614 | 0.347 | 0 | 1 | ABCA1 |
| 0 | 0.49438331 | 0.337 | 0.074 | 0 | 1 | NMRK1 |
| 0 | 0.49435241 | 0.886 | 0.769 | 0 | 1 | APLP2 |
| 0 | 0.49410521 | 0.414 | 0.142 | 0 | 1 | TGFBR1 |
| 0 | 0.49400654 | 0.506 | 0.212 | 0 | 1 | BAZ2B |
| 0 | 0.49382631 | 0.383 | 0.067 | 0 | 1 | ETV5 |
| 0 | 0.49252846 | 0.472 | 0.178 | 0 | 1 | SETX |
| 0 | 0.49203953 | 0.509 | 0.25 | 0 | 1 | IQGAP2 |
| 0 | 0.49198889 | 0.826 | 0.625 | 0 | 1 | QKI |
| 0 | 0.48952681 | 0.904 | 0.824 | 0 | 1 | SRSF5 |
| 0 | 0.48852012 | 0.315 | 0.017 | 0 | 1 | NCKAP5 |
| 0 | 0.48708605 | 0.384 | 0.075 | 0 | 1 | ABCC5 |
| 0 | 0.48684465 | 0.587 | 0.291 | 0 | 1 | EPB41L3 |
| 0 | 0.48504924 | 0.5 | 0.208 | 0 | 1 | ZRANB2 |
| 0 | 0.48472637 | 0.748 | 0.535 | 0 | 1 | SNX3 |
| 0 | 0.48421792 | 0.433 | 0.184 | 0 | 1 | KIF1B |
| 0 | 0.48405291 | 0.444 | 0.173 | 0 | 1 | PCBP1-AS1 |
| 0 | 0.48357149 | 0.923 | 0.798 | 0 | 1 | RBM39 |
| 0 | 0.48338731 | 0.488 | 0.281 | 0 | 1 | DYNC1H1 |
| 0 | 0.48198752 | 0.429 | 0.188 | 0 | 1 | THUMPD3-AS[illegible] |
| 0 | 0.48037168 | 0.328 | 0.027 | 0 | 1 | AC110995.1 |
| 0 | 0.48014766 | 0.541 | 0.271 | 0 | 1 | MAT2A |
| 0 | 0.47944416 | 0.735 | 0.516 | 0 | 1 | NCL |
| 0 | 0.47879631 | 0.488 | 0.17 | 0 | 1 | TECR |
| 0 | 0.47868442 | 0.38 | 0.138 | 0 | 1 | FAM53C |
| 0 | 0.47835193 | 0.468 | 0.234 | 0 | 1 | CHD4 |
| 0 | 0.4782174 | 0.453 | 0.183 | 0 | 1 | GALNT1 |
| 0 | 0.47800124 | 0.531 | 0.226 | 0 | 1 | PTMS |
| 0 | 0.4779853 | 0.424 | 0.12 | 0 | 1 | ARID3A |
| 0 | 0.4778757 | 0.373 | 0.077 | 0 | 1 | ZBTB38 |
| 0 | 0.47680929 | 0.461 | 0.159 | 0 | 1 | CAB39 |
| 0 | 0.47643084 | 0.505 | 0.225 | 0 | 1 | PHIP |
| 0 | 0.47594145 | 0.429 | 0.24 | 0 | 1 | TENT5A |
| 0 | 0.47583343 | 0.617 | 0.387 | 0 | 1 | FAM133B |
| 0 | 0.47571556 | 0.354 | 0.085 | 0 | 1 | CDKN2AIP |

| | | | | | | |
|---|---|---|---|---|---|---|
| 0 | 0.4753721 | 0.568 | 0.255 | 0 | 1 | SNRNP70 |
| 0 | 0.47510585 | 0.483 | 0.171 | 0 | 1 | KANSL1 |
| 0 | 0.4739983 | 0.735 | 0.575 | 0 | 1 | CPVL |
| 0 | 0.47221775 | 0.491 | 0.206 | 0 | 1 | TM9SF2 |
| 0 | 0.47211882 | 0.511 | 0.23 | 0 | 1 | IL6ST |
| 0 | 0.4716375 | 0.439 | 0.159 | 0 | 1 | EEA1 |
| 0 | 0.47132076 | 0.431 | 0.125 | 0 | 1 | ATF7IP |
| 0 | 0.46991314 | 0.49 | 0.193 | 0 | 1 | OTULINL |
| 0 | 0.4696335 | 0.484 | 0.24 | 0 | 1 | TRIM8 |
| 0 | 0.46673935 | 0.466 | 0.171 | 0 | 1 | ATF6 |
| 0 | 0.46572457 | 0.776 | 0.593 | 0 | 1 | BLVRB |
| 0 | 0.46570722 | 0.406 | 0.11 | 0 | 1 | DNAJC13 |
| 0 | 0.46546161 | 0.526 | 0.246 | 0 | 1 | ITSN2 |
| 0 | 0.46408342 | 0.396 | 0.108 | 0 | 1 | PIK3IP1 |
| 0 | 0.46368695 | 0.34 | 0.046 | 0 | 1 | PLEKHA1 |
| 0 | 0.46079999 | 0.455 | 0.149 | 0 | 1 | TCF12 |
| 0 | 0.46047901 | 0.406 | 0.134 | 0 | 1 | TTC14 |
| 0 | 0.4597265 | 0.288 | 0.052 | 0 | 1 | PPM1N |
| 0 | 0.45950576 | 0.301 | 0.046 | 0 | 1 | ZNF844 |
| 0 | 0.45913002 | 0.608 | 0.337 | 0 | 1 | NCKAP1L |
| 0 | 0.4586927 | 0.599 | 0.333 | 0 | 1 | HP1BP3 |
| 0 | 0.45863295 | 0.464 | 0.184 | 0 | 1 | ASH1L |
| 0 | 0.45848548 | 0.458 | 0.169 | 0 | 1 | HELZ |
| 0 | 0.45684387 | 0.267 | 0.035 | 0 | 1 | SCIN |
| 0 | 0.45676545 | 0.42 | 0.125 | 0 | 1 | AFF1 |
| 0 | 0.4566036 | 0.382 | 0.107 | 0 | 1 | NCOA7 |
| 0 | 0.45626653 | 0.489 | 0.208 | 0 | 1 | RIN2 |
| 0 | 0.45558936 | 0.361 | 0.113 | 0 | 1 | SCML1 |
| 0 | 0.45483123 | 0.369 | 0.094 | 0 | 1 | PHYH |
| 0 | 0.45435674 | 0.438 | 0.167 | 0 | 1 | TLR1 |
| 0 | 0.45382925 | 0.302 | 0.048 | 0 | 1 | ZNF117 |
| 0 | 0.45326976 | 0.331 | 0.056 | 0 | 1 | CHD7 |
| 0 | 0.45303978 | 0.467 | 0.185 | 0 | 1 | SENP6 |
| 0 | 0.45265732 | 0.499 | 0.209 | 0 | 1 | P4HA1 |
| 0 | 0.4515184 | 1 | 1 | 0 | 1 | MT-ND2 |
| 0 | 0.4501195 | 0.49 | 0.206 | 0 | 1 | SDCCAG8 |
| 0 | 0.44966145 | 0.79 | 0.609 | 0 | 1 | TIMP2 |
| 0 | 0.44927576 | 0.63 | 0.375 | 0 | 1 | TRAM1 |
| 0 | 0.44919711 | 0.569 | 0.3 | 0 | 1 | RNASE6 |
| 0 | 0.44890865 | 0.703 | 0.479 | 0 | 1 | TMBIM4 |
| 0 | 0.44882303 | 0.351 | 0.124 | 0 | 1 | OTUD6B-AS1 |
| 0 | 0.44876367 | 0.348 | 0.082 | 0 | 1 | PAX8-AS1 |
| 0 | 0.4484312 | 0.42 | 0.163 | 0 | 1 | CEP350 |

| | | | | | | |
|---|---|---|---|---|---|---|
| 0 | 0.44703312 | 0.441 | 0.179 | 0 | 1 | LPP |
| 0 | 0.44674026 | 0.337 | 0.037 | 0 | 1 | LPAR5 |
| 0 | 0.44670578 | 0.7 | 0.452 | 0 | 1 | LAPTM4A |
| 0 | 0.44575137 | 0.305 | 0.065 | 0 | 1 | OLMALINC |
| 0 | 0.44563461 | 0.336 | 0.079 | 0 | 1 | VCPIP1 |
| 0 | 0.44497496 | 0.439 | 0.162 | 0 | 1 | TENT2 |
| 0 | 0.44484577 | 0.322 | 0.046 | 0 | 1 | TBC1D4 |
| 0 | 0.44451274 | 0.413 | 0.143 | 0 | 1 | KMT2A |
| 0 | 0.44287156 | 0.439 | 0.251 | 0 | 1 | IRS2 |
| 0 | 0.44119468 | 0.281 | 0.015 | 0 | 1 | SYT1 |
| 0 | 0.44117504 | 0.408 | 0.117 | 0 | 1 | FGD2 |
| 0 | 0.44039429 | 0.283 | 0.113 | 0 | 1 | EGR2 |
| 0 | 0.43985489 | 0.345 | 0.062 | 0 | 1 | SEPTIN11 |
| 0 | 0.43897497 | 0.341 | 0.073 | 0 | 1 | PIAS2 |
| 0 | 0.43873283 | 0.373 | 0.079 | 0 | 1 | NISCH |
| 0 | 0.43826015 | 0.299 | 0.022 | 0 | 1 | DISP1 |
| 0 | 0.4376629 | 0.6 | 0.374 | 0 | 1 | VPS13C |
| 0 | 0.4369812 | 0.409 | 0.163 | 0 | 1 | H1F0 |
| 0 | 0.43622238 | 0.882 | 0.735 | 0 | 1 | HSPE1 |
| 0 | 0.4356965 | 0.45 | 0.223 | 0 | 1 | MIS18BP1 |
| 0 | 0.43500251 | 0.402 | 0.12 | 0 | 1 | TBC1D9 |
| 0 | 0.43427157 | 0.32 | 0.061 | 0 | 1 | DAAM1 |
| 0 | 0.43370351 | 0.462 | 0.189 | 0 | 1 | CTBS |
| 0 | 0.43366403 | 0.558 | 0.286 | 0 | 1 | PJA2 |
| 0 | 0.43353257 | 0.304 | 0.036 | 0 | 1 | TNFRSF11A |
| 0 | 0.43351243 | 0.423 | 0.21 | 0 | 1 | Z93930.2 |
| 0 | 0.43275113 | 0.321 | 0.107 | 0 | 1 | GAS2L3 |
| 0 | 0.43238395 | 1 | 1 | 0 | 1 | TPT1 |
| 0 | 0.43226148 | 0.333 | 0.048 | 0 | 1 | ANKS1A |
| 0 | 0.43196873 | 0.62 | 0.381 | 0 | 1 | CD4 |
| 0 | 0.43190388 | 0.671 | 0.458 | 0 | 1 | RSRC2 |
| 0 | 0.43149259 | 0.449 | 0.2 | 0 | 1 | TPR |
| 0 | 0.43148518 | 0.416 | 0.128 | 0 | 1 | RERE |
| 0 | 0.4311755 | 0.405 | 0.123 | 0 | 1 | ZNF280D |
| 0 | 0.43108105 | 0.433 | 0.148 | 0 | 1 | SNX29 |
| 0 | 0.43095051 | 0.485 | 0.22 | 0 | 1 | CHD9 |
| 0 | 0.43023698 | 0.412 | 0.156 | 0 | 1 | SMARCA2 |
| 0 | 0.43014511 | 0.404 | 0.157 | 0 | 1 | BCAT1 |
| 0 | 0.42952475 | 0.448 | 0.189 | 0 | 1 | CEBPA |
| 0 | 0.42917824 | 0.527 | 0.245 | 0 | 1 | SEC14L1 |
| 0 | 0.42698303 | 0.334 | 0.051 | 0 | 1 | WDR91 |
| 0 | 0.42648773 | 0.449 | 0.188 | 0 | 1 | OSBPL11 |
| 0 | 0.42594225 | 0.357 | 0.081 | 0 | 1 | DIP2B |

| | | | | | | |
|---|---|---|---|---|---|---|
| 0 | 0.42532375 | 0.331 | 0.057 | 0 | 1 | FMN1 |
| 0 | 0.42529651 | 0.462 | 0.239 | 0 | 1 | ATP2A2 |
| 0 | 0.42500321 | 0.288 | 0.031 | 0 | 1 | MLF1 |
| 0 | 0.42494977 | 0.325 | 0.064 | 0 | 1 | SRGAP3 |
| 0 | 0.42480661 | 0.382 | 0.113 | 0 | 1 | DSC2 |
| 0 | 0.42326257 | 0.332 | 0.104 | 0 | 1 | AL450998.2 |
| 0 | 0.42324149 | 0.355 | 0.09 | 0 | 1 | EPB41L2 |
| 0 | 0.422509 | 0.666 | 0.457 | 0 | 1 | CD93 |
| 0 | 0.42247893 | 0.31 | 0.137 | 0 | 1 | KCNQ1OT1 |
| 0 | 0.4221663 | 0.31 | 0.034 | 0 | 1 | IGF2BP3 |
| 0 | 0.42109687 | 0.5 | 0.257 | 0 | 1 | CACYBP |
| 0 | 0.420786 | 0.381 | 0.151 | 0 | 1 | ING1 |
| 0 | 0.42015902 | 0.418 | 0.155 | 0 | 1 | TOR1AIP2 |
| 0 | 0.42004366 | 0.415 | 0.154 | 0 | 1 | TMEM173 |
| 0 | 0.41964634 | 0.436 | 0.17 | 0 | 1 | NECAP2 |
| 0 | 0.4189498 | 0.314 | 0.051 | 0 | 1 | PER3 |
| 0 | 0.41855378 | 0.311 | 0.029 | 0 | 1 | ICA1 |
| 0 | 0.41831461 | 0.376 | 0.114 | 0 | 1 | ARHGAP21 |
| 0 | 0.41773592 | 0.646 | 0.429 | 0 | 1 | KTN1 |
| 0 | 0.41734097 | 0.453 | 0.205 | 0 | 1 | TNRC6B |
| 0 | 0.41690791 | 0.54 | 0.279 | 0 | 1 | GTF2I |
| 0 | 0.41528865 | 0.522 | 0.256 | 0 | 1 | TMED9 |
| 0 | 0.41509781 | 0.288 | 0.024 | 0 | 1 | LRRC4 |
| 0 | 0.41418167 | 0.967 | 0.929 | 0 | 1 | GNAS |
| 0 | 0.41384654 | 0.472 | 0.221 | 0 | 1 | BPTF |
| 0 | 0.41334494 | 0.349 | 0.102 | 0 | 1 | NHLRC3 |
| 0 | 0.41316221 | 0.268 | 0.088 | 0 | 1 | HIST1H2BG |
| 0 | 0.41215222 | 0.418 | 0.2 | 0 | 1 | WDFY2 |
| 0 | 0.41212949 | 0.571 | 0.345 | 0 | 1 | HSPA9 |
| 0 | 0.41167933 | 0.322 | 0.043 | 0 | 1 | NCK2 |
| 0 | 0.41073256 | 0.276 | 0.018 | 0 | 1 | USP9Y |
| 0 | 0.41020068 | 0.365 | 0.089 | 0 | 1 | CHID1 |
| 0 | 0.40918292 | 0.491 | 0.226 | 0 | 1 | DDAH2 |
| 0 | 0.40862422 | 0.538 | 0.28 | 0 | 1 | MDM4 |
| 0 | 0.40691627 | 0.326 | 0.086 | 0 | 1 | TRIP11 |
| 0 | 0.40669974 | 0.604 | 0.362 | 0 | 1 | SERINC1 |
| 0 | 0.40636867 | 0.266 | 0.013 | 0 | 1 | LGI2 |
| 0 | 0.40508561 | 0.429 | 0.164 | 0 | 1 | SRGAP2 |
| 0 | 0.40453599 | 0.29 | 0.046 | 0 | 1 | CCDC170 |
| 0 | 0.40453278 | 0.283 | 0.019 | 0 | 1 | CTTNBP2 |
| 0 | 0.40426934 | 0.414 | 0.154 | 0 | 1 | ZNF638 |
| 0 | 0.40407407 | 0.319 | 0.071 | 0 | 1 | AC020911.2 |
| 0 | 0.402694 | 0.323 | 0.062 | 0 | 1 | APPL2 |

| | | | | | | |
|---|---|---|---|---|---|---|
| 0 | 0.40250671 | 0.388 | 0.119 | 0 | 1 | TBC1D12 |
| 0 | 0.40178355 | 0.393 | 0.147 | 0 | 1 | HDAC9 |
| 0 | 0.400683 | 0.308 | 0.112 | 0 | 1 | ANKH |
| 0 | 0.39993919 | 0.473 | 0.222 | 0 | 1 | ARID1B |
| 0 | 0.39984889 | 0.517 | 0.275 | 0 | 1 | TMEM165 |
| 0 | 0.39943817 | 0.348 | 0.094 | 0 | 1 | PARP1 |
| 0 | 0.39861283 | 0.394 | 0.156 | 0 | 1 | CDKN1B |
| 0 | 0.39846423 | 0.289 | 0.107 | 0 | 1 | PLK2 |
| 0 | 0.39731928 | 0.327 | 0.054 | 0 | 1 | RGL2 |
| 0 | 0.39584909 | 0.306 | 0.048 | 0 | 1 | GIMAP8 |
| 0 | 0.39539582 | 0.424 | 0.16 | 0 | 1 | ASPH |
| 0 | 0.39488512 | 0.278 | 0.017 | 0 | 1 | GRIN2C |
| 0 | 0.39423087 | 0.28 | 0.018 | 0 | 1 | GNG12 |
| 0 | 0.39395125 | 0.41 | 0.157 | 0 | 1 | LBR |
| 0 | 0.39371705 | 0.37 | 0.091 | 0 | 1 | KCNQ1 |
| 0 | 0.39326969 | 0.374 | 0.122 | 0 | 1 | CNTRL |
| 0 | 0.39280782 | 0.424 | 0.177 | 0 | 1 | PHF3 |
| 0 | 0.39276581 | 0.44 | 0.173 | 0 | 1 | ACIN1 |
| 0 | 0.39260483 | 0.329 | 0.082 | 0 | 1 | ROCK2 |
| 0 | 0.39231707 | 0.394 | 0.138 | 0 | 1 | MILR1 |
| 0 | 0.39145245 | 0.408 | 0.162 | 0 | 1 | PCMTD1 |
| 0 | 0.39030587 | 0.519 | 0.303 | 0 | 1 | RPN2 |
| 0 | 0.38968703 | 0.401 | 0.148 | 0 | 1 | GATAD1 |
| 0 | 0.38852807 | 0.367 | 0.1 | 0 | 1 | TNRC18 |
| 0 | 0.38810432 | 0.286 | 0.086 | 0 | 1 | CDKN1C |
| 0 | 0.38751564 | 0.567 | 0.334 | 0 | 1 | PRKAR1A |
| 0 | 0.38745872 | 0.403 | 0.135 | 0 | 1 | FLI1 |
| 0 | 0.3870172 | 0.361 | 0.125 | 0 | 1 | IRF2BPL |
| 0 | 0.38686342 | 0.252 | 0.018 | 0 | 1 | ITM2C |
| 0 | 0.38614705 | 0.425 | 0.162 | 0 | 1 | RBM6 |
| 0 | 0.38586489 | 0.271 | 0.047 | 0 | 1 | PRKCA |
| 0 | 0.38581249 | 0.363 | 0.149 | 0 | 1 | SNHG1 |
| 0 | 0.38529037 | 0.759 | 0.59 | 0 | 1 | CAPZB |
| 0 | 0.38487665 | 0.303 | 0.077 | 0 | 1 | CCNG2 |
| 0 | 0.38475629 | 0.364 | 0.103 | 0 | 1 | CCNL2 |
| 0 | 0.3846515 | 0.466 | 0.223 | 0 | 1 | U2SURP |
| 0 | 0.38416613 | 0.278 | 0.022 | 0 | 1 | CNRIP1 |
| 0 | 0.38393623 | 0.285 | 0.027 | 0 | 1 | SHMT1 |
| 0 | 0.3837929 | 0.322 | 0.074 | 0 | 1 | RAPGEF6 |
| 0 | 0.38273852 | 0.291 | 0.041 | 0 | 1 | EPHX1 |
| 0 | 0.38263291 | 0.276 | 0.024 | 0 | 1 | SKP2 |
| 0 | 0.38242967 | 0.33 | 0.075 | 0 | 1 | SRGAP2C |
| 0 | 0.38221678 | 0.411 | 0.167 | 0 | 1 | CCDC47 |

| | | | | | | |
|---|---|---|---|---|---|---|
| 0 | 0.38169307 | 0.516 | 0.269 | 0 | 1 | GAA |
| 0 | 0.38091679 | 0.491 | 0.237 | 0 | 1 | GCA |
| 0 | 0.3797245 | 0.451 | 0.219 | 0 | 1 | PPP1R12A |
| 0 | 0.37825029 | 0.335 | 0.086 | 0 | 1 | SPIN1 |
| 0 | 0.37731368 | 0.49 | 0.25 | 0 | 1 | KLF3 |
| 0 | 0.37664372 | 0.457 | 0.222 | 0 | 1 | MRPL18 |
| 0 | 0.37658372 | 0.688 | 0.462 | 0 | 1 | PLEKHO1 |
| 0 | 0.37633617 | 0.381 | 0.138 | 0 | 1 | MESD |
| 0 | 0.37620127 | 0.405 | 0.154 | 0 | 1 | EPN1 |
| 0 | 0.3755863 | 0.378 | 0.133 | 0 | 1 | RBM26 |
| 0 | 0.37557679 | 0.437 | 0.178 | 0 | 1 | INPP5D |
| 0 | 0.37470725 | 0.735 | 0.531 | 0 | 1 | CIRBP |
| 0 | 0.37459054 | 0.279 | 0.037 | 0 | 1 | RHOBTB3 |
| 0 | 0.37388047 | 0.33 | 0.076 | 0 | 1 | DISC1 |
| 0 | 0.37369944 | 0.395 | 0.16 | 0 | 1 | CALU |
| 0 | 0.3722212 | 0.409 | 0.169 | 0 | 1 | FGD5-AS1 |
| 0 | 0.37162722 | 0.448 | 0.206 | 0 | 1 | RAB14 |
| 0 | 0.3714407 | 0.431 | 0.221 | 0 | 1 | TRIM38 |
| 0 | 0.37102853 | 0.426 | 0.206 | 0 | 1 | FAM91A1 |
| 0 | 0.37043746 | 0.539 | 0.288 | 0 | 1 | SH2B3 |
| 0 | 0.36947759 | 0.362 | 0.119 | 0 | 1 | UPF2 |
| 0 | 0.36892809 | 0.361 | 0.134 | 0 | 1 | SEL1L |
| 0 | 0.36852988 | 0.3 | 0.082 | 0 | 1 | SEPHS2 |
| 0 | 0.36843409 | 0.661 | 0.449 | 0 | 1 | MEF2A |
| 0 | 0.36835118 | 0.387 | 0.141 | 0 | 1 | VPS36 |
| 0 | 0.36834508 | 0.352 | 0.142 | 0 | 1 | DIPK2A |
| 0 | 0.36820765 | 0.488 | 0.253 | 0 | 1 | SYK |
| 0 | 0.36710931 | 0.257 | 0.014 | 0 | 1 | GIPC2 |
| 0 | 0.36611498 | 0.283 | 0.044 | 0 | 1 | MGAT5 |
| 0 | 0.36562274 | 0.302 | 0.062 | 0 | 1 | C20orf194 |
| 0 | 0.36542346 | 0.326 | 0.081 | 0 | 1 | NFE2L1 |
| 0 | 0.36525312 | 0.425 | 0.186 | 0 | 1 | STX16 |
| 0 | 0.36449718 | 0.414 | 0.173 | 0 | 1 | ADCY7 |
| 0 | 0.36345926 | 0.347 | 0.104 | 0 | 1 | RSBN1L |
| 0 | 0.36341217 | 0.313 | 0.068 | 0 | 1 | HERC2 |
| 0 | 0.3633306 | 0.291 | 0.051 | 0 | 1 | SGPP1 |
| 0 | 0.36314574 | 0.519 | 0.296 | 0 | 1 | SRSF10 |
| 0 | 0.36301768 | 0.296 | 0.058 | 0 | 1 | SDC3 |
| 0 | 0.36176899 | 0.315 | 0.101 | 0 | 1 | LRRC58 |
| 0 | 0.3617168 | 0.447 | 0.228 | 0 | 1 | POLR2A |
| 0 | 0.361147 | 0.572 | 0.349 | 0 | 1 | MAN2B1 |
| 0 | 0.36112575 | 0.466 | 0.245 | 0 | 1 | CAT |
| 0 | 0.36044434 | 0.272 | 0.026 | 0 | 1 | MYO7A |

| | | | | | | |
|---|---|---|---|---|---|---|
| 0 | 0.36018951 | 0.37 | 0.131 | 0 | 1 | MLXIP |
| 0 | 0.35988749 | 0.307 | 0.086 | 0 | 1 | OFD1 |
| 0 | 0.3593274 | 0.387 | 0.172 | 0 | 1 | SLC11A2 |
| 0 | 0.35893347 | 0.327 | 0.084 | 0 | 1 | RNF216 |
| 0 | 0.35889976 | 0.254 | 0.019 | 0 | 1 | KANK2 |
| 0 | 0.35870825 | 0.539 | 0.344 | 0 | 1 | PDIA6 |
| 0 | 0.35860233 | 0.337 | 0.109 | 0 | 1 | TOB2 |
| 0 | 0.35856982 | 0.301 | 0.074 | 0 | 1 | BAG5 |
| 0 | 0.3581781 | 0.391 | 0.181 | 0 | 1 | CREBRF |
| 0 | 0.35741846 | 0.324 | 0.089 | 0 | 1 | SNX13 |
| 0 | 0.35695477 | 0.482 | 0.277 | 0 | 1 | PPIG |
| 0 | 0.3555991 | 0.415 | 0.175 | 0 | 1 | NPTN |
| 0 | 0.35505507 | 0.462 | 0.23 | 0 | 1 | DEGS1 |
| 0 | 0.354515 | 0.378 | 0.134 | 0 | 1 | SIPA1 |
| 0 | 0.3544318 | 0.34 | 0.105 | 0 | 1 | RAP2A |
| 0 | 0.35410612 | 0.317 | 0.078 | 0 | 1 | TSPAN33 |
| 0 | 0.35386631 | 0.449 | 0.214 | 0 | 1 | RCSD1 |
| 0 | 0.35326233 | 0.441 | 0.212 | 0 | 1 | CTNND1 |
| 0 | 0.35322539 | 0.343 | 0.119 | 0 | 1 | HERC4 |
| 0 | 0.35298503 | 0.472 | 0.235 | 0 | 1 | SRSF6 |
| 0 | 0.35286249 | 0.489 | 0.289 | 0 | 1 | MGAT4A |
| 0 | 0.3527453 | 0.39 | 0.161 | 0 | 1 | OGA |
| 0 | 0.3525494 | 0.306 | 0.094 | 0 | 1 | PURA |
| 0 | 0.35235953 | 0.321 | 0.089 | 0 | 1 | SENP7 |
| 0 | 0.35177166 | 0.272 | 0.048 | 0 | 1 | NPHP3 |
| 0 | 0.35165487 | 0.316 | 0.083 | 0 | 1 | YTHDC2 |
| 0 | 0.35103397 | 0.29 | 0.083 | 0 | 1 | SESN3 |
| 0 | 0.35092085 | 0.261 | 0.026 | 0 | 1 | MAN1C1 |
| 0 | 0.34996185 | 0.303 | 0.064 | 0 | 1 | MMP14 |
| 0 | 0.34949545 | 0.495 | 0.269 | 0 | 1 | DRAM2 |
| 0 | 0.34873228 | 0.368 | 0.138 | 0 | 1 | GIMAP1 |
| 0 | 0.34819729 | 0.56 | 0.332 | 0 | 1 | HNRNPF |
| 0 | 0.3480154 | 0.281 | 0.06 | 0 | 1 | EPAS1 |
| 0 | 0.34733408 | 0.4 | 0.175 | 0 | 1 | NEMF |
| 0 | 0.34716787 | 0.346 | 0.108 | 0 | 1 | TRPM2 |
| 0 | 0.34684258 | 0.384 | 0.141 | 0 | 1 | DGKZ |
| 0 | 0.34663946 | 0.378 | 0.162 | 0 | 1 | PI4K2A |
| 0 | 0.34638711 | 0.403 | 0.213 | 0 | 1 | TOB1 |
| 0 | 0.34630039 | 0.517 | 0.294 | 0 | 1 | TMEM14C |
| 0 | 0.34525253 | 0.405 | 0.182 | 0 | 1 | CREBL2 |
| 0 | 0.34481608 | 0.382 | 0.186 | 0 | 1 | PDIA4 |
| 0 | 0.34477396 | 0.486 | 0.279 | 0 | 1 | HNRNPUL1 |
| 0 | 0.34448673 | 0.267 | 0.064 | 0 | 1 | HLX |

| | | | | | | |
|---|---|---|---|---|---|---|
| 0 | 0.34431577 | 0.416 | 0.183 | 0 | 1 | LYL1 |
| 0 | 0.34411396 | 0.39 | 0.176 | 0 | 1 | APPL1 |
| 0 | 0.3439672 | 0.321 | 0.106 | 0 | 1 | ZNF518A |
| 0 | 0.34391564 | 0.272 | 0.047 | 0 | 1 | PON2 |
| 0 | 0.34305817 | 0.364 | 0.144 | 0 | 1 | DNAJB14 |
| 0 | 0.34219201 | 0.266 | 0.055 | 0 | 1 | TP53INP1 |
| 0 | 0.34106088 | 0.382 | 0.154 | 0 | 1 | TMEM35B |
| 0 | 0.34102213 | 0.401 | 0.195 | 0 | 1 | TRPS1 |
| 0 | 0.3406839 | 0.517 | 0.297 | 0 | 1 | OS9 |
| 0 | 0.3403986 | 0.294 | 0.08 | 0 | 1 | APC |
| 0 | 0.34017224 | 0.335 | 0.111 | 0 | 1 | PAXBP1 |
| 0 | 0.33974992 | 0.453 | 0.249 | 0 | 1 | NIPBL |
| 0 | 0.33971077 | 0.437 | 0.216 | 0 | 1 | XPO1 |
| 0 | 0.33967191 | 0.347 | 0.116 | 0 | 1 | MSL1 |
| 0 | 0.33913933 | 0.304 | 0.088 | 0 | 1 | ZNF148 |
| 0 | 0.33913371 | 0.277 | 0.047 | 0 | 1 | ZMYND11 |
| 0 | 0.3387606 | 0.291 | 0.091 | 0 | 1 | SIRT1 |
| 0 | 0.33824336 | 0.466 | 0.24 | 0 | 1 | NUCB1 |
| 0 | 0.3377954 | 0.356 | 0.133 | 0 | 1 | SETD2 |
| 0 | 0.33773165 | 0.516 | 0.284 | 0 | 1 | DENND3 |
| 0 | 0.33754126 | 0.322 | 0.103 | 0 | 1 | VPS53 |
| 0 | 0.33750638 | 0.291 | 0.065 | 0 | 1 | AKAP10 |
| 0 | 0.33748217 | 0.309 | 0.085 | 0 | 1 | CEPT1 |
| 0 | 0.33696993 | 0.271 | 0.067 | 0 | 1 | TNFSF9 |
| 0 | 0.33675979 | 0.355 | 0.14 | 0 | 1 | LMAN1 |
| 0 | 0.33670794 | 0.516 | 0.301 | 0 | 1 | ACAP2 |
| 0 | 0.33547281 | 0.292 | 0.063 | 0 | 1 | NRROS |
| 0 | 0.33541139 | 0.339 | 0.134 | 0 | 1 | FKBP4 |
| 0 | 0.33501433 | 0.396 | 0.185 | 0 | 1 | PBRM1 |
| 0 | 0.33458036 | 0.39 | 0.188 | 0 | 1 | UGP2 |
| 0 | 0.33436153 | 0.499 | 0.291 | 0 | 1 | EIF3D |
| 0 | 0.33405884 | 0.344 | 0.147 | 0 | 1 | RSF1 |
| 0 | 0.33385935 | 0.434 | 0.197 | 0 | 1 | EVL |
| 0 | 0.33381267 | 0.538 | 0.344 | 0 | 1 | MAML2 |
| 0 | 0.33369667 | 0.341 | 0.118 | 0 | 1 | PHKB |
| 0 | 0.33301695 | 0.376 | 0.145 | 0 | 1 | CYBC1 |
| 0 | 0.33245781 | 0.335 | 0.113 | 0 | 1 | TSPAN3 |
| 0 | 0.33240614 | 0.474 | 0.262 | 0 | 1 | SLTM |
| 0 | 0.33212187 | 0.355 | 0.151 | 0 | 1 | KIDINS220 |
| 0 | 0.33182491 | 0.423 | 0.223 | 0 | 1 | TTC3 |
| 0 | 0.3313558 | 0.362 | 0.147 | 0 | 1 | ZNF330 |
| 0 | 0.33105025 | 0.395 | 0.177 | 0 | 1 | NAGA |
| 0 | 0.33099587 | 0.317 | 0.116 | 0 | 1 | ZNF292 |

| | | | | | | |
|---|---|---|---|---|---|---|
| 0 | 0.33099484 | 0.404 | 0.198 | 0 | 1 | SFT2D2 |
| 0 | 0.33058207 | 0.313 | 0.1 | 0 | 1 | RAPH1 |
| 0 | 0.33054793 | 0.3 | 0.076 | 0 | 1 | TMC8 |
| 0 | 0.33031724 | 0.262 | 0.055 | 0 | 1 | SHPRH |
| 0 | 0.32991685 | 0.351 | 0.131 | 0 | 1 | TMC6 |
| 0 | 0.3296446 | 0.272 | 0.082 | 0 | 1 | GPR155 |
| 0 | 0.32916859 | 0.577 | 0.355 | 0 | 1 | DPP7 |
| 0 | 0.32862453 | 0.29 | 0.077 | 0 | 1 | NOL8 |
| 0 | 0.32851139 | 0.304 | 0.088 | 0 | 1 | NSL1 |
| 0 | 0.32844543 | 0.565 | 0.366 | 0 | 1 | TLN1 |
| 0 | 0.32819042 | 0.324 | 0.107 | 0 | 1 | TBC1D2B |
| 0 | 0.32811797 | 0.325 | 0.104 | 0 | 1 | TPCN1 |
| 0 | 0.32752331 | 0.322 | 0.113 | 0 | 1 | NOP56 |
| 0 | 0.32729852 | 0.307 | 0.118 | 0 | 1 | CGAS |
| 0 | 0.32723738 | 0.297 | 0.095 | 0 | 1 | ERV3-1 |
| 0 | 0.32625566 | 0.295 | 0.078 | 0 | 1 | LARP4 |
| 0 | 0.32587685 | 0.309 | 0.135 | 0 | 1 | ODC1 |
| 0 | 0.32573656 | 0.393 | 0.188 | 0 | 1 | PTAFR |
| 0 | 0.32532701 | 0.307 | 0.084 | 0 | 1 | SYNRG |
| 0 | 0.32503242 | 0.335 | 0.134 | 0 | 1 | KDM5A |
| 0 | 0.32493569 | 0.333 | 0.148 | 0 | 1 | ABCE1 |
| 0 | 0.32490927 | 0.616 | 0.429 | 0 | 1 | LAMP2 |
| 0 | 0.32486488 | 0.385 | 0.173 | 0 | 1 | ZNF644 |
| 0 | 0.32465956 | 0.279 | 0.068 | 0 | 1 | CCDC14 |
| 0 | 0.32460241 | 0.254 | 0.041 | 0 | 1 | RALGPS2 |
| 0 | 0.32460193 | 0.562 | 0.338 | 0 | 1 | NAGK |
| 0 | 0.32432636 | 0.502 | 0.275 | 0 | 1 | EHD4 |
| 0 | 0.32400864 | 0.433 | 0.205 | 0 | 1 | TMEM259 |
| 0 | 0.32399815 | 0.344 | 0.13 | 0 | 1 | TNRC6A |
| 0 | 0.32352339 | 0.294 | 0.084 | 0 | 1 | ARAP2 |
| 0 | 0.3235136 | 0.282 | 0.061 | 0 | 1 | RUBCNL |
| 0 | 0.32331941 | 0.285 | 0.104 | 0 | 1 | RHOU |
| 0 | 0.32284256 | 0.567 | 0.366 | 0 | 1 | TRA2A |
| 0 | 0.32228433 | 0.503 | 0.295 | 0 | 1 | CEP170 |
| 0 | 0.32222353 | 0.332 | 0.119 | 0 | 1 | ZNF24 |
| 0 | 0.3219783 | 0.415 | 0.213 | 0 | 1 | SREK1 |
| 0 | 0.3219368 | 0.281 | 0.067 | 0 | 1 | STX2 |
| 0 | 0.32141151 | 0.346 | 0.125 | 0 | 1 | TOR1AIP1 |
| 0 | 0.32132852 | 0.603 | 0.41 | 0 | 1 | RAB5C |
| 0 | 0.32113068 | 0.254 | 0.043 | 0 | 1 | GIMAP6 |
| 0 | 0.32097889 | 0.464 | 0.266 | 0 | 1 | DDX46 |
| 0 | 0.32092614 | 0.352 | 0.144 | 0 | 1 | EFCAB14 |
| 0 | 0.31981424 | 0.347 | 0.126 | 0 | 1 | VAV1 |

| | | | | | | |
|---|---|---|---|---|---|---|
| 0 | 0.31944265 | 0.382 | 0.159 | 0 | 1 | GRAMD1A |
| 0 | 0.31908104 | 0.36 | 0.159 | 0 | 1 | PTPRJ |
| 0 | 0.31896539 | 0.391 | 0.19 | 0 | 1 | TAOK1 |
| 0 | 0.31896513 | 0.268 | 0.096 | 0 | 1 | AC008875.3 |
| 0 | 0.31859801 | 0.295 | 0.082 | 0 | 1 | MERTK |
| 0 | 0.3183462 | 0.527 | 0.331 | 0 | 1 | UBXN4 |
| 0 | 0.3177795 | 0.412 | 0.21 | 0 | 1 | PSMA3-AS1 |
| 0 | 0.31755941 | 0.464 | 0.278 | 0 | 1 | SBDS |
| 0 | 0.31747745 | 0.448 | 0.229 | 0 | 1 | ITGAM |
| 0 | 0.31715068 | 0.318 | 0.11 | 0 | 1 | GTF2A1 |
| 0 | 0.31703517 | 0.52 | 0.307 | 0 | 1 | TBXAS1 |
| 0 | 0.31686005 | 0.304 | 0.091 | 0 | 1 | APMAP |
| 0 | 0.31683368 | 0.407 | 0.208 | 0 | 1 | TMEM30A |
| 0 | 0.31653454 | 0.42 | 0.201 | 0 | 1 | PUM1 |
| 0 | 0.31619332 | 0.311 | 0.093 | 0 | 1 | IKBKB |
| 0 | 0.31598395 | 0.337 | 0.115 | 0 | 1 | OSBPL9 |
| 0 | 0.31559799 | 0.52 | 0.313 | 0 | 1 | OSTF1 |
| 0 | 0.31487401 | 0.306 | 0.094 | 0 | 1 | JKAMP |
| 0 | 0.31484725 | 0.486 | 0.286 | 0 | 1 | WDR26 |
| 0 | 0.31403544 | 0.41 | 0.196 | 0 | 1 | TMED7 |
| 0 | 0.3138784 | 0.371 | 0.165 | 0 | 1 | SRSF1 |
| 0 | 0.31371571 | 0.564 | 0.341 | 0 | 1 | NRIP1 |
| 0 | 0.3132317 | 0.428 | 0.216 | 0 | 1 | SNAP23 |
| 0 | 0.3132208 | 0.442 | 0.241 | 0 | 1 | PRMT2 |
| 0 | 0.31307997 | 0.366 | 0.179 | 0 | 1 | SUGT1 |
| 0 | 0.31275125 | 0.297 | 0.115 | 0 | 1 | MTHFR |
| 0 | 0.31201369 | 0.667 | 0.471 | 0 | 1 | ARL6IP1 |
| 0 | 0.31175965 | 0.374 | 0.157 | 0 | 1 | KIAA0930 |
| 0 | 0.31167728 | 0.39 | 0.196 | 0 | 1 | ZMYM2 |
| 0 | 0.31148264 | 0.381 | 0.182 | 0 | 1 | SPEN |
| 0 | 0.31137865 | 0.279 | 0.084 | 0 | 1 | KCNK6 |
| 0 | 0.31077831 | 0.307 | 0.125 | 0 | 1 | EPM2AIP1 |
| 0 | 0.31063574 | 0.252 | 0.043 | 0 | 1 | ABCD4 |
| 0 | 0.31057465 | 0.285 | 0.071 | 0 | 1 | YTHDF1 |
| 0 | 0.31033487 | 0.308 | 0.107 | 0 | 1 | RABEP1 |
| 0 | 0.30913373 | 0.429 | 0.239 | 0 | 1 | RAD23A |
| 0 | 0.30897011 | 0.288 | 0.098 | 0 | 1 | SMC3 |
| 0 | 0.3087419 | 0.337 | 0.125 | 0 | 1 | PUM2 |
| 0 | 0.3084488 | 0.305 | 0.101 | 0 | 1 | KIAA2026 |
| 0 | 0.30804376 | 0.363 | 0.153 | 0 | 1 | PPP3R1 |
| 0 | 0.3078097 | 0.337 | 0.126 | 0 | 1 | NSD1 |
| 0 | 0.30737935 | 0.329 | 0.116 | 0 | 1 | SLC39A1 |
| 0 | 0.30707968 | 0.295 | 0.103 | 0 | 1 | EVI5 |

| | | | | | | |
|---|---|---|---|---|---|---|
| 0 | 0.30690283 | 0.475 | 0.26 | 0 | 1 | ISCU |
| 0 | 0.30673584 | 0.326 | 0.115 | 0 | 1 | PRRC2B |
| 0 | 0.30651729 | 0.498 | 0.289 | 0 | 1 | SIRPA |
| 0 | 0.30602459 | 0.274 | 0.072 | 0 | 1 | TM7SF3 |
| 0 | 0.30564022 | 0.381 | 0.177 | 0 | 1 | MXD4 |
| 0 | 0.30538789 | 0.767 | 0.573 | 0 | 1 | IFITM2 |
| 0 | 0.30520892 | 0.343 | 0.159 | 0 | 1 | LINC01094 |
| 0 | 0.30515479 | 0.319 | 0.108 | 0 | 1 | GPR137B |
| 0 | 0.30491305 | 0.27 | 0.076 | 0 | 1 | FEM1B |
| 0 | 0.30459035 | 0.273 | 0.075 | 0 | 1 | CYBRD1 |
| 0 | 0.30388965 | 0.422 | 0.221 | 0 | 1 | ASAP1 |
| 0 | 0.30382897 | 0.256 | 0.078 | 0 | 1 | CBX4 |
| 0 | 0.30378751 | 0.343 | 0.156 | 0 | 1 | TMF1 |
| 0 | 0.30357885 | 0.269 | 0.076 | 0 | 1 | C5orf24 |
| 0 | 0.30344575 | 0.285 | 0.084 | 0 | 1 | REV1 |
| 0 | 0.30339743 | 0.271 | 0.071 | 0 | 1 | ANKFY1 |
| 0 | 0.30314254 | 0.347 | 0.146 | 0 | 1 | GARS-DT |
| 0 | 0.30270125 | 0.284 | 0.08 | 0 | 1 | CLN6 |
| 0 | 0.30239792 | 0.431 | 0.226 | 0 | 1 | FNBP4 |
| 0 | 0.3021584 | 0.407 | 0.223 | 0 | 1 | MYCBP2 |
| 0 | 0.30162935 | 0.357 | 0.157 | 0 | 1 | CDK12 |
| 0 | 0.30133719 | 0.358 | 0.177 | 0 | 1 | G3BP1 |
| 0 | 0.30130579 | 0.401 | 0.202 | 0 | 1 | FGFR1OP2 |
| 0 | 0.30121715 | 0.361 | 0.163 | 0 | 1 | DIAPH2 |
| 0 | 0.30121538 | 0.313 | 0.113 | 0 | 1 | GAPVD1 |
| 0 | 0.30107101 | 0.317 | 0.129 | 0 | 1 | CENPC |
| 0 | 0.3009319 | 0.314 | 0.102 | 0 | 1 | GON4L |
| 0 | 0.30065568 | 0.294 | 0.096 | 0 | 1 | PHF14 |
| 0 | 0.3006397 | 0.287 | 0.088 | 0 | 1 | BAZ1B |
| 0 | 0.30063075 | 0.279 | 0.073 | 0 | 1 | SSBP2 |
| 0 | 0.30060189 | 0.412 | 0.236 | 0 | 1 | DDX18 |
| 0 | 0.29970082 | 0.491 | 0.289 | 0 | 1 | ZFYVE16 |
| 0 | 0.29957939 | 0.337 | 0.133 | 0 | 1 | SAFB2 |
| 0 | 0.29840495 | 0.361 | 0.157 | 0 | 1 | NAA20 |
| 0 | 0.2975762 | 0.34 | 0.161 | 0 | 1 | ATXN7L3B |
| 0 | 0.29716911 | 0.405 | 0.221 | 0 | 1 | SPPL2A |
| 0 | 0.2966826 | 0.367 | 0.17 | 0 | 1 | DUSP3 |
| 0 | 0.29538259 | 0.387 | 0.198 | 0 | 1 | CMTM3 |
| 0 | 0.29531748 | 0.255 | 0.057 | 0 | 1 | SLC45A4 |
| 0 | 0.29511303 | 0.332 | 0.155 | 0 | 1 | SLC25A36 |
| 0 | 0.2943456 | 0.409 | 0.204 | 0 | 1 | LMO2 |
| 0 | 0.29408921 | 0.268 | 0.074 | 0 | 1 | SRFBP1 |
| 0 | 0.29391604 | 0.397 | 0.192 | 0 | 1 | CARD8 |

| | | | | | | |
|---|---|---|---|---|---|---|
| 0 | 0.29372883 | 0.295 | 0.104 | 0 | 1 | CD2AP |
| 0 | 0.29356553 | 0.369 | 0.183 | 0 | 1 | LMO4 |
| 0 | 0.29338488 | 0.337 | 0.135 | 0 | 1 | DHRS3 |
| 0 | 0.29304011 | 0.364 | 0.172 | 0 | 1 | PPP3CA |
| 0 | 0.29303374 | 0.284 | 0.091 | 0 | 1 | ATXN2 |
| 0 | 0.29242949 | 0.456 | 0.266 | 0 | 1 | GNAI3 |
| 0 | 0.29210997 | 0.301 | 0.104 | 0 | 1 | TK2 |
| 0 | 0.29208919 | 0.268 | 0.079 | 0 | 1 | ZNF703 |
| 0 | 0.29150967 | 0.328 | 0.131 | 0 | 1 | LONP2 |
| 0 | 0.29128273 | 0.259 | 0.083 | 0 | 1 | ESF1 |
| 0 | 0.29096415 | 0.285 | 0.094 | 0 | 1 | GIGYF2 |
| 0 | 0.29091003 | 0.292 | 0.089 | 0 | 1 | ACP2 |
| 0 | 0.29082817 | 0.279 | 0.087 | 0 | 1 | C1GALT1 |
| 0 | 0.29072706 | 0.315 | 0.12 | 0 | 1 | CDK13 |
| 0 | 0.29018636 | 0.285 | 0.09 | 0 | 1 | AC012368.1 |
| 0 | 0.29005964 | 0.261 | 0.066 | 0 | 1 | EI24 |
| 0 | 0.29003891 | 0.364 | 0.174 | 0 | 1 | TBL1XR1 |
| 0 | 0.28992413 | 0.299 | 0.107 | 0 | 1 | POGZ |
| 0 | 0.28980834 | 0.373 | 0.184 | 0 | 1 | PEPD |
| 0 | 0.28941823 | 0.344 | 0.14 | 0 | 1 | SELPLG |
| 0 | 0.28930641 | 0.33 | 0.137 | 0 | 1 | LGALS8 |
| 0 | 0.28823575 | 0.36 | 0.157 | 0 | 1 | VEGFB |
| 0 | 0.28820097 | 0.325 | 0.119 | 0 | 1 | DENND4B |
| 0 | 0.28796902 | 0.48 | 0.282 | 0 | 1 | NXF1 |
| 0 | 0.28783599 | 0.264 | 0.07 | 0 | 1 | DCAF12 |
| 0 | 0.28765215 | 0.3 | 0.109 | 0 | 1 | DELE1 |
| 0 | 0.28760707 | 0.41 | 0.212 | 0 | 1 | RHBDF2 |
| 0 | 0.28734857 | 0.323 | 0.134 | 0 | 1 | PPIP5K2 |
| 0 | 0.28682212 | 0.264 | 0.087 | 0 | 1 | SMPDL3A |
| 0 | 0.28667462 | 0.313 | 0.121 | 0 | 1 | ST3GAL5 |
| 0 | 0.28627384 | 0.253 | 0.068 | 0 | 1 | SLF2 |
| 0 | 0.28604671 | 0.316 | 0.115 | 0 | 1 | IRF3 |
| 0 | 0.28596931 | 0.306 | 0.13 | 0 | 1 | CCDC82 |
| 0 | 0.28545476 | 0.45 | 0.264 | 0 | 1 | LEPROT |
| 0 | 0.28526775 | 0.341 | 0.165 | 0 | 1 | SREK1IP1 |
| 0 | 0.28486163 | 0.475 | 0.289 | 0 | 1 | SCARB2 |
| 0 | 0.28483665 | 0.327 | 0.128 | 0 | 1 | ANKRD13D |
| 0 | 0.28455956 | 0.322 | 0.131 | 0 | 1 | AHSA1 |
| 0 | 0.28391008 | 0.417 | 0.236 | 0 | 1 | SMARCA5 |
| 0 | 0.2835988 | 0.299 | 0.127 | 0 | 1 | SLC39A10 |
| 0 | 0.28294215 | 0.275 | 0.083 | 0 | 1 | RASA2 |
| 0 | 0.28265125 | 0.484 | 0.279 | 0 | 1 | DDX39B |
| 0 | 0.28263242 | 0.253 | 0.058 | 0 | 1 | ANKRD9 |

| | | | | | | |
|---|---|---|---|---|---|---|
| 0 | 0.28257008 | 0.28 | 0.12 | 0 | 1 | SIAH2 |
| 0 | 0.28256743 | 0.301 | 0.119 | 0 | 1 | PIK3CG |
| 0 | 0.28255798 | 0.27 | 0.089 | 0 | 1 | KMT5A |
| 0 | 0.28253226 | 0.359 | 0.164 | 0 | 1 | LMBRD1 |
| 0 | 0.28201948 | 0.317 | 0.139 | 0 | 1 | NR1D2 |
| 0 | 0.28151691 | 0.372 | 0.175 | 0 | 1 | ELF2 |
| 0 | 0.28140505 | 0.289 | 0.12 | 0 | 1 | SETDB2 |
| 0 | 0.28122921 | 0.352 | 0.166 | 0 | 1 | TMEM70 |
| 0 | 0.28090735 | 0.447 | 0.254 | 0 | 1 | CNDP2 |
| 0 | 0.28090227 | 0.277 | 0.088 | 0 | 1 | PIKFYVE |
| 0 | 0.2807501 | 0.384 | 0.213 | 0 | 1 | RALBP1 |
| 0 | 0.28074207 | 0.321 | 0.135 | 0 | 1 | GNPDA1 |
| 0 | 0.28067496 | 0.277 | 0.092 | 0 | 1 | CCDC91 |
| 0 | 0.28052056 | 0.285 | 0.088 | 0 | 1 | CRYL1 |
| 0 | 0.28049572 | 0.285 | 0.092 | 0 | 1 | SLC12A9 |
| 0 | 0.28044758 | 0.267 | 0.08 | 0 | 1 | ICE2 |
| 0 | 0.27936045 | 0.384 | 0.198 | 0 | 1 | DHX36 |
| 0 | 0.27913366 | 0.283 | 0.133 | 0 | 1 | CXorf21 |
| 0 | 0.27910889 | 0.301 | 0.112 | 0 | 1 | PRDX2 |
| 0 | 0.278744 | 0.29 | 0.117 | 0 | 1 | RIF1 |
| 0 | 0.27838738 | 0.389 | 0.216 | 0 | 1 | PDCD4 |
| 0 | 0.27826426 | 0.268 | 0.091 | 0 | 1 | CWF19L2 |
| 0 | 0.27791132 | 0.367 | 0.187 | 0 | 1 | USP8 |
| 0 | 0.27751034 | 0.388 | 0.199 | 0 | 1 | FKBP15 |
| 0 | 0.27736625 | 0.339 | 0.156 | 0 | 1 | ADI1 |
| 0 | 0.27728496 | 0.41 | 0.222 | 0 | 1 | THRAP3 |
| 0 | 0.27687785 | 0.327 | 0.147 | 0 | 1 | YIPF4 |
| 0 | 0.2767581 | 0.322 | 0.156 | 0 | 1 | ZC3H13 |
| 0 | 0.27659645 | 0.358 | 0.163 | 0 | 1 | CD33 |
| 0 | 0.27597819 | 0.292 | 0.104 | 0 | 1 | NBR1 |
| 0 | 0.27586343 | 0.303 | 0.134 | 0 | 1 | GOLIM4 |
| 0 | 0.27550206 | 0.344 | 0.155 | 0 | 1 | MAP3K1 |
| 0 | 0.2754532 | 0.336 | 0.158 | 0 | 1 | CCPG1 |
| 0 | 0.27506455 | 0.386 | 0.191 | 0 | 1 | TIAL1 |
| 0 | 0.27436153 | 0.277 | 0.093 | 0 | 1 | LUZP1 |
| 0 | 0.27397128 | 0.374 | 0.193 | 0 | 1 | FAR1 |
| 0 | 0.27358177 | 0.266 | 0.078 | 0 | 1 | AMDHD2 |
| 0 | 0.27293346 | 0.398 | 0.202 | 0 | 1 | TPD52L2 |
| 0 | 0.27218499 | 0.297 | 0.11 | 0 | 1 | ANKRD17 |
| 0 | 0.27119924 | 0.313 | 0.153 | 0 | 1 | ITPRIPL2 |
| 0 | 0.27117024 | 0.258 | 0.085 | 0 | 1 | PRKACB |
| 0 | 0.27115089 | 0.364 | 0.179 | 0 | 1 | TPP2 |
| 0 | 0.2708203 | 0.261 | 0.077 | 0 | 1 | ALDH9A1 |

| | | | | | | |
|---|---|---|---|---|---|---|
| 0 | 0.27022561 | 0.42 | 0.243 | 0 | 1 | MGST2 |
| 0 | 0.26933099 | 0.318 | 0.134 | 0 | 1 | ZNRF2 |
| 0 | 0.26908673 | 0.304 | 0.122 | 0 | 1 | CREB3L2 |
| 0 | 0.26874122 | 0.305 | 0.124 | 0 | 1 | TBC1D15 |
| 0 | 0.26864609 | 0.337 | 0.178 | 0 | 1 | LARP7 |
| 0 | 0.26809405 | 0.339 | 0.15 | 0 | 1 | MAP3K11 |
| 0 | 0.26807793 | 0.349 | 0.165 | 0 | 1 | JAK2 |
| 0 | 0.26802451 | 0.32 | 0.144 | 0 | 1 | MIA3 |
| 0 | 0.26802428 | 0.318 | 0.136 | 0 | 1 | DOCK11 |
| 0 | 0.26796982 | 0.338 | 0.161 | 0 | 1 | CDC5L |
| 0 | 0.2669767 | 0.386 | 0.198 | 0 | 1 | EP300 |
| 0 | 0.26681525 | 0.294 | 0.119 | 0 | 1 | SLC38A6 |
| 0 | 0.26629154 | 0.358 | 0.176 | 0 | 1 | TMEM9B |
| 0 | 0.26618215 | 0.309 | 0.134 | 0 | 1 | CORO7 |
| 0 | 0.26587339 | 0.319 | 0.142 | 0 | 1 | ZBTB1 |
| 0 | 0.26566549 | 0.273 | 0.095 | 0 | 1 | CRK |
| 0 | 0.2650651 | 0.379 | 0.205 | 0 | 1 | MRPL14 |
| 0 | 0.2642668 | 0.417 | 0.231 | 0 | 1 | ARID1A |
| 0 | 0.26423034 | 0.292 | 0.107 | 0 | 1 | SLC12A7 |
| 0 | 0.26400063 | 0.261 | 0.093 | 0 | 1 | MCUR1 |
| 0 | 0.26335935 | 0.381 | 0.193 | 0 | 1 | RBM5 |
| 0 | 0.26238352 | 0.376 | 0.197 | 0 | 1 | RABGAP1L |
| 0 | 0.26234035 | 0.25 | 0.081 | 0 | 1 | CREBZF |
| 0 | 0.26226044 | 0.306 | 0.129 | 0 | 1 | MOB1B |
| 0 | 0.26220509 | 0.301 | 0.123 | 0 | 1 | MYSM1 |
| 0 | 0.26206339 | 0.304 | 0.134 | 0 | 1 | TLNRD1 |
| 0 | 0.26150024 | 0.265 | 0.094 | 0 | 1 | CYP20A1 |
| 0 | 0.26135255 | 0.266 | 0.093 | 0 | 1 | ERLEC1 |
| 0 | 0.26101069 | 0.388 | 0.208 | 0 | 1 | VCP |
| 0 | 0.26067645 | 0.288 | 0.117 | 0 | 1 | CLN5 |
| 0 | 0.26042196 | 0.28 | 0.098 | 0 | 1 | CNOT4 |
| 0 | 0.2603533 | 0.277 | 0.101 | 0 | 1 | STAG1 |
| 0 | 0.26020621 | 0.286 | 0.122 | 0 | 1 | TUT4 |
| 0 | 0.25961591 | 0.294 | 0.14 | 0 | 1 | ILF3-DT |
| 0 | 0.25961556 | 0.308 | 0.141 | 0 | 1 | CEBPZ |
| 0 | 0.25939704 | 0.288 | 0.121 | 0 | 1 | MKLN1 |
| 0 | 0.25928218 | 0.3 | 0.119 | 0 | 1 | TTC17 |
| 0 | 0.25860081 | 0.299 | 0.122 | 0 | 1 | RPL7L1 |
| 0 | 0.25823955 | 0.299 | 0.122 | 0 | 1 | SPTLC2 |
| 0 | 0.25785933 | 0.313 | 0.139 | 0 | 1 | GIT2 |
| 0 | 0.2574062 | 0.26 | 0.085 | 0 | 1 | RELL1 |
| 0 | 0.25668037 | 0.285 | 0.109 | 0 | 1 | KATNBL1 |
| 0 | 0.25618905 | 0.25 | 0.087 | 0 | 1 | TCEAL3 |

| | | | | | | |
|---|---|---|---|---|---|---|
| 0 | 0.25605739 | 0.348 | 0.158 | 0 | 1 | ARHGEF1 |
| 0 | 0.25591394 | 0.269 | 0.105 | 0 | 1 | CREB1 |
| 0 | 0.25579653 | 0.319 | 0.138 | 0 | 1 | SUMF2 |
| 0 | 0.25569133 | 0.381 | 0.21 | 0 | 1 | NCOR1 |
| 0 | 0.25561607 | 0.312 | 0.142 | 0 | 1 | CACUL1 |
| 0 | 0.25553333 | 0.369 | 0.181 | 0 | 1 | ARAP1 |
| 0 | 0.25525299 | 0.253 | 0.084 | 0 | 1 | CCDC18-AS1 |
| 0 | 0.25519924 | 0.281 | 0.109 | 0 | 1 | SMC5 |
| 0 | 0.25511976 | 0.386 | 0.2 | 0 | 1 | WBP2 |
| 0 | 0.2547625 | 0.319 | 0.152 | 0 | 1 | EFR3A |
| 0 | 0.25468251 | 0.282 | 0.114 | 0 | 1 | PIK3C2A |
| 0 | 0.25400136 | 0.258 | 0.086 | 0 | 1 | RASA1 |
| 0 | 0.2532655 | 0.283 | 0.133 | 0 | 1 | UPF3A |
| 0 | 0.25318692 | 0.25 | 0.09 | 0 | 1 | RAD50 |
| 0 | 0.25299588 | 0.294 | 0.125 | 0 | 1 | LPIN2 |
| 0 | 0.25274487 | 0.317 | 0.143 | 0 | 1 | RBM33 |
| 0 | 0.25269182 | 0.308 | 0.129 | 0 | 1 | TPST2 |
| 0 | 0.2526611 | 0.334 | 0.161 | 0 | 1 | WAPL |
| 0 | 0.25265028 | 0.315 | 0.142 | 0 | 1 | YTHDF2 |
| 0 | 0.25246354 | 0.315 | 0.159 | 0 | 1 | UGCG |
| 0 | 0.25229182 | 0.298 | 0.122 | 0 | 1 | POLK |
| 0 | 0.25199597 | 0.337 | 0.162 | 0 | 1 | IRF2 |
| 0 | 0.25161144 | 0.328 | 0.153 | 0 | 1 | IRF9 |
| 0 | 0.25145889 | 0.314 | 0.148 | 0 | 1 | DYNLL2 |
| 0 | 0.25060376 | 0.254 | 0.093 | 0 | 1 | PSIP1 |
| 0 | 0.25049457 | 0.291 | 0.12 | 0 | 1 | EXOC1 |
| 0 | 0.25034854 | 0.334 | 0.16 | 0 | 1 | LARP1 |
| 0 | 0.25000799 | 0.376 | 0.201 | 0 | 1 | APH1A |
| 4.26E-307 | 0.26522428 | 0.358 | 0.195 | 1.03E-302 | 1 | DICER1 |
| 1.43E-303 | 0.29476827 | 0.48 | 0.3 | 3.46E-299 | 1 | SSB |
| 2.99E-302 | 0.30245602 | 0.609 | 0.425 | 7.24E-298 | 1 | CAPZA2 |
| 1.15E-300 | 0.28505579 | 0.42 | 0.242 | 2.79E-296 | 1 | HIST1H4C |
| 4.07E-300 | 0.29231792 | 0.438 | 0.26 | 9.86E-296 | 1 | SNHG9 |
| 7.57E-298 | 0.28114969 | 0.423 | 0.252 | 1.83E-293 | 1 | IFNAR1 |
| 1.59E-297 | 0.25489581 | 0.319 | 0.166 | 3.85E-293 | 1 | ARMCX3 |
| 4.37E-297 | 0.34914161 | 0.699 | 0.534 | 1.06E-292 | 1 | LY96 |
| 5.79E-297 | 0.31028645 | 0.375 | 0.211 | 1.40E-292 | 1 | GPATCH2L |
| 5.16E-296 | 0.25415571 | 0.341 | 0.184 | 1.25E-291 | 1 | RDX |
| 5.09E-289 | 0.30509151 | 0.852 | 0.721 | 1.23E-284 | 1 | IFITM3 |
| 1.77E-288 | 0.29650844 | 0.538 | 0.357 | 4.30E-284 | 1 | ROCK1 |
| 1.38E-287 | 0.29077086 | 0.435 | 0.266 | 3.34E-283 | 1 | AIMP1 |
| 2.08E-287 | 0.25575326 | 0.44 | 0.266 | 5.05E-283 | 1 | SYNCRIP |
| 2.02E-285 | 0.71057517 | 0.718 | 0.659 | 4.90E-281 | 1 | NR4A2 |

| | | | | | | |
|---|---|---|---|---|---|---|
| 2.31E-285 | 0.26811034 | 0.43 | 0.26 | 5.59E-281 | 1 | HOOK3 |
| 1.48E-283 | 0.31289454 | 0.537 | 0.361 | 3.59E-279 | 1 | TAF1D |
| 4.01E-282 | 0.29704312 | 0.402 | 0.24 | 9.73E-278 | 1 | EIF5B |
| 1.10E-279 | 0.37965168 | 0.88 | 0.816 | 2.66E-275 | 1 | HCST |
| 9.43E-277 | 0.28386498 | 0.395 | 0.233 | 2.29E-272 | 1 | TUT7 |
| 6.78E-275 | 0.27913545 | 1 | 1 | 1.64E-270 | 1 | MT-CO1 |
| 2.85E-271 | 0.26597298 | 0.456 | 0.286 | 6.90E-267 | 1 | YY1 |
| 1.45E-270 | 0.25934158 | 0.436 | 0.268 | 3.51E-266 | 1 | DDX6 |
| 2.09E-270 | 0.27480841 | 0.601 | 0.416 | 5.08E-266 | 1 | RBM47 |
| 4.37E-270 | 0.25528237 | 0.397 | 0.234 | 1.06E-265 | 1 | MKNK2 |
| 5.70E-268 | 0.27937096 | 0.487 | 0.316 | 1.38E-263 | 1 | PRPF40A |
| 6.73E-267 | 0.25290007 | 0.546 | 0.361 | 1.63E-262 | 1 | PRNP |
| 3.69E-266 | 0.26910115 | 0.462 | 0.292 | 8.95E-262 | 1 | SHTN1 |
| 2.17E-262 | 0.28529573 | 0.301 | 0.161 | 5.26E-258 | 1 | AC104695.4 |
| 1.93E-258 | 0.2590381 | 0.509 | 0.335 | 4.67E-254 | 1 | CD46 |
| 4.01E-257 | 0.29726861 | 0.742 | 0.591 | 9.71E-253 | 1 | PRRC2C |
| 1.39E-256 | 0.4229112 | 0.515 | 0.359 | 3.36E-252 | 1 | WDR74 |
| 2.60E-255 | 0.28794081 | 0.685 | 0.525 | 6.29E-251 | 1 | SARAF |
| 6.88E-253 | 0.51426877 | 0.738 | 0.625 | 1.67E-248 | 1 | SLC38A2 |
| 2.42E-252 | 0.37895542 | 0.558 | 0.394 | 5.86E-248 | 1 | GLS |
| 6.67E-251 | 0.25942003 | 0.484 | 0.317 | 1.62E-246 | 1 | TM9SF3 |
| 3.97E-247 | 0.25418364 | 0.52 | 0.35 | 9.63E-243 | 1 | PTTG1IP |
| 2.29E-245 | 0.30854442 | 0.738 | 0.598 | 5.55E-241 | 1 | CD302 |
| 7.96E-244 | 0.40750906 | 0.715 | 0.556 | 1.93E-239 | 1 | EIF4A3 |
| 4.38E-242 | 0.27657062 | 0.571 | 0.406 | 1.06E-237 | 1 | TMEM59 |
| 1.82E-241 | 0.25952518 | 0.514 | 0.348 | 4.41E-237 | 1 | SP100 |
| 4.14E-240 | 0.2711919 | 0.495 | 0.33 | 1.00E-235 | 1 | TMED5 |
| 7.36E-239 | 0.26276253 | 0.352 | 0.208 | 1.79E-234 | 1 | EIF2AK2 |
| 3.12E-236 | 0.32048549 | 0.742 | 0.614 | 7.56E-232 | 1 | CANX |
| 8.06E-236 | 0.29387507 | 0.656 | 0.502 | 1.95E-231 | 1 | ATP6AP2 |
| 2.06E-235 | 0.30512289 | 0.621 | 0.46 | 5.00E-231 | 1 | CCDC88A |
| 2.18E-234 | 0.25687849 | 0.421 | 0.265 | 5.27E-230 | 1 | MT-ND6 |
| 5.55E-234 | 0.29561484 | 0.631 | 0.475 | 1.35E-229 | 1 | SEC62 |
| 6.54E-225 | 0.27154733 | 0.266 | 0.142 | 1.59E-220 | 1 | GABPB1-AS1 |
| 9.30E-224 | 0.25369431 | 0.277 | 0.148 | 2.25E-219 | 1 | TSC22D1 |
| 4.82E-213 | 0.27378903 | 0.61 | 0.457 | 1.17E-208 | 1 | MTDH |
| 1.45E-205 | 0.27666125 | 0.913 | 0.879 | 3.51E-201 | 1 | CD68 |
| 6.70E-199 | 0.33274012 | 0.868 | 0.819 | 1.62E-194 | 1 | SRSF3 |
| 5.36E-187 | 0.25574557 | 0.719 | 0.58 | 1.30E-182 | 1 | PTGES3 |
| 1.92E-184 | 0.53350343 | 0.877 | 0.891 | 4.64E-180 | 1 | AC020916.1 |
| 9.96E-183 | 0.40888367 | 0.478 | 0.342 | 2.41E-178 | 1 | ARL4C |
| 3.56E-181 | 0.26658732 | 0.479 | 0.335 | 8.64E-177 | 1 | BCAS2 |
| 1.87E-180 | 0.28606919 | 0.703 | 0.577 | 4.52E-176 | 1 | CDV3 |

| | | | | | | |
|---|---|---|---|---|---|---|
| 2.68E-177 | 0.58409075 | 0.683 | 0.575 | 6.50E-173 | 1 | DUSP6 |
| 3.82E-177 | 0.37870441 | 0.452 | 0.318 | 9.26E-173 | 1 | RBBP6 |
| 2.65E-176 | 0.26747036 | 0.407 | 0.27 | 6.41E-172 | 1 | ZC3HAV1 |
| 1.15E-172 | 0.25972166 | 0.418 | 0.289 | 2.79E-168 | 1 | NANS |
| 1.70E-168 | 0.29778215 | 0.382 | 0.253 | 4.13E-164 | 1 | ADM |
| 5.89E-166 | 0.25363301 | 0.432 | 0.3 | 1.43E-161 | 1 | CCNH |
| 3.22E-159 | 0.25920368 | 0.746 | 0.631 | 7.80E-155 | 1 | PDIA3 |
| 1.45E-147 | 0.26334999 | 0.556 | 0.42 | 3.51E-143 | 1 | PARP14 |
| 8.86E-135 | 0.33366657 | 0.503 | 0.386 | 2.15E-130 | 1 | AL021155.5 |
| 1.85E-130 | 0.50159667 | 0.64 | 0.551 | 4.48E-126 | 1 | HBEGF |
| 7.47E-120 | 0.40267611 | 0.594 | 0.509 | 1.81E-115 | 1 | LINC-PINT |
| 6.02E-115 | 0.25690089 | 0.419 | 0.314 | 1.46E-110 | 1 | XAF1 |
| 4.69E-113 | 0.32575852 | 0.316 | 0.216 | 1.14E-108 | 1 | MDM2 |
| 9.58E-102 | 0.38578744 | 0.733 | 0.672 | 2.32E-97 | 1 | SKP1 |
| 5.40E-90 | 0.39032499 | 0.748 | 0.713 | 1.31E-85 | 1 | MAP1LC3B |
| 4.20E-73 | 0.28599372 | 0.998 | 0.999 | 1.02E-68 | 1 | NEAT1 |
| 3.89E-65 | 0.29754435 | 0.825 | 0.79 | 9.44E-61 | 1 | SQSTM1 |
| 2.69E-55 | 0.32712168 | 0.895 | 0.945 | 6.53E-51 | 1 | NFKBIZ |
| 3.73E-51 | 0.3285913 | 0.682 | 0.661 | 9.03E-47 | 1 | FCGR3A |
| 1.41E-31 | 0.35043056 | 0.973 | 0.978 | 3.41E-27 | 1 | MTRNR2L12 |
| 3.15E-30 | 0.27406064 | 0.896 | 0.894 | 7.63E-26 | 1 | GLUL |
| 3.76E-22 | 0.27283647 | 0.787 | 0.777 | 9.12E-18 | 1 | DNAJB6 |
| 0 | 3.08160591 | 0.996 | 0.842 | 0 | 2 | S100A9 |
| 0 | 3.01984661 | 0.993 | 0.569 | 0 | 2 | S100A8 |
| 0 | 2.20830678 | 0.837 | 0.171 | 0 | 2 | S100A12 |
| 0 | 1.96954037 | 0.413 | 0.086 | 0 | 2 | SERPINB2 |
| 0 | 1.77348706 | 0.449 | 0.108 | 0 | 2 | THBS1 |
| 0 | 1.76736948 | 0.968 | 0.427 | 0 | 2 | FCN1 |
| 0 | 1.75301417 | 0.837 | 0.241 | 0 | 2 | VCAN |
| 0 | 1.70675299 | 1 | 0.765 | 0 | 2 | LYZ |
| 0 | 1.3249064 | 0.939 | 0.591 | 0 | 2 | TIMP1 |
| 0 | 1.20442957 | 0.997 | 0.973 | 0 | 2 | CXCL8 |
| 0 | 1.16181095 | 1 | 0.728 | 0 | 2 | S100A4 |
| 0 | 1.16090646 | 0.869 | 0.459 | 0 | 2 | AC020656.1 |
| 0 | 1.15843551 | 0.83 | 0.444 | 0 | 2 | CRIP1 |
| 0 | 1.1240498 | 0.831 | 0.488 | 0 | 2 | CD55 |
| 0 | 1.10925099 | 0.905 | 0.539 | 0 | 2 | EREG |
| 0 | 1.07491012 | 1 | 0.973 | 0 | 2 | CTSS |
| 0 | 1.0610215 | 0.698 | 0.312 | 0 | 2 | APOBEC3A |
| 0 | 1.0606673 | 1 | 0.699 | 0 | 2 | S100A6 |
| 0 | 1.04020115 | 0.769 | 0.359 | 0 | 2 | ITGAX |
| 0 | 0.99185362 | 0.804 | 0.488 | 0 | 2 | TKT |
| 0 | 0.98692677 | 0.84 | 0.593 | 0 | 2 | ATP2B1 |

| | | | | | | |
|---|---|---|---|---|---|---|
| 0 | 0.98250361 | 0.985 | 0.752 | 0 | 2 | IL1B |
| 0 | 0.97413721 | 0.792 | 0.384 | 0 | 2 | CSTA |
| 0 | 0.96568604 | 0.94 | 0.72 | 0 | 2 | ATP2B1-AS1 |
| 0 | 0.95752511 | 0.91 | 0.624 | 0 | 2 | SLC2A3 |
| 0 | 0.94626305 | 0.675 | 0.258 | 0 | 2 | CD300E |
| 0 | 0.93330974 | 0.748 | 0.579 | 0 | 2 | H1FX |
| 0 | 0.93309085 | 0.744 | 0.362 | 0 | 2 | TREM1 |
| 0 | 0.92914801 | 0.889 | 0.579 | 0 | 2 | GPCPD1 |
| 0 | 0.91856811 | 0.784 | 0.445 | 0 | 2 | EHD1 |
| 0 | 0.90463998 | 0.699 | 0.398 | 0 | 2 | ACSL1 |
| 0 | 0.89398085 | 0.62 | 0.289 | 0 | 2 | LUCAT1 |
| 0 | 0.88463399 | 0.787 | 0.423 | 0 | 2 | CD52 |
| 0 | 0.88391215 | 0.748 | 0.426 | 0 | 2 | SLC25A37 |
| 0 | 0.87379859 | 0.607 | 0.23 | 0 | 2 | LGALS2 |
| 0 | 0.87012021 | 0.701 | 0.381 | 0 | 2 | ATP13A3 |
| 0 | 0.85977897 | 0.856 | 0.589 | 0 | 2 | NLRP3 |
| 0 | 0.84977393 | 0.76 | 0.423 | 0 | 2 | SLC11A1 |
| 0 | 0.83651684 | 0.995 | 0.659 | 0 | 2 | S100A10 |
| 0 | 0.83396442 | 0.985 | 0.715 | 0 | 2 | CD44 |
| 0 | 0.83299842 | 0.707 | 0.335 | 0 | 2 | CSF3R |
| 0 | 0.83076612 | 0.804 | 0.494 | 0 | 2 | PPIF |
| 0 | 0.82556216 | 0.489 | 0.092 | 0 | 2 | SELL |
| 0 | 0.82234648 | 0.948 | 0.711 | 0 | 2 | COTL1 |
| 0 | 0.81126304 | 0.673 | 0.324 | 0 | 2 | AQP9 |
| 0 | 0.80725679 | 0.786 | 0.544 | 0 | 2 | SERPINB1 |
| 0 | 0.80683264 | 0.995 | 0.921 | 0 | 2 | GAPDH |
| 0 | 0.78568711 | 1 | 0.739 | 0 | 2 | VIM |
| 0 | 0.75893113 | 0.787 | 0.509 | 0 | 2 | INSIG1 |
| 0 | 0.75599777 | 1 | 0.972 | 0 | 2 | SRGN |
| 0 | 0.74829175 | 0.5 | 0.102 | 0 | 2 | CFP |
| 0 | 0.74222217 | 0.531 | 0.285 | 0 | 2 | BASP1 |
| 0 | 0.73975389 | 0.692 | 0.34 | 0 | 2 | CD48 |
| 0 | 0.73677772 | 0.766 | 0.559 | 0 | 2 | PTGS2 |
| 0 | 0.72885098 | 0.896 | 0.584 | 0 | 2 | SERPINA1 |
| 0 | 0.72869346 | 0.669 | 0.394 | 0 | 2 | JARID2 |
| 0 | 0.72415352 | 0.721 | 0.348 | 0 | 2 | FLNA |
| 0 | 0.71746373 | 0.758 | 0.476 | 0 | 2 | EVI2B |
| 0 | 0.71224828 | 0.744 | 0.537 | 0 | 2 | NCF1 |
| 0 | 0.71131507 | 0.973 | 0.814 | 0 | 2 | LST1 |
| 0 | 0.69291827 | 0.936 | 0.707 | 0 | 2 | TSPO |
| 0 | 0.69150943 | 0.761 | 0.506 | 0 | 2 | SERPINB9 |
| 0 | 0.68650718 | 0.537 | 0.238 | 0 | 2 | CDC42EP3 |
| 0 | 0.68444701 | 0.973 | 0.754 | 0 | 2 | EMP3 |

| | | | | | | |
|---|---|---|---|---|---|---|
| 0 | 0.68017391 | 0.851 | 0.662 | 0 | 2 | LRRFIP1 |
| 0 | 0.67332306 | 0.723 | 0.484 | 0 | 2 | AHNAK |
| 0 | 0.67203393 | 0.564 | 0.193 | 0 | 2 | FGR |
| 0 | 0.67134233 | 0.992 | 0.943 | 0 | 2 | ZEB2 |
| 0 | 0.66774403 | 0.998 | 0.89 | 0 | 2 | SH3BGRL3 |
| 0 | 0.66118331 | 1 | 0.986 | 0 | 2 | RPS9 |
| 0 | 0.65366502 | 0.987 | 0.807 | 0 | 2 | PLAUR |
| 0 | 0.65223676 | 0.549 | 0.372 | 0 | 2 | HIST1H1C |
| 0 | 0.6421488 | 0.537 | 0.26 | 0 | 2 | ZBTB43 |
| 0 | 0.63852991 | 0.65 | 0.381 | 0 | 2 | CARD16 |
| 0 | 0.63767972 | 0.732 | 0.46 | 0 | 2 | TAGLN2 |
| 0 | 0.6356692 | 0.631 | 0.394 | 0 | 2 | TALDO1 |
| 0 | 0.63218742 | 0.949 | 0.781 | 0 | 2 | LCP1 |
| 0 | 0.63014761 | 0.706 | 0.457 | 0 | 2 | IL1RN |
| 0 | 0.6287573 | 0.539 | 0.197 | 0 | 2 | MYO1G |
| 0 | 0.6137158 | 0.962 | 0.852 | 0 | 2 | MAP3K8 |
| 0 | 0.6135056 | 0.868 | 0.559 | 0 | 2 | UPP1 |
| 0 | 0.61149338 | 0.992 | 0.751 | 0 | 2 | BCL2A1 |
| 0 | 0.60627751 | 0.625 | 0.353 | 0 | 2 | STXBP2 |
| 0 | 0.60077322 | 0.58 | 0.326 | 0 | 2 | PNP |
| 0 | 0.58240107 | 0.507 | 0.246 | 0 | 2 | AC015912.3 |
| 0 | 0.5806967 | 1 | 0.987 | 0 | 2 | H3F3A |
| 0 | 0.58052493 | 0.787 | 0.602 | 0 | 2 | CTNNB1 |
| 0 | 0.57382124 | 1 | 0.984 | 0 | 2 | RPS13 |
| 0 | 0.57278662 | 0.537 | 0.256 | 0 | 2 | WARS |
| 0 | 0.57166609 | 0.51 | 0.249 | 0 | 2 | TNIP1 |
| 0 | 0.56305152 | 0.82 | 0.649 | 0 | 2 | GMFG |
| 0 | 0.56209319 | 0.635 | 0.396 | 0 | 2 | SLC43A2 |
| 0 | 0.56202156 | 0.652 | 0.445 | 0 | 2 | FGL2 |
| 0 | 0.56149567 | 1 | 0.977 | 0 | 2 | RPL35A |
| 0 | 0.55726637 | 0.63 | 0.412 | 0 | 2 | PHLDA1 |
| 0 | 0.55062094 | 0.553 | 0.252 | 0 | 2 | SMIM25 |
| 0 | 0.54788717 | 0.989 | 0.864 | 0 | 2 | ANXA1 |
| 0 | 0.54252816 | 0.718 | 0.537 | 0 | 2 | AP1S2 |
| 0 | 0.53453755 | 0.417 | 0.154 | 0 | 2 | MCEMP1 |
| 0 | 0.53351582 | 0.696 | 0.497 | 0 | 2 | GABARAPL1 |
| 0 | 0.53202479 | 0.997 | 0.948 | 0 | 2 | RPL7 |
| 0 | 0.5307372 | 0.998 | 0.969 | 0 | 2 | NACA |
| 0 | 0.52681429 | 1 | 0.996 | 0 | 2 | RPL39 |
| 0 | 0.5260095 | 0.665 | 0.457 | 0 | 2 | PIM3 |
| 0 | 0.52046008 | 0.508 | 0.253 | 0 | 2 | CARD19 |
| 0 | 0.51805077 | 0.627 | 0.417 | 0 | 2 | MAP2K3 |
| 0 | 0.51643782 | 1 | 0.994 | 0 | 2 | RPL34 |

| | | | | | | |
|---|---|---|---|---|---|---|
| 0 | 0.51545481 | 0.878 | 0.679 | 0 | 2 | GSTP1 |
| 0 | 0.51377378 | 0.455 | 0.179 | 0 | 2 | PLP2 |
| 0 | 0.51201952 | 0.921 | 0.802 | 0 | 2 | RPL4 |
| 0 | 0.51047047 | 0.617 | 0.415 | 0 | 2 | AHR |
| 0 | 0.5104498 | 0.369 | 0.098 | 0 | 2 | FCAR |
| 0 | 0.50870016 | 1 | 0.99 | 0 | 2 | RPL26 |
| 0 | 0.5080946 | 0.34 | 0.06 | 0 | 2 | CDA |
| 0 | 0.50612924 | 0.379 | 0.133 | 0 | 2 | CPD |
| 0 | 0.50288189 | 1 | 0.991 | 0 | 2 | RPS14 |
| 0 | 0.50119113 | 0.569 | 0.404 | 0 | 2 | SAMSN1 |
| 0 | 0.49646984 | 0.451 | 0.211 | 0 | 2 | MARCKSL1 |
| 0 | 0.49329685 | 0.999 | 0.982 | 0 | 2 | SOD2 |
| 0 | 0.49253852 | 0.999 | 0.972 | 0 | 2 | RPL6 |
| 0 | 0.49083585 | 0.489 | 0.262 | 0 | 2 | VSIR |
| 0 | 0.49002806 | 0.311 | 0.051 | 0 | 2 | LILRA5 |
| 0 | 0.48936147 | 0.933 | 0.765 | 0 | 2 | CDKN1A |
| 0 | 0.4858591 | 0.624 | 0.357 | 0 | 2 | LSP1 |
| 0 | 0.48246516 | 1 | 0.982 | 0 | 2 | RPL29 |
| 0 | 0.48187542 | 1 | 0.999 | 0 | 2 | RPLP1 |
| 0 | 0.47990575 | 0.985 | 0.929 | 0 | 2 | CEBPB |
| 0 | 0.47971447 | 0.637 | 0.428 | 0 | 2 | RILPL2 |
| 0 | 0.47963836 | 1 | 0.997 | 0 | 2 | RPS24 |
| 0 | 0.4795101 | 1 | 0.986 | 0 | 2 | RPS11 |
| 0 | 0.47905941 | 0.499 | 0.257 | 0 | 2 | ITGA5 |
| 0 | 0.47793592 | 0.461 | 0.224 | 0 | 2 | JAML |
| 0 | 0.47752183 | 0.997 | 0.97 | 0 | 2 | OAZ1 |
| 0 | 0.47464103 | 1 | 0.982 | 0 | 2 | UBA52 |
| 0 | 0.4737228 | 0.513 | 0.296 | 0 | 2 | MAPK6 |
| 0 | 0.4703117 | 1 | 0.995 | 0 | 2 | RPL37 |
| 0 | 0.46960871 | 0.734 | 0.587 | 0 | 2 | TNFRSF1B |
| 0 | 0.46869492 | 0.993 | 0.935 | 0 | 2 | NAMPT |
| 0 | 0.46501414 | 0.998 | 0.966 | 0 | 2 | PFDN5 |
| 0 | 0.46470998 | 0.84 | 0.613 | 0 | 2 | VEGFA |
| 0 | 0.46342582 | 1 | 0.993 | 0 | 2 | RPL32 |
| 0 | 0.46302863 | 1 | 0.997 | 0 | 2 | RPL13 |
| 0 | 0.46276317 | 0.503 | 0.282 | 0 | 2 | CSGALNACT2 |
| 0 | 0.46131906 | 0.394 | 0.126 | 0 | 2 | C19orf38 |
| 0 | 0.45932028 | 0.42 | 0.245 | 0 | 2 | PHLDA2 |
| 0 | 0.45868645 | 0.622 | 0.429 | 0 | 2 | PRELID1 |
| 0 | 0.45529921 | 0.484 | 0.269 | 0 | 2 | PRKAG2 |
| 0 | 0.45461979 | 0.384 | 0.187 | 0 | 2 | RBP7 |
| 0 | 0.45457577 | 0.596 | 0.408 | 0 | 2 | ADGRE5 |
| 0 | 0.45413815 | 0.772 | 0.647 | 0 | 2 | EFHD2 |

| | | | | | | |
|---|---|---|---|---|---|---|
| 0 | 0.45222995 | 0.64 | 0.455 | 0 | 2 | FAM107B |
| 0 | 0.45184703 | 0.466 | 0.209 | 0 | 2 | GLIPR2 |
| 0 | 0.45110432 | 0.385 | 0.169 | 0 | 2 | ANPEP |
| 0 | 0.44838615 | 1 | 0.998 | 0 | 2 | TMSB10 |
| 0 | 0.44589257 | 0.994 | 0.95 | 0 | 2 | RACK1 |
| 0 | 0.44431771 | 0.973 | 0.896 | 0 | 2 | COX4I1 |
| 0 | 0.4435123 | 0.468 | 0.27 | 0 | 2 | METTL9 |
| 0 | 0.44269631 | 0.385 | 0.158 | 0 | 2 | NDRG1 |
| 0 | 0.44077892 | 0.563 | 0.382 | 0 | 2 | PNPLA8 |
| 0 | 0.43884337 | 0.441 | 0.208 | 0 | 2 | RAC2 |
| 0 | 0.43573546 | 0.721 | 0.556 | 0 | 2 | KDM6B |
| 0 | 0.43565202 | 0.986 | 0.928 | 0 | 2 | RPL31 |
| 0 | 0.43334298 | 1 | 0.994 | 0 | 2 | RPL28 |
| 0 | 0.43259255 | 0.428 | 0.203 | 0 | 2 | IRAK3 |
| 0 | 0.43236673 | 0.763 | 0.651 | 0 | 2 | TPM4 |
| 0 | 0.42974762 | 0.769 | 0.66 | 0 | 2 | IRF2BP2 |
| 0 | 0.4293207 | 0.778 | 0.608 | 0 | 2 | FOSL2 |
| 0 | 0.42869293 | 0.999 | 0.97 | 0 | 2 | RPL22 |
| 0 | 0.42790399 | 0.593 | 0.428 | 0 | 2 | FPR1 |
| 0 | 0.42767715 | 0.999 | 0.974 | 0 | 2 | RPL18 |
| 0 | 0.42693897 | 0.594 | 0.43 | 0 | 2 | CORO1A |
| 0 | 0.42683596 | 0.815 | 0.642 | 0 | 2 | MXD1 |
| 0 | 0.4213708 | 0.41 | 0.244 | 0 | 2 | PLBD1 |
| 0 | 0.41500078 | 0.997 | 0.955 | 0 | 2 | RPL27 |
| 0 | 0.41437856 | 0.353 | 0.131 | 0 | 2 | SMAD3 |
| 0 | 0.41079825 | 0.331 | 0.138 | 0 | 2 | PLAC8 |
| 0 | 0.41038111 | 0.545 | 0.348 | 0 | 2 | GBP2 |
| 0 | 0.40977388 | 0.407 | 0.207 | 0 | 2 | PGD |
| 0 | 0.40965336 | 0.304 | 0.075 | 0 | 2 | ICAM3 |
| 0 | 0.40928071 | 0.621 | 0.477 | 0 | 2 | NCF2 |
| 0 | 0.40911821 | 0.472 | 0.277 | 0 | 2 | SLC7A5 |
| 0 | 0.4084834 | 0.48 | 0.306 | 0 | 2 | RIOK3 |
| 0 | 0.408006 | 0.432 | 0.247 | 0 | 2 | IL6R |
| 0 | 0.40790355 | 0.428 | 0.213 | 0 | 2 | LIMD2 |
| 0 | 0.40781687 | 1 | 0.99 | 0 | 2 | RPL30 |
| 0 | 0.40644834 | 0.466 | 0.259 | 0 | 2 | CLEC12A |
| 0 | 0.40585983 | 0.999 | 0.975 | 0 | 2 | RPL24 |
| 0 | 0.40536868 | 1 | 0.994 | 0 | 2 | RPL11 |
| 0 | 0.40535759 | 0.359 | 0.161 | 0 | 2 | LDLR |
| 0 | 0.40456134 | 1 | 0.991 | 0 | 2 | RPL19 |
| 0 | 0.40343878 | 0.501 | 0.307 | 0 | 2 | KYNU |
| 0 | 0.40333252 | 0.425 | 0.246 | 0 | 2 | GK |
| 0 | 0.40325034 | 0.999 | 0.973 | 0 | 2 | RPS7 |

| | | | | | | |
|---|---|---|---|---|---|---|
| 0 | 0.40301931 | 0.568 | 0.404 | 0 | 2 | BACH1 |
| 0 | 0.40261354 | 1 | 0.992 | 0 | 2 | RPLP2 |
| 0 | 0.40232994 | 0.925 | 0.594 | 0 | 2 | G0S2 |
| 0 | 0.40189171 | 0.784 | 0.572 | 0 | 2 | XIST |
| 0 | 0.40136914 | 0.268 | 0.056 | 0 | 2 | RIPOR2 |
| 0 | 0.39820509 | 0.939 | 0.784 | 0 | 2 | TYMP |
| 0 | 0.39809933 | 0.305 | 0.07 | 0 | 2 | SULF2 |
| 0 | 0.39772333 | 0.999 | 0.989 | 0 | 2 | RPS15A |
| 0 | 0.39747582 | 0.901 | 0.777 | 0 | 2 | SERP1 |
| 0 | 0.39619161 | 0.863 | 0.718 | 0 | 2 | PLCG2 |
| 0 | 0.3953656 | 0.679 | 0.543 | 0 | 2 | PTP4A2 |
| 0 | 0.39416032 | 0.719 | 0.575 | 0 | 2 | CD37 |
| 0 | 0.39286161 | 0.925 | 0.86 | 0 | 2 | HIF1A |
| 0 | 0.39279374 | 0.756 | 0.576 | 0 | 2 | GSTO1 |
| 0 | 0.39202689 | 1 | 0.988 | 0 | 2 | RPS3A |
| 0 | 0.39201555 | 0.368 | 0.138 | 0 | 2 | CAPN2 |
| 0 | 0.38929384 | 0.462 | 0.284 | 0 | 2 | ANKRD28 |
| 0 | 0.38608599 | 0.441 | 0.255 | 0 | 2 | AC016831.1 |
| 0 | 0.38535833 | 0.392 | 0.198 | 0 | 2 | EAF1 |
| 0 | 0.38429957 | 1 | 0.983 | 0 | 2 | RPL18A |
| 0 | 0.38229769 | 0.255 | 0.052 | 0 | 2 | ECE1 |
| 0 | 0.37970816 | 0.436 | 0.255 | 0 | 2 | RABGEF1 |
| 0 | 0.3793776 | 0.963 | 0.862 | 0 | 2 | ATP5MG |
| 0 | 0.37790564 | 0.868 | 0.768 | 0 | 2 | SLC25A6 |
| 0 | 0.37725077 | 0.591 | 0.39 | 0 | 2 | STX11 |
| 0 | 0.37226484 | 1 | 0.974 | 0 | 2 | ATP5F1E |
| 0 | 0.37045112 | 0.501 | 0.333 | 0 | 2 | UBE2R2 |
| 0 | 0.36875427 | 0.324 | 0.119 | 0 | 2 | CLEC4E |
| 0 | 0.36742988 | 0.328 | 0.129 | 0 | 2 | CPPED1 |
| 0 | 0.36403451 | 0.427 | 0.238 | 0 | 2 | PLEC |
| 0 | 0.36275261 | 1 | 0.993 | 0 | 2 | RPS8 |
| 0 | 0.35885505 | 0.4 | 0.21 | 0 | 2 | GCH1 |
| 0 | 0.35772527 | 0.815 | 0.733 | 0 | 2 | EEF2 |
| 0 | 0.35323646 | 1 | 0.987 | 0 | 2 | RPS15 |
| 0 | 0.34970382 | 0.438 | 0.253 | 0 | 2 | NFIL3 |
| 0 | 0.34877218 | 0.999 | 0.982 | 0 | 2 | RPL17 |
| 0 | 0.34796646 | 0.86 | 0.725 | 0 | 2 | ELOB |
| 0 | 0.34791901 | 0.995 | 0.981 | 0 | 2 | PABPC1 |
| 0 | 0.34647023 | 0.897 | 0.771 | 0 | 2 | MTRNR2L8 |
| 0 | 0.34613796 | 0.428 | 0.257 | 0 | 2 | LYST |
| 0 | 0.34597017 | 1 | 0.985 | 0 | 2 | FAU |
| 0 | 0.34473429 | 0.332 | 0.155 | 0 | 2 | SEMA6B |
| 0 | 0.33762051 | 0.376 | 0.188 | 0 | 2 | SLCO3A1 |

| | | | | | | |
|---|---|---|---|---|---|---|
| 0 | 0.33583355 | 1 | 0.995 | 0 | 2 | RPS12 |
| 0 | 0.33570338 | 0.319 | 0.141 | 0 | 2 | SPATA13 |
| 0 | 0.3348271 | 1 | 1 | 0 | 2 | FTH1 |
| 0 | 0.33463922 | 1 | 0.994 | 0 | 2 | RPS28 |
| 0 | 0.3340796 | 0.3 | 0.117 | 0 | 2 | PTGER2 |
| 0 | 0.33393347 | 0.306 | 0.139 | 0 | 2 | LRRK2 |
| 0 | 0.3325861 | 0.998 | 0.968 | 0 | 2 | RPL8 |
| 0 | 0.33011991 | 0.285 | 0.097 | 0 | 2 | BST1 |
| 0 | 0.32983545 | 0.992 | 0.942 | 0 | 2 | RPL5 |
| 0 | 0.32186041 | 0.999 | 0.977 | 0 | 2 | RPS21 |
| 0 | 0.32082087 | 0.981 | 0.911 | 0 | 2 | RPS17 |
| 0 | 0.31776791 | 0.299 | 0.113 | 0 | 2 | PRAM1 |
| 0 | 0.3156412 | 1 | 0.994 | 0 | 2 | RPL37A |
| 0 | 0.31507114 | 0.318 | 0.132 | 0 | 2 | DIAPH1 |
| 0 | 0.31223789 | 0.364 | 0.197 | 0 | 2 | HEBP2 |
| 0 | 0.31146486 | 1 | 0.994 | 0 | 2 | RPS18 |
| 0 | 0.31104586 | 0.323 | 0.153 | 0 | 2 | HK2 |
| 0 | 0.30577515 | 0.999 | 0.979 | 0 | 2 | RPL7A |
| 0 | 0.30513914 | 0.338 | 0.164 | 0 | 2 | RARA |
| 0 | 0.29959864 | 0.297 | 0.143 | 0 | 2 | QSOX1 |
| 0 | 0.29453461 | 0.998 | 0.98 | 0 | 2 | RPL36A |
| 0 | 0.29429234 | 0.999 | 0.99 | 0 | 2 | RPS16 |
| 0 | 0.28977094 | 0.999 | 0.988 | 0 | 2 | RPS6 |
| 0 | 0.2861103 | 0.258 | 0.092 | 0 | 2 | MGST1 |
| 0 | 0.28206447 | 1 | 1 | 0 | 2 | EIF1 |
| 0 | 0.28087797 | 1 | 0.997 | 0 | 2 | HLA-B |
| 0 | 0.27983916 | 0.853 | 0.638 | 0 | 2 | ANXA2 |
| 0 | 0.27729755 | 0.28 | 0.128 | 0 | 2 | CD300LB |
| 0 | 0.27704739 | 0.999 | 0.987 | 0 | 2 | RPL15 |
| 0 | 0.2765916 | 1 | 0.989 | 0 | 2 | RPL38 |
| 0 | 0.27532542 | 1 | 0.989 | 0 | 2 | RPL9 |
| 3.68E-307 | 0.3968689 | 0.495 | 0.33 | 8.91E-303 | 2 | SH3BP2 |
| 1.17E-306 | 0.41544244 | 0.642 | 0.475 | 2.84E-302 | 2 | CD93 |
| 5.45E-306 | 0.29155166 | 0.346 | 0.182 | 1.32E-301 | 2 | PRKCB |
| 1.08E-304 | 0.27268999 | 0.999 | 0.981 | 2.61E-300 | 2 | RPL21 |
| 3.58E-302 | 0.32178613 | 0.893 | 0.811 | 8.67E-298 | 2 | CLIC1 |
| 1.64E-301 | 0.27470006 | 0.999 | 0.976 | 3.97E-297 | 2 | RPS4X |
| 6.97E-300 | 0.27055218 | 1 | 0.991 | 1.69E-295 | 2 | RPS2 |
| 1.16E-297 | 0.31717397 | 0.431 | 0.257 | 2.82E-293 | 2 | ZNF385A |
| 6.53E-297 | 0.41180788 | 0.521 | 0.356 | 1.58E-292 | 2 | EZR |
| 2.52E-295 | 0.46904152 | 0.886 | 0.732 | 6.11E-291 | 2 | CXCL2 |
| 6.92E-295 | 0.2512994 | 1 | 0.999 | 1.68E-290 | 2 | RPL41 |
| 8.61E-292 | 0.26861609 | 0.996 | 0.97 | 2.09E-287 | 2 | RPL23A |

| | | | | | | |
|---|---|---|---|---|---|---|
| 3.28E-291 | 0.34511187 | 0.38 | 0.218 | 7.96E-287 | 2 | RYBP |
| 3.39E-291 | 0.36738394 | 0.759 | 0.633 | 8.21E-287 | 2 | ATP5MPL |
| 3.55E-291 | 0.27730425 | 0.996 | 0.976 | 8.60E-287 | 2 | RPL23 |
| 1.08E-290 | 0.36473713 | 0.379 | 0.209 | 2.62E-286 | 2 | RETN |
| 5.32E-290 | 0.34275626 | 0.361 | 0.199 | 1.29E-285 | 2 | GPR132 |
| 1.82E-289 | 0.33803514 | 0.69 | 0.506 | 4.41E-285 | 2 | BHLHE40 |
| 4.27E-289 | 0.37499031 | 0.504 | 0.341 | 1.03E-284 | 2 | BID |
| 2.99E-288 | 0.34979912 | 0.431 | 0.266 | 7.24E-284 | 2 | DENND5A |
| 3.17E-285 | 0.34308677 | 0.769 | 0.656 | 7.69E-281 | 2 | H2AFY |
| 1.86E-284 | 0.29336522 | 0.931 | 0.844 | 4.52E-280 | 2 | EEF1D |
| 1.93E-284 | 0.35416702 | 0.475 | 0.309 | 4.68E-280 | 2 | UBE2J1 |
| 9.16E-284 | 0.36174327 | 0.498 | 0.333 | 2.22E-279 | 2 | BCL3 |
| 8.22E-283 | 0.3643912 | 0.466 | 0.307 | 1.99E-278 | 2 | NUMB |
| 4.95E-278 | 0.48244558 | 0.634 | 0.514 | 1.20E-273 | 2 | EIF1B |
| 8.65E-278 | 0.2832897 | 0.343 | 0.184 | 2.10E-273 | 2 | TES |
| 4.53E-275 | 0.35765789 | 0.398 | 0.241 | 1.10E-270 | 2 | NFAT5 |
| 1.03E-273 | 0.35664629 | 0.718 | 0.572 | 2.49E-269 | 2 | DUSP6 |
| 5.41E-273 | 0.33434854 | 0.396 | 0.239 | 1.31E-268 | 2 | PURB |
| 1.27E-272 | 0.50081445 | 0.687 | 0.595 | 3.07E-268 | 2 | MNDA |
| 6.55E-268 | 0.25339397 | 0.991 | 0.954 | 1.59E-263 | 2 | LAPTM5 |
| 1.10E-267 | 0.2534498 | 0.999 | 0.984 | 2.67E-263 | 2 | RPL36 |
| 2.66E-267 | 0.39299019 | 0.781 | 0.686 | 6.45E-263 | 2 | MBNL1 |
| 6.57E-267 | 0.33871147 | 0.418 | 0.26 | 1.59E-262 | 2 | SH3BP5 |
| 2.39E-265 | 0.32362172 | 0.374 | 0.22 | 5.79E-261 | 2 | NOTCH2 |
| 3.88E-265 | 0.33296281 | 0.407 | 0.252 | 9.41E-261 | 2 | ZBTB7A |
| 3.29E-264 | 0.35979353 | 0.867 | 0.8 | 7.97E-260 | 2 | PTPRC |
| 2.82E-260 | 0.3446875 | 0.739 | 0.599 | 6.83E-256 | 2 | PDE4B |
| 3.87E-260 | 0.3268964 | 0.407 | 0.256 | 9.38E-256 | 2 | AGTRAP |
| 1.77E-259 | 0.28009039 | 0.949 | 0.894 | 4.30E-255 | 2 | CDC42 |
| 1.51E-258 | 0.27474165 | 0.988 | 0.928 | 3.66E-254 | 2 | RPL10A |
| 4.50E-254 | 0.25989967 | 0.983 | 0.938 | 1.09E-249 | 2 | GABARAP |
| 7.56E-254 | 0.26575649 | 0.955 | 0.851 | 1.83E-249 | 2 | ZFAS1 |
| 9.01E-253 | 0.35792394 | 0.698 | 0.593 | 2.18E-248 | 2 | CAST |
| 1.03E-248 | 0.36234116 | 0.675 | 0.561 | 2.49E-244 | 2 | IFNGR2 |
| 1.10E-248 | 0.28351432 | 0.33 | 0.186 | 2.66E-244 | 2 | SNN |
| 4.82E-248 | 0.3195853 | 0.763 | 0.667 | 1.17E-243 | 2 | SUB1 |
| 1.03E-247 | 0.29327349 | 0.392 | 0.24 | 2.50E-243 | 2 | USP3 |
| 5.44E-247 | 0.27237974 | 0.967 | 0.919 | 1.32E-242 | 2 | BRI3 |
| 1.86E-244 | 0.28897929 | 0.901 | 0.834 | 4.50E-240 | 2 | BTF3 |
| 6.82E-244 | 0.27198626 | 0.987 | 0.955 | 1.65E-239 | 2 | MCL1 |
| 6.95E-244 | 0.27949474 | 0.892 | 0.804 | 1.69E-239 | 2 | YWHAZ |
| 5.37E-242 | 0.32404461 | 0.692 | 0.57 | 1.30E-237 | 2 | MSN |
| 4.66E-240 | 0.36863468 | 0.494 | 0.353 | 1.13E-235 | 2 | DMXL2 |

| | | | | | | |
|---|---|---|---|---|---|---|
| 8.79E-237 | 0.30057761 | 0.382 | 0.231 | 2.13E-232 | 2 | B4GALT5 |
| 1.74E-236 | 0.3533049 | 0.459 | 0.315 | 4.22E-232 | 2 | AGFG1 |
| 3.75E-236 | 0.30894119 | 0.639 | 0.481 | 9.09E-232 | 2 | PHACTR1 |
| 1.79E-231 | 0.25950753 | 0.274 | 0.143 | 4.35E-227 | 2 | AGO2 |
| 5.06E-231 | 0.29002915 | 0.916 | 0.855 | 1.23E-226 | 2 | HNRNPC |
| 9.07E-231 | 0.33432255 | 0.532 | 0.386 | 2.20E-226 | 2 | ADAM17 |
| 1.41E-230 | 0.30165859 | 0.835 | 0.73 | 3.41E-226 | 2 | CLEC7A |
| 4.36E-230 | 0.33439907 | 0.449 | 0.296 | 1.06E-225 | 2 | RNF19B |
| 6.12E-230 | 0.37966721 | 0.576 | 0.457 | 1.48E-225 | 2 | SPAG9 |
| 1.79E-229 | 0.31600322 | 0.447 | 0.284 | 4.33E-225 | 2 | BIRC3 |
| 5.05E-228 | 0.32770432 | 0.696 | 0.596 | 1.22E-223 | 2 | TRAPPC5 |
| 7.76E-227 | 0.2512775 | 0.278 | 0.144 | 1.88E-222 | 2 | HCAR3 |
| 2.55E-224 | 0.31548497 | 0.454 | 0.312 | 6.18E-220 | 2 | CASP1 |
| 4.29E-223 | 0.28136073 | 0.56 | 0.386 | 1.04E-218 | 2 | LTA4H |
| 1.97E-222 | 0.31108386 | 0.45 | 0.306 | 4.76E-218 | 2 | LILRB2 |
| 6.36E-220 | 0.35820693 | 0.546 | 0.407 | 1.54E-215 | 2 | MYADM |
| 4.17E-217 | 0.26108544 | 0.369 | 0.224 | 1.01E-212 | 2 | PTK2B |
| 6.34E-217 | 0.25835816 | 0.34 | 0.202 | 1.54E-212 | 2 | TOM1 |
| 1.96E-211 | 0.30828315 | 0.729 | 0.621 | 4.75E-207 | 2 | PSME1 |
| 2.08E-211 | 0.26250546 | 0.934 | 0.853 | 5.05E-207 | 2 | TOMM7 |
| 5.22E-211 | 0.27799758 | 0.795 | 0.685 | 1.27E-206 | 2 | EIF3K |
| 8.91E-211 | 0.32181329 | 0.46 | 0.322 | 2.16E-206 | 2 | UBE2D1 |
| 3.85E-210 | 0.2847383 | 0.708 | 0.587 | 9.33E-206 | 2 | MIDN |
| 4.16E-210 | 0.36877394 | 0.652 | 0.53 | 1.01E-205 | 2 | NFKB1 |
| 1.83E-209 | 0.27097333 | 1 | 0.996 | 4.44E-205 | 2 | JUND |
| 1.95E-209 | 0.25551578 | 0.9 | 0.814 | 4.73E-205 | 2 | TMA7 |
| 2.85E-207 | 0.27318655 | 0.339 | 0.205 | 6.91E-203 | 2 | OXSR1 |
| 2.23E-202 | 0.25441606 | 0.297 | 0.17 | 5.40E-198 | 2 | LRRFIP2 |
| 1.06E-200 | 0.28566501 | 0.412 | 0.276 | 2.57E-196 | 2 | C9orf72 |
| 2.50E-199 | 0.30161054 | 0.452 | 0.294 | 6.06E-195 | 2 | CYTOR |
| 2.30E-198 | 0.25887994 | 0.37 | 0.236 | 5.58E-194 | 2 | RIN3 |
| 2.69E-198 | 0.30901601 | 0.462 | 0.333 | 6.53E-194 | 2 | RAB5IF |
| 1.69E-197 | 0.32942717 | 0.661 | 0.515 | 4.09E-193 | 2 | AC253572.2 |
| 6.90E-195 | 0.27204638 | 0.846 | 0.78 | 1.67E-190 | 2 | PCBP1 |
| 2.11E-194 | 0.30039893 | 0.412 | 0.276 | 5.12E-190 | 2 | NEDD9 |
| 3.00E-194 | 0.30532272 | 0.558 | 0.429 | 7.26E-190 | 2 | PLK3 |
| 3.39E-194 | 0.43007567 | 0.418 | 0.283 | 8.21E-190 | 2 | THBD |
| 6.50E-193 | 0.27446075 | 0.364 | 0.232 | 1.58E-188 | 2 | TLE3 |
| 2.19E-190 | 0.26845501 | 0.766 | 0.658 | 5.32E-186 | 2 | UQCRH |
| 6.49E-189 | 0.25350731 | 0.864 | 0.781 | 1.57E-184 | 2 | APLP2 |
| 1.35E-186 | 0.28247249 | 0.42 | 0.287 | 3.27E-182 | 2 | IRAK1 |
| 1.68E-186 | 0.31449798 | 0.527 | 0.406 | 4.06E-182 | 2 | NUP214 |
| 5.83E-186 | 0.31314245 | 0.886 | 0.832 | 1.41E-181 | 2 | NINJ1 |

| | | | | | | |
|---|---|---|---|---|---|---|
| 7.48E-184 | 0.29298848 | 0.739 | 0.654 | 1.81E-179 | 2 | LCP2 |
| 9.65E-183 | 0.27774107 | 0.31 | 0.181 | 2.34E-178 | 2 | DNAAF1 |
| 9.70E-178 | 0.289761 | 0.589 | 0.486 | 2.35E-173 | 2 | EIF3H |
| 2.60E-177 | 0.32106461 | 0.83 | 0.747 | 6.29E-173 | 2 | C5AR1 |
| 2.68E-175 | 0.31413091 | 0.633 | 0.532 | 6.50E-171 | 2 | ETS2 |
| 3.90E-175 | 0.25382446 | 0.872 | 0.806 | 9.44E-171 | 2 | ITGB2 |
| 6.26E-174 | 0.28152961 | 0.358 | 0.236 | 1.52E-169 | 2 | SERTAD2 |
| 2.82E-172 | 0.25466632 | 0.688 | 0.574 | 6.83E-168 | 2 | FXYD5 |
| 7.52E-172 | 0.26590664 | 0.411 | 0.287 | 1.82E-167 | 2 | ARHGAP26 |
| 2.62E-170 | 0.29336233 | 0.568 | 0.46 | 6.35E-166 | 2 | VASP |
| 7.59E-169 | 0.29820613 | 0.415 | 0.293 | 1.84E-164 | 2 | ARFGAP3 |
| 1.37E-168 | 0.29557758 | 0.624 | 0.531 | 3.32E-164 | 2 | LYN |
| 1.62E-166 | 0.28007922 | 0.47 | 0.351 | 3.94E-162 | 2 | FMNL1 |
| 1.10E-164 | 0.25449715 | 0.458 | 0.334 | 2.66E-160 | 2 | PILRA |
| 2.57E-163 | 0.25818158 | 0.537 | 0.412 | 6.23E-159 | 2 | PKM |
| 8.72E-162 | 0.25135964 | 0.396 | 0.272 | 2.11E-157 | 2 | DPYD |
| 1.09E-161 | 0.27802804 | 0.479 | 0.359 | 2.64E-157 | 2 | RELT |
| 1.13E-161 | 0.27713389 | 0.492 | 0.369 | 2.73E-157 | 2 | HLA-F |
| 4.25E-155 | 0.27851748 | 0.53 | 0.424 | 1.03E-150 | 2 | EIF2S3 |
| 9.31E-145 | 0.32517989 | 0.25 | 0.147 | 2.26E-140 | 2 | HES4 |
| 1.95E-144 | 0.26976818 | 0.454 | 0.348 | 4.73E-140 | 2 | HCK |
| 2.46E-143 | 0.25614497 | 0.553 | 0.434 | 5.95E-139 | 2 | PFKFB3 |
| 5.89E-143 | 0.26391391 | 0.577 | 0.48 | 1.43E-138 | 2 | GRINA |
| 2.70E-141 | 0.25694766 | 0.459 | 0.348 | 6.55E-137 | 2 | CHMP4B |
| 2.99E-139 | 0.2531134 | 0.63 | 0.536 | 7.25E-135 | 2 | PTPRE |
| 2.59E-136 | 0.27866804 | 0.694 | 0.631 | 6.29E-132 | 2 | METRNL |
| 2.19E-132 | 0.26264612 | 0.594 | 0.505 | 5.32E-128 | 2 | ARF6 |
| 2.16E-131 | 0.26780576 | 0.737 | 0.646 | 5.24E-127 | 2 | AC007952.4 |
| 2.70E-128 | 0.27963005 | 0.576 | 0.493 | 6.53E-124 | 2 | BZW1 |
| 1.62E-123 | 0.26262626 | 0.355 | 0.252 | 3.92E-119 | 2 | ACSL4 |
| 4.51E-123 | 0.27703617 | 0.462 | 0.367 | 1.09E-118 | 2 | TLR2 |
| 1.15E-116 | 0.27953061 | 0.571 | 0.503 | 2.79E-112 | 2 | RAP1B |
| 1.22E-115 | 0.26541516 | 0.443 | 0.342 | 2.96E-111 | 2 | FNDC3B |
| 1.34E-115 | 0.25921394 | 0.475 | 0.383 | 3.25E-111 | 2 | MED13L |
| 3.77E-105 | 0.3193798 | 0.461 | 0.38 | 9.13E-101 | 2 | NRIP1 |
| 3.65E-103 | 0.26024228 | 0.677 | 0.607 | 8.86E-99 | 2 | IFITM2 |
| 2.47E-101 | 0.25019916 | 0.449 | 0.36 | 6.00E-97 | 2 | IVNS1ABP |
| 3.40E-100 | 0.28377153 | 0.343 | 0.256 | 8.25E-96 | 2 | RB1CC1 |
| 3.62E-92 | 0.26516556 | 0.623 | 0.561 | 8.77E-88 | 2 | TNFAIP2 |
| 1.14E-73 | 0.27767061 | 0.562 | 0.506 | 2.77E-69 | 2 | TUBA1A |
| 0 | 3.25844801 | 0.978 | 0.599 | 0 | 3 | APOE |
| 0 | 2.83727876 | 0.96 | 0.476 | 0 | 3 | APOC1 |
| 0 | 1.91300193 | 0.414 | 0.114 | 0 | 3 | APOC2 |

| | | | | | | |
|---|---|---|---|---|---|---|
| 0 | 1.50630687 | 0.865 | 0.299 | 0 | 3 | GPNMB |
| 0 | 1.27120596 | 0.924 | 0.61 | 0 | 3 | RGS1 |
| 0 | 1.2583294 | 0.997 | 0.814 | 0 | 3 | HLA-DPB1 |
| 0 | 1.22248713 | 0.945 | 0.541 | 0 | 3 | HLA-DQA1 |
| 0 | 1.1640414 | 1 | 0.931 | 0 | 3 | HLA-DRB1 |
| 0 | 1.15804297 | 0.999 | 0.86 | 0 | 3 | HLA-DPA1 |
| 0 | 1.13078061 | 0.767 | 0.206 | 0 | 3 | TREM2 |
| 0 | 1.07265698 | 1 | 0.963 | 0 | 3 | HLA-DRA |
| 0 | 1.06536014 | 0.982 | 0.696 | 0 | 3 | HLA-DQB1 |
| 0 | 0.9330079 | 0.526 | 0.161 | 0 | 3 | GCHFR |
| 0 | 0.92153579 | 0.385 | 0.106 | 0 | 3 | HAMP |
| 0 | 0.91808624 | 0.976 | 0.92 | 0 | 3 | CTSD |
| 0 | 0.89522553 | 1 | 0.991 | 0 | 3 | CD74 |
| 0 | 0.89083259 | 0.858 | 0.458 | 0 | 3 | CAPG |
| 0 | 0.84601195 | 0.746 | 0.288 | 0 | 3 | CD9 |
| 0 | 0.81354108 | 0.921 | 0.617 | 0 | 3 | LGALS3 |
| 0 | 0.80997529 | 0.955 | 0.705 | 0 | 3 | HLA-DMA |
| 0 | 0.80674824 | 0.966 | 0.723 | 0 | 3 | LGALS1 |
| 0 | 0.79282602 | 0.609 | 0.252 | 0 | 3 | C3 |
| 0 | 0.78480839 | 0.959 | 0.739 | 0 | 3 | CD81 |
| 0 | 0.77275738 | 0.384 | 0.103 | 0 | 3 | NUPR1 |
| 0 | 0.76504494 | 0.993 | 0.829 | 0 | 3 | IFI30 |
| 0 | 0.76418913 | 0.876 | 0.665 | 0 | 3 | DUSP2 |
| 0 | 0.75510042 | 0.569 | 0.197 | 0 | 3 | SDS |
| 0 | 0.74475425 | 0.764 | 0.374 | 0 | 3 | JPT1 |
| 0 | 0.74197279 | 0.963 | 0.849 | 0 | 3 | PLIN2 |
| 0 | 0.73127182 | 0.999 | 0.926 | 0 | 3 | S100A11 |
| 0 | 0.70568766 | 0.916 | 0.696 | 0 | 3 | GPR183 |
| 0 | 0.7043464 | 1 | 1 | 0 | 3 | FTL |
| 0 | 0.69701729 | 0.507 | 0.233 | 0 | 3 | CD69 |
| 0 | 0.68086226 | 0.652 | 0.272 | 0 | 3 | IL18 |
| 0 | 0.67913489 | 0.858 | 0.565 | 0 | 3 | CXCR4 |
| 0 | 0.67277393 | 0.865 | 0.651 | 0 | 3 | PRDX1 |
| 0 | 0.67071597 | 0.937 | 0.834 | 0 | 3 | SGK1 |
| 0 | 0.66256814 | 0.409 | 0.185 | 0 | 3 | HLA-DQA2 |
| 0 | 0.66189132 | 0.493 | 0.163 | 0 | 3 | ACP5 |
| 0 | 0.6518279 | 0.995 | 0.981 | 0 | 3 | PSAP |
| 0 | 0.57615594 | 0.831 | 0.517 | 0 | 3 | CTSH |
| 0 | 0.5759602 | 0.957 | 0.8 | 0 | 3 | NME2 |
| 0 | 0.56094363 | 0.998 | 0.971 | 0 | 3 | NPC2 |
| 0 | 0.56085931 | 0.914 | 0.705 | 0 | 3 | ATP6V1F |
| 0 | 0.55525913 | 0.647 | 0.372 | 0 | 3 | LIPA |
| 0 | 0.55306999 | 0.543 | 0.24 | 0 | 3 | DNASE2 |

| | | | | | | |
|---|---|---|---|---|---|---|
| 0 | 0.55079484 | 0.821 | 0.603 | 0 | 3 | DBI |
| 0 | 0.54627871 | 1 | 0.99 | 0 | 3 | RPS19 |
| 0 | 0.51487547 | 1 | 1 | 0 | 3 | B2M |
| 0 | 0.51368448 | 0.944 | 0.811 | 0 | 3 | RPSA |
| 0 | 0.50331651 | 0.788 | 0.524 | 0 | 3 | HLA-DMB |
| 0 | 0.49897394 | 0.615 | 0.311 | 0 | 3 | ALDH2 |
| 0 | 0.49444709 | 0.331 | 0.058 | 0 | 3 | CADM1 |
| 0 | 0.48420727 | 0.998 | 0.979 | 0 | 3 | RPS4X |
| 0 | 0.48322695 | 0.679 | 0.42 | 0 | 3 | MGST3 |
| 0 | 0.48258211 | 0.999 | 0.995 | 0 | 3 | RPS28 |
| 0 | 0.47904356 | 0.589 | 0.322 | 0 | 3 | ARL5A |
| 0 | 0.47712817 | 1 | 0.998 | 0 | 3 | RPL10 |
| 0 | 0.47194905 | 0.956 | 0.843 | 0 | 3 | GPX4 |
| 0 | 0.46288616 | 0.999 | 0.992 | 0 | 3 | RPS2 |
| 0 | 0.45404277 | 0.724 | 0.463 | 0 | 3 | CHCHD10 |
| 0 | 0.44725518 | 0.748 | 0.486 | 0 | 3 | SSR3 |
| 0 | 0.44490934 | 0.965 | 0.891 | 0 | 3 | RPS5 |
| 0 | 0.44474739 | 0.996 | 0.972 | 0 | 3 | HLA-C |
| 0 | 0.43639751 | 0.994 | 0.967 | 0 | 3 | CD63 |
| 0 | 0.43147855 | 0.999 | 0.989 | 0 | 3 | HLA-A |
| 0 | 0.41761031 | 0.999 | 0.994 | 0 | 3 | RPS18 |
| 0 | 0.412545 | 0.999 | 0.995 | 0 | 3 | TYROBP |
| 0 | 0.41033677 | 0.979 | 0.894 | 0 | 3 | COX7C |
| 0 | 0.40911933 | 0.999 | 0.996 | 0 | 3 | CST3 |
| 0 | 0.40541491 | 0.999 | 0.989 | 0 | 3 | RPL12 |
| 0 | 0.39553713 | 1 | 0.999 | 0 | 3 | RPL41 |
| 0 | 0.39459701 | 0.995 | 0.972 | 0 | 3 | RPS3 |
| 0 | 0.3905783 | 1 | 0.995 | 0 | 3 | RPS27 |
| 0 | 0.37991044 | 1 | 0.999 | 0 | 3 | RPLP1 |
| 0 | 0.37949965 | 0.361 | 0.135 | 0 | 3 | HGF |
| 0 | 0.37544422 | 1 | 1 | 0 | 3 | TMSB4X |
| 0 | 0.34814414 | 0.414 | 0.18 | 0 | 3 | FPR3 |
| 0 | 0.3055943 | 0.296 | 0.095 | 0 | 3 | HLA-DOA |
| 0 | 0.29636111 | 0.277 | 0.082 | 0 | 3 | GAL3ST4 |
| 7.15E-304 | 0.38173245 | 1 | 0.995 | 1.73E-299 | 3 | RPL28 |
| 7.30E-302 | 0.46233262 | 0.733 | 0.481 | 1.77E-297 | 3 | YWHAH |
| 1.52E-301 | 0.49585806 | 0.737 | 0.502 | 3.69E-297 | 3 | ATOX1 |
| 1.03E-300 | 0.45725032 | 0.767 | 0.521 | 2.49E-296 | 3 | PLXDC2 |
| 2.17E-294 | 0.4600604 | 0.666 | 0.394 | 5.27E-290 | 3 | LTA4H |
| 6.54E-292 | 0.43184617 | 0.885 | 0.704 | 1.59E-287 | 3 | NDUFA4 |
| 2.01E-290 | 0.36323703 | 0.996 | 0.978 | 4.87E-286 | 3 | RPL3 |
| 5.11E-290 | 0.55459772 | 0.891 | 0.759 | 1.24E-285 | 3 | CSTB |
| 6.11E-288 | 0.41335234 | 0.91 | 0.776 | 1.48E-283 | 3 | VAMP8 |

| | | | | | | |
|---|---|---|---|---|---|---|
| 4.82E-285 | 0.35176155 | 0.999 | 0.996 | 1.17E-280 | 3 | RPS12 |
| 3.76E-284 | 0.42026455 | 0.746 | 0.475 | 9.13E-280 | 3 | MSR1 |
| 1.58E-283 | 0.42836649 | 0.851 | 0.644 | 3.83E-279 | 3 | UQCR10 |
| 1.09E-281 | 0.32100611 | 0.999 | 0.993 | 2.63E-277 | 3 | RPL27A |
| 2.98E-280 | 0.36135963 | 1 | 0.995 | 7.23E-276 | 3 | RPS29 |
| 1.23E-279 | 0.6097585 | 0.762 | 0.547 | 2.98E-275 | 3 | FABP5 |
| 8.19E-278 | 0.36456074 | 0.979 | 0.92 | 1.99E-273 | 3 | RPLP0 |
| 1.69E-277 | 0.33455214 | 0.998 | 0.986 | 4.11E-273 | 3 | RPL36 |
| 1.38E-271 | 0.35408211 | 0.998 | 0.989 | 3.35E-267 | 3 | RPS27A |
| 1.46E-267 | 0.33449948 | 1 | 0.994 | 3.54E-263 | 3 | RPL11 |
| 6.15E-265 | 0.40724615 | 0.903 | 0.774 | 1.49E-260 | 3 | NPM1 |
| 1.45E-263 | 0.32657149 | 0.997 | 0.981 | 3.51E-259 | 3 | RPL35 |
| 3.92E-261 | 0.39898814 | 0.855 | 0.657 | 9.50E-257 | 3 | SSR4 |
| 8.62E-251 | 0.27399839 | 0.308 | 0.128 | 2.09E-246 | 3 | GAPT |
| 1.48E-247 | 0.35556114 | 0.946 | 0.844 | 3.58E-243 | 3 | ATP5MC2 |
| 1.58E-246 | 0.3936238 | 0.735 | 0.509 | 3.83E-242 | 3 | ATP5MC3 |
| 1.03E-245 | 0.32113581 | 0.998 | 0.989 | 2.51E-241 | 3 | RPS15 |
| 2.47E-244 | 0.3454173 | 1 | 1 | 5.98E-240 | 3 | FTH1 |
| 3.17E-242 | 0.42072541 | 0.797 | 0.589 | 7.67E-238 | 3 | NDUFB2 |
| 7.99E-241 | 0.36776736 | 0.861 | 0.628 | 1.94E-236 | 3 | C1orf162 |
| 8.97E-236 | 0.36678434 | 0.996 | 0.974 | 2.17E-231 | 3 | SERF2 |
| 2.38E-235 | 0.29487409 | 1 | 0.996 | 5.77E-231 | 3 | RPS23 |
| 2.61E-234 | 0.28333471 | 0.315 | 0.139 | 6.32E-230 | 3 | ALDH1A1 |
| 2.37E-230 | 0.46218696 | 0.718 | 0.496 | 5.74E-226 | 3 | ZNF331 |
| 3.20E-227 | 0.28620942 | 1 | 0.997 | 7.75E-223 | 3 | HLA-B |
| 4.16E-227 | 0.27024045 | 0.29 | 0.125 | 1.01E-222 | 3 | SERPINF1 |
| 1.18E-225 | 0.34799953 | 0.952 | 0.881 | 2.87E-221 | 3 | EEF1B2 |
| 3.80E-224 | 0.29970768 | 0.999 | 0.987 | 9.21E-220 | 3 | FAU |
| 1.23E-223 | 0.39474927 | 0.86 | 0.703 | 2.99E-219 | 3 | RGS10 |
| 2.03E-222 | 0.38906699 | 0.766 | 0.546 | 4.91E-218 | 3 | RPS27L |
| 6.60E-222 | 0.31356588 | 1 | 0.993 | 1.60E-217 | 3 | RPS8 |
| 1.31E-219 | 0.30030431 | 0.998 | 0.989 | 3.19E-215 | 3 | RPL15 |
| 3.43E-218 | 0.3256103 | 0.603 | 0.363 | 8.32E-214 | 3 | PPDPF |
| 8.11E-218 | 0.39285161 | 0.744 | 0.55 | 1.97E-213 | 3 | SLC25A5 |
| 3.42E-216 | 0.34493182 | 0.926 | 0.735 | 8.28E-212 | 3 | TSPO |
| 2.91E-215 | 0.31064884 | 0.999 | 0.977 | 7.06E-211 | 3 | RPL18 |
| 6.16E-215 | 0.35699778 | 0.836 | 0.637 | 1.49E-210 | 3 | ATP5MD |
| 1.08E-214 | 0.36867919 | 0.641 | 0.421 | 2.63E-210 | 3 | TUBB |
| 1.78E-214 | 0.29921316 | 0.998 | 0.99 | 4.31E-210 | 3 | RPS15A |
| 3.79E-213 | 0.37613955 | 0.996 | 0.769 | 9.20E-209 | 3 | VIM |
| 3.95E-211 | 0.45141943 | 0.841 | 0.709 | 9.57E-207 | 3 | CFD |
| 4.95E-211 | 0.31609306 | 0.954 | 0.871 | 1.20E-206 | 3 | SRP14 |
| 2.05E-208 | 0.31524418 | 0.599 | 0.374 | 4.98E-204 | 3 | POLD4 |

| | | | | | | |
|---|---|---|---|---|---|---|
| 3.22E-208 | 0.29855575 | 0.996 | 0.982 | 7.82E-204 | 3 | RPL7A |
| 5.83E-207 | 0.44648386 | 0.442 | 0.245 | 1.41E-202 | 3 | PLAU |
| 6.02E-207 | 0.33239229 | 0.789 | 0.575 | 1.46E-202 | 3 | FXYD5 |
| 7.44E-207 | 0.36184492 | 0.824 | 0.652 | 1.80E-202 | 3 | SUMO2 |
| 4.90E-204 | 0.32327987 | 0.999 | 0.977 | 1.19E-199 | 3 | ATP5F1E |
| 1.57E-203 | 0.29163978 | 0.997 | 0.98 | 3.81E-199 | 3 | RPS21 |
| 2.81E-202 | 0.31224678 | 0.98 | 0.935 | 6.82E-198 | 3 | RPL10A |
| 1.32E-201 | 0.3391888 | 0.7 | 0.486 | 3.20E-197 | 3 | HMGN1 |
| 2.46E-200 | 0.32505929 | 0.359 | 0.184 | 5.97E-196 | 3 | CPM |
| 3.05E-199 | 0.27451608 | 0.997 | 0.982 | 7.40E-195 | 3 | RPL14 |
| 5.75E-197 | 0.2791857 | 0.501 | 0.282 | 1.39E-192 | 3 | SORL1 |
| 6.61E-197 | 0.3310597 | 0.705 | 0.49 | 1.60E-192 | 3 | ATP5PF |
| 1.38E-195 | 0.63674952 | 0.553 | 0.365 | 3.35E-191 | 3 | HLA-DRB5 |
| 5.49E-193 | 0.29224381 | 0.999 | 0.991 | 1.33E-188 | 3 | RPL30 |
| 1.55E-192 | 0.32954761 | 0.923 | 0.824 | 3.76E-188 | 3 | ASAH1 |
| 2.72E-192 | 0.33845556 | 0.812 | 0.642 | 6.58E-188 | 3 | ATP6V0E1 |
| 1.50E-190 | 0.29005428 | 0.348 | 0.177 | 3.63E-186 | 3 | SLC1A3 |
| 1.13E-189 | 0.3022148 | 0.427 | 0.24 | 2.75E-185 | 3 | GM2A |
| 2.43E-189 | 0.28032924 | 0.998 | 0.976 | 5.90E-185 | 3 | RPS7 |
| 1.65E-188 | 0.34407484 | 0.841 | 0.674 | 3.99E-184 | 3 | COX6C |
| 4.06E-188 | 0.32216449 | 0.555 | 0.355 | 9.85E-184 | 3 | LRPAP1 |
| 5.24E-188 | 0.33558668 | 0.84 | 0.663 | 1.27E-183 | 3 | NDUFA1 |
| 7.21E-185 | 0.35753706 | 0.792 | 0.618 | 1.75E-180 | 3 | DYNLL1 |
| 4.03E-184 | 0.31705566 | 0.678 | 0.47 | 9.76E-180 | 3 | ATP5IF1 |
| 6.36E-180 | 0.46977688 | 0.882 | 0.795 | 1.54E-175 | 3 | TXNIP |
| 1.36E-179 | 0.25165062 | 0.999 | 0.995 | 3.29E-175 | 3 | RPL37A |
| 4.15E-179 | 0.3078112 | 0.613 | 0.408 | 1.01E-174 | 3 | LY86 |
| 4.37E-177 | 0.45471658 | 0.724 | 0.54 | 1.06E-172 | 3 | LMNA |
| 1.07E-176 | 0.32055021 | 0.792 | 0.606 | 2.60E-172 | 3 | TMEM258 |
| 1.25E-175 | 0.2772745 | 0.999 | 0.993 | 3.03E-171 | 3 | RPS14 |
| 1.31E-174 | 0.25009142 | 0.999 | 0.992 | 3.17E-170 | 3 | RPL19 |
| 5.63E-172 | 0.26624433 | 0.997 | 0.99 | 1.37E-167 | 3 | RPS3A |
| 9.28E-171 | 0.29084454 | 0.918 | 0.786 | 2.25E-166 | 3 | COX6B1 |
| 1.26E-169 | 0.30461044 | 0.548 | 0.341 | 3.06E-165 | 3 | SMAD7 |
| 1.57E-167 | 0.30152221 | 0.857 | 0.705 | 3.80E-163 | 3 | OST4 |
| 2.22E-167 | 0.29952748 | 0.928 | 0.829 | 5.38E-163 | 3 | RPL36AL |
| 6.96E-167 | 0.33169651 | 0.982 | 0.956 | 1.69E-162 | 3 | ID2 |
| 8.54E-167 | 0.27500015 | 0.65 | 0.442 | 2.07E-162 | 3 | MZT2B |
| 3.68E-166 | 0.45292318 | 0.859 | 0.74 | 8.92E-162 | 3 | ATF3 |
| 1.37E-165 | 0.2744881 | 0.498 | 0.309 | 3.33E-161 | 3 | OSTC |
| 3.59E-164 | 0.30636666 | 0.87 | 0.703 | 8.69E-160 | 3 | GSTP1 |
| 5.43E-164 | 0.30131642 | 0.713 | 0.52 | 1.32E-159 | 3 | KRTCAP2 |
| 8.49E-164 | 0.27219333 | 0.483 | 0.297 | 2.06E-159 | 3 | ALKBH7 |

| | | | | | | |
|---|---|---|---|---|---|---|
| 2.45E-163 | 0.30090592 | 0.861 | 0.705 | 5.94E-159 | 3 | COX5B |
| 2.63E-158 | 0.29703779 | 0.447 | 0.26 | 6.37E-154 | 3 | MIR155HG |
| 2.74E-158 | 0.29599649 | 0.591 | 0.4 | 6.64E-154 | 3 | SYNGR2 |
| 1.19E-157 | 0.29250278 | 0.531 | 0.349 | 2.88E-153 | 3 | APBB1IP |
| 1.41E-157 | 0.34768178 | 0.796 | 0.644 | 3.42E-153 | 3 | PPIB |
| 7.31E-157 | 0.38202739 | 0.919 | 0.868 | 1.77E-152 | 3 | GRN |
| 2.14E-155 | 0.3935968 | 0.703 | 0.503 | 5.19E-151 | 3 | TXN |
| 2.79E-155 | 0.27637497 | 0.841 | 0.662 | 6.75E-151 | 3 | UQCRH |
| 9.18E-154 | 0.38692073 | 0.99 | 0.978 | 2.23E-149 | 3 | IER3 |
| 2.03E-153 | 0.3052309 | 0.866 | 0.731 | 4.92E-149 | 3 | UQCR11 |
| 3.15E-152 | 0.26537422 | 0.631 | 0.432 | 7.64E-148 | 3 | NDUFB4 |
| 3.33E-152 | 0.28650749 | 0.847 | 0.692 | 8.08E-148 | 3 | EIF3K |
| 1.28E-151 | 0.2827854 | 0.578 | 0.389 | 3.11E-147 | 3 | DAD1 |
| 1.60E-151 | 0.26292324 | 0.83 | 0.614 | 3.88E-147 | 3 | C15orf48 |
| 9.28E-150 | 0.25302745 | 1 | 0.999 | 2.25E-145 | 3 | TMSB10 |
| 9.51E-150 | 0.26535328 | 0.946 | 0.87 | 2.31E-145 | 3 | UQCRB |
| 2.11E-149 | 0.26060814 | 0.566 | 0.373 | 5.12E-145 | 3 | CHURC1 |
| 5.32E-149 | 0.29451465 | 0.803 | 0.637 | 1.29E-144 | 3 | POMP |
| 1.00E-148 | 0.25299636 | 0.996 | 0.984 | 2.43E-144 | 3 | RPL17 |
| 5.34E-148 | 0.25811733 | 0.985 | 0.949 | 1.29E-143 | 3 | RPL5 |
| 1.52E-144 | 0.2746166 | 0.8 | 0.632 | 3.68E-140 | 3 | SNHG29 |
| 1.60E-144 | 0.25754002 | 0.525 | 0.342 | 3.88E-140 | 3 | RABAC1 |
| 1.07E-143 | 0.28641862 | 0.661 | 0.473 | 2.60E-139 | 3 | NDUFA3 |
| 1.29E-143 | 0.30013277 | 0.813 | 0.659 | 3.14E-139 | 3 | NAP1L1 |
| 2.39E-139 | 0.273789 | 0.954 | 0.863 | 5.79E-135 | 3 | ZFAS1 |
| 7.79E-139 | 0.27496344 | 0.558 | 0.373 | 1.89E-134 | 3 | PHPT1 |
| 1.80E-138 | 0.2911398 | 0.996 | 0.992 | 4.36E-134 | 3 | CCL3 |
| 7.61E-138 | 0.27408724 | 0.643 | 0.458 | 1.84E-133 | 3 | RBX1 |
| 9.76E-138 | 0.27242862 | 0.785 | 0.621 | 2.37E-133 | 3 | SEC11A |
| 4.63E-136 | 0.28105661 | 0.742 | 0.548 | 1.12E-131 | 3 | CLEC2B |
| 4.71E-136 | 0.26334782 | 0.853 | 0.711 | 1.14E-131 | 3 | LAMTOR4 |
| 7.79E-136 | 0.27812162 | 0.785 | 0.616 | 1.89E-131 | 3 | SEC61B |
| 3.10E-135 | 0.28136657 | 0.814 | 0.643 | 7.52E-131 | 3 | COX7B |
| 2.57E-132 | 0.27797619 | 0.871 | 0.742 | 6.22E-128 | 3 | GNG5 |
| 2.72E-132 | 0.25183918 | 0.362 | 0.208 | 6.59E-128 | 3 | FBP1 |
| 2.21E-131 | 0.25031093 | 0.968 | 0.925 | 5.35E-127 | 3 | BRI3 |
| 3.83E-131 | 0.27399343 | 0.875 | 0.739 | 9.28E-127 | 3 | ELOB |
| 4.28E-131 | 0.2995693 | 0.719 | 0.524 | 1.04E-126 | 3 | BHLHE40 |
| 6.94E-131 | 0.26748328 | 0.633 | 0.447 | 1.68E-126 | 3 | ADA2 |
| 6.17E-130 | 0.27915041 | 0.822 | 0.673 | 1.50E-125 | 3 | NDUFA13 |
| 2.77E-127 | 0.25515376 | 0.782 | 0.613 | 6.72E-123 | 3 | COX6A1 |
| 8.18E-126 | 0.27115705 | 0.865 | 0.724 | 1.98E-121 | 3 | ATP5ME |
| 1.71E-125 | 0.28077038 | 0.588 | 0.393 | 4.15E-121 | 3 | SNX10 |

| | | | | | | |
|---|---|---|---|---|---|---|
| 4.04E-123 | 0.25016061 | 0.467 | 0.308 | 9.81E-119 | 3 | MYDGF |
| 4.47E-123 | 0.29571305 | 0.625 | 0.447 | 1.08E-118 | 3 | PTGER4 |
| 1.98E-122 | 0.26529655 | 0.711 | 0.546 | 4.79E-118 | 3 | ARL6IP5 |
| 1.91E-120 | 0.27691517 | 0.775 | 0.615 | 4.64E-116 | 3 | COX7A2 |
| 5.10E-119 | 0.2784164 | 0.997 | 0.993 | 1.24E-114 | 3 | NFKBIA |
| 2.75E-117 | 0.25842188 | 0.78 | 0.627 | 6.66E-113 | 3 | ATP5MF |
| 9.56E-107 | 0.2787483 | 0.707 | 0.555 | 2.32E-102 | 3 | NOP10 |
| 1.15E-103 | 0.2748917 | 0.82 | 0.667 | 2.78E-99 | 3 | ANXA2 |
| 9.23E-100 | 0.27686565 | 0.491 | 0.34 | 2.24E-95 | 3 | PRDM1 |
| 2.07E-99 | 0.26512329 | 0.559 | 0.398 | 5.02E-95 | 3 | IFI6 |
| 6.31E-99 | 0.2977334 | 0.912 | 0.825 | 1.53E-94 | 3 | TNFAIP3 |
| 1.82E-91 | 0.25182542 | 0.923 | 0.873 | 4.42E-87 | 3 | CTSZ |
| 4.01E-91 | 0.37111563 | 0.927 | 0.9 | 9.72E-87 | 3 | CCL4L2 |
| 6.66E-90 | 0.25526668 | 0.697 | 0.549 | 1.62E-85 | 3 | TAGAP |
| 9.73E-88 | 0.35672714 | 0.959 | 0.945 | 2.36E-83 | 3 | CCL4 |
| 7.64E-79 | 0.36452517 | 0.587 | 0.466 | 1.85E-74 | 3 | CXCL3 |
| 2.25E-42 | 0.32690842 | 0.76 | 0.652 | 5.47E-38 | 3 | G0S2 |
| 0 | 3.54733525 | 0.999 | 0.307 | 0 | 4 | CCL20 |
| 0 | 1.32947454 | 1 | 0.79 | 0 | 4 | IL1B |
| 0 | 1.21273102 | 0.902 | 0.586 | 0 | 4 | PTGS2 |
| 0 | 1.18730087 | 0.95 | 0.597 | 0 | 4 | EREG |
| 0 | 1.09771196 | 0.993 | 0.695 | 0 | 4 | OLR1 |
| 0 | 1.05597365 | 0.962 | 0.647 | 0 | 4 | G0S2 |
| 0 | 0.91657153 | 0.999 | 0.977 | 0 | 4 | CXCL8 |
| 0 | 0.90306529 | 0.982 | 0.705 | 0 | 4 | GPR183 |
| 0 | 0.88846277 | 0.999 | 0.791 | 0 | 4 | BCL2A1 |
| 0 | 0.81755628 | 0.998 | 0.836 | 0 | 4 | PLAUR |
| 0 | 0.74806849 | 1 | 0.977 | 0 | 4 | SRGN |
| 0 | 0.72262197 | 1 | 0.984 | 0 | 4 | SOD2 |
| 0 | 0.72152143 | 0.739 | 0.379 | 0 | 4 | AQP9 |
| 0 | 0.71270053 | 0.99 | 0.759 | 0 | 4 | CD44 |
| 0 | 0.54949702 | 1 | 1 | 0 | 4 | FTH1 |
| 5.80E-300 | 0.88286068 | 0.996 | 0.914 | 1.41E-295 | 4 | CCL3L1 |
| 3.90E-297 | 0.74063293 | 0.846 | 0.544 | 9.45E-293 | 4 | SERPINB9 |
| 9.43E-285 | 0.89415987 | 0.913 | 0.621 | 2.29E-280 | 4 | C15orf48 |
| 2.35E-276 | 0.60953897 | 0.98 | 0.87 | 5.70E-272 | 4 | MAP3K8 |
| 1.62E-269 | 0.93233109 | 0.794 | 0.494 | 3.93E-265 | 4 | IL1RN |
| 6.93E-257 | 0.64295015 | 0.803 | 0.5 | 1.68E-252 | 4 | EHD1 |
| 5.28E-254 | 0.79093216 | 0.947 | 0.754 | 1.28E-249 | 4 | CXCL2 |
| 9.41E-247 | 0.59203452 | 0.999 | 0.967 | 2.28E-242 | 4 | BTG1 |
| 2.40E-235 | 0.68737442 | 0.649 | 0.342 | 5.81E-231 | 4 | LUCAT1 |
| 5.70E-235 | 0.66574059 | 0.866 | 0.633 | 1.38E-230 | 4 | NLRP3 |
| 2.63E-230 | 0.77442111 | 0.453 | 0.195 | 6.37E-226 | 4 | DNAAF1 |

| | | | | | | |
|---|---|---|---|---|---|---|
| 6.36E-226 | 0.59437696 | 1 | 0.782 | 1.54E-221 | 4 | VIM |
| 8.19E-220 | 0.44060368 | 0.389 | 0.152 | 1.99E-215 | 4 | TRAF1 |
| 6.26E-219 | 0.63339777 | 0.788 | 0.543 | 1.52E-214 | 4 | NFKB1 |
| 2.51E-214 | 0.4507593 | 1 | 0.995 | 6.09E-210 | 4 | RPS28 |
| 1.88E-213 | 0.56388363 | 0.978 | 0.826 | 4.55E-209 | 4 | TNFAIP3 |
| 1.68E-210 | 0.63710724 | 0.596 | 0.322 | 4.08E-206 | 4 | BASP1 |
| 1.14E-208 | 0.44549541 | 1 | 0.995 | 2.76E-204 | 4 | RPS27 |
| 3.87E-208 | 0.66907367 | 0.806 | 0.554 | 9.37E-204 | 4 | INSIG1 |
| 3.21E-206 | 0.55930406 | 0.889 | 0.609 | 7.78E-202 | 4 | UPP1 |
| 2.67E-202 | 0.4794905 | 0.996 | 0.98 | 6.47E-198 | 4 | PNRC1 |
| 2.21E-193 | 0.7886133 | 0.35 | 0.139 | 5.35E-189 | 4 | AREG |
| 4.91E-193 | 0.55797072 | 0.806 | 0.591 | 1.19E-188 | 4 | TNFAIP8 |
| 6.61E-193 | 0.39051887 | 1 | 0.999 | 1.60E-188 | 4 | RPL41 |
| 1.01E-191 | 0.56792037 | 0.85 | 0.686 | 2.45E-187 | 4 | CFLAR |
| 2.24E-185 | 0.40405785 | 1 | 0.995 | 5.43E-181 | 4 | RPS29 |
| 1.89E-180 | 0.54000453 | 0.832 | 0.634 | 4.57E-176 | 4 | ATP2B1 |
| 3.34E-180 | 0.4042129 | 1 | 0.995 | 8.09E-176 | 4 | RPL28 |
| 6.16E-180 | 0.51064485 | 1 | 0.804 | 1.49E-175 | 4 | LYZ |
| 2.23E-178 | 0.55598568 | 0.692 | 0.434 | 5.41E-174 | 4 | ATP13A3 |
| 5.25E-176 | 0.42895156 | 0.279 | 0.1 | 1.27E-171 | 4 | IL1A |
| 1.46E-175 | 0.38148099 | 1 | 0.997 | 3.55E-171 | 4 | RPL39 |
| 2.18E-175 | 0.46859294 | 0.561 | 0.304 | 5.29E-171 | 4 | SLC7A5 |
| 5.52E-174 | 0.40621083 | 1 | 0.981 | 1.34E-169 | 4 | RPL35A |
| 5.47E-173 | 0.48769081 | 0.965 | 0.868 | 1.33E-168 | 4 | HIF1A |
| 6.35E-172 | 0.52167104 | 0.723 | 0.481 | 1.54E-167 | 4 | FAM107B |
| 4.95E-169 | 0.57212053 | 0.669 | 0.419 | 1.20E-164 | 4 | NR4A3 |
| 2.97E-168 | 0.38677722 | 0.999 | 0.992 | 7.20E-164 | 4 | RPL30 |
| 6.12E-168 | 0.55854802 | 0.552 | 0.305 | 1.48E-163 | 4 | BIRC3 |
| 4.50E-166 | 0.37903661 | 1 | 0.993 | 1.09E-161 | 4 | RPS14 |
| 8.83E-166 | 0.38064117 | 1 | 0.995 | 2.14E-161 | 4 | RPL34 |
| 1.10E-164 | 0.519845 | 0.872 | 0.649 | 2.67E-160 | 4 | VEGFA |
| 6.40E-163 | 0.71287476 | 0.976 | 0.899 | 1.55E-158 | 4 | CCL4L2 |
| 7.94E-161 | 0.3631063 | 1 | 0.985 | 1.92E-156 | 4 | UBA52 |
| 5.23E-160 | 0.43690176 | 0.446 | 0.221 | 1.27E-155 | 4 | GPR132 |
| 9.97E-160 | 0.37014161 | 1 | 0.996 | 2.42E-155 | 4 | RPS12 |
| 6.14E-159 | 0.36365602 | 1 | 0.987 | 1.49E-154 | 4 | FAU |
| 6.92E-159 | 0.58309733 | 0.878 | 0.652 | 1.68E-154 | 4 | TIMP1 |
| 3.01E-157 | 0.45582016 | 0.986 | 0.716 | 7.29E-153 | 4 | S100A10 |
| 3.20E-153 | 0.37714843 | 0.999 | 0.978 | 7.77E-149 | 4 | RPL18 |
| 5.46E-152 | 0.48475787 | 0.841 | 0.633 | 1.32E-147 | 4 | GPCPD1 |
| 3.19E-150 | 0.51215459 | 0.601 | 0.381 | 7.73E-146 | 4 | ELL2 |
| 1.60E-149 | 0.37062812 | 1 | 0.981 | 3.88E-145 | 4 | RPS21 |
| 2.07E-149 | 0.54416276 | 1 | 0.992 | 5.03E-145 | 4 | CCL3 |

| | | | | | | |
|---|---|---|---|---|---|---|
| 1.66E-148 | 0.3353289 | 1 | 0.999 | 4.01E-144 | 4 | RPLP1 |
| 4.22E-146 | 0.31740138 | 1 | 0.993 | 1.02E-141 | 4 | RPLP2 |
| 4.92E-146 | 0.35030557 | 1 | 0.999 | 1.19E-141 | 4 | RPL10 |
| 1.77E-144 | 0.44089317 | 0.814 | 0.618 | 4.30E-140 | 4 | PDE4B |
| 2.67E-144 | 0.3523625 | 1 | 0.995 | 6.46E-140 | 4 | RPL11 |
| 1.70E-143 | 0.42845235 | 0.415 | 0.208 | 4.13E-139 | 4 | THAP2 |
| 2.07E-142 | 0.49225814 | 0.706 | 0.481 | 5.01E-138 | 4 | SLC25A37 |
| 5.20E-140 | 0.35272804 | 0.998 | 0.99 | 1.26E-135 | 4 | RPS27A |
| 6.20E-139 | 0.35195096 | 0.997 | 0.991 | 1.50E-134 | 4 | RPS15A |
| 9.79E-139 | 0.33835273 | 1 | 0.989 | 2.37E-134 | 4 | RPS15 |
| 2.54E-137 | 0.34088134 | 0.999 | 0.987 | 6.15E-133 | 4 | RPL36 |
| 1.30E-136 | 0.46799794 | 0.671 | 0.46 | 3.16E-132 | 4 | RILPL2 |
| 1.49E-135 | 0.34805492 | 0.999 | 0.986 | 3.61E-131 | 4 | RPS13 |
| 6.41E-135 | 0.31577847 | 1 | 0.997 | 1.56E-130 | 4 | RPS24 |
| 3.63E-134 | 0.34265463 | 1 | 0.986 | 8.79E-130 | 4 | RPL18A |
| 3.79E-134 | 0.34670754 | 1 | 0.99 | 9.19E-130 | 4 | RPS19 |
| 1.64E-133 | 0.47164417 | 0.765 | 0.547 | 3.97E-129 | 4 | PPIF |
| 1.92E-133 | 0.45330828 | 0.909 | 0.758 | 4.66E-129 | 4 | ATP2B1-AS1 |
| 8.69E-131 | 0.37464754 | 0.953 | 0.792 | 2.11E-126 | 4 | CDKN1A |
| 2.55E-130 | 0.4159959 | 0.465 | 0.256 | 6.17E-126 | 4 | CCRL2 |
| 8.39E-130 | 0.4796232 | 0.685 | 0.445 | 2.03E-125 | 4 | PHLDA1 |
| 6.62E-129 | 0.57282023 | 0.831 | 0.685 | 1.60E-124 | 4 | CHMP1B |
| 2.75E-128 | 0.33418025 | 0.999 | 0.992 | 6.67E-124 | 4 | RPL26 |
| 1.71E-127 | 0.32620242 | 1 | 0.988 | 4.16E-123 | 4 | RPS9 |
| 3.76E-127 | 0.42955587 | 0.999 | 0.749 | 9.12E-123 | 4 | S100A6 |
| 4.50E-127 | 0.41130083 | 0.994 | 0.773 | 1.09E-122 | 4 | S100A4 |
| 1.36E-126 | 0.31274919 | 1 | 0.994 | 3.29E-122 | 4 | RPL32 |
| 6.55E-126 | 0.44948545 | 0.537 | 0.315 | 1.59E-121 | 4 | CYTOR |
| 4.61E-125 | 0.3077436 | 1 | 0.991 | 1.12E-120 | 4 | RPL9 |
| 1.05E-122 | 0.34838041 | 0.999 | 0.98 | 2.54E-118 | 4 | RPS4X |
| 9.21E-122 | 0.43868356 | 0.474 | 0.269 | 2.23E-117 | 4 | MIR155HG |
| 1.37E-120 | 0.29190543 | 1 | 0.996 | 3.31E-116 | 4 | RPL37 |
| 5.31E-120 | 0.39812773 | 0.849 | 0.638 | 1.29E-115 | 4 | SERPINA1 |
| 5.34E-120 | 0.39034401 | 0.492 | 0.283 | 1.29E-115 | 4 | AC016831.1 |
| 6.24E-120 | 0.28585678 | 1 | 0.998 | 1.51E-115 | 4 | RPL13 |
| 1.02E-117 | 0.34184415 | 1 | 0.965 | 2.47E-113 | 4 | HLA-DRA |
| 3.74E-117 | 0.31096538 | 1 | 0.979 | 9.06E-113 | 4 | RPL24 |
| 6.65E-116 | 0.3149416 | 1 | 0.994 | 1.61E-111 | 4 | RPS8 |
| 5.56E-115 | 0.33533515 | 0.999 | 0.93 | 1.35E-110 | 4 | S100A11 |
| 2.95E-110 | 0.29904998 | 0.998 | 0.985 | 7.16E-106 | 4 | RPL29 |
| 3.52E-110 | 0.32703133 | 0.36 | 0.183 | 8.53E-106 | 4 | SEMA6B |
| 9.13E-110 | 0.40381644 | 0.884 | 0.758 | 2.21E-105 | 4 | C5AR1 |
| 9.16E-107 | 0.29590292 | 0.999 | 0.99 | 2.22E-102 | 4 | RPS3A |

| | | | | | | |
|---|---|---|---|---|---|---|
| 1.49E-106 | 0.38731066 | 0.534 | 0.338 | 3.62E-102 | 4 | KYNU |
| 3.54E-106 | 0.29583143 | 0.999 | 0.984 | 8.57E-102 | 4 | RPL17 |
| 7.09E-105 | 0.35358406 | 0.993 | 0.944 | 1.72E-100 | 4 | NAMPT |
| 7.84E-105 | 0.30277121 | 0.999 | 0.978 | 1.90E-100 | 4 | ATP5F1E |
| 1.80E-103 | 0.37116403 | 0.402 | 0.223 | 4.36E-99 | 4 | RASGRP3 |
| 2.44E-103 | 0.33696362 | 0.969 | 0.826 | 5.91E-99 | 4 | HLA-DPB1 |
| 6.96E-103 | 0.26436612 | 0.513 | 0.311 | 1.69E-98 | 4 | FN1 |
| 2.24E-102 | 0.35029345 | 0.513 | 0.318 | 5.44E-98 | 4 | RNF19B |
| 6.34E-102 | 0.39208028 | 0.637 | 0.463 | 1.54E-97 | 4 | ARL8B |
| 1.15E-101 | 0.44326363 | 0.766 | 0.577 | 2.78E-97 | 4 | RGCC |
| 2.68E-101 | 0.2844575 | 0.999 | 0.99 | 6.49E-97 | 4 | RPL12 |
| 9.41E-101 | 0.41813741 | 0.658 | 0.494 | 2.28E-96 | 4 | WTAP |
| 5.01E-100 | 0.35048322 | 0.999 | 0.979 | 1.21E-95 | 4 | IER3 |
| 5.52E-100 | 0.28436523 | 1 | 0.989 | 1.34E-95 | 4 | RPL15 |
| 7.16E-100 | 0.27038273 | 1 | 0.991 | 1.74E-95 | 4 | RPL38 |
| 1.31E-98 | 0.38629179 | 0.827 | 0.67 | 3.18E-94 | 4 | MXD1 |
| 4.05E-98 | 0.39758043 | 0.538 | 0.354 | 9.82E-94 | 4 | FNDC3B |
| 4.73E-98 | 0.39398724 | 0.513 | 0.33 | 1.15E-93 | 4 | USP12 |
| 6.60E-98 | 0.2875354 | 1 | 0.977 | 1.60E-93 | 4 | RPS7 |
| 8.81E-98 | 0.28899704 | 0.999 | 0.984 | 2.14E-93 | 4 | RPL21 |
| 1.12E-97 | 0.28903042 | 1 | 0.993 | 2.72E-93 | 4 | RPS2 |
| 1.61E-97 | 0.33761732 | 0.434 | 0.253 | 3.89E-93 | 4 | B4GALT5 |
| 1.63E-97 | 0.26256368 | 1 | 0.995 | 3.96E-93 | 4 | RPL37A |
| 4.53E-97 | 0.31302183 | 0.979 | 0.736 | 1.10E-92 | 4 | LGALS1 |
| 4.71E-96 | 0.26726051 | 1 | 0.997 | 1.14E-91 | 4 | RPS23 |
| 7.46E-96 | 0.28568751 | 0.998 | 0.974 | 1.81E-91 | 4 | RPS3 |
| 2.81E-95 | 0.26235787 | 0.999 | 0.988 | 6.82E-91 | 4 | RPS11 |
| 5.80E-95 | 0.37906562 | 0.806 | 0.682 | 1.41E-90 | 4 | DSE |
| 7.63E-95 | 0.28369699 | 0.998 | 0.971 | 1.85E-90 | 4 | PFDN5 |
| 6.76E-94 | 0.2640479 | 1 | 0.985 | 1.64E-89 | 4 | RPS25 |
| 9.47E-94 | 0.37448154 | 0.775 | 0.632 | 2.30E-89 | 4 | LITAF |
| 1.29E-93 | 0.3776126 | 0.974 | 0.855 | 3.14E-89 | 4 | PLIN2 |
| 1.54E-93 | 0.42508751 | 0.59 | 0.382 | 3.73E-89 | 4 | APOBEC3A |
| 2.53E-93 | 0.37096095 | 0.624 | 0.442 | 6.12E-89 | 4 | JARID2 |
| 5.25E-93 | 0.3448385 | 0.791 | 0.636 | 1.27E-88 | 4 | FOSL2 |
| 8.54E-93 | 0.35467067 | 0.751 | 0.533 | 2.07E-88 | 4 | AC020656.1 |
| 2.89E-91 | 0.27656623 | 1 | 0.995 | 7.01E-87 | 4 | RPS18 |
| 3.00E-91 | 0.3764786 | 0.62 | 0.456 | 7.27E-87 | 4 | BIRC2 |
| 4.51E-90 | 0.28803982 | 0.985 | 0.869 | 1.09E-85 | 4 | HLA-DPA1 |
| 5.58E-90 | 0.2804996 | 0.998 | 0.973 | 1.35E-85 | 4 | RPL8 |
| 3.55E-89 | 0.36311427 | 0.674 | 0.496 | 8.60E-85 | 4 | LIMS1 |
| 1.07E-87 | 0.73177309 | 0.626 | 0.471 | 2.60E-83 | 4 | CXCL3 |
| 3.15E-87 | 0.34472391 | 0.622 | 0.415 | 7.63E-83 | 4 | FLNA |

| | | | | | | |
|---|---|---|---|---|---|---|
| 1.74E-86 | 0.25979784 | 0.998 | 0.982 | 4.21E-82 | 4 | RPL35 |
| 2.98E-86 | 0.34204742 | 0.633 | 0.451 | 7.22E-82 | 4 | MAP2K3 |
| 1.78E-84 | 0.27672952 | 0.996 | 0.962 | 4.31E-80 | 4 | RPL27 |
| 3.88E-84 | 0.27310766 | 0.993 | 0.956 | 9.41E-80 | 4 | RPL7 |
| 6.22E-84 | 0.26331063 | 1 | 0.999 | 1.51E-79 | 4 | SAT1 |
| 1.54E-83 | 0.31301205 | 0.452 | 0.284 | 3.74E-79 | 4 | RABGEF1 |
| 6.37E-83 | 0.2832015 | 0.935 | 0.792 | 1.55E-78 | 4 | EMP3 |
| 1.57E-82 | 0.33551049 | 0.56 | 0.38 | 3.80E-78 | 4 | GBP2 |
| 1.58E-82 | 0.38282668 | 0.764 | 0.639 | 3.83E-78 | 4 | LDHA |
| 4.85E-82 | 0.41113202 | 0.343 | 0.196 | 1.17E-77 | 4 | NAF1 |
| 2.09E-81 | 0.28659788 | 0.252 | 0.122 | 5.08E-77 | 4 | PALM2-AKAP |
| 2.71E-81 | 0.31398597 | 0.631 | 0.432 | 6.58E-77 | 4 | TREM1 |
| 3.99E-81 | 0.40342685 | 0.631 | 0.47 | 9.68E-77 | 4 | EIF4E |
| 6.11E-81 | 0.32150142 | 0.634 | 0.434 | 1.48E-76 | 4 | ITGAX |
| 1.18E-79 | 0.33676787 | 0.464 | 0.295 | 2.86E-75 | 4 | TNIP1 |
| 1.05E-78 | 0.36296363 | 0.52 | 0.336 | 2.55E-74 | 4 | CD300E |
| 2.43E-78 | 0.31737459 | 0.667 | 0.484 | 5.90E-74 | 4 | SLC11A1 |
| 1.57E-76 | 0.26419552 | 0.855 | 0.638 | 3.82E-72 | 4 | LGALS3 |
| 2.29E-76 | 0.29734565 | 0.511 | 0.326 | 5.54E-72 | 4 | CD9 |
| 3.39E-76 | 0.33017606 | 0.727 | 0.583 | 8.22E-72 | 4 | KDM6B |
| 1.04E-75 | 0.27783475 | 0.987 | 0.938 | 2.53E-71 | 4 | RPL10A |
| 3.07E-74 | 0.31423819 | 0.538 | 0.377 | 7.43E-70 | 4 | ZNF267 |
| 8.41E-74 | 0.30353427 | 0.833 | 0.676 | 2.04E-69 | 4 | SLC2A3 |
| 1.43E-73 | 0.28476734 | 0.893 | 0.798 | 3.48E-69 | 4 | SERP1 |
| 2.62E-73 | 0.39491386 | 0.574 | 0.431 | 6.36E-69 | 4 | SAMSN1 |
| 5.21E-73 | 0.30928804 | 0.797 | 0.668 | 1.26E-68 | 4 | COMMD6 |
| 6.56E-73 | 0.35821548 | 0.615 | 0.452 | 1.59E-68 | 4 | ACSL1 |
| 9.81E-73 | 0.35422488 | 0.735 | 0.62 | 2.38E-68 | 4 | ICAM1 |
| 2.01E-72 | 0.33073646 | 0.704 | 0.552 | 4.88E-68 | 4 | CD55 |
| 4.29E-71 | 0.27696503 | 0.919 | 0.865 | 1.04E-66 | 4 | HNRNPC |
| 6.59E-70 | 0.27033072 | 0.337 | 0.195 | 1.60E-65 | 4 | CD109 |
| 2.56E-69 | 0.28086167 | 0.656 | 0.492 | 6.20E-65 | 4 | PIM3 |
| 2.67E-69 | 0.25563682 | 0.988 | 0.951 | 6.47E-65 | 4 | RPL5 |
| 4.90E-69 | 0.32959281 | 0.312 | 0.172 | 1.19E-64 | 4 | THBS1 |
| 5.55E-69 | 0.29186799 | 0.456 | 0.297 | 1.35E-64 | 4 | SORL1 |
| 5.79E-69 | 0.2795914 | 0.928 | 0.867 | 1.40E-64 | 4 | TOMM7 |
| 8.15E-69 | 0.2628075 | 0.954 | 0.868 | 1.98E-64 | 4 | ZFAS1 |
| 6.48E-68 | 0.42005862 | 0.653 | 0.512 | 1.57E-63 | 4 | ZNF331 |
| 2.24E-67 | 0.31461968 | 0.41 | 0.264 | 5.43E-63 | 4 | EDEM1 |
| 6.62E-67 | 0.25636629 | 0.962 | 0.899 | 1.61E-62 | 4 | COX7C |
| 6.82E-67 | 0.29730942 | 0.496 | 0.337 | 1.65E-62 | 4 | GRASP |
| 2.88E-66 | 0.25974242 | 0.585 | 0.404 | 6.98E-62 | 4 | CD48 |
| 7.69E-66 | 0.28109118 | 0.474 | 0.317 | 1.87E-61 | 4 | AZIN1-AS1 |

| | | | | | | |
|---|---|---|---|---|---|---|
| 1.38E-65 | 0.29161335 | 0.918 | 0.839 | 3.35E-61 | 4 | NINJ1 |
| 6.05E-65 | 0.27088384 | 0.962 | 0.895 | 1.47E-60 | 4 | RPS5 |
| 7.55E-64 | 0.38870228 | 0.82 | 0.73 | 1.83E-59 | 4 | PMAIP1 |
| 7.84E-64 | 0.28787812 | 0.423 | 0.278 | 1.90E-59 | 4 | FNIP2 |
| 1.25E-63 | 0.30111413 | 0.604 | 0.451 | 3.04E-59 | 4 | PFKFB3 |
| 4.47E-62 | 0.28671205 | 0.566 | 0.405 | 1.08E-57 | 4 | SNX10 |
| 2.27E-61 | 0.34598647 | 0.658 | 0.544 | 5.50E-57 | 4 | NFE2L2 |
| 3.25E-61 | 0.28329051 | 0.42 | 0.276 | 7.87E-57 | 4 | GK |
| 3.93E-61 | 0.29585633 | 0.75 | 0.635 | 9.54E-57 | 4 | CTNNB1 |
| 5.77E-61 | 0.25545584 | 0.924 | 0.851 | 1.40E-56 | 4 | GPX4 |
| 6.61E-61 | 0.26162273 | 0.486 | 0.333 | 1.60E-56 | 4 | MAPK6 |
| 1.14E-60 | 0.27900834 | 0.936 | 0.882 | 2.76E-56 | 4 | PLEK |
| 1.15E-60 | 0.25955695 | 0.59 | 0.423 | 2.79E-56 | 4 | STX11 |
| 1.80E-59 | 0.26955344 | 0.338 | 0.202 | 4.36E-55 | 4 | MCEMP1 |
| 2.58E-59 | 0.27738988 | 0.734 | 0.607 | 6.26E-55 | 4 | GSTO1 |
| 8.23E-59 | 0.26871191 | 0.34 | 0.207 | 1.99E-54 | 4 | ANPEP |
| 3.02E-56 | 0.2562617 | 0.903 | 0.821 | 7.32E-52 | 4 | RPSA |
| 1.53E-55 | 0.29436768 | 0.363 | 0.228 | 3.70E-51 | 4 | SDS |
| 1.84E-54 | 0.3595927 | 0.513 | 0.39 | 4.47E-50 | 4 | RANBP2 |
| 5.26E-54 | 0.26640104 | 0.406 | 0.27 | 1.28E-49 | 4 | OASL |
| 1.99E-53 | 0.28524282 | 0.57 | 0.439 | 4.82E-49 | 4 | SLC43A2 |
| 2.60E-53 | 0.30282344 | 0.729 | 0.656 | 6.32E-49 | 4 | REL |
| 1.60E-52 | 0.25754736 | 0.763 | 0.667 | 3.89E-48 | 4 | LCP2 |
| 1.71E-52 | 0.26234215 | 0.55 | 0.405 | 4.14E-48 | 4 | CSF3R |
| 1.49E-49 | 0.25142909 | 0.437 | 0.307 | 3.62E-45 | 4 | ETV3 |
| 1.95E-49 | 0.31290555 | 0.434 | 0.304 | 4.73E-45 | 4 | THBD |
| 2.15E-48 | 0.3054537 | 0.669 | 0.539 | 5.22E-44 | 4 | AC253572.2 |
| 2.12E-47 | 0.32559906 | 0.362 | 0.238 | 5.14E-43 | 4 | RETN |
| 9.01E-46 | 0.2670028 | 0.41 | 0.282 | 2.19E-41 | 4 | C3 |
| 6.22E-45 | 0.26740512 | 0.492 | 0.372 | 1.51E-40 | 4 | IVNS1ABP |
| 3.08E-44 | 0.38219282 | 0.667 | 0.587 | 7.48E-40 | 4 | SELENOK |
| 3.28E-44 | 0.30763825 | 0.618 | 0.515 | 7.94E-40 | 4 | PELI1 |
| 5.64E-44 | 0.46002438 | 0.965 | 0.946 | 1.37E-39 | 4 | CCL4 |
| 8.82E-44 | 0.25125209 | 0.438 | 0.32 | 2.14E-39 | 4 | PTPN1 |
| 1.35E-43 | 0.26278051 | 0.609 | 0.496 | 3.26E-39 | 4 | TIPARP |
| 6.05E-43 | 0.28722441 | 0.679 | 0.59 | 1.47E-38 | 4 | SERPINB1 |
| 4.44E-37 | 0.27149138 | 0.592 | 0.498 | 1.08E-32 | 4 | MAP3K2 |
| 2.69E-35 | 0.28975843 | 0.617 | 0.535 | 6.52E-31 | 4 | EIF1B |
| 5.50E-31 | 0.27085741 | 0.479 | 0.379 | 1.33E-26 | 4 | HLA-DRB5 |
| 2.34E-27 | 0.30769159 | 0.679 | 0.611 | 5.68E-23 | 4 | H1FX |
| 6.37E-16 | 0.26314057 | 0.469 | 0.406 | 1.54E-11 | 4 | HIST1H1C |
| 3.18E-14 | 0.29247359 | 0.266 | 0.213 | 7.70E-10 | 4 | TWISTNB |
| 0 | 3.97478437 | 0.995 | 0.321 | 0 | 5 | HSPA6 |

| | | | | | | |
|---|---|---|---|---|---|---|
| 0 | 1.66854111 | 0.977 | 0.76 | 0 | 5 | HSPB1 |
| 0 | 1.57814568 | 0.968 | 0.62 | 0 | 5 | HSPH1 |
| 0 | 1.53927466 | 0.798 | 0.307 | 0 | 5 | BAG3 |
| 0 | 1.50371115 | 1 | 0.918 | 0 | 5 | HSPA1A |
| 0 | 1.49936587 | 1 | 0.797 | 0 | 5 | HSPA1B |
| 0 | 1.48934359 | 1 | 0.831 | 0 | 5 | DNAJB1 |
| 0 | 1.44562966 | 0.966 | 0.714 | 0 | 5 | HSPD1 |
| 0 | 1.40413071 | 0.973 | 0.812 | 0 | 5 | IER5 |
| 0 | 1.38057489 | 0.998 | 0.956 | 0 | 5 | JUN |
| 0 | 1.34648578 | 1 | 0.989 | 0 | 5 | HSP90AA1 |
| 0 | 1.34282217 | 0.772 | 0.372 | 0 | 5 | ZFAND2A |
| 0 | 1.22373279 | 0.931 | 0.661 | 0 | 5 | RHOB |
| 0 | 1.2086893 | 0.974 | 0.789 | 0 | 5 | DNAJA1 |
| 0 | 1.20521314 | 0.966 | 0.807 | 0 | 5 | KLF2 |
| 0 | 1.19364037 | 0.997 | 0.964 | 0 | 5 | HSP90AB1 |
| 0 | 1.17786658 | 0.808 | 0.431 | 0 | 5 | SELENOP |
| 0 | 1.09602169 | 0.997 | 0.982 | 0 | 5 | FOS |
| 0 | 1.04115512 | 0.958 | 0.763 | 0 | 5 | HSPE1 |
| 0 | 1.02688709 | 0.977 | 0.839 | 0 | 5 | TRA2B |
| 0 | 0.97565007 | 0.595 | 0.226 | 0 | 5 | DNAJA4 |
| 0 | 0.92669829 | 0.998 | 0.993 | 0 | 5 | DUSP1 |
| 0 | 0.89877496 | 0.992 | 0.949 | 0 | 5 | JUNB |
| 0 | 0.8864784 | 0.996 | 0.992 | 0 | 5 | KLF6 |
| 0 | 0.76947728 | 0.725 | 0.366 | 0 | 5 | CHORDC1 |
| 5.67E-300 | 0.83415228 | 0.898 | 0.652 | 1.37E-295 | 5 | CLK1 |
| 2.76E-295 | 0.94396752 | 0.982 | 0.917 | 6.70E-291 | 5 | IER2 |
| 1.36E-294 | 0.9261235 | 0.958 | 0.829 | 3.29E-290 | 5 | HSPA8 |
| 2.39E-284 | 1.05328578 | 0.568 | 0.236 | 5.78E-280 | 5 | DDX3Y |
| 1.48E-283 | 1.17414161 | 0.902 | 0.741 | 3.60E-279 | 5 | RGS2 |
| 1.13E-282 | 0.90873348 | 0.992 | 0.969 | 2.75E-278 | 5 | GADD45B |
| 1.85E-282 | 1.00148151 | 0.905 | 0.673 | 4.49E-278 | 5 | C1QA |
| 1.66E-272 | 0.79013492 | 0.983 | 0.926 | 4.03E-268 | 5 | UBB |
| 3.96E-269 | 0.75037208 | 0.638 | 0.298 | 9.60E-265 | 5 | DNAJB4 |
| 2.77E-264 | 0.87633004 | 0.844 | 0.53 | 6.72E-260 | 5 | C1QC |
| 7.26E-264 | 0.92062101 | 0.876 | 0.6 | 1.76E-259 | 5 | C1QB |
| 3.20E-259 | 0.63354913 | 1 | 1 | 7.77E-255 | 5 | UBC |
| 3.01E-253 | 0.97035065 | 0.879 | 0.643 | 7.29E-249 | 5 | EGR1 |
| 1.10E-251 | 0.87675882 | 0.657 | 0.333 | 2.68E-247 | 5 | NEU1 |
| 4.03E-251 | 0.76217509 | 0.716 | 0.355 | 9.78E-247 | 5 | FOLR2 |
| 2.67E-250 | 0.86615547 | 0.762 | 0.413 | 6.47E-246 | 5 | MAF |
| 1.02E-248 | 0.85030233 | 0.659 | 0.301 | 2.46E-244 | 5 | LYVE1 |
| 1.65E-247 | 0.98921351 | 0.733 | 0.398 | 4.00E-243 | 5 | PLTP |
| 1.14E-241 | 1.00942155 | 0.807 | 0.502 | 2.76E-237 | 5 | RNASE1 |

| | | | | | | |
|---|---|---|---|---|---|---|
| 3.71E-236 | 0.68635854 | 0.609 | 0.291 | 8.98E-232 | 5 | MYLIP |
| 9.70E-234 | 0.83084264 | 0.843 | 0.594 | 2.35E-229 | 5 | BTG2 |
| 2.22E-225 | 0.85962982 | 0.737 | 0.412 | 5.39E-221 | 5 | F13A1 |
| 4.37E-225 | 0.88234697 | 0.793 | 0.499 | 1.06E-220 | 5 | DAB2 |
| 4.68E-222 | 0.65822788 | 0.697 | 0.392 | 1.13E-217 | 5 | TCP1 |
| 2.37E-220 | 0.71763713 | 0.764 | 0.447 | 5.74E-216 | 5 | STAB1 |
| 5.97E-214 | 1.00593719 | 0.848 | 0.632 | 1.45E-209 | 5 | HMOX1 |
| 9.01E-213 | 0.8517325 | 0.963 | 0.911 | 2.19E-208 | 5 | KLF4 |
| 1.75E-211 | 0.66437595 | 0.618 | 0.312 | 4.24E-207 | 5 | ATP1B1 |
| 4.07E-210 | 0.70040131 | 0.735 | 0.43 | 9.87E-206 | 5 | MS4A4A |
| 4.03E-204 | 0.52246972 | 0.498 | 0.212 | 9.77E-200 | 5 | LILRB5 |
| 6.08E-204 | 0.48557772 | 0.615 | 0.287 | 1.47E-199 | 5 | GPR34 |
| 8.69E-203 | 0.82697681 | 0.804 | 0.545 | 2.11E-198 | 5 | LGMN |
| 1.25E-202 | 0.81670942 | 0.866 | 0.682 | 3.02E-198 | 5 | HERPUD1 |
| 1.11E-200 | 0.62970465 | 0.58 | 0.307 | 2.69E-196 | 5 | CACYBP |
| 3.83E-195 | 0.47948164 | 0.389 | 0.149 | 9.29E-191 | 5 | CD209 |
| 8.11E-191 | 0.83610724 | 0.786 | 0.558 | 1.97E-186 | 5 | IFRD1 |
| 2.47E-189 | 0.84416839 | 0.93 | 0.8 | 5.99E-185 | 5 | MAFB |
| 1.84E-186 | 1.0666797 | 0.929 | 0.837 | 4.45E-182 | 5 | HSPA5 |
| 2.02E-186 | 0.51653523 | 0.647 | 0.342 | 4.90E-182 | 5 | FRMD4B |
| 1.70E-185 | 0.70242248 | 0.988 | 0.978 | 4.11E-181 | 5 | FOSB |
| 1.83E-181 | 0.56152936 | 0.629 | 0.336 | 4.44E-177 | 5 | SLCO2B1 |
| 2.08E-180 | 0.58149214 | 0.99 | 0.963 | 5.05E-176 | 5 | ITM2B |
| 9.63E-179 | 0.38364136 | 0.285 | 0.093 | 2.33E-174 | 5 | FCGBP |
| 5.63E-175 | 0.47197997 | 0.497 | 0.223 | 1.37E-170 | 5 | COLEC12 |
| 6.14E-175 | 0.43422642 | 0.645 | 0.318 | 1.49E-170 | 5 | MRC1 |
| 1.59E-174 | 0.55739536 | 0.908 | 0.746 | 3.85E-170 | 5 | EIF4A2 |
| 4.91E-173 | 0.49682771 | 0.693 | 0.388 | 1.19E-168 | 5 | LPAR6 |
| 1.60E-170 | 0.56147743 | 0.645 | 0.354 | 3.87E-166 | 5 | VSIG4 |
| 4.67E-169 | 0.47172461 | 0.479 | 0.218 | 1.13E-164 | 5 | FSCN1 |
| 1.45E-168 | 0.60952857 | 0.587 | 0.321 | 3.53E-164 | 5 | DDIT3 |
| 5.19E-167 | 0.41602561 | 0.528 | 0.251 | 1.26E-162 | 5 | GAS6 |
| 8.55E-163 | 0.41016771 | 0.474 | 0.21 | 2.07E-158 | 5 | EGFL7 |
| 4.70E-160 | 0.53778873 | 0.544 | 0.292 | 1.14E-155 | 5 | DOK2 |
| 8.46E-160 | 0.5170706 | 0.741 | 0.466 | 2.05E-155 | 5 | ADAP2 |
| 8.47E-156 | 0.40229263 | 0.507 | 0.241 | 2.05E-151 | 5 | ITSN1 |
| 6.25E-155 | 0.38073367 | 0.421 | 0.187 | 1.52E-150 | 5 | OLFML2B |
| 3.40E-154 | 0.56221228 | 0.564 | 0.298 | 8.25E-150 | 5 | PMP22 |
| 1.25E-152 | 0.57605694 | 0.888 | 0.722 | 3.02E-148 | 5 | CTSC |
| 1.16E-151 | 0.73795494 | 0.874 | 0.706 | 2.81E-147 | 5 | SRSF7 |
| 1.74E-151 | 0.42809793 | 0.395 | 0.176 | 4.23E-147 | 5 | FKBP4 |
| 3.58E-151 | 0.3989369 | 0.436 | 0.194 | 8.68E-147 | 5 | SLC40A1 |
| 7.70E-149 | 0.36368153 | 0.415 | 0.18 | 1.87E-144 | 5 | IGF1 |

| | | | | | | |
|---|---|---|---|---|---|---|
| 1.45E-148 | 0.62542744 | 0.538 | 0.293 | 3.50E-144 | 5 | IER5L |
| 2.06E-148 | 0.42217753 | 0.535 | 0.281 | 4.99E-144 | 5 | PIK3R1 |
| 2.37E-147 | 0.48049429 | 0.514 | 0.271 | 5.75E-143 | 5 | MRPL18 |
| 3.24E-146 | 0.41053234 | 0.523 | 0.257 | 7.86E-142 | 5 | NRP1 |
| 3.32E-146 | 0.44511937 | 0.437 | 0.204 | 8.05E-142 | 5 | GATM |
| 9.86E-146 | 0.42470903 | 1 | 1 | 2.39E-141 | 5 | MALAT1 |
| 2.32E-144 | 0.48271839 | 0.999 | 0.991 | 5.63E-140 | 5 | DDX5 |
| 1.06E-143 | 0.30426889 | 0.322 | 0.127 | 2.57E-139 | 5 | TNFRSF25 |
| 2.57E-143 | 0.4852271 | 0.755 | 0.508 | 6.24E-139 | 5 | SNX6 |
| 4.51E-143 | 0.54703832 | 0.905 | 0.793 | 1.09E-138 | 5 | MS4A7 |
| 2.18E-142 | 0.50533086 | 0.907 | 0.789 | 5.29E-138 | 5 | FCGR2A |
| 2.45E-140 | 0.58862423 | 0.807 | 0.599 | 5.94E-136 | 5 | KCTD12 |
| 4.85E-139 | 0.82802791 | 0.857 | 0.756 | 1.18E-134 | 5 | OTUD1 |
| 2.06E-138 | 0.36747281 | 0.442 | 0.203 | 5.00E-134 | 5 | SCN9A |
| 4.25E-137 | 0.40370042 | 0.416 | 0.196 | 1.03E-132 | 5 | FAM13A |
| 8.30E-136 | 0.36293504 | 0.357 | 0.157 | 2.01E-131 | 5 | CGAS |
| 9.57E-136 | 0.54209138 | 0.964 | 0.92 | 2.32E-131 | 5 | CCNL1 |
| 3.63E-135 | 0.30546814 | 0.434 | 0.196 | 8.80E-131 | 5 | HPGDS |
| 7.56E-135 | 0.63013291 | 0.714 | 0.488 | 1.83E-130 | 5 | NUFIP2 |
| 6.88E-134 | 0.44699626 | 0.396 | 0.187 | 1.67E-129 | 5 | TSPYL2 |
| 3.17E-132 | 0.29119736 | 0.317 | 0.129 | 7.67E-128 | 5 | RAB3IL1 |
| 2.43E-131 | 0.64096046 | 0.514 | 0.271 | 5.88E-127 | 5 | PDK4 |
| 2.46E-131 | 0.40228055 | 0.538 | 0.292 | 5.97E-127 | 5 | MKNK1 |
| 5.27E-131 | 0.36491519 | 0.507 | 0.258 | 1.28E-126 | 5 | C2 |
| 7.03E-131 | 0.32317738 | 0.295 | 0.118 | 1.70E-126 | 5 | USPL1 |
| 1.59E-130 | 0.60104496 | 0.933 | 0.874 | 3.85E-126 | 5 | CD14 |
| 2.06E-130 | 0.415776 | 0.605 | 0.359 | 5.00E-126 | 5 | NCF4 |
| 6.44E-129 | 0.52890922 | 0.768 | 0.557 | 1.56E-124 | 5 | MFSD1 |
| 1.86E-125 | 0.32158854 | 0.292 | 0.118 | 4.52E-121 | 5 | OLMALINC |
| 5.55E-125 | 0.43386977 | 0.606 | 0.359 | 1.35E-120 | 5 | MTSS1 |
| 7.65E-125 | 0.40988787 | 0.62 | 0.369 | 1.86E-120 | 5 | C3AR1 |
| 7.41E-124 | 0.32039535 | 0.36 | 0.161 | 1.80E-119 | 5 | FAM20A |
| 8.19E-124 | 0.58120313 | 0.983 | 0.962 | 1.98E-119 | 5 | PPP1R15A |
| 9.65E-124 | 0.36875853 | 0.384 | 0.182 | 2.34E-119 | 5 | ABHD3 |
| 4.98E-123 | 0.30155538 | 0.384 | 0.174 | 1.21E-118 | 5 | CD28 |
| 2.93E-120 | 0.34229064 | 0.502 | 0.265 | 7.09E-116 | 5 | CD59 |
| 5.68E-119 | 0.26045647 | 0.267 | 0.105 | 1.38E-114 | 5 | AC129507.1 |
| 1.68E-118 | 0.32991015 | 0.263 | 0.104 | 4.07E-114 | 5 | PPM1N |
| 2.80E-117 | 0.41350648 | 0.759 | 0.523 | 6.79E-113 | 5 | MEF2C |
| 7.87E-117 | 0.52577151 | 0.757 | 0.561 | 1.91E-112 | 5 | CSF1R |
| 1.97E-116 | 0.31472548 | 0.496 | 0.259 | 4.77E-112 | 5 | SLC9A9 |
| 4.12E-116 | 0.37082798 | 0.379 | 0.178 | 1.00E-111 | 5 | ENPP2 |
| 5.51E-116 | 0.61976604 | 0.532 | 0.294 | 1.34E-111 | 5 | CCL2 |

| | | | | | | |
|---|---|---|---|---|---|---|
| 9.77E-116 | 0.56711292 | 0.45 | 0.244 | 2.37E-111 | 5 | ZBTB10 |
| 2.70E-115 | 0.28971529 | 0.373 | 0.176 | 6.56E-111 | 5 | FRMD4A |
| 2.98E-115 | 0.49030356 | 0.278 | 0.114 | 7.23E-111 | 5 | CH25H |
| 1.01E-112 | 0.62098101 | 0.609 | 0.396 | 2.45E-108 | 5 | CPEB4 |
| 4.96E-112 | 0.3298587 | 0.395 | 0.194 | 1.20E-107 | 5 | TM2D2 |
| 5.88E-112 | 0.48269396 | 0.569 | 0.346 | 1.43E-107 | 5 | GOLGA4 |
| 7.25E-112 | 0.4890161 | 0.541 | 0.325 | 1.76E-107 | 5 | DNAJB11 |
| 5.06E-110 | 0.34011843 | 0.503 | 0.275 | 1.23E-105 | 5 | SIGLEC1 |
| 1.29E-109 | 0.30255372 | 0.438 | 0.223 | 3.14E-105 | 5 | SPRED1 |
| 2.37E-109 | 0.44500537 | 0.567 | 0.347 | 5.74E-105 | 5 | ATP6V0A1 |
| 8.53E-109 | 0.58762053 | 0.925 | 0.856 | 2.07E-104 | 5 | HNRNPU |
| 2.09E-108 | 0.38946905 | 0.644 | 0.416 | 5.08E-104 | 5 | RASSF4 |
| 5.26E-108 | 0.4327376 | 0.597 | 0.387 | 1.28E-103 | 5 | SNHG32 |
| 7.14E-108 | 0.47026109 | 0.794 | 0.6 | 1.73E-103 | 5 | WSB1 |
| 1.67E-106 | 0.34553422 | 0.559 | 0.321 | 4.06E-102 | 5 | ITPR2 |
| 2.06E-106 | 0.31826568 | 0.33 | 0.154 | 4.99E-102 | 5 | AL450998.2 |
| 1.42E-104 | 0.40058944 | 0.461 | 0.253 | 3.45E-100 | 5 | AL499604.1 |
| 2.53E-104 | 0.62851245 | 0.756 | 0.577 | 6.14E-100 | 5 | ARRDC3 |
| 9.97E-104 | 0.31358734 | 0.39 | 0.194 | 2.42E-99 | 5 | RGL1 |
| 1.00E-103 | 0.36000963 | 0.431 | 0.227 | 2.43E-99 | 5 | NFIA |
| 6.23E-103 | 0.3442447 | 0.398 | 0.208 | 1.51E-98 | 5 | CDKN1B |
| 1.97E-102 | 0.28439166 | 0.33 | 0.156 | 4.77E-98 | 5 | PHYH |
| 3.10E-102 | 0.37888164 | 0.759 | 0.542 | 7.52E-98 | 5 | IFI16 |
| 8.81E-102 | 0.46010487 | 0.86 | 0.74 | 2.14E-97 | 5 | EIF5 |
| 5.12E-99 | 0.30658989 | 0.483 | 0.26 | 1.24E-94 | 5 | WWP1 |
| 1.53E-98 | 0.27828407 | 0.363 | 0.177 | 3.71E-94 | 5 | RPS4Y1 |
| 2.49E-98 | 0.37311951 | 0.454 | 0.255 | 6.04E-94 | 5 | Z93930.2 |
| 7.64E-98 | 0.27221471 | 0.338 | 0.162 | 1.85E-93 | 5 | GIMAP7 |
| 4.74E-97 | 0.58613535 | 0.638 | 0.444 | 1.15E-92 | 5 | NABP1 |
| 5.30E-97 | 0.27180652 | 0.426 | 0.225 | 1.29E-92 | 5 | SPATS2L |
| 7.65E-96 | 0.4069826 | 0.693 | 0.483 | 1.86E-91 | 5 | CD163 |
| 2.50E-95 | 0.33623881 | 0.498 | 0.293 | 6.07E-91 | 5 | PLXND1 |
| 2.88E-95 | 0.44259859 | 0.985 | 0.961 | 6.97E-91 | 5 | ZFP36L1 |
| 3.00E-95 | 0.40550407 | 0.506 | 0.302 | 7.28E-91 | 5 | MAN1A1 |
| 3.43E-95 | 0.47911852 | 0.329 | 0.155 | 8.31E-91 | 5 | CCL8 |
| 5.06E-95 | 0.72125007 | 0.655 | 0.463 | 1.23E-90 | 5 | CITED2 |
| 7.38E-95 | 0.36795327 | 0.495 | 0.29 | 1.79E-90 | 5 | MCOLN1 |
| 7.61E-95 | 0.29425354 | 0.462 | 0.257 | 1.85E-90 | 5 | FCHSD2 |
| 2.52E-94 | 0.30283885 | 0.416 | 0.219 | 6.10E-90 | 5 | FUCA1 |
| 9.11E-94 | 0.60800767 | 0.869 | 0.775 | 2.21E-89 | 5 | DNAJB6 |
| 9.14E-93 | 0.4201804 | 0.553 | 0.356 | 2.21E-88 | 5 | TMEM176A |
| 6.72E-92 | 0.26657403 | 0.386 | 0.198 | 1.63E-87 | 5 | ARHGAP5 |
| 1.13E-91 | 0.56150613 | 0.912 | 0.85 | 2.73E-87 | 5 | CEBPD |

| | | | | | | |
|---|---|---|---|---|---|---|
| 2.89E-91 | 0.4466465 | 0.7 | 0.521 | 7.02E-87 | 5 | TMEM176B |
| 8.08E-91 | 0.4034286 | 0.931 | 0.861 | 1.96E-86 | 5 | MS4A6A |
| 8.89E-91 | 0.27508837 | 0.424 | 0.229 | 2.15E-86 | 5 | EPS8 |
| 2.47E-89 | 0.36019818 | 0.281 | 0.13 | 5.99E-85 | 5 | CDKN1C |
| 4.89E-86 | 0.39642993 | 0.253 | 0.112 | 1.19E-81 | 5 | RND3 |
| 7.57E-86 | 0.43922072 | 0.769 | 0.613 | 1.83E-81 | 5 | SFPQ |
| 1.77E-85 | 0.3730288 | 0.643 | 0.437 | 4.30E-81 | 5 | PLD3 |
| 5.35E-85 | 0.366958 | 0.468 | 0.277 | 1.30E-80 | 5 | PYGL |
| 9.70E-85 | 0.31466663 | 0.557 | 0.35 | 2.35E-80 | 5 | CYTH4 |
| 1.04E-84 | 0.26513064 | 0.43 | 0.242 | 2.51E-80 | 5 | TECR |
| 1.71E-84 | 0.33120394 | 0.269 | 0.124 | 4.13E-80 | 5 | ZNF503 |
| 2.05E-83 | 0.43875182 | 0.731 | 0.586 | 4.97E-79 | 5 | RBPJ |
| 4.07E-83 | 0.47518774 | 0.679 | 0.498 | 9.87E-79 | 5 | YME1L1 |
| 4.86E-83 | 0.38883606 | 0.985 | 0.967 | 1.18E-78 | 5 | HNRNPA2B1 |
| 7.61E-83 | 0.32543597 | 0.526 | 0.328 | 1.85E-78 | 5 | GYPC |
| 2.93E-82 | 0.27357405 | 0.399 | 0.217 | 7.09E-78 | 5 | SESN1 |
| 5.74E-82 | 0.28067141 | 0.257 | 0.118 | 1.39E-77 | 5 | CSH1 |
| 9.76E-82 | 0.25598256 | 0.377 | 0.2 | 2.37E-77 | 5 | CR1 |
| 1.25E-81 | 0.31987033 | 0.538 | 0.337 | 3.03E-77 | 5 | 1-Mar |
| 1.33E-81 | 0.32376986 | 0.594 | 0.393 | 3.22E-77 | 5 | ARHGAP18 |
| 6.04E-81 | 0.25561385 | 0.298 | 0.145 | 1.46E-76 | 5 | SLC7A8 |
| 1.22E-80 | 0.44580545 | 0.783 | 0.662 | 2.96E-76 | 5 | TSC22D3 |
| 2.37E-79 | 0.32600411 | 0.295 | 0.146 | 5.74E-75 | 5 | HHEX |
| 9.33E-79 | 0.27803267 | 0.556 | 0.341 | 2.26E-74 | 5 | A2M |
| 5.22E-78 | 0.27170275 | 0.462 | 0.274 | 1.27E-73 | 5 | MPHOSPH8 |
| 3.81E-77 | 0.25517273 | 0.277 | 0.134 | 9.24E-73 | 5 | NMRK1 |
| 5.16E-77 | 0.27304944 | 0.338 | 0.18 | 1.25E-72 | 5 | TLR7 |
| 1.40E-76 | 0.32335203 | 0.602 | 0.408 | 3.38E-72 | 5 | PRPF38B |
| 7.83E-76 | 0.34988957 | 0.356 | 0.196 | 1.90E-71 | 5 | CEMIP2 |
| 8.29E-76 | 0.4162913 | 0.765 | 0.608 | 2.01E-71 | 5 | PTGES3 |
| 1.44E-75 | 0.37120548 | 0.662 | 0.486 | 3.49E-71 | 5 | DDX24 |
| 2.08E-75 | 0.52040687 | 0.599 | 0.419 | 5.05E-71 | 5 | NRP2 |
| 6.34E-75 | 0.2611702 | 0.322 | 0.169 | 1.54E-70 | 5 | SCML1 |
| 6.85E-75 | 0.25416625 | 0.329 | 0.173 | 1.66E-70 | 5 | PIK3IP1 |
| 9.65E-75 | 0.28579875 | 1 | 1 | 2.34E-70 | 5 | MT-CYB |
| 1.31E-74 | 0.25715541 | 0.427 | 0.249 | 3.18E-70 | 5 | FCGR1B |
| 2.29E-74 | 0.37613973 | 0.637 | 0.46 | 5.55E-70 | 5 | FCGR1A |
| 7.19E-74 | 0.47485153 | 0.519 | 0.348 | 1.74E-69 | 5 | CREM |
| 1.31E-72 | 0.28208988 | 0.392 | 0.226 | 3.18E-68 | 5 | TCF4 |
| 3.56E-72 | 0.4764534 | 0.655 | 0.507 | 8.63E-68 | 5 | TRIB1 |
| 5.89E-72 | 0.38085865 | 0.472 | 0.297 | 1.43E-67 | 5 | ZC3HAV1 |
| 7.27E-72 | 0.29660926 | 0.6 | 0.409 | 1.76E-67 | 5 | SNX2 |
| 2.50E-71 | 0.49402217 | 0.537 | 0.363 | 6.06E-67 | 5 | BCAS2 |

| | | | | | | |
|---|---|---|---|---|---|---|
| 7.36E-70 | 0.54247758 | 0.771 | 0.67 | 1.78E-65 | 5 | NR4A2 |
| 1.09E-69 | 0.26377863 | 0.436 | 0.263 | 2.64E-65 | 5 | STOM |
| 1.83E-69 | 0.35970533 | 0.931 | 0.888 | 4.44E-65 | 5 | FCGRT |
| 3.24E-69 | 0.46040262 | 0.9 | 0.848 | 7.85E-65 | 5 | DDX3X |
| 4.45E-69 | 0.28622168 | 0.486 | 0.309 | 1.08E-64 | 5 | NASP |
| 5.42E-69 | 0.26030751 | 0.403 | 0.236 | 1.31E-64 | 5 | SERPING1 |
| 3.75E-68 | 0.28244725 | 0.443 | 0.27 | 9.08E-64 | 5 | GASK1B |
| 1.04E-67 | 0.2548767 | 0.309 | 0.164 | 2.51E-63 | 5 | Z99127.4 |
| 2.49E-66 | 0.51632076 | 0.721 | 0.591 | 6.03E-62 | 5 | EIF4A3 |
| 4.99E-66 | 0.2717853 | 0.574 | 0.382 | 1.21E-61 | 5 | NAIP |
| 5.07E-66 | 0.27972059 | 0.34 | 0.191 | 1.23E-61 | 5 | UGCG |
| 5.35E-66 | 0.38577094 | 0.623 | 0.457 | 1.30E-61 | 5 | AFF4 |
| 1.30E-65 | 0.26873192 | 1 | 1 | 3.16E-61 | 5 | MT-ATP6 |
| 2.15E-65 | 0.27021537 | 0.547 | 0.365 | 5.21E-61 | 5 | BMP2K |
| 5.61E-64 | 0.38262963 | 0.983 | 0.981 | 1.36E-59 | 5 | ZFP36 |
| 9.43E-63 | 0.40782474 | 0.69 | 0.534 | 2.29E-58 | 5 | AMD1 |
| 1.00E-61 | 0.29738845 | 0.511 | 0.341 | 2.42E-57 | 5 | CCR1 |
| 2.76E-61 | 0.25667006 | 0.347 | 0.198 | 6.69E-57 | 5 | BHLHE41 |
| 5.71E-61 | 0.32657137 | 0.591 | 0.418 | 1.38E-56 | 5 | PPP1R10 |
| 1.07E-60 | 0.3992494 | 0.936 | 0.885 | 2.58E-56 | 5 | AC020916.1 |
| 1.26E-60 | 0.30982594 | 0.814 | 0.712 | 3.07E-56 | 5 | RNASET2 |
| 1.41E-60 | 0.26316904 | 0.552 | 0.374 | 3.41E-56 | 5 | RB1 |
| 1.54E-60 | 0.38439135 | 0.464 | 0.312 | 3.73E-56 | 5 | PLEKHG2 |
| 1.55E-60 | 0.46888743 | 0.765 | 0.662 | 3.76E-56 | 5 | FCGR3A |
| 3.56E-60 | 0.34140689 | 0.636 | 0.471 | 8.63E-56 | 5 | TACC1 |
| 5.63E-60 | 0.27811415 | 0.462 | 0.297 | 1.36E-55 | 5 | LTC4S |
| 6.50E-59 | 0.25421355 | 0.5 | 0.329 | 1.58E-54 | 5 | NKTR |
| 7.33E-59 | 0.25164274 | 0.501 | 0.332 | 1.78E-54 | 5 | RRBP1 |
| 2.71E-58 | 0.29492344 | 0.676 | 0.521 | 6.56E-54 | 5 | CREG1 |
| 2.71E-58 | 0.25839105 | 0.403 | 0.249 | 6.57E-54 | 5 | SNHG12 |
| 2.00E-57 | 0.36882649 | 1 | 0.999 | 4.84E-53 | 5 | H3F3B |
| 1.78E-56 | 0.28304629 | 0.324 | 0.19 | 4.31E-52 | 5 | AC104695.4 |
| 2.30E-56 | 0.34328735 | 0.551 | 0.387 | 5.58E-52 | 5 | PNN |
| 3.26E-56 | 0.30866709 | 0.424 | 0.267 | 7.91E-52 | 5 | AC091271.1 |
| 3.30E-56 | 0.27283603 | 0.905 | 0.826 | 8.01E-52 | 5 | RBM39 |
| 4.33E-56 | 0.43971234 | 0.656 | 0.525 | 1.05E-51 | 5 | LINC-PINT |
| 2.39E-55 | 0.26041111 | 0.771 | 0.613 | 5.79E-51 | 5 | SON |
| 1.35E-53 | 0.4272687 | 0.759 | 0.615 | 3.27E-49 | 5 | ZFP36L2 |
| 4.13E-53 | 0.34362805 | 0.436 | 0.292 | 1.00E-48 | 5 | IRS2 |
| 5.56E-53 | 0.34041017 | 0.653 | 0.506 | 1.35E-48 | 5 | RNF149 |
| 1.26E-52 | 0.26260928 | 0.518 | 0.357 | 3.05E-48 | 5 | TNFRSF1A |
| 3.47E-52 | 0.2909745 | 0.614 | 0.461 | 8.42E-48 | 5 | BCLAF1 |
| 2.08E-51 | 0.38025417 | 0.629 | 0.497 | 5.04E-47 | 5 | ELF1 |

| | | | | | | |
|---|---|---|---|---|---|---|
| 2.32E-51 | 0.29251968 | 0.524 | 0.369 | 5.63E-47 | 5 | HEXA |
| 4.63E-51 | 0.2560445 | 0.647 | 0.497 | 1.12E-46 | 5 | CLTA |
| 1.67E-50 | 0.26668814 | 0.332 | 0.203 | 4.06E-46 | 5 | ING1 |
| 3.00E-50 | 0.31742117 | 0.475 | 0.326 | 7.27E-46 | 5 | IDI1 |
| 7.47E-49 | 0.28640544 | 0.546 | 0.395 | 1.81E-44 | 5 | HSPA9 |
| 2.69E-48 | 0.40879755 | 0.417 | 0.282 | 6.53E-44 | 5 | TENT5A |
| 4.86E-48 | 0.25119697 | 0.5 | 0.353 | 1.18E-43 | 5 | RBMS1 |
| 1.37E-47 | 0.25738845 | 0.545 | 0.389 | 3.32E-43 | 5 | ANKRD12 |
| 1.48E-47 | 0.28944108 | 0.538 | 0.387 | 3.59E-43 | 5 | BRD2 |
| 3.64E-46 | 0.2738674 | 0.616 | 0.465 | 8.83E-42 | 5 | RBM25 |
| 9.29E-46 | 0.29521466 | 0.897 | 0.829 | 2.25E-41 | 5 | SRSF3 |
| 4.33E-45 | 0.47907117 | 0.822 | 0.761 | 1.05E-40 | 5 | HSP90B1 |
| 3.10E-44 | 0.27410544 | 0.548 | 0.402 | 7.51E-40 | 5 | STAT3 |
| 9.12E-44 | 0.25872687 | 0.589 | 0.45 | 2.21E-39 | 5 | LAMP1 |
| 1.83E-43 | 0.38326715 | 0.751 | 0.659 | 4.43E-39 | 5 | NR4A1 |
| 3.24E-43 | 0.31900663 | 0.806 | 0.718 | 7.85E-39 | 5 | FUS |
| 4.37E-41 | 0.37379032 | 0.5 | 0.37 | 1.06E-36 | 5 | ARL4C |
| 4.42E-41 | 0.29021776 | 0.576 | 0.439 | 1.07E-36 | 5 | FAM133B |
| 2.97E-40 | 0.29585229 | 0.614 | 0.472 | 7.20E-36 | 5 | RAP2B |
| 5.57E-39 | 0.31316569 | 0.943 | 0.932 | 1.35E-34 | 5 | NFKBIZ |
| 2.51E-36 | 0.33229874 | 0.566 | 0.434 | 6.09E-32 | 5 | TLR4 |
| 8.53E-34 | 0.49179153 | 0.484 | 0.384 | 2.07E-29 | 5 | TFRC |
| 3.16E-31 | 0.33752721 | 0.439 | 0.331 | 7.65E-27 | 5 | TCOF1 |
| 6.73E-31 | 0.3297642 | 0.809 | 0.749 | 1.63E-26 | 5 | ATF3 |
| 2.43E-30 | 0.27717001 | 0.736 | 0.649 | 5.88E-26 | 5 | XBP1 |
| 3.99E-30 | 0.45535052 | 0.958 | 0.958 | 9.68E-26 | 5 | ID2 |
| 5.92E-26 | 0.32849539 | 0.335 | 0.242 | 1.43E-21 | 5 | CKS2 |
| 1.22E-23 | 0.3108438 | 0.78 | 0.72 | 2.97E-19 | 5 | MAP1LC3B |
| 4.21E-19 | 0.2548119 | 0.415 | 0.329 | 1.02E-14 | 5 | DYNC1H1 |
| 8.59E-17 | 0.31222574 | 0.356 | 0.282 | 2.08E-12 | 5 | ADM |
| 2.49E-16 | 0.48328775 | 0.606 | 0.572 | 6.04E-12 | 5 | HBEGF |
| 1.19E-13 | 0.28469791 | 0.626 | 0.585 | 2.89E-09 | 5 | SOCS3 |
| 6.24E-11 | 0.30101159 | 0.633 | 0.595 | 1.51E-06 | 5 | YBX3 |
| 9.75E-06 | 0.26009855 | 0.894 | 0.894 | 0.23648353 | 5 | GLUL |
| 0 | 3.55907458 | 0.939 | 0.797 | 0 | 6 | SPP1 |
| 0 | 3.39179712 | 0.849 | 0.305 | 0 | 6 | FN1 |
| 0 | 2.6327284 | 0.943 | 0.558 | 0 | 6 | FABP5 |
| 0 | 1.62910411 | 0.966 | 0.767 | 0 | 6 | CSTB |
| 0 | 1.53027279 | 0.997 | 0.741 | 0 | 6 | LGALS1 |
| 0 | 1.30747751 | 0.912 | 0.443 | 0 | 6 | PHLDA1 |
| 0 | 1.30143596 | 0.478 | 0.037 | 0 | 6 | LPL |
| 0 | 1.2854528 | 0.805 | 0.344 | 0 | 6 | GPNMB |
| 0 | 1.28289461 | 0.841 | 0.32 | 0 | 6 | CD9 |

| | | | | | | |
|---|---|---|---|---|---|---|
| 0 | 1.2760225 | 0.974 | 0.741 | 0 | 6 | MIF |
| 0 | 1.27328024 | 0.995 | 0.721 | 0 | 6 | S100A10 |
| 0 | 1.26984784 | 0.999 | 0.787 | 0 | 6 | VIM |
| 0 | 1.24599633 | 0.985 | 0.924 | 0 | 6 | CTSD |
| 0 | 1.21741908 | 0.663 | 0.185 | 0 | 6 | GCHFR |
| 0 | 1.21570275 | 0.969 | 0.639 | 0 | 6 | LGALS3 |
| 0 | 1.16970464 | 0.983 | 0.674 | 0 | 6 | ANXA2 |
| 0 | 1.14442242 | 0.746 | 0.195 | 0 | 6 | ALCAM |
| 0 | 1.1332844 | 0.57 | 0.191 | 0 | 6 | MARCO |
| 0 | 1.1259212 | 0.999 | 0.932 | 0 | 6 | S100A11 |
| 0 | 1.07367287 | 0.746 | 0.25 | 0 | 6 | TREM2 |
| 0 | 1.04407027 | 0.9 | 0.565 | 0 | 6 | ENO1 |
| 0 | 0.99913156 | 0.998 | 0.91 | 0 | 6 | SH3BGRL3 |
| 0 | 0.9622547 | 0.634 | 0.211 | 0 | 6 | FBP1 |
| 0 | 0.92206674 | 0.896 | 0.488 | 0 | 6 | CAPG |
| 0 | 0.91497598 | 0.998 | 0.961 | 0 | 6 | MYL6 |
| 0 | 0.86692728 | 1 | 0.999 | 0 | 6 | ACTB |
| 0 | 0.81731591 | 0.509 | 0.145 | 0 | 6 | CLEC5A |
| 0 | 0.75995492 | 0.403 | 0.086 | 0 | 6 | SDC2 |
| 0 | 0.7580553 | 0.471 | 0.085 | 0 | 6 | HTRA1 |
| 0 | 0.73426133 | 0.428 | 0.089 | 0 | 6 | CST6 |
| 0 | 0.6273834 | 0.451 | 0.113 | 0 | 6 | EMP1 |
| 0 | 0.59134655 | 0.37 | 0.07 | 0 | 6 | MMP19 |
| 0 | 0.57825775 | 0.477 | 0.124 | 0 | 6 | ST14 |
| 0 | 0.56889441 | 0.382 | 0.083 | 0 | 6 | SLC16A10 |
| 2.75E-304 | 1.65015734 | 0.85 | 0.499 | 6.67E-300 | 6 | IL1RN |
| 7.10E-302 | 1.0222105 | 0.997 | 0.755 | 1.72E-297 | 6 | S100A6 |
| 2.24E-301 | 0.8697301 | 0.833 | 0.426 | 5.43E-297 | 6 | PKM |
| 1.08E-300 | 0.70645868 | 0.999 | 0.976 | 2.62E-296 | 6 | SERF2 |
| 8.80E-297 | 0.91306593 | 0.999 | 0.842 | 2.13E-292 | 6 | IFI30 |
| 3.20E-295 | 0.53959481 | 0.427 | 0.114 | 7.77E-291 | 6 | NME1 |
| 5.50E-285 | 1.01342641 | 0.91 | 0.548 | 1.33E-280 | 6 | LMNA |
| 9.12E-284 | 0.74792918 | 0.977 | 0.861 | 2.21E-279 | 6 | ANXA5 |
| 1.16E-282 | 0.82288009 | 0.885 | 0.493 | 2.81E-278 | 6 | LIMS1 |
| 1.84E-278 | 0.78427315 | 1 | 0.982 | 4.45E-274 | 6 | PSAP |
| 1.97E-277 | 1.95839503 | 0.829 | 0.516 | 4.78E-273 | 6 | APOC1 |
| 1.12E-272 | 0.79406708 | 0.847 | 0.474 | 2.72E-268 | 6 | SEC61G |
| 5.02E-270 | 0.72259916 | 0.992 | 0.93 | 1.22E-265 | 6 | PFN1 |
| 2.77E-266 | 0.94973989 | 0.783 | 0.437 | 6.72E-262 | 6 | MGST3 |
| 1.16E-259 | 0.79138461 | 0.945 | 0.72 | 2.82E-255 | 6 | ATP6V1F |
| 3.02E-256 | 0.64630159 | 0.992 | 0.937 | 7.32E-252 | 6 | CFL1 |
| 5.98E-255 | 0.32048768 | 0.262 | 0.051 | 1.45E-250 | 6 | SEMA3C |
| 4.08E-254 | 0.71630465 | 0.983 | 0.925 | 9.89E-250 | 6 | YBX1 |

| | | | | | | |
|---|---|---|---|---|---|---|
| 1.86E-251 | 0.87126239 | 0.638 | 0.279 | 4.50E-247 | 6 | GSN |
| 1.57E-247 | 0.80596071 | 0.717 | 0.315 | 3.80E-243 | 6 | CYTOR |
| 3.19E-245 | 0.70323871 | 0.652 | 0.293 | 7.74E-241 | 6 | PPP1R14B |
| 7.50E-239 | 0.78572367 | 0.894 | 0.602 | 1.82E-234 | 6 | NDUFB2 |
| 8.18E-219 | 0.70047091 | 0.902 | 0.605 | 1.98E-214 | 6 | GSTO1 |
| 1.98E-217 | 0.70437018 | 0.988 | 0.946 | 4.80E-213 | 6 | ACTG1 |
| 3.33E-214 | 0.73349022 | 0.997 | 0.934 | 8.09E-210 | 6 | GAPDH |
| 5.24E-213 | 0.33426509 | 0.263 | 0.059 | 1.27E-208 | 6 | SERPINE1 |
| 8.94E-208 | 0.49746004 | 0.54 | 0.207 | 2.17E-203 | 6 | ACTN1 |
| 1.53E-207 | 0.81268328 | 0.76 | 0.402 | 3.70E-203 | 6 | SNX10 |
| 3.08E-206 | 0.61511177 | 0.701 | 0.355 | 7.46E-202 | 6 | VDAC1 |
| 4.17E-206 | 0.72180756 | 0.997 | 0.969 | 1.01E-201 | 6 | CD63 |
| 5.10E-205 | 0.8460216 | 0.888 | 0.577 | 1.24E-200 | 6 | RGCC |
| 3.33E-204 | 0.61819523 | 0.612 | 0.279 | 8.08E-200 | 6 | CCT5 |
| 4.37E-204 | 0.38568879 | 0.333 | 0.093 | 1.06E-199 | 6 | RMDN3 |
| 1.15E-203 | 0.59213796 | 0.593 | 0.252 | 2.78E-199 | 6 | GLIPR2 |
| 2.07E-202 | 0.81750524 | 0.874 | 0.618 | 5.01E-198 | 6 | DBI |
| 2.59E-200 | 0.76478287 | 0.896 | 0.637 | 6.28E-196 | 6 | LDHA |
| 3.15E-200 | 0.59157465 | 0.69 | 0.331 | 7.63E-196 | 6 | TUBA1C |
| 4.98E-200 | 0.74087925 | 0.904 | 0.666 | 1.21E-195 | 6 | PRDX1 |
| 8.31E-200 | 0.46993651 | 0.308 | 0.083 | 2.01E-195 | 6 | SLC39A8 |
| 5.04E-197 | 0.70846029 | 0.891 | 0.624 | 1.22E-192 | 6 | COX7A2 |
| 1.88E-196 | 0.64260369 | 0.958 | 0.812 | 4.55E-192 | 6 | NME2 |
| 2.31E-195 | 0.49027349 | 0.489 | 0.184 | 5.59E-191 | 6 | RAB13 |
| 4.16E-195 | 0.64804387 | 0.796 | 0.414 | 1.01E-190 | 6 | FLNA |
| 3.06E-193 | 0.49264179 | 0.384 | 0.121 | 7.43E-189 | 6 | PALM2-AKAP |
| 1.77E-192 | 0.57204397 | 0.986 | 0.91 | 4.29E-188 | 6 | PPIA |
| 7.31E-192 | 0.61616598 | 0.947 | 0.749 | 1.77E-187 | 6 | GNG5 |
| 1.68E-191 | 0.60075391 | 0.733 | 0.377 | 4.08E-187 | 6 | PPDPF |
| 8.16E-190 | 0.50713391 | 1 | 1 | 1.98E-185 | 6 | TMSB4X |
| 1.25E-185 | 0.63293224 | 0.853 | 0.552 | 3.02E-181 | 6 | POLR2L |
| 2.82E-184 | 0.73673443 | 0.803 | 0.494 | 6.83E-180 | 6 | MSR1 |
| 2.36E-180 | 0.65743767 | 0.751 | 0.435 | 5.73E-176 | 6 | TUBB |
| 2.88E-180 | 0.70541636 | 0.987 | 0.701 | 6.99E-176 | 6 | OLR1 |
| 1.31E-179 | 0.59384228 | 0.713 | 0.37 | 3.17E-175 | 6 | HMGA1 |
| 7.05E-179 | 0.64910626 | 0.812 | 0.524 | 1.71E-174 | 6 | ATP5MC3 |
| 4.08E-176 | 0.58642452 | 0.726 | 0.398 | 9.90E-172 | 6 | PDXK |
| 8.67E-175 | 0.50318024 | 0.493 | 0.195 | 2.10E-170 | 6 | FPR3 |
| 2.22E-174 | 0.66562728 | 0.955 | 0.626 | 5.39E-170 | 6 | C15orf48 |
| 4.44E-170 | 0.43201576 | 0.468 | 0.183 | 1.08E-165 | 6 | SLC43A3 |
| 7.16E-170 | 0.4631328 | 0.411 | 0.15 | 1.74E-165 | 6 | NCEH1 |
| 2.52E-165 | 0.57295757 | 0.852 | 0.574 | 6.12E-161 | 6 | NDUFS5 |
| 3.27E-165 | 0.63386996 | 0.878 | 0.653 | 7.93E-161 | 6 | PPIB |

| | | | | | | |
|---|---|---|---|---|---|---|
| 6.60E-164 | 0.51750169 | 0.992 | 0.96 | 1.60E-159 | 6 | LAPTM5 |
| 1.80E-161 | 0.58572964 | 0.956 | 0.821 | 4.37E-157 | 6 | RPSA |
| 3.40E-161 | 0.43617002 | 1 | 0.998 | 8.23E-157 | 6 | PTMA |
| 3.07E-160 | 0.61449904 | 0.84 | 0.507 | 7.44E-156 | 6 | TAGLN2 |
| 5.44E-158 | 0.35026801 | 0.377 | 0.132 | 1.32E-153 | 6 | MCRIP2 |
| 2.58E-157 | 0.33578554 | 0.25 | 0.067 | 6.26E-153 | 6 | CYP27A1 |
| 4.87E-157 | 0.44944486 | 0.53 | 0.236 | 1.18E-152 | 6 | COX20 |
| 5.31E-157 | 0.30466189 | 0.287 | 0.084 | 1.29E-152 | 6 | BOLA3 |
| 3.37E-156 | 0.51082714 | 0.619 | 0.317 | 8.18E-152 | 6 | BCAP31 |
| 4.19E-155 | 0.608744 | 0.487 | 0.205 | 1.02E-150 | 6 | ANPEP |
| 1.47E-151 | 0.33484817 | 0.314 | 0.101 | 3.58E-147 | 6 | SRM |
| 7.63E-150 | 0.41094864 | 0.347 | 0.12 | 1.85E-145 | 6 | MGST1 |
| 3.62E-148 | 0.43340796 | 0.384 | 0.142 | 8.78E-144 | 6 | IL4I1 |
| 4.18E-148 | 1.04460028 | 0.822 | 0.633 | 1.01E-143 | 6 | APOE |
| 7.55E-148 | 0.58405389 | 0.782 | 0.492 | 1.83E-143 | 6 | PGK1 |
| 2.03E-147 | 0.55442263 | 0.88 | 0.651 | 4.92E-143 | 6 | ATP5MD |
| 3.63E-147 | 0.63117075 | 0.905 | 0.635 | 8.79E-143 | 6 | RGS1 |
| 4.56E-147 | 0.54574166 | 0.933 | 0.783 | 1.11E-142 | 6 | ATP6V0B |
| 3.50E-146 | 0.45911341 | 0.551 | 0.263 | 8.49E-142 | 6 | TXNDC17 |
| 1.15E-145 | 0.49618852 | 0.704 | 0.382 | 2.78E-141 | 6 | PHPT1 |
| 2.22E-144 | 0.55321065 | 0.866 | 0.626 | 5.39E-140 | 6 | SEC61B |
| 7.83E-144 | 0.45451103 | 0.565 | 0.273 | 1.90E-139 | 6 | TIMM8B |
| 1.84E-143 | 0.43398522 | 0.451 | 0.191 | 4.47E-139 | 6 | NHP2 |
| 2.93E-142 | 0.44999902 | 0.585 | 0.285 | 7.10E-138 | 6 | CORO1C |
| 4.08E-142 | 0.54270691 | 0.824 | 0.515 | 9.90E-138 | 6 | TXN |
| 6.04E-142 | 0.50693071 | 0.723 | 0.417 | 1.47E-137 | 6 | NDUFS6 |
| 1.95E-141 | 0.45663233 | 0.478 | 0.214 | 4.74E-137 | 6 | C1QBP |
| 8.71E-139 | 0.46457063 | 0.694 | 0.382 | 2.11E-134 | 6 | ROMO1 |
| 2.23E-138 | 0.47050783 | 0.557 | 0.275 | 5.40E-134 | 6 | ATP5MC1 |
| 5.63E-138 | 0.42992355 | 0.48 | 0.214 | 1.37E-133 | 6 | PDCD5 |
| 2.60E-137 | 0.27947385 | 0.263 | 0.079 | 6.30E-133 | 6 | PPARG |
| 3.21E-137 | 0.33428581 | 0.317 | 0.108 | 7.79E-133 | 6 | MYO1E |
| 3.42E-137 | 0.49632506 | 0.62 | 0.327 | 8.29E-133 | 6 | SCARB2 |
| 3.49E-137 | 0.50000469 | 0.614 | 0.326 | 8.46E-133 | 6 | SLIRP |
| 2.14E-133 | 0.68113393 | 0.663 | 0.392 | 5.19E-129 | 6 | LIPA |
| 1.66E-131 | 0.5612684 | 0.813 | 0.57 | 4.03E-127 | 6 | ATP6V1G1 |
| 4.45E-131 | 0.43691634 | 0.536 | 0.262 | 1.08E-126 | 6 | TOMM5 |
| 1.07E-130 | 0.48399208 | 0.728 | 0.403 | 2.58E-126 | 6 | LSP1 |
| 1.82E-130 | 0.50779023 | 0.848 | 0.609 | 4.42E-126 | 6 | COX8A |
| 7.17E-129 | 0.48866213 | 0.907 | 0.682 | 1.74E-124 | 6 | NDUFA13 |
| 7.42E-128 | 0.4470796 | 0.998 | 0.978 | 1.80E-123 | 6 | ATP5F1E |
| 2.27E-127 | 0.47060775 | 1 | 1 | 5.49E-123 | 6 | FTH1 |
| 2.24E-126 | 1.47350948 | 0.594 | 0.335 | 5.43E-122 | 6 | CCL20 |

| | | | | | | |
|---|---|---|---|---|---|---|
| 2.97E-126 | 0.64182966 | 0.891 | 0.681 | 7.20E-122 | 6 | DUSP2 |
| 4.62E-126 | 0.42604085 | 0.574 | 0.294 | 1.12E-121 | 6 | SNRPE |
| 2.17E-125 | 0.50449176 | 0.888 | 0.654 | 5.27E-121 | 6 | COX7B |
| 4.04E-124 | 0.50268129 | 0.772 | 0.504 | 9.78E-120 | 6 | ATP5PF |
| 8.17E-124 | 0.60754741 | 0.834 | 0.63 | 1.98E-119 | 6 | DYNLL1 |
| 1.40E-123 | 0.46532782 | 0.92 | 0.758 | 3.40E-119 | 6 | CHCHD2 |
| 2.64E-123 | 0.54635429 | 0.938 | 0.826 | 6.41E-119 | 6 | CALM1 |
| 2.84E-123 | 0.51477689 | 0.878 | 0.686 | 6.87E-119 | 6 | COX6C |
| 3.34E-123 | 0.56239608 | 0.889 | 0.693 | 8.11E-119 | 6 | CALR |
| 4.90E-123 | 0.62273412 | 0.965 | 0.794 | 1.19E-118 | 6 | EMP3 |
| 1.39E-121 | 0.50056455 | 0.667 | 0.401 | 3.36E-117 | 6 | DAD1 |
| 1.45E-121 | 0.46326635 | 0.584 | 0.317 | 3.51E-117 | 6 | MYDGF |
| 4.36E-121 | 0.48610926 | 0.934 | 0.757 | 1.06E-116 | 6 | CD81 |
| 1.38E-120 | 0.42646951 | 0.528 | 0.267 | 3.34E-116 | 6 | TIMM13 |
| 2.97E-120 | 0.51425558 | 0.818 | 0.58 | 7.21E-116 | 6 | TPI1 |
| 3.07E-120 | 0.43707718 | 0.62 | 0.331 | 7.45E-116 | 6 | PEA15 |
| 4.01E-120 | 0.39840579 | 0.489 | 0.229 | 9.72E-116 | 6 | RALA |
| 4.40E-120 | 0.49975961 | 0.783 | 0.518 | 1.07E-115 | 6 | ATOX1 |
| 1.88E-119 | 0.43157753 | 0.727 | 0.387 | 4.57E-115 | 6 | AQP9 |
| 1.27E-118 | 0.53109878 | 0.811 | 0.561 | 3.07E-114 | 6 | RPS27L |
| 3.54E-118 | 0.54422383 | 0.985 | 0.765 | 8.58E-114 | 6 | CD44 |
| 5.00E-118 | 0.49411318 | 0.839 | 0.623 | 1.21E-113 | 6 | AP2S1 |
| 5.96E-118 | 0.40068791 | 0.608 | 0.306 | 1.44E-113 | 6 | SMIM25 |
| 6.95E-118 | 0.65348386 | 0.968 | 0.857 | 1.68E-113 | 6 | PLIN2 |
| 6.54E-117 | 0.42535774 | 0.566 | 0.3 | 1.59E-112 | 6 | MICOS13 |
| 1.48E-116 | 0.37564089 | 0.435 | 0.197 | 3.59E-112 | 6 | ANXA4 |
| 2.34E-116 | 0.46838498 | 0.878 | 0.671 | 5.67E-112 | 6 | SSR4 |
| 8.99E-116 | 0.33004068 | 0.341 | 0.133 | 2.18E-111 | 6 | CD82 |
| 1.69E-115 | 0.47599908 | 0.635 | 0.355 | 4.10E-111 | 6 | FNDC3B |
| 1.40E-114 | 0.45445002 | 0.9 | 0.706 | 3.39E-110 | 6 | UBL5 |
| 3.24E-114 | 0.47754063 | 0.863 | 0.648 | 7.87E-110 | 6 | POMP |
| 2.59E-113 | 0.27773319 | 0.31 | 0.115 | 6.29E-109 | 6 | PAPSS1 |
| 4.01E-113 | 0.53398745 | 0.889 | 0.734 | 9.72E-109 | 6 | ATP5ME |
| 6.00E-113 | 0.45579843 | 0.636 | 0.354 | 1.46E-108 | 6 | SMAD7 |
| 1.19E-112 | 0.41306665 | 0.982 | 0.924 | 2.89E-108 | 6 | RPLP0 |
| 1.69E-112 | 0.2575761 | 0.261 | 0.088 | 4.09E-108 | 6 | MAP4K3-DT |
| 3.69E-111 | 0.28796531 | 0.262 | 0.089 | 8.95E-107 | 6 | MRAS |
| 6.24E-110 | 0.46142554 | 0.612 | 0.349 | 1.51E-105 | 6 | EIF5A |
| 3.61E-109 | 0.46319227 | 0.909 | 0.731 | 8.74E-105 | 6 | EIF4G2 |
| 6.07E-109 | 0.37967657 | 0.485 | 0.241 | 1.47E-104 | 6 | NDUFAB1 |
| 6.18E-109 | 0.39334656 | 0.545 | 0.289 | 1.50E-104 | 6 | ACADVL |
| 2.35E-108 | 0.46558711 | 0.914 | 0.748 | 5.70E-104 | 6 | ELOB |
| 1.48E-107 | 0.34773826 | 0.486 | 0.229 | 3.59E-103 | 6 | PLP2 |

| | | | | | | |
|---|---|---|---|---|---|---|
| 1.54E-107 | 0.39168603 | 0.606 | 0.338 | 3.72E-103 | 6 | PSMD8 |
| 4.68E-107 | 0.39725419 | 0.373 | 0.161 | 1.13E-102 | 6 | ARID5B |
| 1.36E-106 | 1.33483322 | 0.331 | 0.139 | 3.29E-102 | 6 | APOC2 |
| 1.47E-106 | 0.43308332 | 0.784 | 0.484 | 3.57E-102 | 6 | SLC11A1 |
| 1.56E-106 | 0.44638079 | 0.789 | 0.548 | 3.77E-102 | 6 | MICOS10 |
| 1.99E-106 | 0.41144577 | 0.575 | 0.319 | 4.82E-102 | 6 | MRPL52 |
| 5.35E-106 | 0.70161445 | 0.35 | 0.143 | 1.30E-101 | 6 | AREG |
| 7.75E-106 | 0.47811381 | 0.475 | 0.228 | 1.88E-101 | 6 | SDS |
| 7.88E-106 | 0.45312501 | 0.847 | 0.636 | 1.91E-101 | 6 | ATP5MF |
| 2.21E-105 | 0.43087892 | 0.946 | 0.749 | 5.36E-101 | 6 | TSPO |
| 6.01E-105 | 0.30552458 | 0.33 | 0.133 | 1.46E-100 | 6 | GNA12 |
| 1.01E-104 | 0.44699738 | 0.947 | 0.824 | 2.45E-100 | 6 | CLIC1 |
| 2.18E-104 | 0.45524942 | 0.64 | 0.384 | 5.28E-100 | 6 | ATP6V0D1 |
| 1.03E-103 | 0.40479837 | 0.538 | 0.272 | 2.49E-99 | 6 | MIR155HG |
| 1.72E-103 | 0.42042047 | 0.628 | 0.37 | 4.17E-99 | 6 | HM13 |
| 3.41E-103 | 0.5422602 | 0.508 | 0.274 | 8.26E-99 | 6 | ME2 |
| 3.43E-103 | 0.42756336 | 0.896 | 0.74 | 8.31E-99 | 6 | UQCR11 |
| 4.55E-103 | 0.55019279 | 1 | 1 | 1.10E-98 | 6 | FTL |
| 6.97E-103 | 0.70054059 | 0.883 | 0.607 | 1.69E-98 | 6 | EREG |
| 1.21E-102 | 0.34830361 | 0.473 | 0.225 | 2.94E-98 | 6 | RASGRP3 |
| 1.63E-102 | 0.34975474 | 0.427 | 0.203 | 3.95E-98 | 6 | PHB |
| 3.33E-102 | 0.54724244 | 0.665 | 0.407 | 8.07E-98 | 6 | IFI6 |
| 4.17E-102 | 0.3116124 | 0.314 | 0.126 | 1.01E-97 | 6 | FAM20C |
| 4.79E-102 | 0.39344376 | 0.585 | 0.33 | 1.16E-97 | 6 | PSMB6 |
| 1.92E-101 | 0.39602634 | 0.642 | 0.377 | 4.67E-97 | 6 | TMED10 |
| 6.66E-101 | 0.3425695 | 0.507 | 0.26 | 1.61E-96 | 6 | GTF3C6 |
| 1.60E-100 | 0.40635826 | 0.699 | 0.43 | 3.87E-96 | 6 | CALM3 |
| 2.83E-99 | 0.38075945 | 0.64 | 0.373 | 6.85E-95 | 6 | ANAPC11 |
| 3.09E-99 | 0.40035453 | 0.642 | 0.384 | 7.48E-95 | 6 | SSBP1 |
| 3.52E-99 | 0.29646094 | 0.384 | 0.169 | 8.52E-95 | 6 | CYSTM1 |
| 6.77E-99 | 0.39392664 | 0.523 | 0.281 | 1.64E-94 | 6 | ENG |
| 1.14E-98 | 0.35731541 | 0.444 | 0.219 | 2.77E-94 | 6 | RANBP1 |
| 1.17E-98 | 0.45015096 | 0.786 | 0.539 | 2.84E-94 | 6 | PLXDC2 |
| 3.11E-98 | 0.38133025 | 0.998 | 0.973 | 7.53E-94 | 6 | RPL8 |
| 6.02E-98 | 0.42379656 | 0.675 | 0.426 | 1.46E-93 | 6 | SPCS2 |
| 8.69E-97 | 0.36666886 | 0.997 | 0.983 | 2.11E-92 | 6 | RPL35 |
| 1.36E-96 | 0.4882019 | 0.985 | 0.946 | 3.31E-92 | 6 | CTSB |
| 3.01E-96 | 0.41734584 | 0.722 | 0.468 | 7.29E-92 | 6 | APRT |
| 8.02E-96 | 0.51387334 | 0.781 | 0.564 | 1.94E-91 | 6 | NOP10 |
| 2.02E-95 | 0.35533649 | 0.496 | 0.251 | 4.90E-91 | 6 | SLC6A6 |
| 3.32E-95 | 0.365191 | 0.518 | 0.282 | 8.05E-91 | 6 | KDELR2 |
| 3.97E-95 | 0.3775006 | 0.945 | 0.818 | 9.63E-91 | 6 | RHOA |
| 6.47E-95 | 0.33101327 | 0.4 | 0.188 | 1.57E-90 | 6 | C1orf122 |

| | | | | | | |
|---|---|---|---|---|---|---|
| 7.88E-95 | 0.52424885 | 0.934 | 0.712 | 1.91E-90 | 6 | GPR183 |
| 1.39E-94 | 0.40250796 | 0.931 | 0.85 | 3.36E-90 | 6 | ARPC2 |
| 3.68E-94 | 0.27072946 | 0.332 | 0.14 | 8.93E-90 | 6 | ORMDL2 |
| 4.53E-94 | 0.48437927 | 0.507 | 0.269 | 1.10E-89 | 6 | OASL |
| 1.05E-93 | 0.39470262 | 0.786 | 0.539 | 2.54E-89 | 6 | GUK1 |
| 1.67E-93 | 0.40164926 | 0.654 | 0.401 | 4.06E-89 | 6 | COX5A |
| 1.69E-93 | 0.40523178 | 0.563 | 0.326 | 4.10E-89 | 6 | SNRPF |
| 2.54E-93 | 0.44323 | 0.316 | 0.129 | 6.16E-89 | 6 | HAMP |
| 3.48E-93 | 0.38059735 | 0.706 | 0.448 | 8.44E-89 | 6 | ARPC4 |
| 5.37E-93 | 0.25298687 | 0.285 | 0.109 | 1.30E-88 | 6 | SDC4 |
| 8.24E-93 | 0.38618809 | 0.694 | 0.432 | 2.00E-88 | 6 | PSMA6 |
| 1.53E-92 | 0.41295881 | 0.884 | 0.715 | 3.71E-88 | 6 | OST4 |
| 3.29E-92 | 0.3840818 | 0.474 | 0.248 | 7.98E-88 | 6 | DOCK10 |
| 3.70E-92 | 0.47169575 | 0.862 | 0.719 | 8.97E-88 | 6 | NDUFA4 |
| 3.80E-92 | 0.37388134 | 0.496 | 0.262 | 9.21E-88 | 6 | SDF2L1 |
| 1.65E-91 | 0.38539627 | 0.561 | 0.322 | 4.00E-87 | 6 | OSTC |
| 6.85E-91 | 0.38969801 | 0.67 | 0.413 | 1.66E-86 | 6 | LSM7 |
| 9.37E-91 | 0.41331156 | 0.831 | 0.624 | 2.27E-86 | 6 | COX6A1 |
| 1.63E-90 | 0.38241121 | 0.558 | 0.319 | 3.96E-86 | 6 | HNRNPAB |
| 2.36E-90 | 0.31091123 | 0.399 | 0.19 | 5.73E-86 | 6 | FUOM |
| 1.01E-89 | 0.34495232 | 0.404 | 0.191 | 2.44E-85 | 6 | ACP5 |
| 4.09E-89 | 0.40392675 | 0.865 | 0.675 | 9.91E-85 | 6 | NDUFA1 |
| 7.74E-89 | 0.31505342 | 0.395 | 0.187 | 1.88E-84 | 6 | GPI |
| 9.86E-89 | 0.30301966 | 0.409 | 0.198 | 2.39E-84 | 6 | TMEM208 |
| 1.37E-88 | 0.41599207 | 0.615 | 0.379 | 3.32E-84 | 6 | P4HB |
| 1.77E-88 | 0.29397567 | 0.426 | 0.209 | 4.30E-84 | 6 | EIF6 |
| 4.69E-88 | 0.45021705 | 0.824 | 0.662 | 1.14E-83 | 6 | PSMA7 |
| 6.79E-88 | 0.33015184 | 0.469 | 0.242 | 1.65E-83 | 6 | SQOR |
| 1.95E-87 | 0.3191762 | 0.435 | 0.218 | 4.74E-83 | 6 | MFSD12 |
| 2.17E-87 | 0.37467964 | 0.603 | 0.355 | 5.26E-83 | 6 | MPP1 |
| 3.73E-87 | 0.32273479 | 0.377 | 0.178 | 9.05E-83 | 6 | RHOC |
| 8.09E-87 | 0.33583934 | 0.562 | 0.317 | 1.96E-82 | 6 | COX16 |
| 8.90E-87 | 0.42949263 | 0.861 | 0.668 | 2.16E-82 | 6 | TPM4 |
| 1.08E-86 | 0.42611915 | 0.599 | 0.368 | 2.63E-82 | 6 | ELOC |
| 8.14E-86 | 0.42175342 | 0.917 | 0.784 | 1.97E-81 | 6 | NPM1 |
| 2.13E-85 | 0.44422096 | 0.727 | 0.493 | 5.17E-81 | 6 | CCDC88A |
| 3.80E-85 | 0.38941911 | 0.735 | 0.499 | 9.20E-81 | 6 | PARK7 |
| 4.38E-85 | 0.35701463 | 0.732 | 0.481 | 1.06E-80 | 6 | SEM1 |
| 5.00E-85 | 0.3401088 | 0.569 | 0.313 | 1.21E-80 | 6 | ANKRD28 |
| 5.85E-85 | 0.3262193 | 0.998 | 0.975 | 1.42E-80 | 6 | OAZ1 |
| 1.23E-84 | 0.37779789 | 0.985 | 0.924 | 2.97E-80 | 6 | RPS17 |
| 2.50E-84 | 0.35221396 | 0.469 | 0.248 | 6.05E-80 | 6 | LHFPL2 |
| 3.05E-84 | 0.3900965 | 0.962 | 0.812 | 7.40E-80 | 6 | TYMP |

| | | | | | | |
|---|---|---|---|---|---|---|
| 6.86E-84 | 0.32114116 | 0.424 | 0.213 | 1.66E-79 | 6 | CALU |
| 1.45E-83 | 0.33940066 | 0.996 | 0.973 | 3.51E-79 | 6 | NPC2 |
| 4.50E-83 | 0.3679673 | 0.764 | 0.538 | 1.09E-78 | 6 | NEDD8 |
| 8.30E-83 | 0.37992379 | 0.66 | 0.407 | 2.01E-78 | 6 | JPT1 |
| 1.43E-82 | 0.338485 | 0.419 | 0.211 | 3.46E-78 | 6 | ATF5 |
| 1.81E-82 | 0.26678385 | 0.351 | 0.158 | 4.39E-78 | 6 | PDLIM7 |
| 1.94E-82 | 0.26698517 | 0.352 | 0.162 | 4.71E-78 | 6 | PSMD14 |
| 2.34E-82 | 0.31914014 | 0.428 | 0.218 | 5.68E-78 | 6 | MANF |
| 3.35E-82 | 0.33626107 | 0.603 | 0.36 | 8.11E-78 | 6 | RHEB |
| 5.20E-82 | 0.30325911 | 0.415 | 0.209 | 1.26E-77 | 6 | SLC25A24 |
| 7.53E-82 | 0.40262347 | 0.736 | 0.506 | 1.83E-77 | 6 | SSR3 |
| 3.51E-81 | 0.49644163 | 0.991 | 0.778 | 8.51E-77 | 6 | S100A4 |
| 5.43E-81 | 0.35346746 | 0.726 | 0.467 | 1.32E-76 | 6 | CIB1 |
| 7.06E-81 | 0.35876832 | 0.643 | 0.408 | 1.71E-76 | 6 | NDUFB8 |
| 8.54E-81 | 0.32726212 | 0.558 | 0.323 | 2.07E-76 | 6 | WDR1 |
| 1.04E-80 | 0.29084635 | 0.432 | 0.221 | 2.51E-76 | 6 | PGAM1 |
| 3.69E-80 | 0.29971016 | 0.372 | 0.177 | 8.94E-76 | 6 | ABL2 |
| 5.39E-80 | 0.60206902 | 0.885 | 0.656 | 1.31E-75 | 6 | TIMP1 |
| 1.80E-79 | 0.37849427 | 0.502 | 0.279 | 4.36E-75 | 6 | FNIP2 |
| 3.62E-79 | 0.321584 | 0.538 | 0.305 | 8.79E-75 | 6 | NDUFB3 |
| 5.78E-79 | 0.35743561 | 0.947 | 0.852 | 1.40E-74 | 6 | GPX4 |
| 7.70E-79 | 0.35925512 | 0.71 | 0.47 | 1.87E-74 | 6 | RBX1 |
| 1.17E-78 | 0.26839819 | 0.386 | 0.187 | 2.83E-74 | 6 | UQCC2 |
| 4.20E-78 | 0.36761152 | 0.797 | 0.57 | 1.02E-73 | 6 | CAP1 |
| 5.29E-78 | 0.2837314 | 0.344 | 0.16 | 1.28E-73 | 6 | GPRIN3 |
| 7.25E-78 | 0.44179443 | 0.671 | 0.425 | 1.76E-73 | 6 | NR4A3 |
| 9.77E-78 | 0.33411093 | 0.593 | 0.359 | 2.37E-73 | 6 | CIAO2B |
| 1.30E-77 | 0.34801005 | 0.683 | 0.445 | 3.15E-73 | 6 | RBM8A |
| 1.47E-76 | 0.27092025 | 0.319 | 0.143 | 3.57E-72 | 6 | MYC |
| 1.66E-76 | 0.40972062 | 0.799 | 0.589 | 4.03E-72 | 6 | MSN |
| 2.04E-76 | 0.36861469 | 0.57 | 0.345 | 4.95E-72 | 6 | YWHAQ |
| 2.24E-76 | 0.29881328 | 0.455 | 0.242 | 5.43E-72 | 6 | NDUFA7 |
| 3.51E-76 | 0.31926817 | 0.608 | 0.372 | 8.50E-72 | 6 | PSMA2 |
| 5.06E-76 | 0.35641868 | 0.648 | 0.412 | 1.23E-71 | 6 | COX17 |
| 8.56E-76 | 0.37654135 | 0.612 | 0.387 | 2.08E-71 | 6 | PDIA6 |
| 8.66E-76 | 0.36588441 | 0.88 | 0.716 | 2.10E-71 | 6 | COX5B |
| 1.00E-75 | 0.30467464 | 0.393 | 0.202 | 2.43E-71 | 6 | SEC11C |
| 1.66E-75 | 0.27860068 | 0.297 | 0.128 | 4.02E-71 | 6 | MIR4435-2H[illegible] |
| 1.84E-75 | 0.27478026 | 0.323 | 0.148 | 4.46E-71 | 6 | MGLL |
| 3.42E-75 | 0.35704148 | 0.645 | 0.421 | 8.30E-71 | 6 | ATP5F1B |
| 4.11E-75 | 0.34580873 | 0.709 | 0.474 | 9.97E-71 | 6 | PSMB1 |
| 1.44E-74 | 0.39408188 | 0.617 | 0.385 | 3.48E-70 | 6 | ELL2 |
| 5.22E-74 | 0.25277967 | 0.266 | 0.111 | 1.27E-69 | 6 | HS3ST1 |

| | | | | | | |
|---|---|---|---|---|---|---|
| 8.26E-74 | 0.35940211 | 0.619 | 0.397 | 2.00E-69 | 6 | RAN |
| 6.72E-73 | 0.36624372 | 0.739 | 0.526 | 1.63E-68 | 6 | UQCRQ |
| 9.56E-73 | 0.34974916 | 0.866 | 0.674 | 2.32E-68 | 6 | H2AFY |
| 1.15E-72 | 0.62508956 | 0.784 | 0.634 | 2.79E-68 | 6 | CTSL |
| 4.04E-72 | 0.29297995 | 0.457 | 0.248 | 9.79E-68 | 6 | RAPGEF1 |
| 7.61E-72 | 0.31237828 | 0.442 | 0.242 | 1.85E-67 | 6 | YWHAG |
| 1.12E-71 | 0.31697383 | 0.63 | 0.389 | 2.72E-67 | 6 | CTSA |
| 5.99E-71 | 0.30212874 | 0.463 | 0.26 | 1.45E-66 | 6 | RPS19BP1 |
| 9.99E-71 | 0.29367142 | 0.999 | 0.991 | 2.42E-66 | 6 | RPS19 |
| 1.60E-70 | 0.29366372 | 0.502 | 0.288 | 3.87E-66 | 6 | TCIRG1 |
| 1.92E-70 | 0.25983084 | 0.421 | 0.221 | 4.66E-66 | 6 | LSM5 |
| 1.17E-69 | 0.34347681 | 0.758 | 0.533 | 2.85E-65 | 6 | KRTCAP2 |
| 3.03E-69 | 0.31903391 | 0.961 | 0.892 | 7.35E-65 | 6 | ATP6V0C |
| 1.23E-68 | 0.35095161 | 0.794 | 0.601 | 2.97E-64 | 6 | RBM3 |
| 1.71E-68 | 0.42092963 | 0.646 | 0.415 | 4.14E-64 | 6 | LTA4H |
| 1.97E-68 | 0.27437528 | 1 | 0.995 | 4.77E-64 | 6 | TYROBP |
| 2.06E-68 | 0.28347053 | 0.513 | 0.299 | 4.99E-64 | 6 | LSM3 |
| 2.18E-68 | 0.26044242 | 0.327 | 0.158 | 5.27E-64 | 6 | ABHD2 |
| 4.27E-68 | 0.29915194 | 0.435 | 0.241 | 1.03E-63 | 6 | SNRPD1 |
| 4.48E-67 | 0.28949676 | 0.543 | 0.323 | 1.09E-62 | 6 | TRAPPC1 |
| 6.92E-67 | 0.35675975 | 0.847 | 0.675 | 1.68E-62 | 6 | UQCRH |
| 1.09E-66 | 0.27441156 | 0.341 | 0.171 | 2.63E-62 | 6 | CD151 |
| 1.73E-66 | 0.33573647 | 0.811 | 0.62 | 4.20E-62 | 6 | TMEM258 |
| 3.51E-66 | 0.34376613 | 0.748 | 0.532 | 8.51E-62 | 6 | PPT1 |
| 3.71E-66 | 0.35575288 | 0.729 | 0.531 | 8.99E-62 | 6 | TOMM6 |
| 8.19E-66 | 0.29658951 | 0.488 | 0.284 | 1.98E-61 | 6 | NDUFB9 |
| 1.07E-65 | 0.36151215 | 0.836 | 0.654 | 2.60E-61 | 6 | PDIA3 |
| 1.25E-65 | 0.33221273 | 0.945 | 0.812 | 3.02E-61 | 6 | LCP1 |
| 2.51E-65 | 0.33645905 | 0.824 | 0.64 | 6.09E-61 | 6 | CANX |
| 2.91E-65 | 0.2996598 | 0.525 | 0.316 | 7.06E-61 | 6 | PA2G4 |
| 4.25E-65 | 0.26390134 | 0.384 | 0.197 | 1.03E-60 | 6 | CD109 |
| 2.01E-64 | 0.28035345 | 0.529 | 0.315 | 4.88E-60 | 6 | NDUFA6 |
| 3.40E-64 | 0.31272311 | 0.602 | 0.388 | 8.24E-60 | 6 | NDUFB7 |
| 3.48E-64 | 0.26221314 | 0.431 | 0.238 | 8.43E-60 | 6 | UQCRC1 |
| 5.17E-64 | 0.31992866 | 0.629 | 0.41 | 1.25E-59 | 6 | TLN1 |
| 6.86E-64 | 0.3326829 | 0.963 | 0.896 | 1.66E-59 | 6 | RPS5 |
| 1.67E-63 | 0.29684315 | 0.517 | 0.308 | 4.06E-59 | 6 | IRAK1 |
| 2.87E-63 | 0.28391736 | 0.526 | 0.318 | 6.96E-59 | 6 | GTF3A |
| 5.59E-63 | 0.31246019 | 0.758 | 0.548 | 1.36E-58 | 6 | YWHAE |
| 6.42E-63 | 0.32252771 | 0.874 | 0.711 | 1.56E-58 | 6 | MYL12B |
| 6.45E-63 | 0.27933998 | 0.404 | 0.22 | 1.56E-58 | 6 | B3GNT2 |
| 9.13E-63 | 0.29582977 | 0.505 | 0.304 | 2.21E-58 | 6 | MRPL20 |
| 5.69E-62 | 0.74618053 | 0.695 | 0.553 | 1.38E-57 | 6 | TGFBI |

| | | | | | | |
|---|---|---|---|---|---|---|
| 6.21E-62 | 0.28710219 | 0.547 | 0.34 | 1.51E-57 | 6 | FIS1 |
| 9.95E-62 | 0.26037323 | 0.528 | 0.318 | 2.41E-57 | 6 | GHITM |
| 1.73E-61 | 0.25781326 | 0.398 | 0.217 | 4.18E-57 | 6 | VKORC1 |
| 1.88E-61 | 0.26995024 | 0.531 | 0.321 | 4.56E-57 | 6 | BANF1 |
| 2.15E-61 | 0.26258637 | 0.424 | 0.238 | 5.21E-57 | 6 | PSMA5 |
| 3.08E-61 | 0.34841943 | 0.95 | 0.876 | 7.46E-57 | 6 | CTSZ |
| 3.83E-61 | 0.26530597 | 0.462 | 0.265 | 9.29E-57 | 6 | GADD45GIP1 |
| 5.17E-61 | 0.29553584 | 0.364 | 0.193 | 1.25E-56 | 6 | STK38L |
| 1.38E-60 | 0.33474937 | 0.753 | 0.491 | 3.35E-56 | 6 | CD52 |
| 2.62E-60 | 0.28787888 | 0.612 | 0.393 | 6.36E-56 | 6 | LGALS9 |
| 2.93E-60 | 0.29178425 | 0.733 | 0.486 | 7.10E-56 | 6 | FAM107B |
| 3.20E-60 | 0.30505889 | 0.576 | 0.37 | 7.77E-56 | 6 | SNRPG |
| 3.49E-60 | 0.30233625 | 0.558 | 0.354 | 8.47E-56 | 6 | RABAC1 |
| 6.60E-60 | 0.32094797 | 0.826 | 0.643 | 1.60E-55 | 6 | ACTR3 |
| 1.74E-59 | 0.28322288 | 0.535 | 0.336 | 4.21E-55 | 6 | AURKAIP1 |
| 3.34E-59 | 0.33549176 | 0.957 | 0.885 | 8.11E-55 | 6 | CD68 |
| 4.01E-59 | 0.3037513 | 0.673 | 0.462 | 9.72E-55 | 6 | SF3B5 |
| 1.07E-58 | 0.31484244 | 0.897 | 0.786 | 2.59E-54 | 6 | VAMP8 |
| 2.02E-58 | 0.59925692 | 0.715 | 0.521 | 4.91E-54 | 6 | ISG15 |
| 5.89E-58 | 0.37670575 | 0.71 | 0.508 | 1.43E-53 | 6 | ITGB1 |
| 6.37E-58 | 0.25219293 | 0.327 | 0.168 | 1.54E-53 | 6 | MYOF |
| 1.38E-57 | 0.26830678 | 0.392 | 0.214 | 3.34E-53 | 6 | DUSP10 |
| 1.63E-57 | 0.29516352 | 0.595 | 0.388 | 3.95E-53 | 6 | SNU13 |
| 4.89E-57 | 0.33399432 | 0.799 | 0.61 | 1.19E-52 | 6 | RASGEF1B |
| 1.02E-56 | 0.55432005 | 0.747 | 0.518 | 2.48E-52 | 6 | CRIP1 |
| 2.36E-56 | 0.34288019 | 0.813 | 0.661 | 5.72E-52 | 6 | UQCR10 |
| 2.85E-56 | 0.25516675 | 0.45 | 0.265 | 6.90E-52 | 6 | TWF2 |
| 5.02E-56 | 0.34594986 | 0.945 | 0.871 | 1.22E-51 | 6 | GRN |
| 5.40E-56 | 0.30173395 | 0.723 | 0.493 | 1.31E-51 | 6 | PIM3 |
| 6.84E-56 | 0.25735914 | 0.462 | 0.275 | 1.66E-51 | 6 | TMEM147 |
| 1.09E-55 | 0.27283603 | 0.475 | 0.288 | 2.64E-51 | 6 | ATP5F1A |
| 1.54E-55 | 0.28583309 | 0.526 | 0.334 | 3.73E-51 | 6 | SELENOW |
| 3.01E-55 | 0.29861238 | 0.699 | 0.499 | 7.29E-51 | 6 | NDUFA11 |
| 5.71E-55 | 0.28935815 | 0.66 | 0.457 | 1.38E-50 | 6 | HSBP1 |
| 7.10E-55 | 0.27351843 | 0.965 | 0.904 | 1.72E-50 | 6 | CDC42 |
| 8.83E-55 | 0.28041673 | 0.567 | 0.368 | 2.14E-50 | 6 | BAX |
| 9.58E-55 | 0.27225057 | 0.549 | 0.351 | 2.32E-50 | 6 | LAMTOR5 |
| 1.74E-54 | 0.28944319 | 0.543 | 0.353 | 4.22E-50 | 6 | PFDN2 |
| 2.23E-54 | 0.26692937 | 0.513 | 0.323 | 5.41E-50 | 6 | LMAN2 |
| 2.72E-54 | 0.28590162 | 0.376 | 0.204 | 6.59E-50 | 6 | MCEMP1 |
| 2.91E-54 | 0.29562918 | 0.651 | 0.449 | 7.05E-50 | 6 | DRAP1 |
| 3.17E-54 | 0.28788286 | 0.681 | 0.479 | 7.69E-50 | 6 | RNH1 |
| 4.27E-54 | 0.3011484 | 0.549 | 0.363 | 1.03E-49 | 6 | COMT |

| | | | | | | |
|---|---|---|---|---|---|---|
| 4.90E-54 | 0.26193095 | 0.999 | 0.993 | 1.19E-49 | 6 | RPS2 |
| 9.54E-54 | 0.26589665 | 0.566 | 0.367 | 2.31E-49 | 6 | SELENOF |
| 1.10E-53 | 0.28858408 | 0.679 | 0.466 | 2.67E-49 | 6 | ARL8B |
| 9.10E-53 | 0.2716084 | 0.67 | 0.466 | 2.21E-48 | 6 | SNRPD2 |
| 1.18E-52 | 0.28571829 | 0.735 | 0.537 | 2.86E-48 | 6 | SNHG8 |
| 2.28E-52 | 0.3011441 | 0.492 | 0.299 | 5.52E-48 | 6 | SORL1 |
| 9.32E-52 | 0.28573159 | 0.447 | 0.268 | 2.26E-47 | 6 | ACSL4 |
| 9.36E-52 | 0.25336905 | 0.468 | 0.283 | 2.27E-47 | 6 | SLC16A3 |
| 1.40E-51 | 0.25314169 | 0.525 | 0.334 | 3.39E-47 | 6 | PPP1CA |
| 1.87E-51 | 0.27045715 | 0.58 | 0.385 | 4.53E-47 | 6 | SNRPB |
| 2.19E-51 | 0.36511357 | 0.936 | 0.831 | 5.31E-47 | 6 | TNFAIP3 |
| 3.59E-51 | 0.3074658 | 0.688 | 0.503 | 8.71E-47 | 6 | HMGN1 |
| 5.42E-51 | 0.2535444 | 0.985 | 0.947 | 1.31E-46 | 6 | ARPC3 |
| 7.43E-51 | 0.25559198 | 0.938 | 0.845 | 1.80E-46 | 6 | TMBIM6 |
| 7.96E-51 | 0.2615467 | 0.596 | 0.394 | 1.93E-46 | 6 | ZYX |
| 8.31E-51 | 0.28083265 | 0.663 | 0.465 | 2.01E-46 | 6 | PSMB3 |
| 1.41E-50 | 0.25197328 | 0.541 | 0.348 | 3.43E-46 | 6 | ZNHIT1 |
| 2.48E-50 | 0.31389025 | 0.824 | 0.692 | 6.02E-46 | 6 | TUBA1B |
| 4.39E-50 | 0.27731432 | 0.931 | 0.851 | 1.06E-45 | 6 | TPM3 |
| 1.53E-49 | 0.25175793 | 0.949 | 0.878 | 3.71E-45 | 6 | SRP14 |
| 2.03E-49 | 0.26843095 | 0.642 | 0.444 | 4.92E-45 | 6 | ARHGDIA |
| 2.49E-49 | 0.26305651 | 0.528 | 0.341 | 6.04E-45 | 6 | NDUFC2 |
| 1.24E-48 | 0.32623808 | 0.722 | 0.547 | 3.00E-44 | 6 | GNA13 |
| 8.31E-48 | 0.82038609 | 0.338 | 0.199 | 2.02E-43 | 6 | IFIT3 |
| 9.90E-48 | 0.26964864 | 0.911 | 0.796 | 2.40E-43 | 6 | COX6B1 |
| 1.15E-47 | 0.25839004 | 0.505 | 0.323 | 2.78E-43 | 6 | IFI27L2 |
| 1.75E-47 | 0.26735716 | 0.987 | 0.84 | 4.25E-43 | 6 | PLAUR |
| 5.77E-47 | 0.30158483 | 0.812 | 0.655 | 1.40E-42 | 6 | ATP5MPL |
| 6.59E-47 | 0.25223411 | 0.615 | 0.42 | 1.60E-42 | 6 | C19orf53 |
| 7.21E-47 | 0.28024272 | 0.96 | 0.901 | 1.75E-42 | 6 | COX7C |
| 9.54E-47 | 0.26404448 | 0.687 | 0.496 | 2.31E-42 | 6 | TBCA |
| 1.85E-46 | 0.2720227 | 0.673 | 0.483 | 4.47E-42 | 6 | ATP5PO |
| 2.39E-46 | 0.27311388 | 0.595 | 0.414 | 5.79E-42 | 6 | SYNGR2 |
| 9.49E-46 | 0.25688267 | 0.663 | 0.471 | 2.30E-41 | 6 | COPE |
| 1.90E-45 | 0.27962643 | 0.741 | 0.564 | 4.60E-41 | 6 | BLOC1S1 |
| 4.38E-45 | 0.2710662 | 0.332 | 0.19 | 1.06E-40 | 6 | LDLRAD4 |
| 8.00E-45 | 0.2772569 | 0.547 | 0.374 | 1.94E-40 | 6 | HAVCR2 |
| 3.37E-44 | 0.25532153 | 0.666 | 0.473 | 8.17E-40 | 6 | C4orf48 |
| 3.39E-44 | 0.73770534 | 0.619 | 0.474 | 8.22E-40 | 6 | CXCL3 |
| 6.77E-44 | 0.25444753 | 0.484 | 0.314 | 1.64E-39 | 6 | ABRACL |
| 7.11E-44 | 0.25867996 | 0.473 | 0.297 | 1.72E-39 | 6 | GPR84 |
| 4.27E-43 | 0.25592107 | 0.767 | 0.583 | 1.04E-38 | 6 | PET100 |
| 1.79E-42 | 0.26001729 | 0.514 | 0.334 | 4.33E-38 | 6 | USP12 |

| | | | | | | |
|---|---|---|---|---|---|---|
| 2.90E-42 | 0.25324834 | 0.727 | 0.541 | 7.02E-38 | 6 | ATP5PD |
| 4.64E-42 | 0.27729908 | 0.833 | 0.696 | 1.13E-37 | 6 | MYL12A |
| 2.80E-41 | 0.25865834 | 0.916 | 0.755 | 6.78E-37 | 6 | COTL1 |
| 3.45E-41 | 0.27447697 | 0.994 | 0.978 | 8.37E-37 | 6 | FCER1G |
| 1.07E-40 | 0.46859331 | 0.706 | 0.536 | 2.59E-36 | 6 | MT2A |
| 2.27E-39 | 0.35864813 | 0.828 | 0.658 | 5.51E-35 | 6 | G0S2 |
| 2.91E-38 | 0.330295 | 0.992 | 0.968 | 7.04E-34 | 6 | BTG1 |
| 6.19E-38 | 0.27174682 | 0.643 | 0.485 | 1.50E-33 | 6 | CHCHD10 |
| 2.05E-36 | 0.25125706 | 0.639 | 0.478 | 4.96E-32 | 6 | VASP |
| 2.90E-36 | 0.4808147 | 0.452 | 0.307 | 7.04E-32 | 6 | THBD |
| 8.47E-36 | 0.30429767 | 0.916 | 0.842 | 2.05E-31 | 6 | SGK1 |
| 1.50E-35 | 0.28406985 | 0.535 | 0.352 | 3.65E-31 | 6 | LUCAT1 |
| 1.86E-35 | 0.26446256 | 0.665 | 0.515 | 4.51E-31 | 6 | SERBP1 |
| 6.41E-29 | 0.28398941 | 0.946 | 0.796 | 1.55E-24 | 6 | IL1B |
| 2.07E-26 | 0.38016218 | 0.409 | 0.282 | 5.02E-22 | 6 | ADM |
| 1.59E-22 | 0.33764829 | 0.659 | 0.538 | 3.86E-18 | 6 | TNF |
| 6.46E-21 | 0.32239174 | 0.534 | 0.425 | 1.57E-16 | 6 | ALOX5AP |
| 4.85E-20 | 0.87809469 | 0.394 | 0.303 | 1.18E-15 | 6 | IFIT2 |
| 1.24E-19 | 0.30470633 | 0.85 | 0.761 | 3.00E-15 | 6 | CXCL2 |
| 1.14E-18 | 0.41472847 | 0.689 | 0.6 | 2.77E-14 | 6 | PTGS2 |
| 2.74E-17 | 0.38711324 | 0.773 | 0.733 | 6.65E-13 | 6 | PMAIP1 |
| 0 | 4.09982623 | 0.835 | 0.088 | 0 | 7 | CXCL10 |
| 0 | 3.35898706 | 0.567 | 0.034 | 0 | 7 | CXCL9 |
| 0 | 1.722189 | 0.704 | 0.206 | 0 | 7 | GBP1 |
| 0 | 1.52447934 | 0.387 | 0.008 | 0 | 7 | CXCL11 |
| 0 | 1.45071159 | 0.38 | 0.031 | 0 | 7 | IDO1 |
| 0 | 1.12746958 | 0.536 | 0.113 | 0 | 7 | GBP5 |
| 0 | 0.82907693 | 0.436 | 0.089 | 0 | 7 | SLAMF7 |
| 1.25E-300 | 0.57466282 | 0.356 | 0.061 | 3.03E-296 | 7 | ANKRD22 |
| 4.73E-286 | 0.92986368 | 0.55 | 0.149 | 1.15E-281 | 7 | GBP4 |
| 1.51E-274 | 1.44709544 | 0.866 | 0.471 | 3.66E-270 | 7 | CALHM6 |
| 1.08E-254 | 1.8652747 | 0.894 | 0.534 | 2.61E-250 | 7 | MT2A |
| 5.96E-245 | 0.66069013 | 1 | 1 | 1.45E-240 | 7 | B2M |
| 9.43E-215 | 1.06911554 | 0.707 | 0.305 | 2.29E-210 | 7 | VAMP5 |
| 1.02E-213 | 1.00953517 | 0.638 | 0.249 | 2.48E-209 | 7 | PPA1 |
| 1.93E-208 | 0.91315956 | 0.755 | 0.354 | 4.68E-204 | 7 | STAT1 |
| 2.10E-204 | 1.11734005 | 0.991 | 0.937 | 5.10E-200 | 7 | HLA-DRB1 |
| 8.99E-198 | 0.98047276 | 0.995 | 0.966 | 2.18E-193 | 7 | HLA-DRA |
| 1.59E-192 | 1.04705209 | 0.981 | 0.872 | 3.86E-188 | 7 | HLA-DPA1 |
| 1.02E-187 | 1.25580283 | 0.88 | 0.577 | 2.48E-183 | 7 | HLA-DQA1 |
| 1.74E-187 | 0.98061368 | 0.838 | 0.486 | 4.23E-183 | 7 | PSME2 |
| 2.73E-186 | 0.73952474 | 0.463 | 0.143 | 6.61E-182 | 7 | IL4I1 |
| 6.64E-178 | 0.75830702 | 0.523 | 0.186 | 1.61E-173 | 7 | PLAAT4 |

| | | | | | | |
|---|---|---|---|---|---|---|
| 8.43E-172 | 0.82301377 | 0.743 | 0.371 | 2.04E-167 | 7 | PSMB9 |
| 1.13E-171 | 1.05279851 | 0.97 | 0.83 | 2.74E-167 | 7 | HLA-DPB1 |
| 3.13E-168 | 0.79834724 | 0.444 | 0.139 | 7.58E-164 | 7 | ISG20 |
| 7.21E-165 | 1.34173814 | 0.885 | 0.631 | 1.75E-160 | 7 | C15orf48 |
| 1.13E-164 | 0.99300482 | 0.942 | 0.721 | 2.73E-160 | 7 | HLA-DQB1 |
| 1.69E-155 | 0.96101595 | 0.658 | 0.308 | 4.10E-151 | 7 | WARS |
| 3.33E-155 | 1.10267217 | 0.582 | 0.259 | 8.08E-151 | 7 | TNFSF10 |
| 1.42E-148 | 0.70804416 | 1 | 0.999 | 3.43E-144 | 7 | TMSB10 |
| 3.88E-146 | 1.78236798 | 0.805 | 0.521 | 9.41E-142 | 7 | ISG15 |
| 2.45E-142 | 0.33745427 | 0.28 | 0.068 | 5.95E-138 | 7 | SLAMF8 |
| 6.10E-137 | 0.63367416 | 0.998 | 0.992 | 1.48E-132 | 7 | CD74 |
| 1.30E-132 | 0.93621696 | 0.498 | 0.197 | 3.15E-128 | 7 | IFIT3 |
| 6.45E-131 | 0.47350766 | 0.418 | 0.145 | 1.56E-126 | 7 | TAP1 |
| 3.30E-124 | 0.7076912 | 0.711 | 0.394 | 7.99E-120 | 7 | TNFSF13B |
| 1.74E-121 | 0.54700486 | 1 | 0.997 | 4.21E-117 | 7 | HLA-B |
| 5.42E-116 | 0.33726135 | 0.255 | 0.067 | 1.31E-111 | 7 | CD80 |
| 2.14E-115 | 0.69167629 | 0.832 | 0.572 | 5.18E-111 | 7 | LAP3 |
| 3.28E-113 | 0.31019962 | 0.256 | 0.068 | 7.95E-109 | 7 | APOL3 |
| 1.28E-110 | 1.11890319 | 0.6 | 0.299 | 3.09E-106 | 7 | IFIT2 |
| 1.36E-110 | 0.78858153 | 0.713 | 0.407 | 3.30E-106 | 7 | SNX10 |
| 6.65E-109 | 1.24362662 | 0.764 | 0.519 | 1.61E-104 | 7 | TXN |
| 1.14E-102 | 0.5076465 | 0.995 | 0.974 | 2.77E-98 | 7 | HLA-C |
| 9.85E-102 | 0.64049399 | 0.809 | 0.544 | 2.39E-97 | 7 | CTSH |
| 4.84E-101 | 0.46030615 | 0.375 | 0.141 | 1.17E-96 | 7 | CD40 |
| 4.16E-99 | 0.51897713 | 0.459 | 0.199 | 1.01E-94 | 7 | PSTPIP2 |
| 7.99E-99 | 0.67649384 | 0.781 | 0.482 | 1.94E-94 | 7 | FGL2 |
| 5.18E-98 | 0.49470865 | 0.545 | 0.265 | 1.26E-93 | 7 | UBE2L6 |
| 5.60E-96 | 0.57470019 | 0.578 | 0.298 | 1.36E-91 | 7 | EPSTI1 |
| 7.88E-94 | 0.61072693 | 0.943 | 0.814 | 1.91E-89 | 7 | TYMP |
| 1.62E-89 | 0.53662526 | 0.465 | 0.212 | 3.92E-85 | 7 | ATF5 |
| 5.09E-88 | 0.6427068 | 0.291 | 0.101 | 1.23E-83 | 7 | RSAD2 |
| 8.83E-88 | 0.48399808 | 0.997 | 0.99 | 2.14E-83 | 7 | HLA-A |
| 6.36E-85 | 0.48632999 | 0.442 | 0.204 | 1.54E-80 | 7 | NUB1 |
| 1.23E-84 | 0.61723144 | 0.629 | 0.349 | 2.99E-80 | 7 | LY6E |
| 2.11E-84 | 0.36619645 | 0.337 | 0.125 | 5.12E-80 | 7 | PALM2-AKAP |
| 6.08E-82 | 1.17250423 | 0.641 | 0.387 | 1.47E-77 | 7 | APOBEC3A |
| 1.27E-80 | 0.59121725 | 0.515 | 0.256 | 3.07E-76 | 7 | CD69 |
| 1.09E-74 | 0.47001662 | 0.572 | 0.327 | 2.65E-70 | 7 | PSMB8 |
| 5.76E-74 | 0.74578278 | 0.741 | 0.526 | 1.40E-69 | 7 | IRF1 |
| 2.81E-72 | 0.46505047 | 0.906 | 0.728 | 6.81E-68 | 7 | HLA-DMA |
| 4.54E-72 | 0.49404091 | 0.981 | 0.933 | 1.10E-67 | 7 | S100A11 |
| 1.24E-71 | 0.44805362 | 0.399 | 0.183 | 3.00E-67 | 7 | IFIH1 |
| 1.39E-69 | 0.62356207 | 0.66 | 0.41 | 3.36E-65 | 7 | IFI6 |

| | | | | | | |
|---|---|---|---|---|---|---|
| 5.67E-69 | 0.34702214 | 0.399 | 0.184 | 1.37E-64 | 7 | LYSMD2 |
| 2.51E-67 | 0.40618509 | 0.473 | 0.239 | 6.08E-63 | 7 | SERPING1 |
| 6.08E-67 | 0.55451615 | 0.744 | 0.522 | 1.47E-62 | 7 | ATOX1 |
| 1.30E-66 | 0.49771884 | 0.491 | 0.267 | 3.15E-62 | 7 | NBN |
| 1.04E-65 | 0.74590969 | 0.882 | 0.751 | 2.51E-61 | 7 | IFITM3 |
| 6.50E-65 | 0.67664054 | 0.885 | 0.715 | 1.58E-60 | 7 | GPR183 |
| 1.96E-64 | 0.5110602 | 0.507 | 0.275 | 4.75E-60 | 7 | MIR155HG |
| 3.93E-64 | 0.50813754 | 0.627 | 0.384 | 9.54E-60 | 7 | GBP2 |
| 5.93E-64 | 0.39945281 | 0.461 | 0.241 | 1.44E-59 | 7 | SMCO4 |
| 1.88E-63 | 0.37117319 | 0.354 | 0.157 | 4.57E-59 | 7 | CLEC4E |
| 2.14E-60 | 0.37064882 | 0.34 | 0.152 | 5.19E-56 | 7 | SAMD9L |
| 3.09E-60 | 0.51353982 | 0.94 | 0.832 | 7.50E-56 | 7 | TNFAIP3 |
| 5.13E-60 | 0.40682409 | 0.572 | 0.335 | 1.24E-55 | 7 | PEA15 |
| 1.27E-58 | 0.47768472 | 0.821 | 0.651 | 3.09E-54 | 7 | POMP |
| 2.18E-58 | 0.44721468 | 0.855 | 0.645 | 5.28E-54 | 7 | SERPINA1 |
| 6.68E-58 | 0.39044287 | 0.634 | 0.389 | 1.62E-53 | 7 | HLA-F |
| 2.77E-56 | 0.46764143 | 0.533 | 0.306 | 6.71E-52 | 7 | IFI44L |
| 5.08E-56 | 0.30861285 | 0.358 | 0.169 | 1.23E-51 | 7 | MYOF |
| 6.30E-55 | 0.31413698 | 0.352 | 0.168 | 1.53E-50 | 7 | IFI35 |
| 3.57E-54 | 0.66402518 | 0.432 | 0.24 | 8.66E-50 | 7 | MX1 |
| 7.03E-54 | 0.50862781 | 0.663 | 0.45 | 1.70E-49 | 7 | PARP14 |
| 1.82E-53 | 0.37667968 | 0.999 | 0.991 | 4.42E-49 | 7 | RPS19 |
| 3.85E-53 | 0.70810103 | 0.947 | 0.902 | 9.35E-49 | 7 | CCL4L2 |
| 5.27E-53 | 0.54063178 | 0.648 | 0.443 | 1.28E-48 | 7 | IRF8 |
| 2.33E-52 | 0.44977701 | 0.944 | 0.799 | 5.64E-48 | 7 | BCL2A1 |
| 1.50E-51 | 0.86319254 | 0.819 | 0.66 | 3.64E-47 | 7 | G0S2 |
| 5.62E-51 | 0.38485391 | 0.674 | 0.451 | 1.36E-46 | 7 | DRAP1 |
| 3.56E-50 | 0.37920123 | 0.822 | 0.64 | 8.62E-46 | 7 | PSME1 |
| 3.66E-50 | 0.42096431 | 0.996 | 0.985 | 8.88E-46 | 7 | SOD2 |
| 8.36E-50 | 0.37733192 | 1 | 1 | 2.03E-45 | 7 | FTH1 |
| 7.11E-49 | 0.50324174 | 0.775 | 0.598 | 1.72E-44 | 7 | TNFAIP8 |
| 1.53E-48 | 0.39586248 | 0.424 | 0.233 | 3.70E-44 | 7 | RALA |
| 2.55E-48 | 0.4867836 | 0.84 | 0.684 | 6.17E-44 | 7 | DUSP2 |
| 4.40E-48 | 0.37650516 | 0.584 | 0.375 | 1.07E-43 | 7 | PSMA2 |
| 4.83E-48 | 0.29431352 | 0.369 | 0.189 | 1.17E-43 | 7 | NMI |
| 1.28E-47 | 0.35413583 | 0.54 | 0.33 | 3.11E-43 | 7 | SCO2 |
| 2.79E-47 | 0.31866628 | 0.421 | 0.231 | 6.76E-43 | 7 | SRI |
| 1.22E-46 | 0.28778995 | 0.309 | 0.147 | 2.95E-42 | 7 | OAS2 |
| 1.53E-46 | 0.39128175 | 0.973 | 0.844 | 3.72E-42 | 7 | IFI30 |
| 3.27E-46 | 0.51885635 | 0.801 | 0.67 | 7.92E-42 | 7 | PRDX1 |
| 5.45E-46 | 0.51151187 | 0.948 | 0.883 | 1.32E-41 | 7 | PLEK |
| 9.48E-46 | 0.43327248 | 0.534 | 0.338 | 2.30E-41 | 7 | ALDH2 |
| 9.74E-46 | 0.71259223 | 0.576 | 0.38 | 2.36E-41 | 7 | HLA-DRB5 |

| | | | | | | |
|---|---|---|---|---|---|---|
| 1.10E-45 | 0.37440146 | 0.484 | 0.29 | 2.67E-41 | 7 | APOL6 |
| 9.17E-45 | 0.38446999 | 0.772 | 0.595 | 2.22E-40 | 7 | BST2 |
| 1.15E-44 | 0.39142109 | 0.574 | 0.382 | 2.79E-40 | 7 | PSMA4 |
| 3.76E-44 | 0.37144103 | 0.695 | 0.503 | 9.11E-40 | 7 | TAPBP |
| 6.01E-44 | 0.27272328 | 1 | 1 | 1.46E-39 | 7 | EEF1A1 |
| 7.72E-44 | 0.36006755 | 0.571 | 0.376 | 1.87E-39 | 7 | PSMB10 |
| 4.64E-43 | 0.37773578 | 0.886 | 0.759 | 1.12E-38 | 7 | CD81 |
| 8.73E-43 | 0.31110622 | 0.379 | 0.201 | 2.12E-38 | 7 | FPR3 |
| 2.20E-41 | 0.30093656 | 0.465 | 0.278 | 5.32E-37 | 7 | ETV6 |
| 2.73E-41 | 0.34043237 | 0.629 | 0.436 | 6.62E-37 | 7 | PSMA6 |
| 2.96E-41 | 0.35002011 | 0.422 | 0.246 | 7.18E-37 | 7 | IRF7 |
| 3.79E-41 | 0.38466524 | 0.734 | 0.536 | 9.20E-37 | 7 | IGSF6 |
| 7.70E-41 | 0.3018378 | 0.426 | 0.245 | 1.87E-36 | 7 | SQOR |
| 1.20E-40 | 0.34124496 | 0.937 | 0.853 | 2.90E-36 | 7 | GPX4 |
| 2.23E-40 | 0.57632419 | 0.971 | 0.946 | 5.40E-36 | 7 | CCL4 |
| 2.32E-40 | 0.34941371 | 0.997 | 0.993 | 5.64E-36 | 7 | NFKBIA |
| 5.63E-40 | 0.30199121 | 0.999 | 0.995 | 1.36E-35 | 7 | RPL28 |
| 9.49E-40 | 0.29275591 | 0.641 | 0.408 | 2.30E-35 | 7 | CD48 |
| 1.05E-38 | 0.34876956 | 0.971 | 0.933 | 2.54E-34 | 7 | CD83 |
| 1.06E-38 | 0.29585868 | 0.425 | 0.246 | 2.57E-34 | 7 | GCH1 |
| 1.97E-38 | 0.34912753 | 0.784 | 0.622 | 4.78E-34 | 7 | DBI |
| 2.81E-38 | 0.36921592 | 0.694 | 0.513 | 6.82E-34 | 7 | RIPK2 |
| 3.47E-38 | 0.35371721 | 0.468 | 0.291 | 8.40E-34 | 7 | DYNLT1 |
| 3.47E-38 | 0.35015198 | 0.751 | 0.564 | 8.42E-34 | 7 | CLEC2B |
| 4.31E-38 | 0.34691337 | 0.998 | 0.978 | 1.05E-33 | 7 | SRGN |
| 2.21E-36 | 0.32399548 | 0.71 | 0.496 | 5.35E-32 | 7 | CAPG |
| 1.03E-35 | 0.39505249 | 0.947 | 0.811 | 2.50E-31 | 7 | LYZ |
| 1.22E-35 | 0.27546112 | 0.992 | 0.976 | 2.97E-31 | 7 | SERF2 |
| 1.48E-35 | 0.39872232 | 0.481 | 0.31 | 3.59E-31 | 7 | PDGFB |
| 2.88E-35 | 0.3452762 | 0.783 | 0.625 | 6.98E-31 | 7 | PDE4B |
| 3.05E-35 | 0.30823093 | 0.973 | 0.931 | 7.39E-31 | 7 | PFN1 |
| 5.88E-35 | 0.52355191 | 0.781 | 0.64 | 1.42E-30 | 7 | RGS1 |
| 1.14E-34 | 0.33777101 | 0.49 | 0.323 | 2.75E-30 | 7 | PTPN1 |
| 1.16E-34 | 0.2953891 | 0.514 | 0.324 | 2.80E-30 | 7 | RNF19B |
| 1.49E-34 | 0.33439273 | 0.718 | 0.548 | 3.61E-30 | 7 | HLA-DMB |
| 2.25E-34 | 0.37180443 | 0.819 | 0.664 | 5.44E-30 | 7 | FCGR3A |
| 3.31E-34 | 0.36171305 | 0.511 | 0.337 | 8.01E-30 | 7 | XAF1 |
| 1.70E-33 | 0.2862591 | 0.789 | 0.607 | 4.11E-29 | 7 | NDUFB2 |
| 2.43E-33 | 0.35669904 | 0.974 | 0.917 | 5.90E-29 | 7 | CCL3L1 |
| 7.24E-33 | 0.27469852 | 0.432 | 0.267 | 1.76E-28 | 7 | TRIM22 |
| 2.32E-32 | 0.31061439 | 0.413 | 0.255 | 5.63E-28 | 7 | SP110 |
| 8.00E-32 | 0.36405306 | 0.454 | 0.285 | 1.94E-27 | 7 | C3 |
| 8.37E-32 | 0.34141088 | 0.637 | 0.456 | 2.03E-27 | 7 | PFKFB3 |

| | | | | | | |
|---|---|---|---|---|---|---|
| 5.30E-31 | 0.31629293 | 0.864 | 0.746 | 1.28E-26 | 7 | MIF |
| 8.91E-31 | 0.29046984 | 0.566 | 0.388 | 2.16E-26 | 7 | PHPT1 |
| 9.33E-31 | 0.34168137 | 0.568 | 0.408 | 2.26E-26 | 7 | GLRX |
| 1.26E-30 | 0.27871615 | 0.465 | 0.304 | 3.05E-26 | 7 | TFEC |
| 1.36E-30 | 0.27488115 | 0.649 | 0.477 | 3.29E-26 | 7 | PSMB1 |
| 3.74E-30 | 0.32383157 | 0.771 | 0.592 | 9.07E-26 | 7 | CXCR4 |
| 8.44E-30 | 0.26242099 | 0.994 | 0.979 | 2.05E-25 | 7 | ATP5F1E |
| 8.68E-30 | 0.2799181 | 0.504 | 0.336 | 2.10E-25 | 7 | LILRB4 |
| 1.84E-29 | 0.30315266 | 0.798 | 0.658 | 4.47E-25 | 7 | COX7B |
| 1.93E-29 | 0.37353 | 0.597 | 0.429 | 4.69E-25 | 7 | NR4A3 |
| 3.88E-29 | 0.273989 | 0.645 | 0.466 | 9.40E-25 | 7 | PRELID1 |
| 5.35E-29 | 0.25318561 | 0.451 | 0.29 | 1.30E-24 | 7 | RAB8A |
| 7.65E-29 | 0.29925436 | 0.831 | 0.717 | 1.86E-24 | 7 | RGS10 |
| 2.20E-28 | 0.26454028 | 0.516 | 0.351 | 5.33E-24 | 7 | CASP4 |
| 2.91E-28 | 0.34286284 | 0.77 | 0.637 | 7.04E-24 | 7 | LITAF |
| 4.68E-28 | 0.27868884 | 0.842 | 0.645 | 1.14E-23 | 7 | LGALS3 |
| 4.85E-28 | 0.25466268 | 0.458 | 0.3 | 1.18E-23 | 7 | DNAJC15 |
| 6.70E-28 | 0.27424288 | 0.425 | 0.272 | 1.62E-23 | 7 | DRAM1 |
| 1.48E-27 | 0.25630111 | 0.898 | 0.814 | 3.58E-23 | 7 | NME2 |
| 1.56E-27 | 0.33678227 | 0.714 | 0.566 | 3.77E-23 | 7 | RPS27L |
| 3.07E-27 | 0.34695138 | 0.698 | 0.505 | 7.44E-23 | 7 | IL1RN |
| 3.33E-27 | 0.31929574 | 0.996 | 0.992 | 8.07E-23 | 7 | CCL3 |
| 1.02E-26 | 0.25847393 | 0.934 | 0.877 | 2.48E-22 | 7 | CTSZ |
| 4.34E-26 | 0.26454112 | 0.514 | 0.36 | 1.05E-21 | 7 | GABARAPL2 |
| 5.93E-26 | 0.32557985 | 0.725 | 0.577 | 1.44E-21 | 7 | NCF1 |
| 7.88E-26 | 0.27791585 | 0.324 | 0.195 | 1.91E-21 | 7 | OAS1 |
| 9.25E-26 | 0.26039356 | 0.725 | 0.54 | 2.24E-21 | 7 | BHLHE40 |
| 1.48E-25 | 0.25402527 | 0.63 | 0.467 | 3.58E-21 | 7 | PSMB3 |
| 1.81E-25 | 0.31815677 | 0.488 | 0.342 | 4.38E-21 | 7 | SSB |
| 1.83E-25 | 0.27950586 | 0.783 | 0.655 | 4.43E-21 | 7 | ATP5MD |
| 2.23E-25 | 0.26381596 | 0.669 | 0.497 | 5.41E-21 | 7 | PIM3 |
| 2.34E-25 | 0.32093665 | 0.359 | 0.221 | 5.67E-21 | 7 | FBP1 |
| 3.95E-25 | 0.31575641 | 0.463 | 0.305 | 9.57E-21 | 7 | LGALS2 |
| 4.04E-25 | 0.27138475 | 0.797 | 0.649 | 9.79E-21 | 7 | C1orf162 |
| 1.30E-24 | 0.32951202 | 0.654 | 0.516 | 3.16E-20 | 7 | PLEKHO1 |
| 3.86E-24 | 0.28804359 | 0.309 | 0.182 | 9.35E-20 | 7 | CCL5 |
| 9.49E-24 | 0.26533778 | 0.816 | 0.685 | 2.30E-19 | 7 | SUB1 |
| 7.39E-23 | 0.28715603 | 0.462 | 0.315 | 1.79E-18 | 7 | BIRC3 |
| 1.23E-22 | 0.25303974 | 1 | 0.999 | 2.97E-18 | 7 | ACTB |
| 2.00E-22 | 0.27400483 | 0.75 | 0.629 | 4.85E-18 | 7 | COX7A2 |
| 4.97E-22 | 0.25131868 | 0.812 | 0.714 | 1.20E-17 | 7 | MYL12B |
| 1.26E-21 | 0.27388267 | 0.689 | 0.547 | 3.06E-17 | 7 | NFE2L2 |
| 2.49E-21 | 0.25994841 | 0.68 | 0.548 | 6.03E-17 | 7 | LYN |

| | | | | | | |
|---|---|---|---|---|---|---|
| 2.74E-21 | 0.32176612 | 0.623 | 0.49 | 6.65E-17 | 7 | RNF213 |
| 3.12E-21 | 0.2638169 | 0.533 | 0.39 | 7.56E-17 | 7 | BAZ1A |
| 5.39E-21 | 0.2510508 | 0.607 | 0.463 | 1.31E-16 | 7 | PTGER4 |
| 9.70E-21 | 0.3566143 | 0.692 | 0.571 | 2.35E-16 | 7 | TNFAIP2 |
| 8.69E-20 | 0.27507344 | 0.98 | 0.981 | 2.11E-15 | 7 | PNRC1 |
| 1.03E-19 | 0.39658257 | 0.776 | 0.691 | 2.49E-15 | 7 | CHMP1B |
| 9.47E-17 | 0.2501887 | 0.424 | 0.302 | 2.30E-12 | 7 | ZC3HAV1 |
| 1.84E-16 | 0.29249522 | 0.988 | 0.979 | 4.45E-12 | 7 | IER3 |
| 3.21E-14 | 0.31335369 | 0.302 | 0.206 | 7.79E-10 | 7 | DNAAF1 |
| 1.27E-12 | 0.30428632 | 0.327 | 0.242 | 3.07E-08 | 7 | MX2 |
| 1.78E-11 | 0.25447199 | 0.888 | 0.844 | 4.32E-07 | 7 | SGK1 |
| 2.93E-11 | 0.25049285 | 0.795 | 0.716 | 7.10E-07 | 7 | CYCS |
| 3.07E-08 | 0.32750197 | 0.607 | 0.54 | 0.00074473 | 7 | TNF |
| 0 | 3.55180766 | 1 | 0.113 | 0 | 8 | IFI27 |
| 1.08E-133 | 1.18787386 | 0.706 | 0.379 | 2.61E-129 | 8 | HLA-DRB5 |
| 1.45E-125 | 0.72302956 | 1 | 0.992 | 3.52E-121 | 8 | CD74 |
| 4.91E-112 | 0.78350045 | 0.99 | 0.872 | 1.19E-107 | 8 | HLA-DPA1 |
| 5.93E-103 | 0.43040211 | 1 | 1 | 1.44E-98 | 8 | B2M |
| 6.03E-98 | 0.71778162 | 0.717 | 0.41 | 1.46E-93 | 8 | IFI6 |
| 9.00E-98 | 0.89293202 | 0.668 | 0.352 | 2.18E-93 | 8 | GPNMB |
| 1.01E-97 | 0.67195911 | 0.994 | 0.933 | 2.46E-93 | 8 | S100A11 |
| 1.10E-93 | 0.64168419 | 1 | 0.966 | 2.68E-89 | 8 | HLA-DRA |
| 4.58E-93 | 0.64839106 | 0.493 | 0.204 | 1.11E-88 | 8 | HLA-DQA2 |
| 9.12E-88 | 0.6494615 | 0.991 | 0.83 | 2.21E-83 | 8 | HLA-DPB1 |
| 8.63E-87 | 0.98336627 | 0.855 | 0.639 | 2.09E-82 | 8 | RGS1 |
| 3.08E-79 | 0.68664865 | 0.996 | 0.938 | 7.47E-75 | 8 | HLA-DRB1 |
| 2.50E-75 | 0.58522389 | 0.951 | 0.722 | 6.07E-71 | 8 | HLA-DQB1 |
| 3.04E-75 | 0.71888755 | 0.939 | 0.745 | 7.38E-71 | 8 | LGALS1 |
| 2.76E-74 | 0.63280891 | 0.621 | 0.35 | 6.70E-70 | 8 | LY6E |
| 3.85E-73 | 0.57548308 | 0.531 | 0.26 | 9.33E-69 | 8 | TREM2 |
| 6.04E-71 | 0.55567279 | 0.792 | 0.545 | 1.46E-66 | 8 | CTSH |
| 3.59E-70 | 0.6069112 | 0.986 | 0.844 | 8.70E-66 | 8 | IFI30 |
| 2.34E-69 | 0.7087923 | 0.857 | 0.579 | 5.68E-65 | 8 | HLA-DQA1 |
| 5.20E-69 | 0.46560828 | 1 | 0.99 | 1.26E-64 | 8 | HLA-A |
| 7.56E-69 | 0.45542579 | 0.995 | 0.976 | 1.83E-64 | 8 | SERF2 |
| 3.59E-67 | 0.61983675 | 0.749 | 0.496 | 8.70E-63 | 8 | CAPG |
| 4.05E-64 | 0.80363841 | 0.766 | 0.523 | 9.83E-60 | 8 | ISG15 |
| 5.03E-63 | 0.79585276 | 0.772 | 0.522 | 1.22E-58 | 8 | APOC1 |
| 6.76E-60 | 0.51683098 | 0.914 | 0.759 | 1.64E-55 | 8 | CD81 |
| 2.07E-58 | 0.73160262 | 0.323 | 0.132 | 5.01E-54 | 8 | HAMP |
| 3.31E-58 | 0.54622984 | 0.465 | 0.231 | 8.03E-54 | 8 | SDS |
| 1.92E-57 | 0.68952761 | 0.951 | 0.926 | 4.66E-53 | 8 | CTSD |
| 1.66E-56 | 0.31056855 | 1 | 1 | 4.02E-52 | 8 | TMSB4X |

| | | | | | | |
|---|---|---|---|---|---|---|
| 3.86E-56 | 1.07980181 | 0.812 | 0.635 | 9.36E-52 | 8 | APOE |
| 8.01E-55 | 0.44544132 | 0.909 | 0.728 | 1.94E-50 | 8 | HLA-DMA |
| 4.03E-54 | 0.5342247 | 0.878 | 0.645 | 9.77E-50 | 8 | LGALS3 |
| 4.94E-54 | 0.38415178 | 1 | 0.999 | 1.20E-49 | 8 | TMSB10 |
| 9.42E-54 | 0.38538523 | 0.416 | 0.201 | 2.28E-49 | 8 | FPR3 |
| 3.92E-52 | 0.5014778 | 0.991 | 0.982 | 9.52E-48 | 8 | PSAP |
| 7.60E-52 | 0.37863983 | 0.999 | 0.991 | 1.84E-47 | 8 | RPS19 |
| 8.16E-48 | 0.49535781 | 0.524 | 0.307 | 1.98E-43 | 8 | IFI44L |
| 1.23E-47 | 0.36689337 | 0.354 | 0.169 | 2.98E-43 | 8 | LGALS3BP |
| 1.39E-47 | 0.46035585 | 0.635 | 0.411 | 3.38E-43 | 8 | JPT1 |
| 1.59E-47 | 0.36122597 | 0.998 | 0.974 | 3.86E-43 | 8 | NPC2 |
| 3.46E-47 | 0.44590302 | 0.57 | 0.331 | 8.40E-43 | 8 | CD9 |
| 2.41E-46 | 0.46701177 | 0.877 | 0.68 | 5.85E-42 | 8 | ANXA2 |
| 2.42E-46 | 0.46043744 | 0.723 | 0.523 | 5.88E-42 | 8 | ATOX1 |
| 4.70E-46 | 0.52870881 | 0.761 | 0.555 | 1.14E-41 | 8 | LMNA |
| 1.64E-45 | 0.38343808 | 1 | 0.996 | 3.98E-41 | 8 | CST3 |
| 1.91E-45 | 0.58081957 | 0.882 | 0.752 | 4.63E-41 | 8 | IFITM3 |
| 2.30E-45 | 0.45594495 | 0.934 | 0.873 | 5.58E-41 | 8 | GRN |
| 1.38E-44 | 0.30668186 | 0.303 | 0.131 | 3.35E-40 | 8 | MTRNR2L1 |
| 7.59E-44 | 0.41965616 | 0.85 | 0.664 | 1.84E-39 | 8 | FCGR3A |
| 2.91E-43 | 0.30648626 | 1 | 1 | 7.06E-39 | 8 | FTL |
| 4.56E-43 | 0.3694622 | 0.995 | 0.974 | 1.11E-38 | 8 | HLA-C |
| 7.53E-43 | 0.50869355 | 0.719 | 0.568 | 1.82E-38 | 8 | NOP10 |
| 3.12E-42 | 0.35987807 | 0.999 | 0.979 | 7.56E-38 | 8 | ATP5F1E |
| 3.25E-42 | 0.36754248 | 0.531 | 0.325 | 7.88E-38 | 8 | IFI27L2 |
| 4.15E-42 | 0.39974665 | 0.833 | 0.663 | 1.01E-37 | 8 | UQCR10 |
| 1.70E-41 | 0.35306427 | 0.944 | 0.814 | 4.13E-37 | 8 | NME2 |
| 2.57E-41 | 0.48611788 | 0.901 | 0.715 | 6.23E-37 | 8 | GPR183 |
| 2.63E-40 | 0.43774035 | 0.741 | 0.566 | 6.37E-36 | 8 | RPS27L |
| 8.92E-40 | 0.40684464 | 0.783 | 0.607 | 2.16E-35 | 8 | NDUFB2 |
| 9.02E-40 | 0.36140756 | 0.894 | 0.746 | 2.19E-35 | 8 | MIF |
| 4.28E-39 | 0.40290732 | 0.823 | 0.649 | 1.04E-34 | 8 | C1orf162 |
| 4.72E-39 | 0.39273828 | 0.869 | 0.725 | 1.14E-34 | 8 | ATP6V1F |
| 8.16E-39 | 0.38420464 | 0.895 | 0.788 | 1.98E-34 | 8 | VAMP8 |
| 1.07E-38 | 0.39943414 | 0.794 | 0.622 | 2.59E-34 | 8 | DBI |
| 1.35E-38 | 0.33163691 | 0.993 | 0.962 | 3.28E-34 | 8 | MYL6 |
| 1.34E-37 | 0.27156986 | 1 | 0.995 | 3.24E-33 | 8 | TYROBP |
| 2.07E-37 | 0.39696458 | 0.485 | 0.285 | 5.01E-33 | 8 | C3 |
| 3.02E-37 | 0.36130006 | 0.929 | 0.814 | 7.32E-33 | 8 | TYMP |
| 3.96E-37 | 0.3593369 | 0.494 | 0.3 | 9.61E-33 | 8 | EPSTI1 |
| 1.70E-36 | 0.32742453 | 0.974 | 0.939 | 4.13E-32 | 8 | CFL1 |
| 1.30E-35 | 0.45129955 | 0.88 | 0.771 | 3.16E-31 | 8 | CSTB |
| 1.59E-34 | 0.63588982 | 0.729 | 0.567 | 3.85E-30 | 8 | FABP5 |

| | | | | | | |
|---|---|---|---|---|---|---|
| 6.25E-34 | 0.43482298 | 0.784 | 0.671 | 1.52E-29 | 8 | PRDX1 |
| 7.37E-34 | 0.33130713 | 0.883 | 0.753 | 1.79E-29 | 8 | GNG5 |
| 1.27E-33 | 0.36000609 | 0.817 | 0.658 | 3.07E-29 | 8 | COX7B |
| 1.76E-33 | 0.34903398 | 0.83 | 0.679 | 4.26E-29 | 8 | NDUFA1 |
| 2.20E-33 | 0.45765022 | 0.358 | 0.196 | 5.32E-29 | 8 | GCHFR |
| 8.66E-33 | 0.3879501 | 0.789 | 0.592 | 2.10E-28 | 8 | CXCR4 |
| 3.52E-32 | 0.35815707 | 0.979 | 0.931 | 8.52E-28 | 8 | PFN1 |
| 1.05E-31 | 0.39894778 | 0.69 | 0.516 | 2.55E-27 | 8 | ZNF331 |
| 5.30E-31 | 0.482096 | 0.939 | 0.859 | 1.29E-26 | 8 | PLIN2 |
| 1.50E-30 | 0.3684483 | 0.435 | 0.258 | 3.63E-26 | 8 | CD69 |
| 2.32E-30 | 0.32045154 | 0.824 | 0.689 | 5.63E-26 | 8 | COX6C |
| 2.33E-30 | 0.32732513 | 0.993 | 0.97 | 5.65E-26 | 8 | CD63 |
| 2.43E-30 | 0.38453621 | 0.67 | 0.505 | 5.89E-26 | 8 | YWHAH |
| 2.44E-30 | 0.30600542 | 0.914 | 0.798 | 5.91E-26 | 8 | COX6B1 |
| 3.19E-30 | 0.35321678 | 0.777 | 0.652 | 7.73E-26 | 8 | POMP |
| 3.41E-30 | 0.27396652 | 0.999 | 0.988 | 8.26E-26 | 8 | CYBA |
| 7.17E-30 | 0.53489361 | 0.704 | 0.538 | 1.74E-25 | 8 | MT2A |
| 1.21E-29 | 0.34284818 | 0.646 | 0.474 | 2.93E-25 | 8 | RBX1 |
| 3.12E-29 | 0.26449108 | 0.963 | 0.912 | 7.57E-25 | 8 | PPIA |
| 8.08E-29 | 0.2845578 | 0.951 | 0.879 | 1.96E-24 | 8 | SRP14 |
| 1.33E-28 | 0.35378497 | 0.769 | 0.634 | 3.24E-24 | 8 | DYNLL1 |
| 1.34E-28 | 0.28654144 | 0.848 | 0.633 | 3.25E-24 | 8 | C15orf48 |
| 1.81E-28 | 0.42627082 | 0.928 | 0.843 | 4.38E-24 | 8 | SGK1 |
| 2.24E-28 | 0.29586301 | 0.57 | 0.384 | 5.42E-24 | 8 | PPDPF |
| 2.78E-28 | 0.26283054 | 1 | 1 | 6.74E-24 | 8 | FTH1 |
| 6.20E-28 | 0.30047085 | 0.928 | 0.854 | 1.50E-23 | 8 | GPX4 |
| 1.76E-27 | 0.31530897 | 0.985 | 0.912 | 4.27E-23 | 8 | SH3BGRL3 |
| 2.62E-27 | 0.31160755 | 0.732 | 0.58 | 6.36E-23 | 8 | NDUFS5 |
| 4.01E-27 | 0.34012 | 0.607 | 0.444 | 9.73E-23 | 8 | MGST3 |
| 6.79E-27 | 0.36974904 | 0.662 | 0.5 | 1.65E-22 | 8 | MSR1 |
| 5.88E-26 | 0.26077693 | 0.914 | 0.752 | 1.42E-21 | 8 | TSPO |
| 6.49E-26 | 0.31252408 | 0.653 | 0.49 | 1.57E-21 | 8 | NDUFA3 |
| 9.24E-26 | 0.29344798 | 0.87 | 0.737 | 2.24E-21 | 8 | ATP5ME |
| 1.39E-25 | 0.28705615 | 0.849 | 0.721 | 3.38E-21 | 8 | NDUFA4 |
| 1.76E-25 | 0.29377341 | 0.994 | 0.978 | 4.27E-21 | 8 | FCER1G |
| 2.83E-25 | 0.32607487 | 0.926 | 0.726 | 6.86E-21 | 8 | S100A10 |
| 3.35E-25 | 0.317268 | 0.799 | 0.676 | 8.11E-21 | 8 | SSR4 |
| 1.94E-24 | 0.32004649 | 0.385 | 0.241 | 4.70E-20 | 8 | MX1 |
| 2.05E-24 | 0.29032704 | 0.556 | 0.389 | 4.98E-20 | 8 | PHPT1 |
| 2.06E-24 | 0.31990741 | 0.696 | 0.544 | 4.98E-20 | 8 | PLXDC2 |
| 2.91E-24 | 0.30132134 | 0.777 | 0.655 | 7.05E-20 | 8 | ATP5MD |
| 3.72E-24 | 0.28491397 | 0.935 | 0.886 | 9.01E-20 | 8 | CD68 |
| 4.22E-24 | 0.31344933 | 0.688 | 0.536 | 1.02E-19 | 8 | PPT1 |

| | | | | | | |
|---|---|---|---|---|---|---|
| 7.63E-24 | 0.27398843 | 0.825 | 0.71 | 1.85E-19 | 8 | UBL5 |
| 8.57E-24 | 0.34725997 | 0.867 | 0.75 | 2.08E-19 | 8 | ATF3 |
| 9.03E-24 | 0.32876362 | 0.772 | 0.658 | 2.19E-19 | 8 | PPIB |
| 2.40E-23 | 0.28000397 | 0.889 | 0.707 | 5.82E-19 | 8 | OLR1 |
| 2.95E-23 | 0.26370051 | 0.985 | 0.961 | 7.15E-19 | 8 | LAPTM5 |
| 7.18E-23 | 0.28042005 | 0.763 | 0.641 | 1.74E-18 | 8 | ATP5MF |
| 1.42E-22 | 0.28596746 | 0.599 | 0.453 | 3.45E-18 | 8 | DRAP1 |
| 1.43E-22 | 0.27544479 | 0.759 | 0.629 | 3.47E-18 | 8 | COX7A2 |
| 1.50E-22 | 0.2785811 | 0.828 | 0.719 | 3.62E-18 | 8 | COX5B |
| 1.15E-21 | 0.30604534 | 0.616 | 0.472 | 2.78E-17 | 8 | CIB1 |
| 1.27E-21 | 0.25731197 | 1 | 0.999 | 3.07E-17 | 8 | ACTB |
| 1.27E-21 | 0.26922889 | 0.655 | 0.51 | 3.08E-17 | 8 | ATP5PF |
| 4.20E-21 | 0.28925716 | 0.906 | 0.828 | 1.02E-16 | 8 | CALM1 |
| 5.49E-21 | 0.3255213 | 0.938 | 0.759 | 1.33E-16 | 8 | S100A6 |
| 1.16E-20 | 0.2778279 | 0.835 | 0.743 | 2.81E-16 | 8 | UQCR11 |
| 2.14E-20 | 0.27548778 | 0.32 | 0.195 | 5.19E-16 | 8 | ACP5 |
| 2.71E-20 | 0.26206992 | 0.678 | 0.53 | 6.58E-16 | 8 | ATP5MC3 |
| 7.40E-20 | 0.27115604 | 0.857 | 0.751 | 1.79E-15 | 8 | ELOB |
| 1.44E-19 | 0.28525473 | 0.602 | 0.464 | 3.49E-15 | 8 | ADA2 |
| 9.95E-19 | 0.25456008 | 0.762 | 0.658 | 2.41E-14 | 8 | ATP6V0E1 |
| 3.61E-18 | 0.26034989 | 0.964 | 0.782 | 8.76E-14 | 8 | S100A4 |
| 4.58E-18 | 0.26377001 | 0.666 | 0.549 | 1.11E-13 | 8 | HLA-DMB |
| 8.07E-17 | 0.26255987 | 0.791 | 0.687 | 1.96E-12 | 8 | NDUFA13 |
| 2.31E-16 | 0.26114171 | 0.633 | 0.511 | 5.59E-12 | 8 | SSR3 |
| 3.48E-16 | 0.32038503 | 0.803 | 0.685 | 8.43E-12 | 8 | DUSP2 |
| 6.95E-16 | 0.30195709 | 0.685 | 0.597 | 1.68E-11 | 8 | BST2 |
| 2.66E-15 | 0.29197823 | 0.807 | 0.721 | 6.46E-11 | 8 | CFD |
| 3.19E-15 | 0.28330602 | 0.668 | 0.56 | 7.75E-11 | 8 | POLR2L |
| 4.23E-15 | 0.27273157 | 0.621 | 0.486 | 1.03E-10 | 8 | FGL2 |
| 6.64E-15 | 0.26532765 | 0.307 | 0.2 | 1.61E-10 | 8 | MARCO |
| 8.97E-15 | 0.25475672 | 0.531 | 0.418 | 2.17E-10 | 8 | SYNGR2 |
| 1.88E-14 | 0.267042 | 0.687 | 0.566 | 4.56E-10 | 8 | CLEC2B |
| 2.44E-14 | 0.26712802 | 0.378 | 0.264 | 5.91E-10 | 8 | PLAU |
| 2.79E-12 | 0.30146833 | 0.784 | 0.691 | 6.76E-08 | 8 | CHMP1B |
| 0 | 1.85503655 | 0.267 | 0.006 | 0 | 9 | CALD1 |
| 0 | 1.83684133 | 0.516 | 0.004 | 0 | 9 | MKI67 |
| 0 | 1.76482503 | 0.452 | 0.014 | 0 | 9 | CENPF |
| 0 | 1.74052056 | 0.262 | 0.007 | 0 | 9 | IGF2 |
| 0 | 1.64641911 | 0.48 | 0.01 | 0 | 9 | TOP2A |
| 0 | 1.16139261 | 0.439 | 0.023 | 0 | 9 | NUSAP1 |
| 0 | 1.09210376 | 0.398 | 0.004 | 0 | 9 | HMMR |
| 0 | 1.08092226 | 0.385 | 0.003 | 0 | 9 | ASPM |
| 0 | 0.96260015 | 0.416 | 0.009 | 0 | 9 | SYNE2 |

| | | | | | | |
|---|---|---|---|---|---|---|
| 0 | 0.91367732 | 0.416 | 0.004 | 0 | 9 | TPX2 |
| 0 | 0.89591572 | 0.376 | 0.005 | 0 | 9 | PCLAF |
| 0 | 0.85210815 | 0.344 | 0.002 | 0 | 9 | UBE2C |
| 0 | 0.80088763 | 0.38 | 0.002 | 0 | 9 | GTSE1 |
| 0 | 0.75410484 | 0.303 | 0.003 | 0 | 9 | TYMS |
| 0 | 0.67535151 | 0.326 | 0.003 | 0 | 9 | ANLN |
| 0 | 0.6657456 | 0.344 | 0.006 | 0 | 9 | KNL1 |
| 0 | 0.66411515 | 0.339 | 0.002 | 0 | 9 | BIRC5 |
| 0 | 0.63901149 | 0.312 | 0.01 | 0 | 9 | CDK1 |
| 0 | 0.62713482 | 0.303 | 0.005 | 0 | 9 | KIF11 |
| 0 | 0.61535241 | 0.339 | 0.004 | 0 | 9 | CEP55 |
| 0 | 0.61398895 | 0.262 | 0.002 | 0 | 9 | HIST1H3B |
| 0 | 0.59707327 | 0.29 | 0.002 | 0 | 9 | AURKB |
| 0 | 0.55652471 | 0.303 | 0.006 | 0 | 9 | CDKN3 |
| 0 | 0.53562283 | 0.308 | 0.01 | 0 | 9 | CENPK |
| 0 | 0.53506639 | 0.29 | 0.005 | 0 | 9 | TK1 |
| 0 | 0.53090557 | 0.285 | 0.005 | 0 | 9 | CCNB2 |
| 0 | 0.52828674 | 0.253 | 0.001 | 0 | 9 | DLGAP5 |
| 0 | 0.49448799 | 0.271 | 0.002 | 0 | 9 | CENPM |
| 0 | 0.47941732 | 0.253 | 0.003 | 0 | 9 | CKAP2L |
| 0 | 0.47408291 | 0.303 | 0.009 | 0 | 9 | ZWINT |
| 0 | 0.45312385 | 0.271 | 0.001 | 0 | 9 | NCAPG |
| 0 | 0.44618046 | 0.29 | 0.003 | 0 | 9 | NUF2 |
| 0 | 0.43778531 | 0.253 | 0.002 | 0 | 9 | NDC80 |
| 0 | 0.43284916 | 0.258 | 0.002 | 0 | 9 | CCNA2 |
| 0 | 0.43258234 | 0.262 | 0.005 | 0 | 9 | KIFC1 |
| 2.43E-296 | 0.6147473 | 0.267 | 0.01 | 5.89E-292 | 9 | CCNB1 |
| 1.70E-279 | 0.62054764 | 0.335 | 0.017 | 4.11E-275 | 9 | SGO2 |
| 9.65E-260 | 0.75551369 | 0.353 | 0.02 | 2.34E-255 | 9 | CENPE |
| 1.08E-259 | 0.5033152 | 0.258 | 0.01 | 2.61E-255 | 9 | CLSPN |
| 8.26E-226 | 0.74935179 | 0.403 | 0.03 | 2.00E-221 | 9 | PRC1 |
| 6.81E-220 | 4.39567183 | 0.253 | 0.012 | 1.65E-215 | 9 | IGFBP3 |
| 2.38E-200 | 0.62520967 | 0.271 | 0.015 | 5.77E-196 | 9 | HELLS |
| 7.12E-193 | 0.83030207 | 0.525 | 0.058 | 1.73E-188 | 9 | CKS1B |
| 1.05E-192 | 1.09178681 | 0.407 | 0.036 | 2.54E-188 | 9 | PTTG1 |
| 2.85E-146 | 0.44369976 | 0.308 | 0.026 | 6.90E-142 | 9 | NCAPD2 |
| 5.93E-128 | 2.65944483 | 0.317 | 0.032 | 1.44E-123 | 9 | H19 |
| 1.86E-114 | 0.41491613 | 0.262 | 0.024 | 4.50E-110 | 9 | TMEM106C |
| 2.11E-114 | 0.63354811 | 0.367 | 0.046 | 5.12E-110 | 9 | CDKN2C |
| 8.76E-111 | 0.4318402 | 0.29 | 0.03 | 2.12E-106 | 9 | RAD51AP1 |
| 3.76E-99 | 0.59816324 | 0.308 | 0.037 | 9.12E-95 | 9 | SPTBN1 |
| 4.71E-93 | 0.51811775 | 0.303 | 0.038 | 1.14E-88 | 9 | ATAD2 |
| 4.84E-93 | 0.64965182 | 0.403 | 0.066 | 1.17E-88 | 9 | SMC2 |

| | | | | | | |
|---|---|---|---|---|---|---|
| 7.64E-92 | 0.74401144 | 0.452 | 0.083 | 1.85E-87 | 9 | H2AFX |
| 1.25E-90 | 0.75792081 | 0.425 | 0.075 | 3.04E-86 | 9 | CKAP2 |
| 4.38E-88 | 1.97667024 | 0.638 | 0.194 | 1.06E-83 | 9 | STMN1 |
| 6.14E-85 | 0.84339935 | 0.407 | 0.073 | 1.49E-80 | 9 | PCNA |
| 3.98E-84 | 0.44406309 | 0.262 | 0.032 | 9.64E-80 | 9 | DHFR |
| 4.43E-75 | 0.55777156 | 0.29 | 0.042 | 1.07E-70 | 9 | HMGB3 |
| 8.74E-75 | 0.40038681 | 0.344 | 0.057 | 2.12E-70 | 9 | NSD2 |
| 1.53E-74 | 1.06607345 | 0.507 | 0.128 | 3.71E-70 | 9 | SMC4 |
| 2.10E-74 | 0.41494906 | 0.258 | 0.034 | 5.10E-70 | 9 | ATAD5 |
| 5.71E-74 | 0.7539709 | 0.425 | 0.089 | 1.38E-69 | 9 | CARHSP1 |
| 7.91E-74 | 0.46517561 | 0.357 | 0.062 | 1.92E-69 | 9 | SKA2 |
| 3.13E-73 | 1.44872993 | 0.824 | 0.398 | 7.59E-69 | 9 | NUCKS1 |
| 4.04E-71 | 0.35924635 | 0.281 | 0.041 | 9.78E-67 | 9 | CDC25B |
| 9.42E-70 | 0.78859933 | 0.335 | 0.059 | 2.28E-65 | 9 | HIST1H1B |
| 4.78E-66 | 0.44072756 | 0.348 | 0.065 | 1.16E-61 | 9 | DTYMK |
| 3.73E-64 | 0.37589836 | 0.271 | 0.042 | 9.04E-60 | 9 | PRR11 |
| 1.96E-63 | 0.43792625 | 0.253 | 0.038 | 4.75E-59 | 9 | CENPU |
| 2.31E-63 | 0.58436474 | 0.367 | 0.074 | 5.60E-59 | 9 | SERPINH1 |
| 5.24E-63 | 0.71230203 | 0.299 | 0.051 | 1.27E-58 | 9 | HSPA2 |
| 6.84E-60 | 0.32322714 | 0.267 | 0.043 | 1.66E-55 | 9 | GMNN |
| 2.44E-58 | 1.58705615 | 0.747 | 0.371 | 5.92E-54 | 9 | HMGB2 |
| 2.91E-57 | 0.94262293 | 0.276 | 0.049 | 7.06E-53 | 9 | CCND2 |
| 4.33E-57 | 1.65533386 | 0.679 | 0.285 | 1.05E-52 | 9 | HIST1H4C |
| 3.83E-55 | 0.39566926 | 0.303 | 0.058 | 9.28E-51 | 9 | CKAP5 |
| 1.37E-54 | 0.78181418 | 0.548 | 0.181 | 3.31E-50 | 9 | TMPO |
| 1.53E-54 | 0.37972575 | 0.344 | 0.072 | 3.70E-50 | 9 | BRD8 |
| 6.48E-52 | 0.66607351 | 0.267 | 0.049 | 1.57E-47 | 9 | EDNRB |
| 2.15E-51 | 0.45324106 | 0.321 | 0.068 | 5.22E-47 | 9 | LMNB1 |
| 5.05E-51 | 0.46804095 | 0.253 | 0.045 | 1.22E-46 | 9 | CFL2 |
| 1.45E-50 | 0.59247612 | 0.367 | 0.088 | 3.51E-46 | 9 | KIF20B |
| 2.68E-50 | 1.50535445 | 0.801 | 0.501 | 6.50E-46 | 9 | H2AFZ |
| 5.01E-50 | 0.54962343 | 0.462 | 0.132 | 1.22E-45 | 9 | PSIP1 |
| 2.15E-49 | 1.07466266 | 0.258 | 0.049 | 5.21E-45 | 9 | PRLR |
| 1.99E-48 | 1.20211782 | 0.968 | 0.926 | 4.81E-44 | 9 | HMGB1 |
| 1.23E-47 | 0.99577949 | 0.303 | 0.066 | 2.98E-43 | 9 | GPX3 |
| 1.42E-47 | 0.34728906 | 0.258 | 0.048 | 3.44E-43 | 9 | NCAPD3 |
| 1.81E-47 | 0.45126301 | 0.38 | 0.096 | 4.40E-43 | 9 | CBX5 |
| 4.41E-47 | 0.32889295 | 0.29 | 0.059 | 1.07E-42 | 9 | PHF19 |
| 1.23E-46 | 0.97912351 | 0.548 | 0.185 | 2.98E-42 | 9 | ENPP2 |
| 1.73E-46 | 0.80106436 | 0.683 | 0.296 | 4.18E-42 | 9 | RAD21 |
| 8.44E-46 | 0.66757679 | 0.548 | 0.194 | 2.05E-41 | 9 | ANP32E |
| 6.93E-45 | 0.30036301 | 0.253 | 0.048 | 1.68E-40 | 9 | POGLUT3 |
| 1.04E-44 | 0.90728616 | 0.276 | 0.059 | 2.51E-40 | 9 | MBNL3 |

| | | | | | | |
|---|---|---|---|---|---|---|
| 3.89E-44 | 0.44843591 | 0.308 | 0.07 | 9.43E-40 | 9 | ETS1 |
| 5.67E-44 | 0.60976788 | 0.398 | 0.114 | 1.38E-39 | 9 | TACC3 |
| 1.41E-43 | 0.34212619 | 0.281 | 0.06 | 3.41E-39 | 9 | DNAJC9 |
| 1.76E-42 | 0.32543907 | 0.271 | 0.057 | 4.27E-38 | 9 | CEP295 |
| 1.24E-41 | 0.36615555 | 0.308 | 0.072 | 3.00E-37 | 9 | ALDH7A1 |
| 6.78E-41 | 0.55435332 | 0.448 | 0.144 | 1.64E-36 | 9 | SMC3 |
| 3.28E-40 | 0.70310571 | 0.271 | 0.062 | 7.94E-36 | 9 | PALLD |
| 5.41E-40 | 0.60132099 | 0.403 | 0.119 | 1.31E-35 | 9 | NUCB2 |
| 2.10E-39 | 0.41853572 | 0.326 | 0.084 | 5.08E-35 | 9 | PGRMC2 |
| 4.19E-39 | 0.88620841 | 0.538 | 0.219 | 1.02E-34 | 9 | DUT |
| 5.32E-39 | 0.60034778 | 0.335 | 0.089 | 1.29E-34 | 9 | BRCA2 |
| 2.97E-38 | 0.26611472 | 0.271 | 0.061 | 7.19E-34 | 9 | TMEM11 |
| 9.14E-38 | 0.75604056 | 0.661 | 0.315 | 2.22E-33 | 9 | NASP |
| 1.10E-37 | 0.42892405 | 0.258 | 0.057 | 2.66E-33 | 9 | HIST1H2AG |
| 2.02E-37 | 0.49335359 | 0.267 | 0.061 | 4.90E-33 | 9 | HIST1H2BN |
| 3.09E-37 | 0.27608943 | 0.253 | 0.055 | 7.50E-33 | 9 | MYO1C |
| 1.03E-36 | 0.69079806 | 0.507 | 0.184 | 2.50E-32 | 9 | RPS4Y1 |
| 1.36E-36 | 0.87277436 | 0.774 | 0.447 | 3.29E-32 | 9 | SEPTIN7 |
| 1.69E-36 | 0.8648964 | 0.502 | 0.186 | 4.09E-32 | 9 | HIST1H1D |
| 2.29E-36 | 0.60004104 | 0.389 | 0.117 | 5.55E-32 | 9 | RND3 |
| 3.13E-36 | 1.22994493 | 0.914 | 0.748 | 7.60E-32 | 9 | HMGN2 |
| 7.11E-36 | 0.34275019 | 0.362 | 0.104 | 1.72E-31 | 9 | NME4 |
| 1.63E-35 | 0.56581421 | 0.339 | 0.097 | 3.94E-31 | 9 | RHOBTB3 |
| 1.68E-35 | 0.4255099 | 0.376 | 0.114 | 4.07E-31 | 9 | DCK |
| 2.09E-35 | 0.30437369 | 0.262 | 0.061 | 5.06E-31 | 9 | KIF22 |
| 2.23E-35 | 0.57370515 | 0.606 | 0.261 | 5.40E-31 | 9 | TPR |
| 3.86E-35 | 0.34370264 | 0.258 | 0.06 | 9.37E-31 | 9 | S100A13 |
| 6.94E-35 | 0.43404569 | 0.416 | 0.137 | 1.68E-30 | 9 | BUB3 |
| 1.83E-34 | 0.50381 | 0.407 | 0.135 | 4.43E-30 | 9 | RPA3 |
| 3.44E-33 | 0.66487138 | 0.348 | 0.107 | 8.33E-29 | 9 | HIST1H3D |
| 4.82E-33 | 0.29293882 | 0.271 | 0.068 | 1.17E-28 | 9 | ATN1 |
| 7.26E-33 | 0.38167111 | 0.29 | 0.077 | 1.76E-28 | 9 | ANKRD36C |
| 8.98E-33 | 0.80260692 | 0.339 | 0.1 | 2.18E-28 | 9 | FCGBP |
| 1.21E-32 | 1.46419343 | 0.887 | 0.695 | 2.94E-28 | 9 | TUBA1B |
| 2.05E-32 | 0.83892186 | 0.959 | 0.816 | 4.96E-28 | 9 | CALM2 |
| 3.79E-32 | 0.78844302 | 0.566 | 0.266 | 9.18E-28 | 9 | UBE2S |
| 4.04E-32 | 0.50814616 | 0.566 | 0.243 | 9.80E-28 | 9 | MRPL51 |
| 4.37E-32 | 0.72313343 | 0.715 | 0.398 | 1.06E-27 | 9 | HP1BP3 |
| 1.06E-31 | 0.40726373 | 0.403 | 0.136 | 2.57E-27 | 9 | RBBP7 |
| 1.25E-31 | 0.82395328 | 0.606 | 0.302 | 3.03E-27 | 9 | PTMS |
| 1.25E-31 | 0.86692967 | 0.792 | 0.529 | 3.04E-27 | 9 | DEK |
| 2.96E-31 | 0.38470734 | 0.398 | 0.132 | 7.18E-27 | 9 | SEPTIN11 |
| 4.26E-31 | 0.27725158 | 0.276 | 0.072 | 1.03E-26 | 9 | SAE1 |

| | | | | | | |
|---|---|---|---|---|---|---|
| 1.23E-30 | 0.70886453 | 0.733 | 0.463 | 2.97E-26 | 9 | MZT2B |
| 1.45E-30 | 2.06501968 | 0.398 | 0.15 | 3.51E-26 | 9 | PHACTR2 |
| 1.49E-30 | 0.35834669 | 0.339 | 0.104 | 3.62E-26 | 9 | KIF2A |
| 1.91E-30 | 0.32246673 | 0.312 | 0.09 | 4.62E-26 | 9 | KIAA0586 |
| 1.14E-29 | 0.39633138 | 0.389 | 0.134 | 2.77E-25 | 9 | CBX1 |
| 1.25E-29 | 0.3368372 | 0.344 | 0.108 | 3.04E-25 | 9 | MGME1 |
| 1.45E-29 | 0.52676054 | 0.462 | 0.184 | 3.52E-25 | 9 | CNTRL |
| 2.13E-29 | 0.33459073 | 0.262 | 0.07 | 5.16E-25 | 9 | CCDC18 |
| 2.99E-29 | 1.3049233 | 0.697 | 0.443 | 7.26E-25 | 9 | TUBB |
| 3.60E-29 | 0.2702053 | 0.267 | 0.071 | 8.72E-25 | 9 | PCNT |
| 5.33E-29 | 0.7205927 | 0.674 | 0.403 | 1.29E-24 | 9 | H2AFV |
| 6.24E-29 | 0.68943944 | 0.448 | 0.187 | 1.51E-24 | 9 | FAM111A |
| 8.31E-29 | 0.40484521 | 0.421 | 0.155 | 2.02E-24 | 9 | MAZ |
| 1.13E-28 | 0.38866543 | 0.362 | 0.122 | 2.74E-24 | 9 | ARPC5L |
| 2.09E-28 | 0.30582683 | 0.891 | 0.445 | 5.07E-24 | 9 | SELENOP |
| 6.03E-28 | 0.68149536 | 0.421 | 0.166 | 1.46E-23 | 9 | ARID5B |
| 1.27E-27 | 0.27624934 | 0.285 | 0.082 | 3.09E-23 | 9 | CDK4 |
| 1.73E-27 | 0.61367915 | 0.71 | 0.402 | 4.20E-23 | 9 | RAN |
| 2.62E-27 | 0.29464678 | 0.326 | 0.102 | 6.36E-23 | 9 | IGF2BP3 |
| 3.02E-27 | 0.53326377 | 0.624 | 0.304 | 7.32E-23 | 9 | PRPF4B |
| 4.63E-27 | 0.38672317 | 0.407 | 0.15 | 1.12E-22 | 9 | NAP1L4 |
| 7.13E-27 | 0.44066185 | 0.498 | 0.21 | 1.73E-22 | 9 | KMT2A |
| 8.25E-27 | 0.32384966 | 0.299 | 0.092 | 2.00E-22 | 9 | MZT1 |
| 1.11E-26 | 1.18756331 | 0.33 | 0.117 | 2.68E-22 | 9 | PLAGL1 |
| 1.51E-26 | 0.68302052 | 0.529 | 0.245 | 3.66E-22 | 9 | CKS2 |
| 2.34E-26 | 0.29456955 | 0.303 | 0.093 | 5.67E-22 | 9 | SYNE1 |
| 2.35E-26 | 0.36159697 | 0.362 | 0.127 | 5.70E-22 | 9 | ADH5 |
| 3.28E-26 | 0.35649166 | 0.362 | 0.125 | 7.96E-22 | 9 | DAAM1 |
| 4.17E-26 | 0.57703392 | 0.679 | 0.354 | 1.01E-21 | 9 | GOLGA4 |
| 4.32E-26 | 0.46496079 | 0.403 | 0.157 | 1.05E-21 | 9 | PARP1 |
| 4.50E-26 | 0.2868214 | 0.262 | 0.075 | 1.09E-21 | 9 | CDK5RAP2 |
| 6.63E-26 | 0.36362027 | 0.425 | 0.164 | 1.61E-21 | 9 | NOP56 |
| 1.01E-25 | 0.50513481 | 0.584 | 0.289 | 2.46E-21 | 9 | ERH |
| 1.16E-25 | 0.37244272 | 0.452 | 0.179 | 2.82E-21 | 9 | UPF2 |
| 1.17E-25 | 0.2540767 | 0.253 | 0.071 | 2.84E-21 | 9 | RPP25 |
| 1.32E-25 | 0.78005245 | 0.624 | 0.308 | 3.21E-21 | 9 | PMP22 |
| 1.40E-25 | 0.44002522 | 0.502 | 0.218 | 3.38E-21 | 9 | CALU |
| 2.15E-25 | 0.30340022 | 0.267 | 0.079 | 5.21E-21 | 9 | RCN2 |
| 3.49E-25 | 1.825676 | 0.348 | 0.135 | 8.47E-21 | 9 | CDKN1C |
| 3.56E-25 | 0.29908955 | 0.262 | 0.076 | 8.62E-21 | 9 | ITM2C |
| 4.67E-25 | 0.46618783 | 0.597 | 0.297 | 1.13E-20 | 9 | HNRNPR |
| 5.13E-25 | 0.34047509 | 0.9 | 0.513 | 1.24E-20 | 9 | RNASE1 |
| 6.41E-25 | 0.29003506 | 0.271 | 0.082 | 1.55E-20 | 9 | USP1 |

| | | | | | | |
|---|---|---|---|---|---|---|
| 6.89E-25 | 0.26263693 | 0.267 | 0.079 | 1.67E-20 | 9 | MFAP1 |
| 7.47E-25 | 0.34064915 | 0.407 | 0.155 | 1.81E-20 | 9 | SUPT16H |
| 8.43E-25 | 0.25993067 | 0.253 | 0.073 | 2.04E-20 | 9 | CEP164 |
| 8.53E-25 | 0.484398 | 0.575 | 0.283 | 2.07E-20 | 9 | PCM1 |
| 9.59E-25 | 0.50688565 | 0.624 | 0.321 | 2.33E-20 | 9 | TMED9 |
| 1.05E-24 | 0.60971624 | 0.81 | 0.538 | 2.54E-20 | 9 | SET |
| 1.15E-24 | 0.30547504 | 0.326 | 0.111 | 2.79E-20 | 9 | NUDCD2 |
| 1.31E-24 | 0.40206208 | 0.339 | 0.12 | 3.17E-20 | 9 | CCDC14 |
| 1.37E-24 | 0.40739576 | 0.466 | 0.198 | 3.33E-20 | 9 | MESD |
| 1.50E-24 | 0.43469226 | 0.475 | 0.207 | 3.63E-20 | 9 | NSD3 |
| 2.27E-24 | 0.59119609 | 0.724 | 0.446 | 5.50E-20 | 9 | ANP32B |
| 3.17E-24 | 0.27452672 | 0.271 | 0.083 | 7.69E-20 | 9 | EMC2 |
| 4.75E-24 | 0.28717982 | 0.312 | 0.104 | 1.15E-19 | 9 | EXOSC8 |
| 6.86E-24 | 0.65606285 | 0.977 | 0.928 | 1.66E-19 | 9 | UBB |
| 7.89E-24 | 0.39537382 | 0.376 | 0.146 | 1.91E-19 | 9 | MZT2A |
| 8.94E-24 | 0.53048549 | 0.778 | 0.507 | 2.17E-19 | 9 | HMGN1 |
| 1.03E-23 | 0.32198158 | 0.29 | 0.095 | 2.49E-19 | 9 | WASL |
| 1.04E-23 | 0.39462374 | 0.398 | 0.156 | 2.52E-19 | 9 | EPB41L2 |
| 1.27E-23 | 0.29118222 | 0.317 | 0.108 | 3.09E-19 | 9 | TP53I13 |
| 2.53E-23 | 0.25141379 | 0.267 | 0.082 | 6.14E-19 | 9 | RANGAP1 |
| 3.00E-23 | 0.30327714 | 0.348 | 0.126 | 7.27E-19 | 9 | PSMB5 |
| 3.31E-23 | 0.45368802 | 0.516 | 0.246 | 8.03E-19 | 9 | SNRPD1 |
| 4.34E-23 | 0.38310597 | 0.367 | 0.141 | 1.05E-18 | 9 | MRPS6 |
| 4.58E-23 | 0.33666713 | 0.489 | 0.205 | 1.11E-18 | 9 | ARHGAP5 |
| 4.96E-23 | 0.47031807 | 0.688 | 0.315 | 1.20E-18 | 9 | LYVE1 |
| 7.23E-23 | 0.72286646 | 1 | 0.958 | 1.75E-18 | 9 | JUN |
| 9.42E-23 | 0.38725289 | 0.525 | 0.228 | 2.28E-18 | 9 | FSCN1 |
| 1.02E-22 | 0.34791809 | 0.326 | 0.115 | 2.48E-18 | 9 | HERC5 |
| 2.14E-22 | 0.42103422 | 0.321 | 0.115 | 5.19E-18 | 9 | LTN1 |
| 3.08E-22 | 0.29468669 | 0.367 | 0.139 | 7.46E-18 | 9 | TFAM |
| 4.57E-22 | 0.25962969 | 0.299 | 0.101 | 1.11E-17 | 9 | SUDS3 |
| 5.14E-22 | 0.31412799 | 0.357 | 0.135 | 1.25E-17 | 9 | ERLEC1 |
| 7.35E-22 | 0.51736647 | 0.674 | 0.331 | 1.78E-17 | 9 | MRC1 |
| 7.46E-22 | 0.3997984 | 0.389 | 0.159 | 1.81E-17 | 9 | PRDX2 |
| 7.56E-22 | 0.47257801 | 0.606 | 0.329 | 1.83E-17 | 9 | NDUFV2 |
| 8.19E-22 | 0.54557942 | 0.67 | 0.393 | 1.99E-17 | 9 | PNN |
| 8.46E-22 | 0.3119033 | 0.303 | 0.106 | 2.05E-17 | 9 | CFAP97 |
| 9.62E-22 | 0.71918124 | 0.683 | 0.446 | 2.33E-17 | 9 | TMEM59 |
| 9.65E-22 | 0.32846781 | 0.448 | 0.193 | 2.34E-17 | 9 | LUC7L2 |
| 1.11E-21 | 0.35042155 | 0.376 | 0.15 | 2.68E-17 | 9 | RFC1 |
| 1.20E-21 | 0.52288664 | 0.855 | 0.614 | 2.92E-17 | 9 | PTGES3 |
| 1.45E-21 | 0.64446499 | 0.38 | 0.161 | 3.52E-17 | 9 | KPNA2 |
| 1.46E-21 | 0.26047522 | 0.258 | 0.082 | 3.54E-17 | 9 | LTV1 |

| | | | | | | |
|---|---|---|---|---|---|---|
| 1.71E-21 | 0.47922576 | 0.796 | 0.427 | 4.15E-17 | 9 | MAF |
| 1.79E-21 | 0.4571954 | 0.516 | 0.233 | 4.33E-17 | 9 | COLEC12 |
| 2.09E-21 | 0.34706852 | 0.262 | 0.087 | 5.06E-17 | 9 | MAGED2 |
| 2.58E-21 | 0.27012906 | 0.308 | 0.108 | 6.26E-17 | 9 | GOPC |
| 2.80E-21 | 0.47556685 | 0.652 | 0.37 | 6.79E-17 | 9 | HNRNPH3 |
| 3.14E-21 | 0.31553591 | 0.326 | 0.118 | 7.61E-17 | 9 | MTUS1 |
| 3.94E-21 | 0.33250496 | 0.299 | 0.107 | 9.56E-17 | 9 | SLC1A5 |
| 4.73E-21 | 0.33217343 | 0.407 | 0.171 | 1.15E-16 | 9 | FKBP3 |
| 4.74E-21 | 0.46507774 | 0.52 | 0.26 | 1.15E-16 | 9 | RTF1 |
| 4.91E-21 | 0.46483843 | 0.466 | 0.205 | 1.19E-16 | 9 | HPGDS |
| 5.20E-21 | 0.51654585 | 0.561 | 0.274 | 1.26E-16 | 9 | CD59 |
| 5.94E-21 | 0.27948863 | 0.276 | 0.092 | 1.44E-16 | 9 | CDKN2D |
| 7.90E-21 | 0.32174012 | 0.253 | 0.082 | 1.92E-16 | 9 | UACA |
| 8.49E-21 | 0.47504591 | 0.466 | 0.225 | 2.06E-16 | 9 | RANBP1 |
| 9.63E-21 | 0.32338269 | 0.385 | 0.157 | 2.34E-16 | 9 | KRR1 |
| 1.20E-20 | 0.25322028 | 0.258 | 0.083 | 2.91E-16 | 9 | AHI1 |
| 1.23E-20 | 0.31771906 | 0.258 | 0.086 | 2.99E-16 | 9 | PXDC1 |
| 1.44E-20 | 0.42481807 | 0.611 | 0.327 | 3.49E-16 | 9 | PPIG |
| 1.50E-20 | 0.26372613 | 0.317 | 0.116 | 3.62E-16 | 9 | ARL1 |
| 1.54E-20 | 0.28632429 | 0.29 | 0.101 | 3.73E-16 | 9 | VGLL4 |
| 3.35E-20 | 0.28458994 | 0.864 | 0.511 | 8.12E-16 | 9 | DAB2 |
| 3.68E-20 | 0.31328123 | 0.38 | 0.158 | 8.92E-16 | 9 | MED10 |
| 3.88E-20 | 0.2721939 | 0.29 | 0.102 | 9.40E-16 | 9 | ZNF22 |
| 4.39E-20 | 0.32811869 | 0.303 | 0.112 | 1.07E-15 | 9 | ZDHHC12 |
| 4.76E-20 | 0.40125289 | 0.439 | 0.2 | 1.15E-15 | 9 | BDP1 |
| 4.90E-20 | 0.2916271 | 0.335 | 0.128 | 1.19E-15 | 9 | ITGB1BP1 |
| 5.48E-20 | 0.39376751 | 0.511 | 0.249 | 1.33E-15 | 9 | TECR |
| 5.90E-20 | 0.41280971 | 0.575 | 0.297 | 1.43E-15 | 9 | LUC7L3 |
| 6.19E-20 | 0.27895165 | 0.398 | 0.164 | 1.50E-15 | 9 | RSBN1L |
| 6.36E-20 | 0.51710478 | 0.529 | 0.279 | 1.54E-15 | 9 | MIS18BP1 |
| 6.98E-20 | 0.25552343 | 0.308 | 0.113 | 1.69E-15 | 9 | MRPL32 |
| 1.18E-19 | 0.42690133 | 0.543 | 0.28 | 2.86E-15 | 9 | EIF5B |
| 1.34E-19 | 0.41170856 | 0.493 | 0.241 | 3.25E-15 | 9 | TAF7 |
| 1.47E-19 | 0.57063174 | 0.471 | 0.218 | 3.57E-15 | 9 | TPM1 |
| 1.74E-19 | 0.47436322 | 0.452 | 0.22 | 4.23E-15 | 9 | LBR |
| 1.79E-19 | 0.57705118 | 0.566 | 0.317 | 4.33E-15 | 9 | CACYBP |
| 1.96E-19 | 0.28591174 | 0.339 | 0.133 | 4.74E-15 | 9 | KMT5A |
| 2.68E-19 | 0.46433114 | 0.493 | 0.236 | 6.49E-15 | 9 | EPS8 |
| 3.15E-19 | 0.28423009 | 0.294 | 0.108 | 7.65E-15 | 9 | ARPC1A |
| 4.28E-19 | 0.25272446 | 0.29 | 0.105 | 1.04E-14 | 9 | MRPL22 |
| 4.80E-19 | 0.49255211 | 0.928 | 0.763 | 1.16E-14 | 9 | HSP90B1 |
| 5.65E-19 | 1.04858053 | 0.697 | 0.52 | 1.37E-14 | 9 | ARL6IP1 |
| 5.71E-19 | 0.32530994 | 0.443 | 0.204 | 1.38E-14 | 9 | PDAP1 |

| | | | | | | |
|---|---|---|---|---|---|---|
| 6.57E-19 | 0.32873477 | 0.326 | 0.127 | 1.59E-14 | 9 | TCEAL3 |
| 7.10E-19 | 0.4056969 | 0.548 | 0.283 | 1.72E-14 | 9 | U2SURP |
| 8.20E-19 | 0.32470695 | 0.367 | 0.152 | 1.99E-14 | 9 | KATNBL1 |
| 8.54E-19 | 0.81383626 | 0.389 | 0.179 | 2.07E-14 | 9 | KCNQ1OT1 |
| 1.14E-18 | 0.38937002 | 0.48 | 0.233 | 2.78E-14 | 9 | SPATS2L |
| 1.19E-18 | 0.35182195 | 0.389 | 0.168 | 2.89E-14 | 9 | PRKDC |
| 1.36E-18 | 0.33207066 | 0.425 | 0.194 | 3.30E-14 | 9 | SYPL1 |
| 1.44E-18 | 0.4187556 | 0.615 | 0.352 | 3.50E-14 | 9 | PNRC2 |
| 1.55E-18 | 0.32509347 | 0.348 | 0.142 | 3.75E-14 | 9 | FGFR1 |
| 1.58E-18 | 0.304053 | 0.498 | 0.22 | 3.83E-14 | 9 | EGFL7 |
| 1.63E-18 | 0.29278737 | 0.321 | 0.126 | 3.96E-14 | 9 | PGRMC1 |
| 1.78E-18 | 0.29713228 | 0.321 | 0.127 | 4.31E-14 | 9 | ESF1 |
| 1.93E-18 | 0.74482021 | 0.62 | 0.403 | 4.69E-14 | 9 | TUBB4B |
| 2.13E-18 | 0.48083652 | 1 | 1 | 5.16E-14 | 9 | MT-CYB |
| 2.54E-18 | 0.35081009 | 0.389 | 0.172 | 6.15E-14 | 9 | DNAJC21 |
| 3.07E-18 | 0.44053133 | 0.552 | 0.307 | 7.45E-14 | 9 | MRFAP1 |
| 3.56E-18 | 0.30490016 | 0.321 | 0.127 | 8.62E-14 | 9 | SAP30 |
| 3.71E-18 | 0.35678274 | 0.434 | 0.201 | 9.00E-14 | 9 | DCTN3 |
| 3.80E-18 | 0.41371825 | 0.457 | 0.226 | 9.21E-14 | 9 | ASPH |
| 3.84E-18 | 0.29488286 | 0.267 | 0.096 | 9.31E-14 | 9 | ADD3 |
| 4.07E-18 | 0.26615372 | 0.262 | 0.093 | 9.86E-14 | 9 | ZC3H14 |
| 5.16E-18 | 0.27001597 | 0.299 | 0.114 | 1.25E-13 | 9 | EI24 |
| 5.75E-18 | 0.50047584 | 0.493 | 0.259 | 1.39E-13 | 9 | PDCD4 |
| 6.56E-18 | 0.30984094 | 0.376 | 0.164 | 1.59E-13 | 9 | CCT7 |
| 8.62E-18 | 0.30948756 | 0.303 | 0.119 | 2.09E-13 | 9 | ERGIC2 |
| 9.67E-18 | 0.47026014 | 0.376 | 0.175 | 2.34E-13 | 9 | CD151 |
| 9.87E-18 | 0.30695492 | 0.416 | 0.191 | 2.39E-13 | 9 | ILF2 |
| 1.20E-17 | 0.33654862 | 0.439 | 0.202 | 2.90E-13 | 9 | BIN1 |
| 1.26E-17 | 0.48008155 | 0.905 | 0.745 | 3.06E-13 | 9 | HNRNPA1 |
| 1.34E-17 | 0.38396131 | 0.516 | 0.269 | 3.25E-13 | 9 | STOM |
| 1.53E-17 | 0.28568411 | 0.294 | 0.113 | 3.71E-13 | 9 | SLF2 |
| 1.55E-17 | 0.28969111 | 0.285 | 0.109 | 3.76E-13 | 9 | CDC123 |
| 2.05E-17 | 0.57976323 | 0.611 | 0.379 | 4.96E-13 | 9 | UBXN4 |
| 3.00E-17 | 0.5336856 | 0.95 | 0.769 | 7.28E-13 | 9 | HSPB1 |
| 4.01E-17 | 0.32405272 | 0.321 | 0.133 | 9.72E-13 | 9 | CTCF |
| 4.04E-17 | 0.28119737 | 0.303 | 0.12 | 9.80E-13 | 9 | TLK1 |
| 4.84E-17 | 0.38663784 | 0.448 | 0.223 | 1.17E-12 | 9 | RDX |
| 6.25E-17 | 0.39607787 | 0.498 | 0.266 | 1.51E-12 | 9 | RAB14 |
| 6.58E-17 | 0.41075339 | 0.457 | 0.228 | 1.59E-12 | 9 | LMO4 |
| 7.03E-17 | 0.27742663 | 0.308 | 0.123 | 1.71E-12 | 9 | C5orf24 |
| 7.11E-17 | 0.34612334 | 0.48 | 0.241 | 1.72E-12 | 9 | CCDC50 |
| 7.67E-17 | 0.26412508 | 0.299 | 0.118 | 1.86E-12 | 9 | CCM2 |
| 8.95E-17 | 0.28340969 | 0.308 | 0.124 | 2.17E-12 | 9 | NUP50 |

| | | | | | | |
|---|---|---|---|---|---|---|
| 9.76E-17 | 0.25862458 | 0.312 | 0.127 | 2.37E-12 | 9 | RASSF1 |
| 1.01E-16 | 0.50311177 | 0.579 | 0.351 | 2.46E-12 | 9 | SRSF10 |
| 1.08E-16 | 0.43320508 | 0.747 | 0.519 | 2.61E-12 | 9 | SERBP1 |
| 1.15E-16 | 0.53790404 | 0.597 | 0.375 | 2.78E-12 | 9 | SOD1 |
| 1.23E-16 | 0.27040991 | 0.308 | 0.124 | 2.99E-12 | 9 | NGRN |
| 1.26E-16 | 0.47237355 | 0.552 | 0.313 | 3.05E-12 | 9 | ISCU |
| 1.32E-16 | 0.32939284 | 0.348 | 0.153 | 3.20E-12 | 9 | SPTAN1 |
| 1.40E-16 | 0.25965121 | 0.344 | 0.145 | 3.39E-12 | 9 | PHF14 |
| 1.87E-16 | 0.34154976 | 0.357 | 0.162 | 4.52E-12 | 9 | ASXL1 |
| 1.93E-16 | 0.38481705 | 0.443 | 0.219 | 4.67E-12 | 9 | SMARCA2 |
| 1.94E-16 | 0.25223531 | 0.299 | 0.119 | 4.70E-12 | 9 | FBXL3 |
| 2.33E-16 | 0.28400771 | 0.344 | 0.148 | 5.65E-12 | 9 | MAP4 |
| 2.34E-16 | 0.26924194 | 0.271 | 0.104 | 5.68E-12 | 9 | ZMYND11 |
| 2.80E-16 | 0.30435046 | 0.335 | 0.143 | 6.80E-12 | 9 | ROCK2 |
| 3.14E-16 | 0.28443664 | 0.326 | 0.137 | 7.61E-12 | 9 | BAZ1B |
| 3.59E-16 | 0.37861244 | 0.457 | 0.239 | 8.71E-12 | 9 | PRDX3 |
| 3.60E-16 | 0.28892252 | 0.339 | 0.147 | 8.73E-12 | 9 | PURA |
| 3.71E-16 | 0.35602503 | 0.475 | 0.25 | 9.00E-12 | 9 | NUDC |
| 3.74E-16 | 0.46543919 | 0.606 | 0.385 | 9.06E-12 | 9 | P4HB |
| 3.80E-16 | 0.43772502 | 0.389 | 0.188 | 9.21E-12 | 9 | DNMT1 |
| 4.19E-16 | 0.30939446 | 0.416 | 0.197 | 1.02E-11 | 9 | ZC3H13 |
| 4.37E-16 | 0.52109157 | 0.76 | 0.571 | 1.06E-11 | 9 | NCL |
| 4.92E-16 | 0.31385141 | 0.308 | 0.13 | 1.19E-11 | 9 | HMG20B |
| 5.14E-16 | 0.53078055 | 0.846 | 0.604 | 1.25E-11 | 9 | BTG2 |
| 5.45E-16 | 0.46053548 | 0.43 | 0.22 | 1.32E-11 | 9 | LDHB |
| 6.14E-16 | 1.97000143 | 0.498 | 0.3 | 1.49E-11 | 9 | IL6ST |
| 6.27E-16 | 0.3068295 | 0.326 | 0.141 | 1.52E-11 | 9 | NUTF2 |
| 6.28E-16 | 0.28471813 | 0.312 | 0.129 | 1.52E-11 | 9 | FARP1 |
| 6.44E-16 | 0.27699733 | 0.33 | 0.141 | 1.56E-11 | 9 | SMIM15 |
| 6.76E-16 | 0.30997562 | 0.403 | 0.191 | 1.64E-11 | 9 | SUZ12 |
| 7.99E-16 | 0.48231118 | 0.995 | 1 | 1.94E-11 | 9 | MT-ATP6 |
| 8.05E-16 | 0.27540126 | 0.267 | 0.103 | 1.95E-11 | 9 | PRKCA |
| 9.65E-16 | 0.27335834 | 0.394 | 0.183 | 2.34E-11 | 9 | ADD1 |
| 9.86E-16 | 0.52399613 | 0.534 | 0.281 | 2.39E-11 | 9 | PDK4 |
| 1.07E-15 | 0.34966424 | 0.452 | 0.23 | 2.59E-11 | 9 | GNG2 |
| 1.08E-15 | 0.33613615 | 0.52 | 0.285 | 2.62E-11 | 9 | RAD23A |
| 1.11E-15 | 0.42512523 | 0.584 | 0.349 | 2.69E-11 | 9 | TMEM14C |
| 1.12E-15 | 0.57112373 | 0.566 | 0.338 | 2.71E-11 | 9 | RRBP1 |
| 1.24E-15 | 0.26434794 | 0.371 | 0.168 | 3.01E-11 | 9 | TRIM28 |
| 2.29E-15 | 0.27585344 | 0.299 | 0.125 | 5.56E-11 | 9 | UBTF |
| 2.41E-15 | 0.46246704 | 0.733 | 0.522 | 5.83E-11 | 9 | SAP18 |
| 2.67E-15 | 0.38063576 | 0.851 | 0.619 | 6.48E-11 | 9 | SON |
| 3.19E-15 | 0.35078691 | 0.308 | 0.133 | 7.75E-11 | 9 | CCNG2 |

| | | | | | | |
|---|---|---|---|---|---|---|
| 4.05E-15 | 0.3767448 | 0.597 | 0.353 | 9.82E-11 | 9 | KIF5B |
| 4.29E-15 | 0.34520182 | 0.52 | 0.267 | 1.04E-10 | 9 | NRP1 |
| 5.07E-15 | 0.2718495 | 0.398 | 0.192 | 1.23E-10 | 9 | LSM14A |
| 5.21E-15 | 0.32472532 | 0.502 | 0.249 | 1.26E-10 | 9 | DDX3Y |
| 6.18E-15 | 0.2661706 | 0.321 | 0.14 | 1.50E-10 | 9 | RECQL |
| 6.56E-15 | 0.36504425 | 0.579 | 0.345 | 1.59E-10 | 9 | NDUFC2 |
| 6.95E-15 | 0.43474639 | 1 | 0.998 | 1.69E-10 | 9 | PTMA |
| 7.13E-15 | 0.27519525 | 0.271 | 0.11 | 1.73E-10 | 9 | REXO2 |
| 7.61E-15 | 0.52881356 | 0.421 | 0.222 | 1.84E-10 | 9 | HIST1H1E |
| 7.78E-15 | 0.2897513 | 0.348 | 0.157 | 1.89E-10 | 9 | SNRNP200 |
| 8.13E-15 | 0.30260209 | 0.326 | 0.146 | 1.97E-10 | 9 | TRIP11 |
| 9.12E-15 | 0.26116965 | 0.376 | 0.175 | 2.21E-10 | 9 | CENPC |
| 9.72E-15 | 0.33747341 | 0.398 | 0.196 | 2.36E-10 | 9 | RSF1 |
| 1.01E-14 | 0.35981742 | 0.443 | 0.224 | 2.44E-10 | 9 | SESN1 |
| 1.02E-14 | 0.26778962 | 0.262 | 0.105 | 2.47E-10 | 9 | UNC50 |
| 1.09E-14 | 0.3426811 | 0.575 | 0.337 | 2.65E-10 | 9 | SF3B6 |
| 1.21E-14 | 0.49186616 | 0.538 | 0.324 | 2.93E-10 | 9 | SBDS |
| 1.26E-14 | 0.79143044 | 0.548 | 0.303 | 3.06E-10 | 9 | CCL2 |
| 1.48E-14 | 0.27287017 | 0.398 | 0.194 | 3.58E-10 | 9 | MPC2 |
| 1.56E-14 | 0.27433624 | 0.312 | 0.138 | 3.78E-10 | 9 | GLRX5 |
| 1.61E-14 | 0.25818262 | 0.321 | 0.141 | 3.90E-10 | 9 | NSL1 |
| 1.79E-14 | 0.29584108 | 0.357 | 0.167 | 4.33E-10 | 9 | OIP5-AS1 |
| 2.13E-14 | 0.27215106 | 0.267 | 0.108 | 5.16E-10 | 9 | TOPORS |
| 2.13E-14 | 0.44602569 | 0.828 | 0.636 | 5.16E-10 | 9 | DYNLL1 |
| 2.19E-14 | 0.34665942 | 0.439 | 0.227 | 5.32E-10 | 9 | CEP350 |
| 2.21E-14 | 0.49464211 | 0.439 | 0.238 | 5.36E-10 | 9 | UGP2 |
| 2.38E-14 | 0.55315418 | 0.516 | 0.291 | 5.77E-10 | 9 | PIK3R1 |
| 2.44E-14 | 0.54635685 | 0.271 | 0.115 | 5.92E-10 | 9 | EPAS1 |
| 2.76E-14 | 0.36756541 | 0.919 | 0.682 | 6.70E-10 | 9 | C1QA |
| 2.77E-14 | 0.30075996 | 0.33 | 0.152 | 6.72E-10 | 9 | SMC5 |
| 2.89E-14 | 0.37213179 | 0.308 | 0.135 | 7.00E-10 | 9 | IGFBP4 |
| 4.10E-14 | 0.26541244 | 0.376 | 0.182 | 9.94E-10 | 9 | EMC7 |
| 4.19E-14 | 0.30154459 | 0.281 | 0.121 | 1.02E-09 | 9 | SUN2 |
| 4.42E-14 | 0.34594846 | 0.484 | 0.267 | 1.07E-09 | 9 | MORF4L2 |
| 4.56E-14 | 0.36047418 | 0.466 | 0.258 | 1.10E-09 | 9 | LSM4 |
| 4.69E-14 | 0.42230651 | 0.597 | 0.374 | 1.14E-09 | 9 | EIF2S2 |
| 5.38E-14 | 0.39652917 | 0.633 | 0.406 | 1.30E-09 | 9 | HNRNPM |
| 5.51E-14 | 0.79935034 | 0.534 | 0.352 | 1.33E-09 | 9 | BSG |
| 5.56E-14 | 0.26897572 | 0.303 | 0.133 | 1.35E-09 | 9 | UBA2 |
| 5.83E-14 | 0.45088699 | 0.71 | 0.46 | 1.41E-09 | 9 | STAB1 |
| 6.12E-14 | 0.47104103 | 0.299 | 0.132 | 1.48E-09 | 9 | HIST1H2BG |
| 6.97E-14 | 0.33269726 | 0.389 | 0.194 | 1.69E-09 | 9 | DIPK2A |
| 7.13E-14 | 0.35613193 | 0.833 | 0.542 | 1.73E-09 | 9 | C1QC |

| | | | | | | |
|---|---|---|---|---|---|---|
| 7.60E-14 | 0.32263282 | 0.353 | 0.17 | 1.84E-09 | 9 | UPF3A |
| 7.76E-14 | 0.25261232 | 0.344 | 0.159 | 1.88E-09 | 9 | BCAP29 |
| 8.60E-14 | 0.34738959 | 0.543 | 0.321 | 2.09E-09 | 9 | TCEA1 |
| 9.01E-14 | 0.25259799 | 0.335 | 0.155 | 2.18E-09 | 9 | GON4L |
| 9.03E-14 | 0.43978174 | 0.873 | 0.611 | 2.19E-09 | 9 | C1QB |
| 9.14E-14 | 0.2694824 | 0.294 | 0.128 | 2.21E-09 | 9 | SLBP |
| 9.69E-14 | 0.36074137 | 0.498 | 0.283 | 2.35E-09 | 9 | BPTF |
| 1.08E-13 | 0.34629977 | 0.339 | 0.161 | 2.62E-09 | 9 | MBNL2 |
| 1.13E-13 | 0.32678721 | 0.385 | 0.193 | 2.73E-09 | 9 | SMC1A |
| 1.21E-13 | 0.26572324 | 0.371 | 0.18 | 2.93E-09 | 9 | OTUD6B-AS1 |
| 1.53E-13 | 0.32534645 | 0.253 | 0.106 | 3.72E-09 | 9 | CYB5A |
| 1.64E-13 | 0.29516042 | 0.357 | 0.17 | 3.97E-09 | 9 | CCDC144A |
| 1.64E-13 | 0.43832815 | 0.543 | 0.338 | 3.97E-09 | 9 | SELENOW |
| 1.82E-13 | 0.28096136 | 0.294 | 0.132 | 4.41E-09 | 9 | TSC22D4 |
| 1.84E-13 | 0.34320588 | 0.615 | 0.39 | 4.47E-09 | 9 | RBMX |
| 1.89E-13 | 0.32413954 | 0.552 | 0.326 | 4.58E-09 | 9 | WASF2 |
| 2.03E-13 | 0.28795633 | 0.339 | 0.16 | 4.91E-09 | 9 | RIF1 |
| 2.19E-13 | 0.29590885 | 0.271 | 0.117 | 5.31E-09 | 9 | PID1 |
| 2.24E-13 | 0.25753359 | 0.353 | 0.169 | 5.42E-09 | 9 | NOP58 |
| 2.26E-13 | 0.35236825 | 0.665 | 0.443 | 5.49E-09 | 9 | TRIR |
| 2.35E-13 | 0.27450142 | 0.656 | 0.369 | 5.69E-09 | 9 | FOLR2 |
| 2.54E-13 | 0.31662472 | 0.357 | 0.178 | 6.15E-09 | 9 | PSMC3 |
| 2.68E-13 | 0.30856767 | 0.421 | 0.221 | 6.50E-09 | 9 | PDCD5 |
| 2.71E-13 | 0.30938365 | 0.398 | 0.205 | 6.56E-09 | 9 | CNPY2 |
| 2.79E-13 | 0.25127672 | 0.326 | 0.151 | 6.76E-09 | 9 | DDX42 |
| 3.49E-13 | 0.32558582 | 0.389 | 0.201 | 8.47E-09 | 9 | TERF2IP |
| 3.52E-13 | 0.4848508 | 0.276 | 0.12 | 8.52E-09 | 9 | ID3 |
| 3.77E-13 | 0.2988817 | 0.371 | 0.186 | 9.14E-09 | 9 | GINM1 |
| 3.81E-13 | 0.35281214 | 0.873 | 0.672 | 9.25E-09 | 9 | RHOB |
| 3.83E-13 | 0.34019457 | 0.403 | 0.21 | 9.29E-09 | 9 | MLEC |
| 3.84E-13 | 0.37596203 | 0.633 | 0.394 | 9.31E-09 | 9 | ANKRD12 |
| 4.26E-13 | 0.29782167 | 0.43 | 0.231 | 1.03E-08 | 9 | PSMG2 |
| 4.54E-13 | 0.34069536 | 0.376 | 0.193 | 1.10E-08 | 9 | LMAN1 |
| 4.57E-13 | 0.36760574 | 0.498 | 0.281 | 1.11E-08 | 9 | MPHOSPH8 |
| 4.64E-13 | 0.30386474 | 0.502 | 0.288 | 1.12E-08 | 9 | NOL7 |
| 5.00E-13 | 0.28921137 | 0.262 | 0.111 | 1.21E-08 | 9 | CEBPG |
| 6.64E-13 | 0.40406161 | 0.466 | 0.266 | 1.61E-08 | 9 | ACTN4 |
| 7.02E-13 | 0.46411241 | 0.71 | 0.511 | 1.70E-08 | 9 | RSRC2 |
| 7.82E-13 | 0.34410983 | 0.706 | 0.477 | 1.89E-08 | 9 | HNRNPD |
| 7.90E-13 | 0.36631254 | 0.525 | 0.317 | 1.92E-08 | 9 | DYNLRB1 |
| 1.15E-12 | 0.25391371 | 0.348 | 0.169 | 2.79E-08 | 9 | PIN1 |
| 1.29E-12 | 0.28489825 | 0.33 | 0.158 | 3.12E-08 | 9 | HSPB11 |
| 1.33E-12 | 0.37750313 | 0.439 | 0.235 | 3.22E-08 | 9 | NFIA |

| | | | | | | |
|---|---|---|---|---|---|---|
| 1.53E-12 | 0.35515184 | 0.57 | 0.364 | 3.71E-08 | 9 | XRCC5 |
| 1.64E-12 | 0.25265771 | 0.276 | 0.123 | 3.96E-08 | 9 | CREBZF |
| 1.89E-12 | 0.27565345 | 0.48 | 0.269 | 4.58E-08 | 9 | THRAP3 |
| 2.02E-12 | 0.30266454 | 0.407 | 0.22 | 4.89E-08 | 9 | SNRPD3 |
| 2.38E-12 | 0.26376701 | 0.281 | 0.125 | 5.78E-08 | 9 | CHD7 |
| 2.40E-12 | 0.28772263 | 0.389 | 0.202 | 5.83E-08 | 9 | SNHG1 |
| 2.61E-12 | 0.26187081 | 0.357 | 0.179 | 6.32E-08 | 9 | HNRNPUL2 |
| 3.09E-12 | 0.27467225 | 0.38 | 0.197 | 7.50E-08 | 9 | SHISA5 |
| 4.23E-12 | 0.27131687 | 0.416 | 0.224 | 1.03E-07 | 9 | ABCF1 |
| 4.55E-12 | 0.3437008 | 0.629 | 0.395 | 1.10E-07 | 9 | SNHG32 |
| 4.72E-12 | 0.25240369 | 0.416 | 0.221 | 1.14E-07 | 9 | POLR2K |
| 4.73E-12 | 0.3371784 | 0.579 | 0.373 | 1.15E-07 | 9 | DNAJC8 |
| 4.86E-12 | 0.28117832 | 0.371 | 0.193 | 1.18E-07 | 9 | COPS6 |
| 4.87E-12 | 0.33243165 | 0.498 | 0.296 | 1.18E-07 | 9 | UBE2I |
| 5.31E-12 | 0.28882621 | 0.543 | 0.312 | 1.29E-07 | 9 | DNAJB4 |
| 5.45E-12 | 0.28211073 | 0.271 | 0.123 | 1.32E-07 | 9 | DDB1 |
| 5.79E-12 | 0.42190541 | 0.353 | 0.18 | 1.40E-07 | 9 | TSC22D1 |
| 6.39E-12 | 0.46435768 | 0.742 | 0.551 | 1.55E-07 | 9 | IFI16 |
| 6.52E-12 | 0.38404853 | 0.697 | 0.502 | 1.58E-07 | 9 | CLTA |
| 7.61E-12 | 0.27422641 | 0.33 | 0.162 | 1.85E-07 | 9 | TUT4 |
| 7.94E-12 | 0.28046837 | 0.33 | 0.164 | 1.93E-07 | 9 | NHLRC3 |
| 8.12E-12 | 0.33565667 | 0.525 | 0.321 | 1.97E-07 | 9 | PEBP1 |
| 8.21E-12 | 0.41610261 | 0.647 | 0.437 | 1.99E-07 | 9 | CALM3 |
| 8.75E-12 | 0.31256982 | 0.412 | 0.226 | 2.12E-07 | 9 | LSM5 |
| 9.33E-12 | 0.2659425 | 0.362 | 0.187 | 2.26E-07 | 9 | PTPRA |
| 9.63E-12 | 0.27990462 | 0.412 | 0.226 | 2.33E-07 | 9 | PSMC5 |
| 1.01E-11 | 0.25851843 | 0.385 | 0.201 | 2.44E-07 | 9 | CCT2 |
| 1.12E-11 | 0.28258105 | 0.407 | 0.221 | 2.72E-07 | 9 | TBL1XR1 |
| 1.19E-11 | 0.26163697 | 0.412 | 0.219 | 2.88E-07 | 9 | ZNF638 |
| 1.20E-11 | 0.42884277 | 0.801 | 0.61 | 2.92E-07 | 9 | HNRNPH1 |
| 1.27E-11 | 0.29677909 | 0.312 | 0.155 | 3.09E-07 | 9 | NFATC2IP |
| 1.50E-11 | 0.26820333 | 0.389 | 0.202 | 3.63E-07 | 9 | CEMIP2 |
| 1.50E-11 | 0.27163496 | 0.367 | 0.191 | 3.65E-07 | 9 | SMARCE1 |
| 1.53E-11 | 0.39261738 | 0.566 | 0.367 | 3.70E-07 | 9 | RHEB |
| 1.54E-11 | 0.28024587 | 0.344 | 0.173 | 3.73E-07 | 9 | GABPB1-AS1 |
| 1.90E-11 | 0.26382442 | 0.353 | 0.178 | 4.59E-07 | 9 | ILF3-DT |
| 1.90E-11 | 0.31371858 | 0.362 | 0.196 | 4.60E-07 | 9 | IDH2 |
| 1.96E-11 | 0.31092759 | 0.398 | 0.217 | 4.75E-07 | 9 | LARP7 |
| 2.45E-11 | 0.60731996 | 0.955 | 0.975 | 5.94E-07 | 9 | RPL23A |
| 2.59E-11 | 0.25521398 | 0.394 | 0.21 | 6.28E-07 | 9 | FUBP1 |
| 2.62E-11 | 0.25038007 | 0.425 | 0.235 | 6.36E-07 | 9 | CCT3 |
| 3.38E-11 | 0.37260184 | 0.674 | 0.483 | 8.20E-07 | 9 | KTN1 |
| 3.71E-11 | 0.33401866 | 0.439 | 0.254 | 8.99E-07 | 9 | ASH1L |

| | | | | | | |
|---|---|---|---|---|---|---|
| 3.72E-11 | 0.40495957 | 0.819 | 0.619 | 9.02E-07 | 9 | SFPQ |
| 3.81E-11 | 0.31555308 | 0.439 | 0.249 | 9.23E-07 | 9 | GOLGB1 |
| 3.86E-11 | 0.65139829 | 0.43 | 0.264 | 9.36E-07 | 9 | DSTN |
| 4.01E-11 | 0.49046307 | 0.33 | 0.162 | 9.72E-07 | 9 | CCL8 |
| 4.02E-11 | 0.27061163 | 0.348 | 0.182 | 9.75E-07 | 9 | CEBPZ |
| 4.24E-11 | 0.29483273 | 0.484 | 0.291 | 1.03E-06 | 9 | PRMT2 |
| 4.25E-11 | 0.37714596 | 0.484 | 0.304 | 1.03E-06 | 9 | TLE5 |
| 4.58E-11 | 0.37427153 | 0.697 | 0.49 | 1.11E-06 | 9 | ATP5IF1 |
| 4.76E-11 | 0.29589478 | 0.534 | 0.325 | 1.15E-06 | 9 | HNRNPAB |
| 5.28E-11 | 0.27091682 | 0.416 | 0.229 | 1.28E-06 | 9 | FGD5-AS1 |
| 6.52E-11 | 0.33913268 | 0.629 | 0.416 | 1.58E-06 | 9 | SNX2 |
| 8.17E-11 | 1.04830166 | 0.339 | 0.184 | 1.98E-06 | 9 | CCL5 |
| 8.30E-11 | 0.25140367 | 0.434 | 0.249 | 2.01E-06 | 9 | PCNP |
| 8.61E-11 | 0.27161116 | 0.457 | 0.271 | 2.09E-06 | 9 | XPO1 |
| 8.65E-11 | 0.31801812 | 0.371 | 0.207 | 2.10E-06 | 9 | DDX39A |
| 9.53E-11 | 0.31143915 | 0.751 | 0.517 | 2.31E-06 | 9 | SNX6 |
| 1.16E-10 | 0.39474621 | 0.633 | 0.428 | 2.82E-06 | 9 | ALOX5AP |
| 1.40E-10 | 0.60188527 | 0.267 | 0.132 | 3.40E-06 | 9 | NUPR1 |
| 1.44E-10 | 0.27678715 | 0.425 | 0.244 | 3.50E-06 | 9 | DHX36 |
| 1.66E-10 | 0.47209277 | 0.733 | 0.57 | 4.03E-06 | 9 | HINT1 |
| 1.88E-10 | 0.31218635 | 0.452 | 0.27 | 4.55E-06 | 9 | POLR2J3 |
| 1.89E-10 | 0.27411827 | 0.588 | 0.367 | 4.58E-06 | 9 | CLTC |
| 1.99E-10 | 0.29597808 | 0.326 | 0.171 | 4.83E-06 | 9 | LGALS3BP |
| 2.00E-10 | 0.28590923 | 0.276 | 0.136 | 4.85E-06 | 9 | EZH2 |
| 2.10E-10 | 0.30447437 | 0.538 | 0.344 | 5.08E-06 | 9 | SSB |
| 2.10E-10 | 0.666572 | 0.484 | 0.341 | 5.10E-06 | 9 | TUBA1C |
| 2.11E-10 | 0.2695962 | 0.471 | 0.282 | 5.12E-06 | 9 | CCT8 |
| 2.14E-10 | 0.25733378 | 0.606 | 0.366 | 5.20E-06 | 9 | VSIG4 |
| 2.36E-10 | 0.32432143 | 0.52 | 0.326 | 5.73E-06 | 9 | BOD1L1 |
| 2.71E-10 | 0.25715763 | 0.412 | 0.236 | 6.57E-06 | 9 | RBBP4 |
| 2.76E-10 | 0.25502312 | 0.367 | 0.196 | 6.68E-06 | 9 | IRF9 |
| 2.84E-10 | 0.27137621 | 0.421 | 0.237 | 6.88E-06 | 9 | NFIC |
| 3.04E-10 | 0.33238072 | 0.462 | 0.281 | 7.38E-06 | 9 | MRPL18 |
| 3.24E-10 | 0.54038469 | 0.914 | 0.796 | 7.85E-06 | 9 | DNAJA1 |
| 3.37E-10 | 0.25694403 | 0.362 | 0.2 | 8.17E-06 | 9 | DDOST |
| 4.36E-10 | 0.25299503 | 0.357 | 0.194 | 1.06E-05 | 9 | AP2B1 |
| 4.54E-10 | 0.41289412 | 0.385 | 0.22 | 1.10E-05 | 9 | HIST2H2AC |
| 4.62E-10 | 0.29138001 | 0.493 | 0.305 | 1.12E-05 | 9 | LSM3 |
| 4.86E-10 | 0.25910791 | 0.389 | 0.218 | 1.18E-05 | 9 | NDUFS4 |
| 5.03E-10 | 0.46914132 | 0.516 | 0.332 | 1.22E-05 | 9 | DDIT3 |
| 5.81E-10 | 0.34858681 | 0.371 | 0.204 | 1.41E-05 | 9 | SLC40A1 |
| 5.99E-10 | 0.34275804 | 0.285 | 0.142 | 1.45E-05 | 9 | CMC1 |
| 6.38E-10 | 0.33163938 | 0.561 | 0.375 | 1.55E-05 | 9 | SNRPG |

| | | | | | | |
|---|---|---|---|---|---|---|
| 6.81E-10 | 0.25918788 | 0.376 | 0.21 | 1.65E-05 | 9 | TSPAN4 |
| 7.59E-10 | 0.30847246 | 0.611 | 0.424 | 1.84E-05 | 9 | C11orf58 |
| 8.23E-10 | 0.34689017 | 0.557 | 0.379 | 2.00E-05 | 9 | RAB2A |
| 8.29E-10 | 0.35164076 | 0.57 | 0.393 | 2.01E-05 | 9 | PDIA6 |
| 8.36E-10 | 0.29027591 | 0.52 | 0.328 | 2.03E-05 | 9 | YY1 |
| 8.49E-10 | 0.27878977 | 0.448 | 0.268 | 2.06E-05 | 9 | CHCHD7 |
| 8.77E-10 | 0.45665206 | 0.579 | 0.404 | 2.13E-05 | 9 | TCP1 |
| 1.13E-09 | 0.52438246 | 0.851 | 0.713 | 2.74E-05 | 9 | SRSF7 |
| 1.16E-09 | 0.40659606 | 0.995 | 0.982 | 2.81E-05 | 9 | FOS |
| 1.30E-09 | 0.27722166 | 0.299 | 0.154 | 3.16E-05 | 9 | KAT6B |
| 1.32E-09 | 0.270272 | 0.344 | 0.19 | 3.20E-05 | 9 | NUDT1 |
| 1.44E-09 | 0.26520605 | 0.52 | 0.325 | 3.48E-05 | 9 | ATRX |
| 1.45E-09 | 0.34044851 | 0.878 | 0.698 | 3.52E-05 | 9 | CALR |
| 1.60E-09 | 0.28679376 | 0.466 | 0.296 | 3.88E-05 | 9 | NUCB1 |
| 1.60E-09 | 0.37291426 | 0.955 | 0.829 | 3.89E-05 | 9 | CALM1 |
| 1.64E-09 | 0.34709978 | 0.846 | 0.753 | 3.97E-05 | 9 | EIF4A2 |
| 1.68E-09 | 0.25325364 | 0.416 | 0.244 | 4.08E-05 | 9 | COX20 |
| 1.70E-09 | 0.39203281 | 0.389 | 0.233 | 4.11E-05 | 9 | TCF4 |
| 2.15E-09 | 0.26390669 | 0.507 | 0.321 | 5.20E-05 | 9 | NDFIP1 |
| 2.38E-09 | 0.27948815 | 0.507 | 0.335 | 5.77E-05 | 9 | KHDRBS1 |
| 2.79E-09 | 0.30401903 | 0.448 | 0.268 | 6.75E-05 | 9 | C2 |
| 2.87E-09 | 0.39467156 | 0.529 | 0.359 | 6.97E-05 | 9 | RBMS1 |
| 2.93E-09 | 0.3603389 | 0.566 | 0.392 | 7.10E-05 | 9 | PTTG1IP |
| 3.41E-09 | 0.25513741 | 0.339 | 0.189 | 8.26E-05 | 9 | PPM1G |
| 3.42E-09 | 0.36672624 | 0.371 | 0.218 | 8.30E-05 | 9 | PCED1B-AS1 |
| 3.81E-09 | 0.29381671 | 0.448 | 0.282 | 9.24E-05 | 9 | SIVA1 |
| 3.95E-09 | 0.45645175 | 0.606 | 0.446 | 9.57E-05 | 9 | PLD3 |
| 4.79E-09 | 0.28537651 | 0.48 | 0.31 | 0.00011619 | 9 | DDX6 |
| 5.37E-09 | 0.35943392 | 0.407 | 0.256 | 0.00013028 | 9 | RAC2 |
| 5.43E-09 | 0.25854934 | 0.33 | 0.185 | 0.00013165 | 9 | BRD7 |
| 6.85E-09 | 0.25437909 | 0.403 | 0.241 | 0.00016609 | 9 | PPP1CC |
| 7.39E-09 | 0.27141748 | 0.67 | 0.47 | 0.00017913 | 9 | SCAF11 |
| 7.52E-09 | 0.28754984 | 0.421 | 0.254 | 0.00018223 | 9 | AKR1B1 |
| 7.62E-09 | 0.25758002 | 0.371 | 0.221 | 0.00018471 | 9 | TAF15 |
| 8.66E-09 | 0.3344002 | 0.493 | 0.304 | 0.00020998 | 9 | MYLIP |
| 8.68E-09 | 0.30727467 | 0.887 | 0.829 | 0.0002104 | 9 | RBM39 |
| 9.23E-09 | 1.38778582 | 0.253 | 0.141 | 0.00022388 | 9 | SERPINF1 |
| 9.85E-09 | 0.41060415 | 0.855 | 0.787 | 0.00023881 | 9 | NPM1 |
| 1.01E-08 | 0.30845677 | 0.303 | 0.166 | 0.00024547 | 9 | ZNF292 |
| 1.12E-08 | 0.31410407 | 0.498 | 0.32 | 0.00027155 | 9 | MACF1 |
| 1.13E-08 | 0.25233662 | 0.317 | 0.174 | 0.00027282 | 9 | PYURF |
| 1.20E-08 | 0.25224241 | 0.525 | 0.335 | 0.00029089 | 9 | NKTR |
| 1.35E-08 | 0.35751086 | 0.95 | 0.912 | 0.00032775 | 9 | PPIA |

| | | | | | | |
|---|---|---|---|---|---|---|
| 1.67E-08 | 0.25543127 | 0.276 | 0.147 | 0.000404 | 9 | POLR3GL |
| 1.68E-08 | 0.25426409 | 0.489 | 0.318 | 0.00040621 | 9 | DYNC1I2 |
| 1.73E-08 | 0.27734622 | 0.448 | 0.285 | 0.00042035 | 9 | SRP9 |
| 1.87E-08 | 0.26843659 | 0.597 | 0.404 | 0.00045324 | 9 | TAF1D |
| 1.87E-08 | 0.37372565 | 0.679 | 0.506 | 0.00045429 | 9 | YME1L1 |
| 1.89E-08 | 0.58499781 | 0.787 | 0.754 | 0.00045854 | 9 | IFITM3 |
| 2.36E-08 | 0.2590999 | 0.765 | 0.588 | 0.00057179 | 9 | SNX3 |
| 2.54E-08 | 0.31461738 | 0.71 | 0.553 | 0.00061512 | 9 | YWHAE |
| 2.74E-08 | 0.42074319 | 0.588 | 0.408 | 0.00066431 | 9 | HIST1H1C |
| 2.97E-08 | 0.40806343 | 0.335 | 0.198 | 0.00072123 | 9 | RNF19A |
| 3.48E-08 | 0.25235332 | 0.425 | 0.262 | 0.00084275 | 9 | ZNF106 |
| 3.98E-08 | 0.47518704 | 0.941 | 0.926 | 0.00096465 | 9 | RPLP0 |
| 4.15E-08 | 0.32805595 | 0.303 | 0.17 | 0.00100553 | 9 | Z99127.4 |
| 4.45E-08 | 0.25881729 | 0.326 | 0.188 | 0.00107858 | 9 | NRBP1 |
| 4.61E-08 | 0.27292067 | 0.416 | 0.266 | 0.00111796 | 9 | USP16 |
| 5.26E-08 | 0.25268245 | 0.394 | 0.244 | 0.00127411 | 9 | PSMD4 |
| 6.20E-08 | 0.34206579 | 0.258 | 0.131 | 0.00150291 | 9 | POLR2B |
| 6.68E-08 | 0.38030555 | 0.995 | 1 | 0.00161938 | 9 | MT-CO3 |
| 6.79E-08 | 0.33792204 | 0.339 | 0.202 | 0.00164605 | 9 | ADI1 |
| 6.91E-08 | 0.26034662 | 0.367 | 0.222 | 0.00167472 | 9 | G3BP1 |
| 7.14E-08 | 0.33753498 | 0.552 | 0.393 | 0.0017317 | 9 | BRD2 |
| 7.29E-08 | 0.28862818 | 0.538 | 0.38 | 0.00176656 | 9 | ANAPC11 |
| 7.43E-08 | 0.26149793 | 0.48 | 0.321 | 0.00180024 | 9 | PA2G4 |
| 7.45E-08 | 0.35874222 | 0.774 | 0.608 | 0.00180541 | 9 | WSB1 |
| 8.32E-08 | 0.30313132 | 0.471 | 0.316 | 0.00201687 | 9 | ARID4B |
| 8.49E-08 | 0.35850029 | 0.995 | 1 | 0.0020577 | 9 | MT-ND4 |
| 8.88E-08 | 0.26200375 | 0.484 | 0.326 | 0.00215364 | 9 | BANF1 |
| 8.92E-08 | 0.26761839 | 0.502 | 0.345 | 0.00216333 | 9 | PSMD8 |
| 9.17E-08 | 0.34692819 | 0.796 | 0.687 | 0.00222282 | 9 | SKP1 |
| 9.27E-08 | 0.38283613 | 0.638 | 0.514 | 0.00224693 | 9 | ITGB1 |
| 9.61E-08 | 0.64178903 | 0.425 | 0.273 | 0.0023293 | 9 | AC091271.1 |
| 1.02E-07 | 0.26762174 | 0.624 | 0.454 | 0.0024717 | 9 | SRSF11 |
| 1.05E-07 | 0.3239669 | 0.742 | 0.613 | 0.00254876 | 9 | MORF4L1 |
| 1.12E-07 | 0.29512149 | 0.43 | 0.281 | 0.00271225 | 9 | SMARCA5 |
| 1.19E-07 | 0.27084262 | 0.548 | 0.39 | 0.0028747 | 9 | CBX3 |
| 1.20E-07 | 0.27951528 | 0.81 | 0.659 | 0.00291032 | 9 | PDIA3 |
| 1.20E-07 | 0.27608835 | 0.534 | 0.378 | 0.00291245 | 9 | EID1 |
| 1.22E-07 | 0.36019952 | 0.416 | 0.27 | 0.00294605 | 9 | PTEN |
| 1.22E-07 | 0.35620335 | 0.262 | 0.15 | 0.00295427 | 9 | RRAS |
| 1.25E-07 | 0.26735475 | 0.452 | 0.301 | 0.00303694 | 9 | TMEM160 |
| 1.30E-07 | 0.4439763 | 0.869 | 0.813 | 0.00314352 | 9 | KLF2 |
| 1.50E-07 | 0.25588577 | 0.258 | 0.139 | 0.0036418 | 9 | AC022868.2 |
| 1.67E-07 | 0.29568373 | 0.656 | 0.497 | 0.00404475 | 9 | NUFIP2 |

| | | | | | | |
|---|---|---|---|---|---|---|
| 1.80E-07 | 0.33633266 | 0.439 | 0.285 | 0.00435292 | 9 | ADM |
| 1.80E-07 | 0.28634772 | 0.584 | 0.432 | 0.00436229 | 9 | SPCS2 |
| 1.85E-07 | 0.25284031 | 0.448 | 0.299 | 0.00449517 | 9 | SMCHD1 |
| 2.24E-07 | 0.34089153 | 0.656 | 0.492 | 0.00544104 | 9 | ARGLU1 |
| 2.58E-07 | 0.33506351 | 0.602 | 0.424 | 0.00625644 | 9 | PPP1R10 |
| 2.80E-07 | 0.25924208 | 0.548 | 0.386 | 0.00678786 | 9 | VPS29 |
| 2.98E-07 | 0.3706248 | 0.873 | 0.771 | 0.00722715 | 9 | HSPE1 |
| 3.06E-07 | 0.25750267 | 0.566 | 0.402 | 0.00741055 | 9 | REEP5 |
| 3.27E-07 | 0.34518911 | 0.48 | 0.332 | 0.0079179 | 9 | DYNC1H1 |
| 3.28E-07 | 0.60914385 | 0.873 | 0.825 | 0.00795555 | 9 | RPSA |
| 3.36E-07 | 0.27103541 | 0.638 | 0.471 | 0.00814324 | 9 | RBM25 |
| 3.56E-07 | 0.29649211 | 0.606 | 0.462 | 0.00862622 | 9 | ST13 |
| 3.82E-07 | 0.26828915 | 0.511 | 0.358 | 0.00925879 | 9 | PRPF40A |
| 3.89E-07 | 0.25701395 | 0.566 | 0.405 | 0.00943857 | 9 | SPCS1 |
| 4.01E-07 | 0.25995439 | 0.425 | 0.28 | 0.00971698 | 9 | DDX18 |
| 4.38E-07 | 0.37113752 | 0.643 | 0.514 | 0.01062362 | 9 | LAPTM4A |
| 5.20E-07 | 0.28289389 | 0.661 | 0.513 | 0.01259505 | 9 | SEC62 |
| 5.73E-07 | 0.41166015 | 0.706 | 0.587 | 0.01390114 | 9 | TPI1 |
| 8.12E-07 | 0.55097822 | 0.765 | 0.653 | 0.01968027 | 9 | EGR1 |
| 1.31E-06 | 0.32786342 | 0.792 | 0.716 | 0.03166133 | 9 | MYL12B |
| 1.37E-06 | 0.28548576 | 0.923 | 0.806 | 0.03325883 | 9 | HSPA1B |
| 1.46E-06 | 0.37190908 | 0.348 | 0.232 | 0.03534433 | 9 | TCEAL4 |
| 1.56E-06 | 0.32340521 | 0.271 | 0.161 | 0.03784327 | 9 | GAS2L3 |
| 1.90E-06 | 0.2535303 | 0.543 | 0.388 | 0.04596208 | 9 | PAIP2 |
| 2.13E-06 | 0.31704577 | 0.977 | 0.974 | 0.05172528 | 9 | RPS20 |
| 2.44E-06 | 0.25029121 | 0.421 | 0.288 | 0.05912376 | 9 | KDELR2 |
| 2.88E-06 | 0.5472344 | 0.498 | 0.35 | 0.06972777 | 9 | HSPA6 |
| 2.96E-06 | 0.2588484 | 0.661 | 0.535 | 0.07171442 | 9 | TMBIM4 |
| 3.00E-06 | 0.26407244 | 0.321 | 0.198 | 0.07262964 | 9 | FAM53C |
| 3.09E-06 | 0.25192234 | 0.652 | 0.508 | 0.0748911 | 9 | SLC25A3 |
| 3.48E-06 | 0.25744491 | 0.281 | 0.169 | 0.08442986 | 9 | GIMAP7 |
| 4.33E-06 | 0.45501139 | 0.692 | 0.568 | 0.10503651 | 9 | IFRD1 |
| 4.74E-06 | 0.29881118 | 0.647 | 0.505 | 0.1149041 | 9 | PARK7 |
| 6.15E-06 | 0.53609141 | 0.643 | 0.557 | 0.14919468 | 9 | TGFBI |
| 6.27E-06 | 0.26482523 | 0.597 | 0.446 | 0.15198373 | 9 | SF1 |
| 6.91E-06 | 0.39177637 | 0.502 | 0.387 | 0.16762072 | 9 | PPDPF |
| 7.22E-06 | 0.25440149 | 0.348 | 0.227 | 0.17504724 | 9 | PGAM1 |
| 7.51E-06 | 0.27806804 | 0.52 | 0.382 | 0.1821343 | 9 | RB1 |
| 8.97E-06 | 0.40320018 | 0.452 | 0.334 | 0.21750618 | 9 | DNAJB11 |
| 1.08E-05 | 0.2722395 | 0.412 | 0.293 | 0.26264898 | 9 | NDUFC1 |
| 1.35E-05 | 0.32428991 | 0.534 | 0.4 | 0.32688006 | 9 | LPAR6 |
| 1.47E-05 | 0.30167544 | 0.367 | 0.248 | 0.35637909 | 9 | THUMPD3-A[illegible] |
| 1.51E-05 | 0.43565144 | 0.955 | 0.948 | 0.36726481 | 9 | ACTG1 |

| | | | | | | |
|---|---|---|---|---|---|---|
| 2.00E-05 | 0.25855154 | 0.48 | 0.365 | 0.48570232 | 9 | KDELR1 |
| 2.08E-05 | 0.25374543 | 0.602 | 0.466 | 0.50425338 | 9 | NOP53 |
| 2.39E-05 | 0.25301166 | 0.778 | 0.659 | 0.57987027 | 9 | PPIB |
| 3.47E-05 | 0.27880003 | 0.9 | 0.845 | 0.84106898 | 9 | TRA2B |
| 3.65E-05 | 0.26298884 | 0.819 | 0.721 | 0.88422074 | 9 | FUS |
| 5.17E-05 | 0.28101294 | 0.602 | 0.469 | 1 | 9 | RSRP1 |
| 6.69E-05 | 0.36855567 | 0.842 | 0.725 | 1 | 9 | HSPD1 |
| 9.93E-05 | 0.32613171 | 0.339 | 0.242 | 1 | 9 | DNAJA4 |
| 0.00011431 | 0.35623456 | 0.756 | 0.667 | 1 | 9 | TSC22D3 |
| 0.00014109 | 0.44299218 | 0.692 | 0.608 | 1 | 9 | CD36 |
| 0.00016723 | 0.3432321 | 0.905 | 0.834 | 1 | 9 | HSPA8 |
| 0.00028386 | 0.26515709 | 0.267 | 0.182 | 1 | 9 | HIST1H2AC |
| 0.00038158 | 0.25090926 | 0.344 | 0.243 | 1 | 9 | SERPING1 |
| 0.00047556 | 0.28500902 | 0.679 | 0.584 | 1 | 9 | ARRDC3 |
| 0.00069396 | 0.37373208 | 0.484 | 0.39 | 1 | 9 | EZR |
| 0.00079828 | 0.41198104 | 1 | 0.99 | 1 | 9 | HSP90AA1 |
| 0.00088661 | 0.26355486 | 0.86 | 0.819 | 1 | 9 | IER5 |
| 0.00093187 | 0.29082021 | 0.801 | 0.7 | 1 | 9 | MYL12A |
| 0.00095244 | 0.4318093 | 0.964 | 0.98 | 1 | 9 | RPL3 |
| 0.00103027 | 0.4544687 | 0.973 | 0.975 | 1 | 9 | RPS3 |
| 0.00118387 | 0.35304326 | 0.824 | 0.758 | 1 | 9 | RPS10 |
| 0.00196969 | 0.27460689 | 0.548 | 0.448 | 1 | 9 | GNG10 |
| 0.00217814 | 0.26406615 | 0.611 | 0.529 | 1 | 9 | TMEM176B |
| 0.00246165 | 0.34444954 | 0.819 | 0.748 | 1 | 9 | MIF |
| 0.00261242 | 0.43755108 | 0.507 | 0.414 | 1 | 9 | IFI6 |
| 0.00283263 | 0.27772162 | 0.991 | 0.999 | 1 | 9 | MT-ND5 |
| 0.00478784 | 0.32270777 | 0.928 | 0.92 | 1 | 9 | IER2 |

51

[illegible]2

[illegible]2

G

2
